# Discovery of the first-efficacious A_2A_R negative allosteric modulators for high adenosine cancer immunotherapies

**DOI:** 10.1101/2024.08.05.606339

**Authors:** Margot Boujut, Margaux Héritier, Aurélie Gouiller, Camille Süess, Alessandro Scapozza, Thibaut De Smedt, Maxime Guibert, Sébastien Tardy, Hesham Hamed, David Pejoski, Leonardo Scapozza

**Affiliations:** School of Pharmaceutical Sciences, University of Geneva, Geneva, Switzerland; Institute of Pharmaceutical Sciences of Western Switzerland, University of Geneva, Geneva, Switzerland; Adoram Therapeutics, Grand-Lancy, Switzerland

**Author notes:** Correspondence should be addressed to Prof. Leonardo Scapozza. These authors contributed equally to this work (Margot Boujut, Margaux Héritier). xSeedD Sàrl, Geneva, Switzerland (Thibaut De Smedt).

**Keywords:** Hit to lead discovery, allosteric modulator, negative allosteric modulator NAM, High throughput screening, A2AR, high adenosine cancer, immunotherapy

## Abstract

Inhibition of adenosine 2A receptor (A2AR) is recognized as a promising immunotherapeutic strategy but is challenged by the ubiquity of A2AR function in the immune system. To develop a safe yet efficacious immunotherapy, the discovery of a novel negative allosteric modulator (NAM) was preferred. Leveraging an in-house, sensitive, high-throughput screening cellular assay, novel A2AR NAM scaffolds were identified followed by an extensive structure-activity relationship (SAR) study, leading to the discovery of potent 2-amino-3,5-dicyanopyridine derivatives. Allosteric mode of action of active compounds was confirmed by shift assay, non-linearity of the Schild plot analysis, biophysical measurements, and retained satisfactory potencies in high-adenosine concentrations. Further correlation of A2AR engagement and downstream signaling was done in a human blood translational assay, clearly showcasing the potential of A2AR allosteric modulation as a novel approach for efficient and safer cancer immunotherapies.

## Introduction

For almost two decades now, the modulation of adenosine receptors has been recognized as a promising therapeutic strategy for many diseases and disorders including, but not limited to, neurological, cardiovascular, renal, intestinal, inflammatory, and pulmonary conditions.^1–4^ This especially broad list is due to the many regulatory mechanisms involving adenosine and the adenosine receptors throughout the body. The adenosine receptor family is part of the G-protein coupled receptor superfamily and is composed of four receptor subtypes: A_1_R and A_3_R, which are G_i_-coupled receptors, and on the other hand A_2A_R and A_2B_R, which are G_s_-coupled receptors.^5^ The adenosine 2A receptor (A_2A_R) for example is found both in the central nervous system and in the periphery, mainly in the immune system (spleen, thymus, leucocytes, and blood platelets) and at intermediate levels in the heart, blood vessels and lungs.^6^ This ubiquity of the adenosine receptors poses a challenge to the development of safe and potent therapies.^1,3^

Adenosine exerts a cytoprotective effect by engaging A_2A_R in the immune system, resulting in an anti-inflammatory and pro-resolving response.^7^ However, accumulation of extracellular adenosine is one of the widespread immunosuppressive mechanisms by which anti-tumor immunity is gained. High adenosine in the tumor microenvironment (TME) indeed correlates with tumor aggressiveness.^8^ It was of interest to discover and optimize safe ligands able to outcompete high adenosine concentrations.

The first A_2A_R orthosteric ligands, such as Preladenant, Imaradenant, Taminadenant or Ciforadenant (Figure 1), were developed for, or derived from, treatments of Parkinson disease where adenosine concentrations are lower than in the TME.^9^ When repurposed as cancer immunotherapies, it resulted in low response rates and/or safety concerns due to high dosing and discontinuation of most clinical trials. Exciting progress was made in the more recent development of these cancer immunotherapies by combination of immune checkpoint inhibitors and A_2A_R antagonists developed to withstand high adenosine concentrations, such as Etrumadenant and Inupedenant (Figure 1).^10^

**Figure 1.**
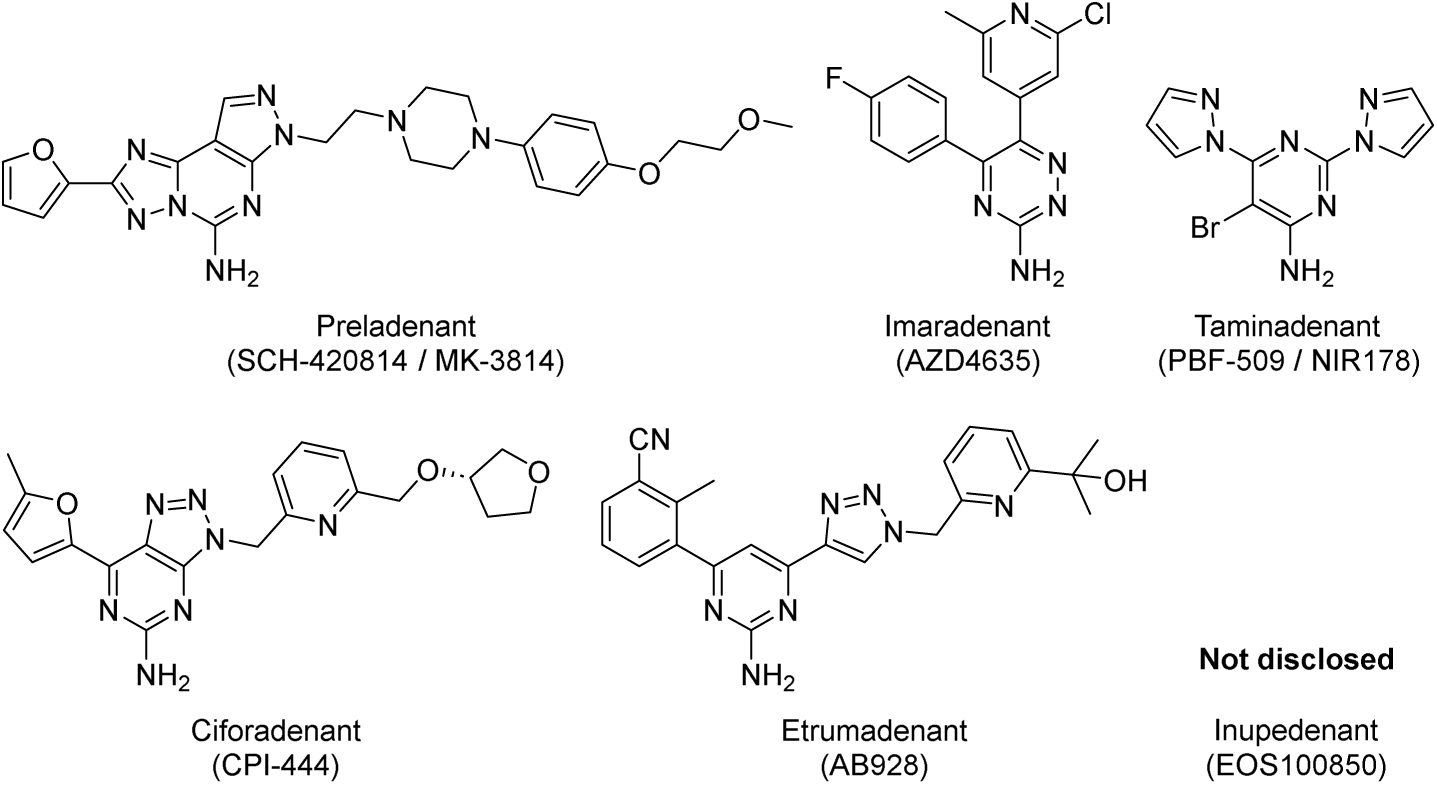
Clinical candidates for cancer immunotherapies by A2AR orthosteric.

One important consideration in the clinical use of A_2A_R antagonists is the safety of the treatments due to the ubiquity of adenosine signaling in physiological processes.^4,7^ Negative allosteric modulators (NAMs) represent an attractive type of ligand to overcome this challenge. NAMs are characterized by their allosteric probe-dependent and saturable mode of action. Negative allosteric modulation is characterized by decreasing of the affinity and/or the efficacy of orthosteric ligands, including the natural ligand(s) of a receptor, which a saturable effect once all allosteric sites are occupied. This saturable effect of NAMs allows to work at the minimum efficacious dosing of a treatment, as increased dosage doesn’t result in increased therapeutic effect, limiting the risk of off-target side-effects.^11,12^ Off-target side effects are also expected to be lowered in the case of allosteric ligands as allosteric pockets are postulated to be less conserved due to their remote positioning with respect to the orthosteric binding site, hence it should be possible to achieve higher receptor selectivity.^12,13^

Despite the now large body of literature surrounding A_2A_R protein structure^14^ (76 structures available in the RCSB protein data bank as of October 4^th^, 2023), only the sodium binding site has been well identified and characterized as an allosteric pocket.^15,16^ Different approaches have been employed to try and propose allosteric sites for A_2A_R^17^ but, to this day, experimental data are lacking to validate these hypothesis. This lack of rational might explain the very small number of known A_2A_R allosteric modulators in comparison to the intense development of its orthosteric ligands. Few positive allosteric modulators have been reported to improve CGS21680 (a specific agonist) A_2A_-mediated pharmacological effects in *ex vivo* experiments,^18^ attenuate inflammation in pre-clinical models of psoriasis-like dermatitis^19,20^ or to be able to induce slow-sleep wave in mania and schizophrenia-like behaving mice.^21,22^ Advances on A_2A_R negative allosteric modulators are more scarce in the literature (Supporting Information Figure S1); amiloride and its derivatives (non-selective A_2A_R binders) were shown to accelerate the dissociation rate of radio-labelled ZM241385, most likely favoring a closed, inactive conformation of A_2A_R. This binding was outcompeted by sodium addition in a dose-dependent manner, hinting that amiloride and sodium act through the same allosteric pocket.^23^ Fragments acting as A_2A_R NAMs, such as Fr754, have been described more recently by Lu *et al.* using a novel affinity mass spectrometry technic and suggesting a different binding pocket.^24^ However, no further developments have been published.

In this work, our in-house dynamic cAMP measurement assay (CamBio assay) was used in a high-throughput screening campaign to discover A_2A_R antagonist scaffolds and novel A_2A_R NAMs. To distinguish between orthosteric inhibitors (and inverse agonists) and fully allosteric hits, these scaffolds were further characterized by shift assays.^12^ This method resulted in the discovery of compound **1**, which demonstrated a desirable NAM profile. We presently describe the first extensive structure-activity relationship (SAR) campaign for A_2A_R allosteric modulators following the discovery of compound **1**, which led to the discovery of potent and A_2A_R-selective 2-amino-3,5-dicyanopyridine derivatives (Figure 2). Rigorous experimentation receptor subtype selectivity. A proof-of-concept of compound activity could be observed in human translational assays, clearly showcasing the potential of these compounds as novel immunotherapies for high adenosine cancers.

**Figure 2.**
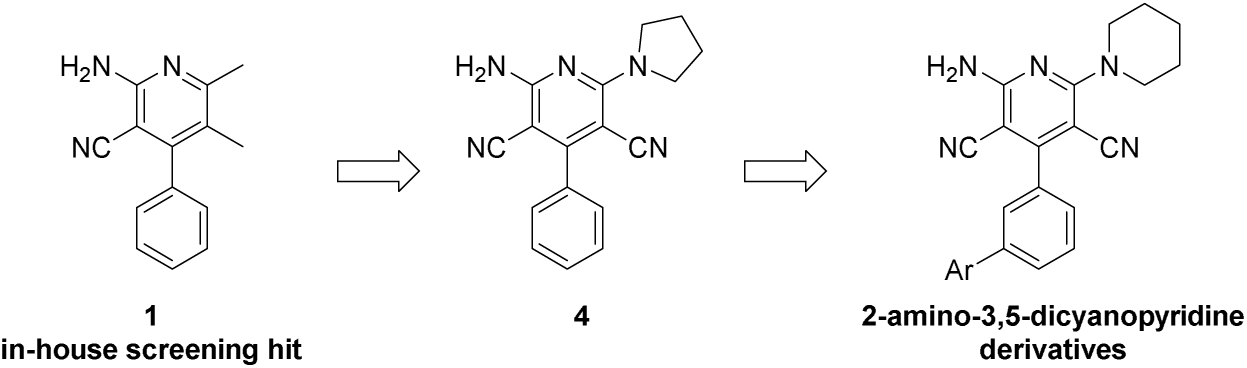
From hit compound 1 to 2-amino-3,4-dicyanopyiridine derivatives.

## Results & discussion

### Synthetic routes

Compounds **4**-**10** were first synthetized *via* a 2-step process^25^ (Scheme 1). Trimethyl orthobenzoate and two equivalents of malononitrile were first heated in pyridine then subjected to strong acidic conditions to give compound **52**. This chloropyridine intermediate **52** can be substituted in a short amount of time by an excess of primary or secondary amine under microwave irradiation to yield the desired compounds **4**-**10**.

**Scheme 1.**
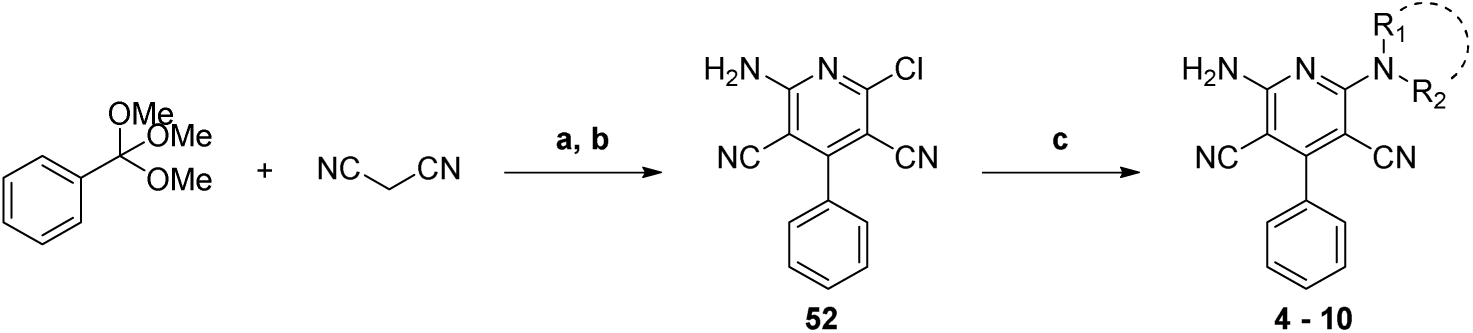
Synthesis of compounds 4-10.

The general one-pot cyclization used for the obtention of compounds **11**-**30** and **32**-**37** is presented Scheme 2; compounds **38**-**43** presented Scheme were also obtained *via* this procedure. This protocol has been adapted from Sarkar *et al.*^26^ with moderate success, however, it happened to be quite tolerant with various chemical function and highly reproducible.

**Scheme 2.**
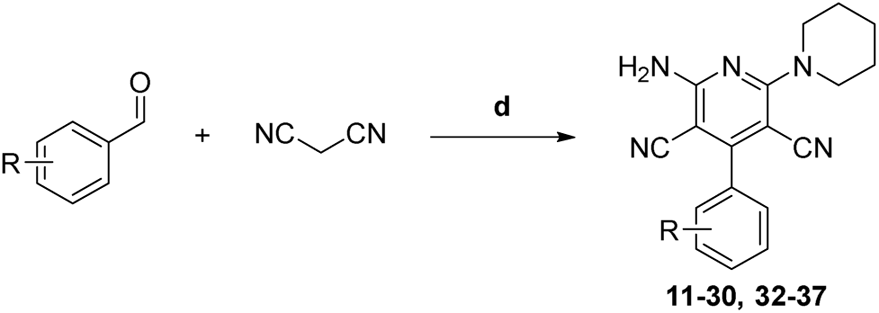
Synthesis of compounds 11-30 and 32-37. d) piperidine, 4-DMAP (cat.) in MeOH from 0°C to r.t., open to air, for 16 h, 7%-30% isolated yields.

Starting from 3-bromobenzaldehyde, various bis aryl derivatives could be isolated after a Suzuki-Miyaura cross-coupling reaction in presence of substituted phenylboronic acids (Scheme 3). These intermediate aldehydes **53**-**58** could be subjected to the general procedure presented above to obtain compounds **38**-**43** with moderate to good yields. Alternatively, 3-bromobenzaldehyde can directly be cyclized to the compound **27** and used as synthetic intermediate

**Scheme 3.**
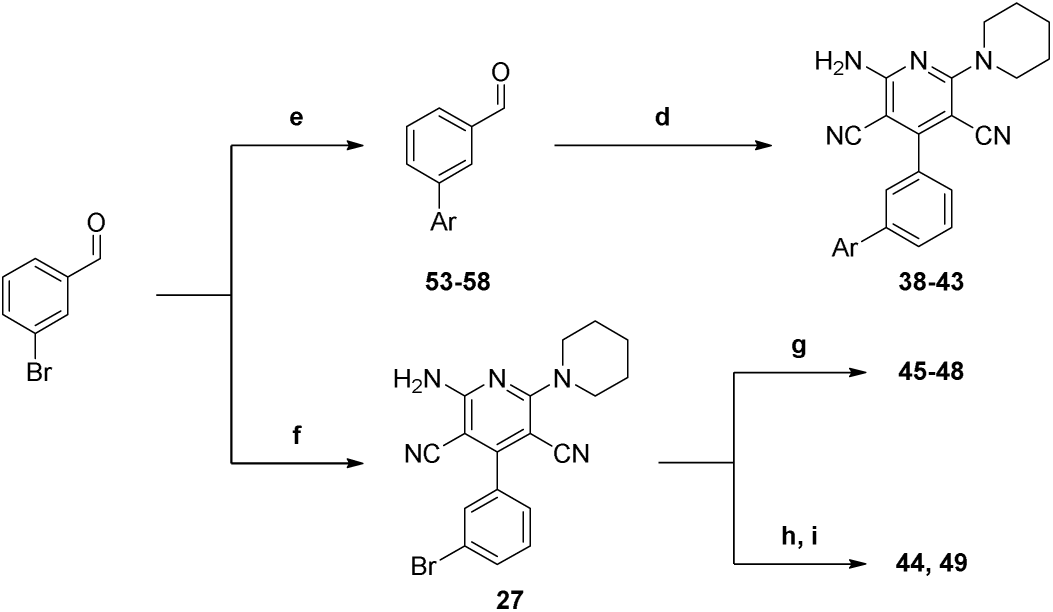
**Synthesis of compounds 38-49.**

A gram-scale protocol can be achieved in a semi-sequential manner, in absence of the catalytic 4-DMAP and by using a slight excess of reactive piperidine instead. The compound **27** was used as a starting aryl bromide in Suzuki-Miyaura cross-coupling activated by microwave irradiation to access other desired compounds **45**-**48**. For more reluctant substrates or non-commercial boronic acids, the boronic pinacol ester could be directly installed on compound **27** under traditional heating and reacted with commercial bromoaryls in similar microwave-activated fashion to yield compounds **44** and **49** (Scheme 3).

The intermediate **27** could also be used in other types of metal cross-couplings like Sonogashira copper-palladium coupling to obtain compound **31** after deprotection of the triisopropylsilyl terminal alkyne with a TBAF solution. Other late-stage intermediates could be employed for further functionalization like compound **25** which was used for the cyclization of an oxadiazole in two steps for the synthesis of compound **50** (Scheme 4).

**Scheme 4.**
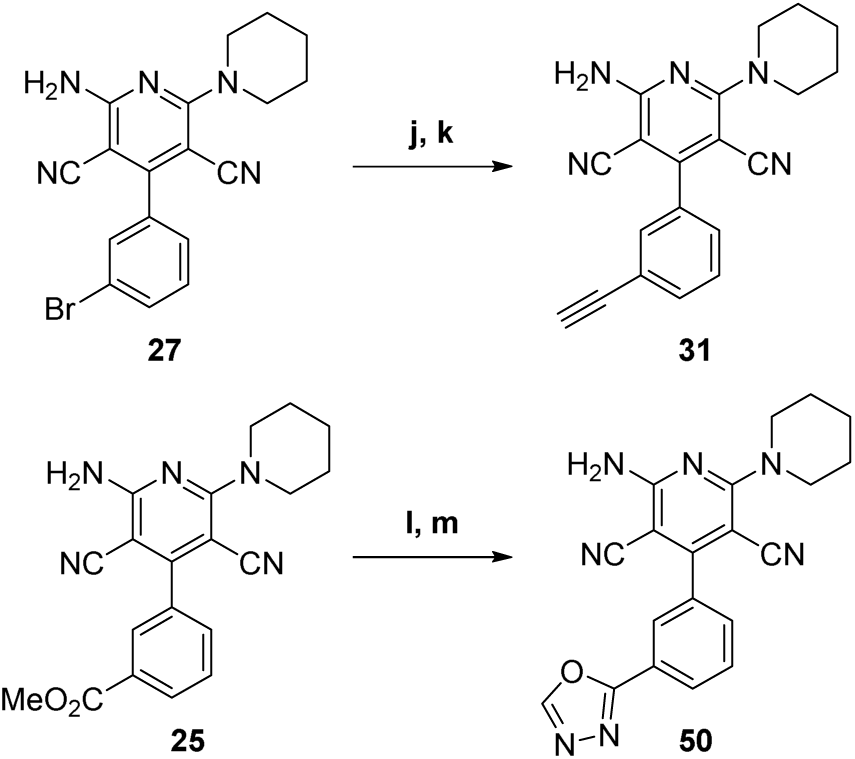
**Synthesis of compounds 31 and 50.**

Finally, some protection/deprotection strategies had to be applied for more reactive derivatives such as the tetrazole **51** which was first built from the 3-formylbenzonitrile to the aldehyde **59** (Scheme 5). After a THP protection, the aldehyde **60** could be further cyclized to the 2-amino-3,5-dicyanopyridine core and deprotected by treatment with a slightly acidic resin to the desired compound **51** (Scheme 5).

**Scheme 5.**
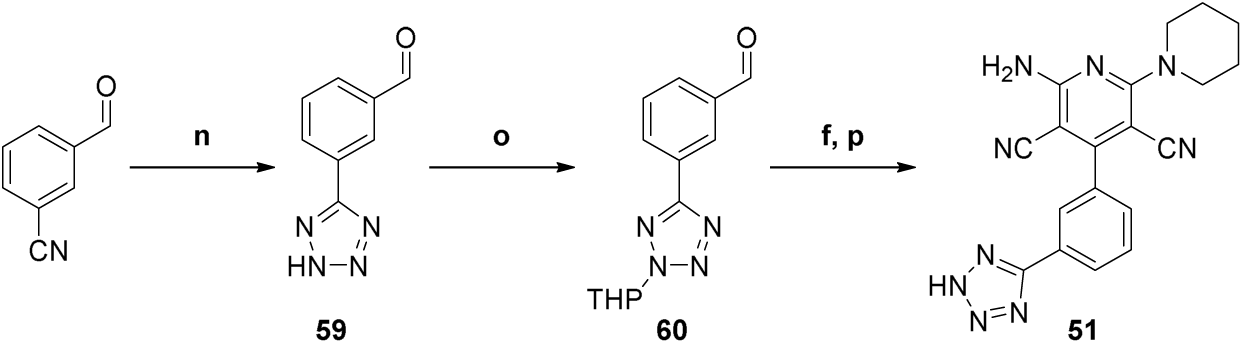
**Synthesis of compounds 51.**

### High throughput screening assay identified novel NAM scaffolds

Though A_2A_R is a well-studied target, the ligands described in the literature are mostly orthosteric inhibitors. To free ourselves from this orthosteric mode of action, novel hits had to be discovered. The strategy of biological assay development followed by an in house high-throughput screening campaign was in line with this need for original scaffolds.

In short, our cAMP measuring (“CamBio”) assay is a high-throughput screening assay, selective, sensitive, and reproducible, developed for allosteric modulators detection. The assay allows the direct, real-time monitoring of intracellular cAMP concentration thanks to a BRET-based biosensor containing a cAMP binding domain co-expressed with A_2A_R, or any GPCR, in CHO cells. Moreover, its BRET-based biosensor enables a dynamic measurement of both G_s_- and G_i_-coupled receptors activation without artificially modifying their coupling mechanism, which is key for allosteric modulator identification. High-throughput screening capabilities are achieved by monitoring using a FDSS/µCELL platform.

For the hit identification campaign, an in house, manually curated library of 2’440 compounds (representative of 6 million of commercially available scaffolds spanning over a large spectrum of biologically relevant targets)^27^ was screened. Testing compounds using a dose of adenosine corresponding to the concentration at which 80% of the receptor is activated (EC_80_, [adenosine] = 400 nM) allowed us to identify novel A_2A_R-antagonising scaffolds (data not shown). Their mode of action was tested on the same cell line for the distinction between orthosteric inhibitors and allosteric modulators by highlighting the non-competitive binding mode of the latter; this is materialized by a stacking of the dose response curves of adenosine in presence of various compound’s concentrations in a shift assay (more details in the following sections). Only the resulting NAMs were labelled as hits.

Compound **1** demonstrated a promising IC_50_ of 0.55 µM even though it was exhibiting only partial inhibition of the receptor (47% at 10 µM of compound; Figure 3). Whether this partial inhibition of the system was due to the compound’s low activity or inherent to its allosteric mode of action could only be solved later in the medicinal chemistry campaign. Compound’s **1** shift assay demonstrated the expected curve stacking pattern for NAMs, thus it was labelled as one of the initial hits.

**Figure 3.**
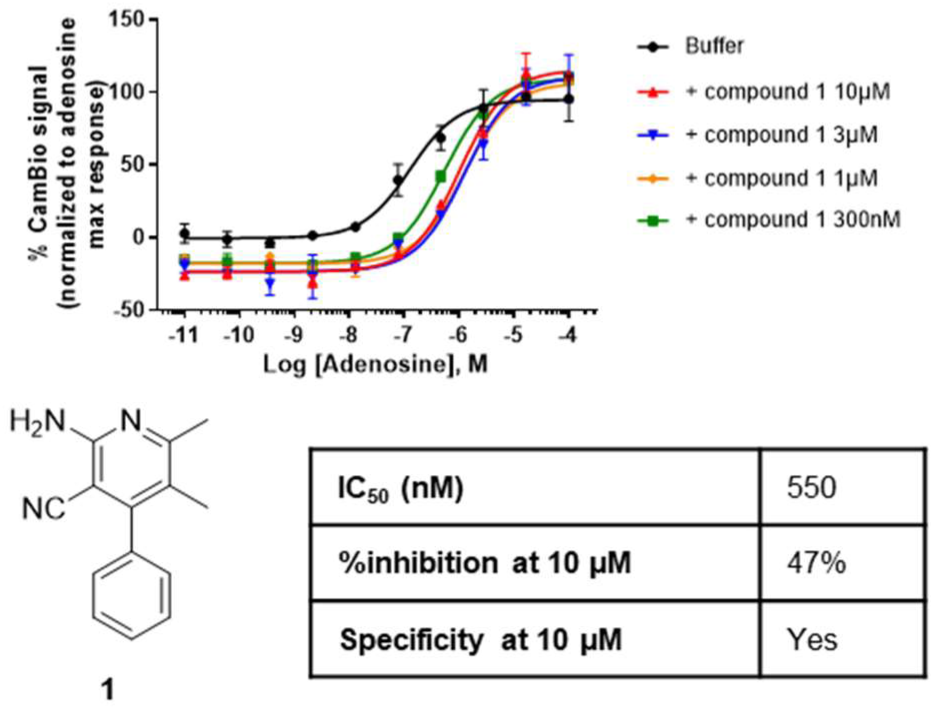
Structure, activity, and confirmation of the allosteric profile by shift assay of hit compound **1**; ^1^ IC_50_ (nM) in presence of 400 nM of adenosine and % inhibition at 10 µM values are given by CamBio assay on CHO cells expressing A_2A_R, experimental repetitions N = 2-5; ^2^ Specificity is given by CamBio assay on the parental CHO cell, experimental repetitions N = 2-3.

### SAR exploration

Using compound **1** as our starting point, a first analoging step was performed thanks to commercially available libraries. Biological effect on A_2A_R was first evaluated using our CamBio assay in presence of an EC_80_ of adenosine and reported in Figure 4. All compounds demonstrated moderately good activity with IC_50_ in the 10^-7^–10^-6^ M range and similar percentage of inhibition at 10 µM of compound (%inh. @10 µM). Formation of a bicyclic system in compound **2** resulted in a 3-fold loss of potency. While two modifications were introduced in compound **3** and direct comparison cannot be made, it demonstrated that both aromatic substitution and larger substituents on the position 6 of the pyridine could be tolerated to a certain extent. Still, compound **3** overly aromatic nature prevented it from being a good starting block and finally, compound **4** was retained for its desirable properties, its maintained allosteric profile (see Supporting Information) and the tractability of its chemistry.

**Figure 4.**
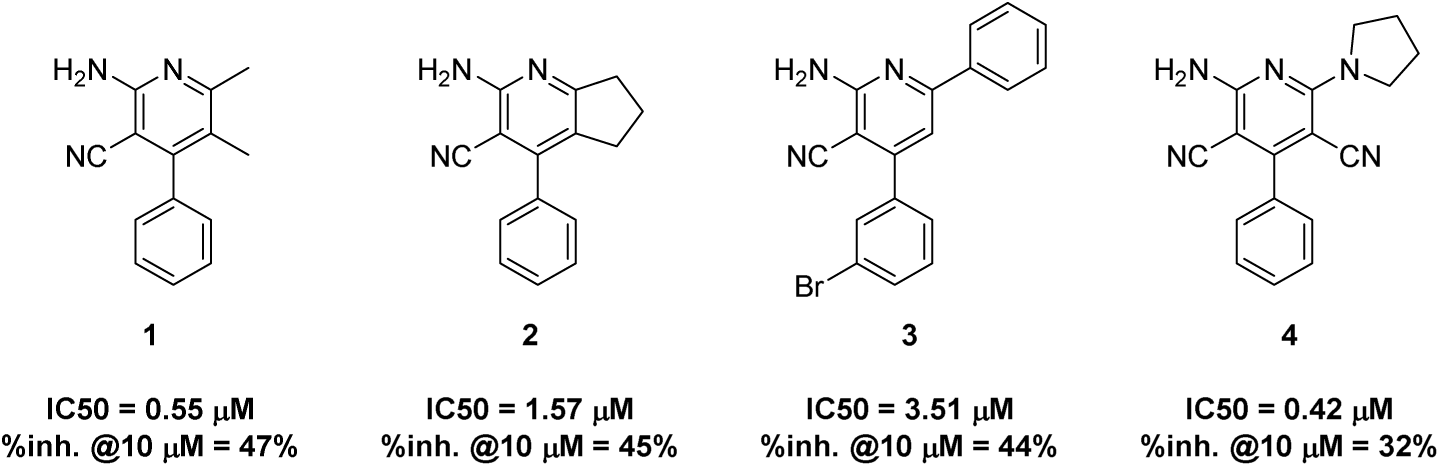
Structure and biological effects on A_2A_R of compounds **1-4** resulting from a rapid scaffold hoping from commercially available analogs.

As it was the largest point of modification between the initial scaffolds, the influence of the cyclic amine substitution was first explored. The nature of the amine, the ring size and ring substituents were varied (Table 1). Going from a 5- to a 6-membered ring proved beneficial with a 2-fold improvement of the IC_50_ (**4**, IC_50_ = 420 nM; **5**, IC_50_ = 220 nM; Table 1) and a satisfactory increase of the % inhibition (**4**, %inh. @10 µM = 32%; **5**, %inh. @10 µM = 70%). Furthermore, ring enlargement only resulted in a drop of activity for compound **6**, most likely due to the overly increased steric hinderance. Similar observations can be made if the nature of the disubstituted amine goes from a restricted cyclic structure to flexible branched amines (**5** vs. **7** and **8**). Hence, the cyclic restriction was kept and substitutions of the piperidine moiety were tested to try increasing either its hydrophobicity or its hydrophilicity. This only resulted in a loss in A_2A_R activity, 12 and 6-fold for compounds **9** and **10** respectively. This further highlighted the likely limited space and hydrophobic nature of the pocket accommodating this part of the molecule. Interestingly, most of the tested scaffolds presented a good selectivity profile compared to the closest isoform A_2B_R; additionally, compound **5** activity on A_3_R was judged as a good starting point while A_1_R selectivity remained to be improved.

**Table 1.** Biological effects of compounds 4-10.

| Compound | R | IC<sub>50</sub> (nM)<sup>1</sup> | %inhibition<br>@10 μM<sup>1</sup> | Specificity<br>@10 μM<sup>2</sup> | Fold select.<br>VS A1R<sup>3</sup> | Fold select.<br>VS A2BR<sup>3</sup> | Fold select.<br>VS A3R<sup>3</sup> |
| --- | --- | --- | --- | --- | --- | --- | --- |
| 4 |  | 420 | 32% | Yes | n.d. | good | n.d. |
| 5 |  | 220 | 70% | Yes | poor | good | agreeable |
| 6 |  | no sat. | 22% | Yes | n.d. | good | n.d. |
| 7 |  | n.a. | n.a. | n.a. | n.d. | n.a. | n.d. |
| 8 |  | no sat. | 24% | Yes | n.d. | good | n.d. |
| 9 |  | 2300 | 50% | Yes | n.d. | good | n.d. |
| 10 |  | 1210 | 54% | Yes | n.d. | good | n.d. |

**n.a.** = non active; **no sat.** = no saturation of the dose response curve was reached, IC<sub>50</sub> cannot be calculated; **n.d.** = not determined.
<sup>1</sup> IC<sub>50</sub> (nM) in presence of 400 nM of adenosine and % inhibition at 10 μM values are given by CamBio assay on CHO cells expressing A<sub>2A</sub>R, experimental repetitions N = 2-5.
<sup>2</sup> Specificity is given by CamBio assay on the parental CHO cell, experimental repetitions N = 2-3.
<sup>3</sup> Fold selectivity is given as a ratio of the IC<sub>50</sub> measured by CamBio assay on CHO cells expressing the corresponding receptor to the IC<sub>50</sub> measured by CamBio assay on CHO cells expressing A<sub>2A</sub>R, experimental repetitions N = 2-3; ratio: 0 ≤ none < 2, 2 ≤ poor < 5, 5 ≤ agreeable < 15, good > 15.

As demonstrated by the early explorations, substituents seemed to be tolerated around the phenyl in position 4 of the pyridine core and called for further inquiries (compound **3,** Figure 4). Still, direct comparison of compounds **3** and **27** was eventually possible and reenforced our choice of the current core as a better and more potent structure (IC_50_ = 3.51 µM and 1.04 µM, respectively). All biological effects for mono- and di-substitution patterns are recapitulated Table 2 and Table 3, respectively.

**Table 2.**
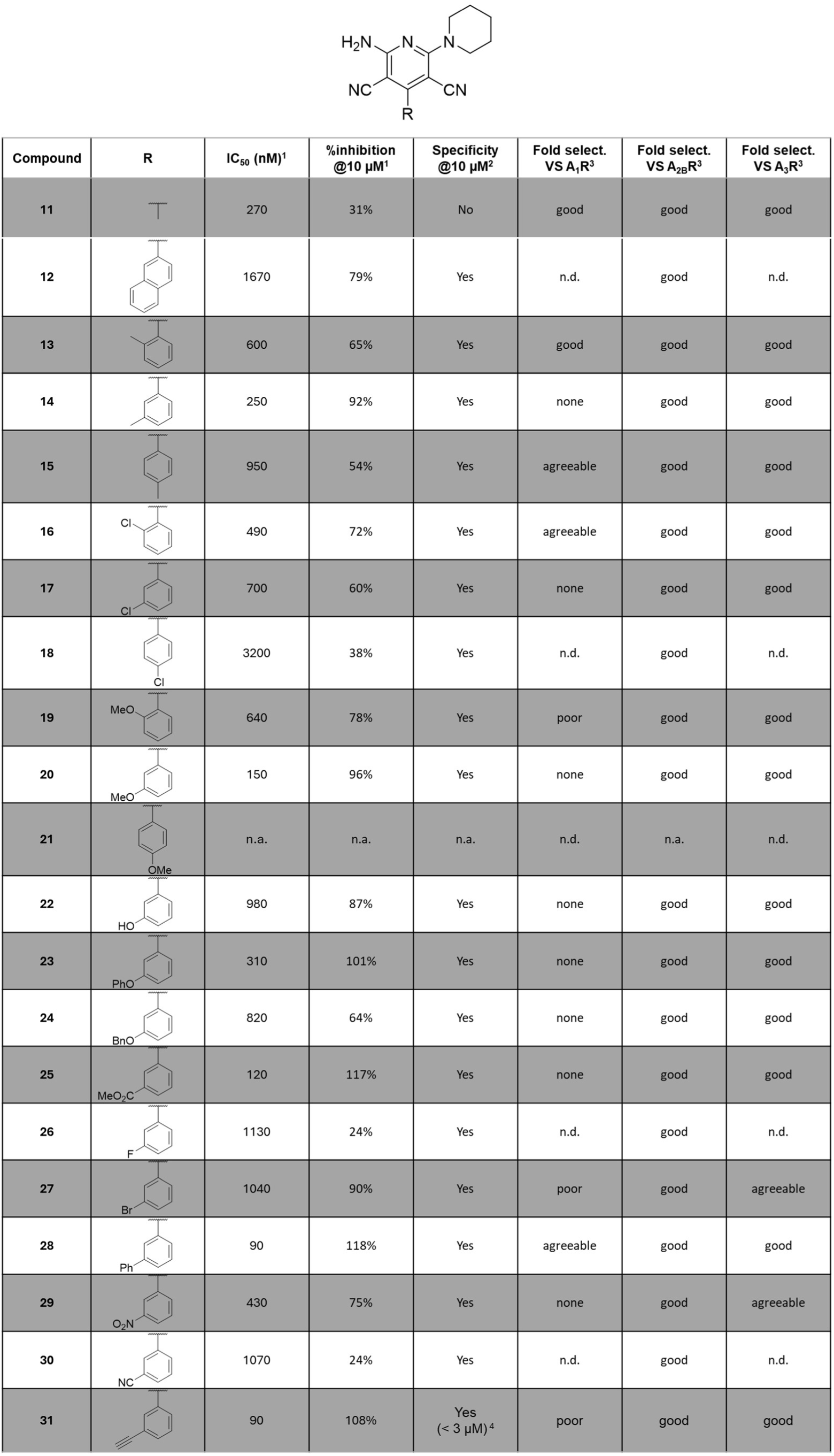
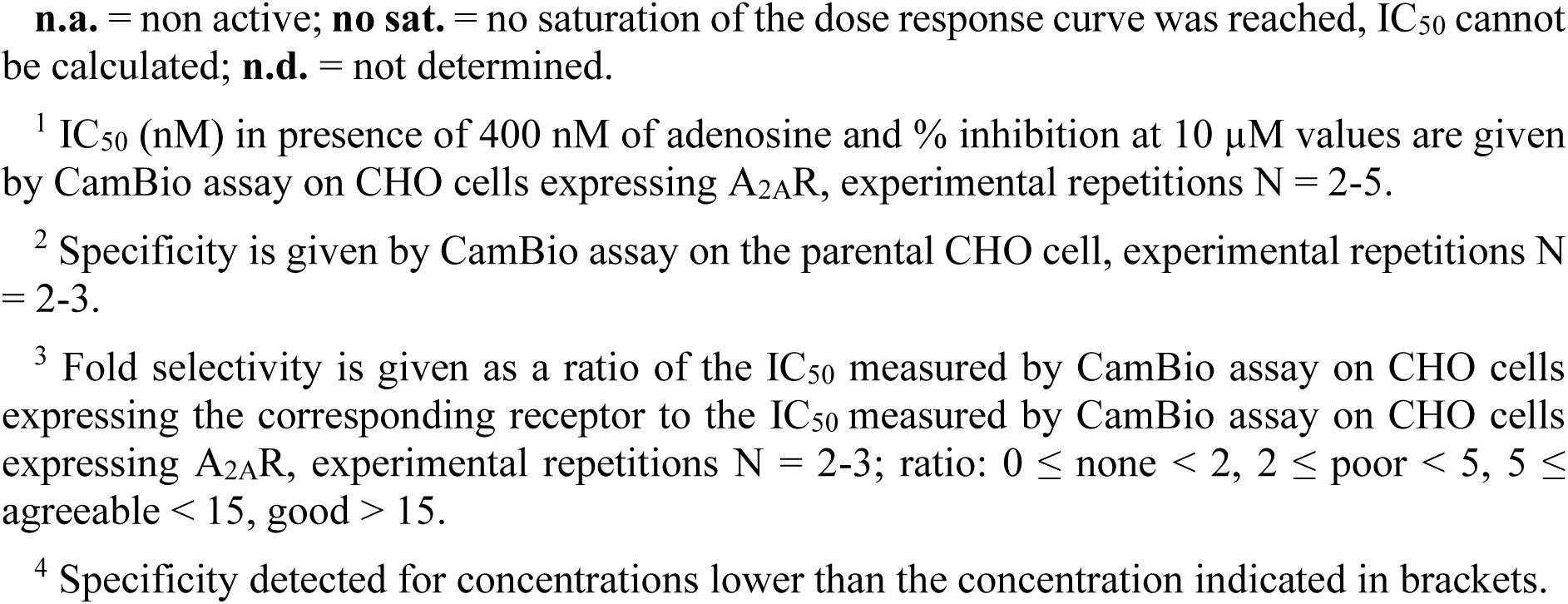
Biological effects of compounds 11-31.

| Compound | R | IC<sub>50</sub> (nM)<sup>1</sup> | %inhibition<br>@10 μM<sup>1</sup> | Specificity<br>@10 μM<sup>2</sup> | Fold select.<br>VS A<sub>1</sub>R<sup>3</sup> | Fold select.<br>VS A<sub>2B</sub>R<sup>3</sup> | Fold select.<br>VS A<sub>3</sub>R<sup>3</sup> |
| --- | --- | --- | --- | --- | --- | --- | --- |
| 11 |  | 270 | 31% | No | good | good | good |
| 12 |  | 1670 | 79% | Yes | n.d. | good | n.d. |
| 13 |  | 600 | 65% | Yes | good | good | good |
| 14 |  | 250 | 92% | Yes | none | good | good |
| 15 |  | 950 | 54% | Yes | agreeable | good | good |
| 16 |  | 490 | 72% | Yes | agreeable | good | good |
| 17 |  | 700 | 60% | Yes | none | good | good |
| 18 |  | 3200 | 38% | Yes | n.d. | good | n.d. |
| 19 |  | 640 | 78% | Yes | poor | good | good |
| 20 |  | 150 | 96% | Yes | none | good | good |
| 21 |  | n.a. | n.a. | n.a. | n.d. | n.a. | n.d. |
| 22 |  | 980 | 87% | Yes | none | good | good |
| 23 |  | 310 | 101% | Yes | none | good | good |
| 24 |  | 820 | 64% | Yes | none | good | good |
| 25 |  | 120 | 117% | Yes | none | good | good |
| 26 |  | 1130 | 24% | Yes | n.d. | good | n.d. |
| 27 |  | 1040 | 90% | Yes | poor | good | agreeable |
| 28 |  | 90 | 118% | Yes | agreeable | good | good |
| 29 |  | 430 | 75% | Yes | none | good | agreeable |
| 30 |  | 1070 | 24% | Yes | n.d. | good | n.d. |
| 31 | | 90 | 108% | Yes<br>($< 3 \mu\text{M}$)<sup>4</sup> | poor | good | good |

**n.a.** = non active; **no sat.** = no saturation of the dose response curve was reached, IC<sub>50</sub> cannot be calculated; **n.d.** = not determined.
<sup>1</sup> IC<sub>50</sub> (nM) in presence of 400 nM of adenosine and % inhibition at 10 μM values are given by CamBio assay on CHO cells expressing A<sub>2A</sub>R, experimental repetitions N = 2-5.
<sup>2</sup> Specificity is given by CamBio assay on the parental CHO cell, experimental repetitions N = 2-3.
<sup>3</sup> Fold selectivity is given as a ratio of the IC<sub>50</sub> measured by CamBio assay on CHO cells expressing the corresponding receptor to the IC<sub>50</sub> measured by CamBio assay on CHO cells expressing A<sub>2A</sub>R, experimental repetitions N = 2-3; ratio: 0 ≤ none < 2, 2 ≤ poor < 5, 5 ≤ agreeable < 15, good > 15.
<sup>4</sup> Specificity detected for concentrations lower than the concentration indicated in brackets.

**Table 3.** Biological effects of mono- (compounds 13, 14 and 17) and di-substituted compounds 32-37.

| Compound | R<sub>1</sub> | R<sub>2</sub> | IC<sub>50</sub> (nM)<sup>1</sup> | %inh@10 μM<sup>1</sup> | Compound | R<sub>1</sub> | R<sub>2</sub> | IC<sub>50</sub> (nM)<sup>1</sup> | %inh@10 μM<sup>1</sup> |
| --- | --- | --- | --- | --- | --- | --- | --- | --- | --- |
| 13 | 2-Me | H | 600 | 65% | 34 | 3-Cl | 2-F | 343 | 83% |
| 14 | 3-Me | H | 250 | 92% | 35 | 3-Cl | 4-F | 599 | 90% |
| 17 | 3-Cl | H | 700 | 60% | 36 | 3-Cl | 5-F | 381 | 89% |
| 32 | 2-Me | 6-Me | n.a. | n.a. | 37 | 5-Cl | 2-F | 207 | 81% |
| 33 | 3-Me | 5-Me | 7280 | 52% |  |  |  |  |  |

<sup>1</sup> IC<sub>50</sub> (nM) in presence of 400 nM of adenosine and % inhibition at 10 μM values are given by CamBio assay on CHO cells expressing A<sub>2A</sub>R, experimental repetitions N = 2-5.

As the biological effect of the aromatic depletion in compound **11** was comparable to the parent hit compound **4** but resulted in a loss of the specific activity towards A_2A_R, the aromatic core in position 4 was maintained (Table 2). Larger groups such as naphthyl (compound **12**) and para substituents (compounds **15**, **18** and **21**) resulted in dropsof 10-fold or more in IC_50_, up to a complete loss of activity (compound **21**). Ortho substitution of the ring was well tolerated and even favored for moderately electron-withdrawing atoms such as chlorine (compound **16**), but meta substitutions yielded the best results, achieving almost full inhibition of the system with the lowest IC_50_ when compared in the same series (compounds **14** vs **13** and **15**, **20** vs **19** and **21**). For this reason, the following efforts were made on the nature of substitutions in the meta position. Medium to strong electron withdrawing groups tended to reduce to both the IC_50_ and maximum potency of the compounds (compounds **26**, **27** and **30**), but this effect appeared to be countered by the presence of hydrogen bond acceptors (compounds **25** and **29**). Surprisingly, larger and more rigid substituents were tolerated for this meta exit vector, with improved IC_50_ in the low nanomolar range and the desired full inhibition of A_2A_R activity (**28**, IC_50_ = 90 nM and %inh. @10 µM = 118%; **31**, IC_50_ = 90 nM and %inh. @10 µM = 108%), resulting in the characterization of the first optimized A_2A_R NAMs.

Most of these compounds were also tested for specific and selective activity and, while A_2B_R and A_3_R selectivity profile remained favorable, meta substituents presented only little selectivity until larger phenyl group was introduced on compound **28**.

Having improved receptor subtype selectivity and encouraged by the current research surrounding A_2A_R orthosteric ligands, we believed that better potencies could be achieved with our NAMs and decided to investigate further this versatile aromatic substitution. Biological activities of some di-substitution patterns are presented Table 3.

Dimethyl substitution patterns of compounds **32** and **33** turned out to be largely disfavored compared to the monosubstituted analog (**32** and **33** vs **13** and **14**), hence the focus shifted to the introduction of single atom differences. As often encountered in the medicinal chemistry campaign,^28^ the change of a hydrogen to a fluorine, or “fluorine walk”, was not benign and influenced both the IC_50_ and % inhibition of the synthetic precursor **17**, with variable results depending on the fluorine position. Interestingly, the presence of an additional fluorine atom positively influenced the compounds’ activities, with IC_50_ cut in half compared to parent compound **17** for all except compound **35** (**34**, IC_50_ = 343 nM, **36**, IC_50_ = 381 nM, **37**, IC_50_ = 207 nM vs. **17**, IC_50_ = 700 nM) and % inhibition at 10 µM increased by 1.5-fold, up to almost full inhibition of the system. These results could be combined with the optimized mono-substitution for the future developments of the lead compounds.

All the previous results combined suggested the existence of a hydrophobic groove/pocket for which the meta position of a phenyl ring creates an ideal orientation. Bisphenyl substitution appeared to be quite potent and offered a solid platform to start exploring further this new space (Table 4). Small substituents of hydrophobic nature, either electron withdrawing or donating, were well accommodated both in ortho and meta (compounds **38**, **39**, **41** and **42**; IC_50_ = 160-290 nM and above full % inhibition), however, para substituents had more than 10-fold losses in IC_50_ (compounds **40** and **43**). Introduction of soft sulfur atoms or various harder heteroatoms in five-member rings did not strongly influence the inhibitory profile either (compounds **47**-**51**). The use of a hydrogen bond donor did not bring anything more to the system (compound **49**) and was even unfavored in case of a strongly acidic proton (tetrazole in compound **51**). On the other hand, when the hydrogen bond donor capabilities of the protein were explored by a simple nitrogen ring road, ortho and meta pyridines were largely favored (**44** and **45**; respectively, 6-fold and 10-fold IC_50_ improvements compared to compound **28**), yielding compounds with IC_50_ in the targeted low nanomolar range. The allosteric mode-of-action of some selected compounds was confirmed once more by shift assay, assuring that the whole family was indeed maintaining its allosteric characteristics (see Supporting Information). Addition of this bisphenyl substitutions conferred a good selectivity profile over A_2B_R and A_3_R overall while A_1_R selectivity profile appeared more unpredictable. Still, the activity and selectivity profiles on cells of compounds **28** and **44-48** were recognized as good and they were pushed for further testing.

**Table 4.**
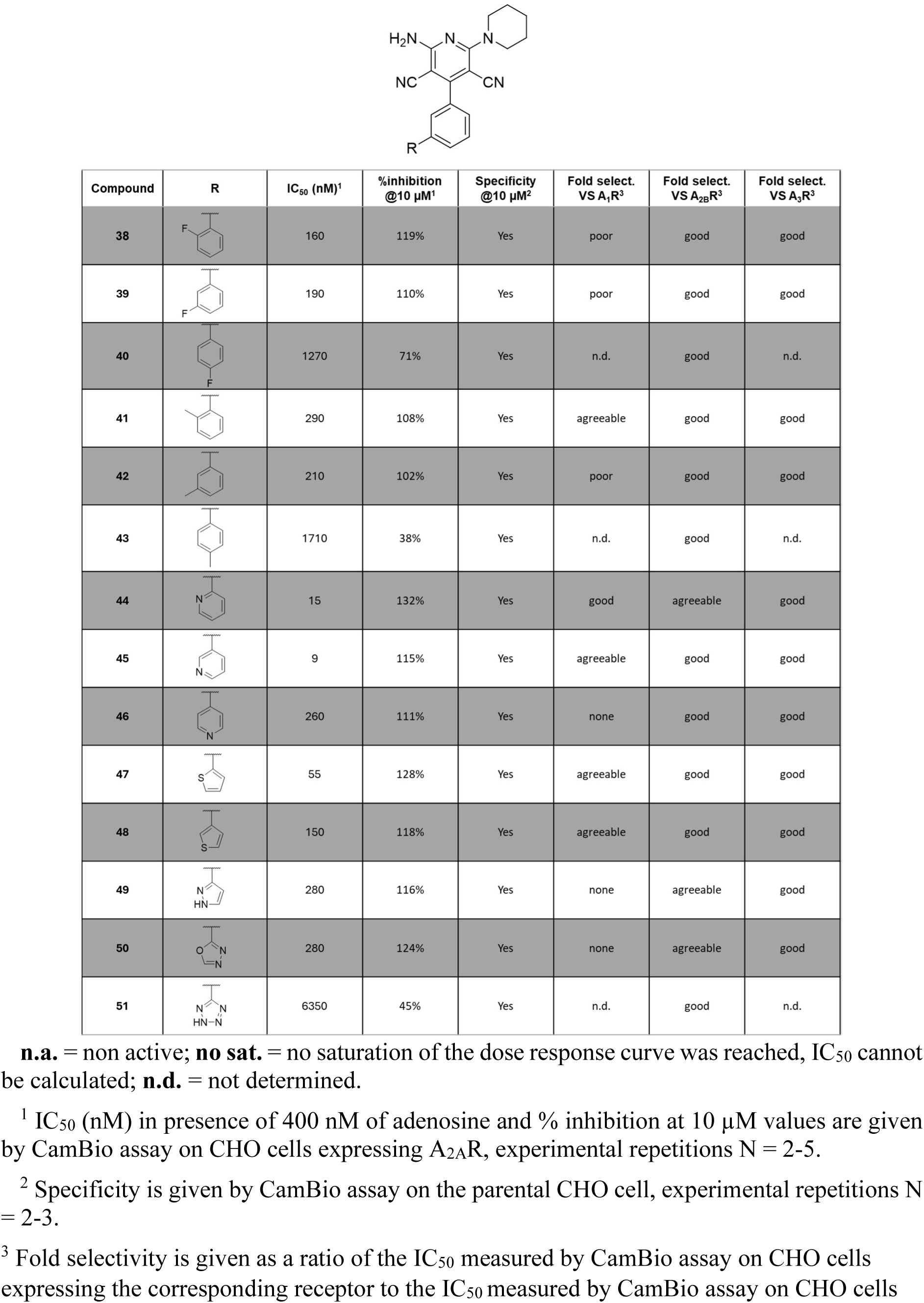
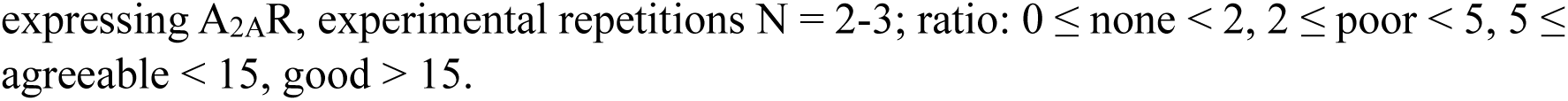
Biological effects of compounds 38-51.

| Compound | R | IC<sub>50</sub> (nM)<sup>1</sup> | %inhibition @10 μM<sup>1</sup> | Specificity @10 μM<sup>2</sup> | Fold select. VS A<sub>1</sub>R<sup>3</sup> | Fold select. VS A<sub>2B</sub>R<sup>3</sup> | Fold select. VS A<sub>3</sub>R<sup>3</sup> |
| --- | --- | --- | --- | --- | --- | --- | --- |
| 38 |  | 160 | 119% | Yes | poor | good | good |
| 39 |  | 190 | 110% | Yes | poor | good | good |
| 40 |  | 1270 | 71% | Yes | n.d. | good | n.d. |
| 41 |  | 290 | 108% | Yes | agreeable | good | good |
| 42 |  | 210 | 102% | Yes | poor | good | good |
| 43 |  | 1710 | 38% | Yes | n.d. | good | n.d. |
| 44 |  | 15 | 132% | Yes | good | agreeable | good |
| 45 |  | 9 | 115% | Yes | agreeable | good | good |
| 46 |  | 260 | 111% | Yes | none | good | good |
| 47 |  | 55 | 128% | Yes | agreeable | good | good |
| 48 |  | 150 | 118% | Yes | agreeable | good | good |
| 49 |  | 280 | 116% | Yes | none | agreeable | good |
| 50 |  | 280 | 124% | Yes | none | agreeable | good |
| 51 |  | 6350 | 45% | Yes | n.d. | good | n.d. |

**n.a.** = non active; **no sat.** = no saturation of the dose response curve was reached, IC<sub>50</sub> cannot be calculated; **n.d.** = not determined.
<sup>1</sup> IC<sub>50</sub> (nM) in presence of 400 nM of adenosine and % inhibition at 10 μM values are given by CamBio assay on CHO cells expressing A<sub>2A</sub>R, experimental repetitions N = 2-5.
<sup>2</sup> Specificity is given by CamBio assay on the parental CHO cell, experimental repetitions N = 2-3.
<sup>3</sup> Fold selectivity is given as a ratio of the IC<sub>50</sub> measured by CamBio assay on CHO cells expressing the corresponding receptor to the IC<sub>50</sub> measured by CamBio assay on CHO cells
expressing A<sub>2A</sub>R, experimental repetitions N = 2-3; ratio: 0 ≤ none < 2, 2 ≤ poor < 5, 5 ≤ agreeable < 15, good > 15.

### Compounds 4 and 5 engage A_2A_R in an allosteric manner

As this discovery program focused on the development of allosteric modulators, extra care was taken to identify the compounds’ mode of action. By definition, allosteric modulators will only exert an effect in presence of any ligand and reach a saturable effect once the system is fully occupied. In the case of negative allosteric modulators, probe-dependency is harder to demonstrate as protein activity will be lowered in cellular assays whether binding is competitive (i.e., orthosteric) or not, hampering many of the NAM discovery programs. However, allosteric modulators exert an effect on ligand binding cooperativity: they affect ligand binding kinetics by affecting the protein’s conformation.

Grating-coupled interferometry (GCI) is a sensitive label-free method to characterize ligand binding to membrane proteins immobilized on a surface. Dissociation constants (K_D_) and kinetic rates (*k*_on_, *k*_off_) of XAC, a xanthine-based A_2A_R orthosteric antagonist, binding to A_2A_R were measured at increasing concentrations of compound **4** present throughout the measurement from the baseline to XAC dissociation. Notably, compound **4** was able to negatively modulate XAC affinity towards A_2A_R, with an approximately four-fold increase in K_D_ (Figure 5 and Supporting Information Figure S2 for compound **5**). Both, association and dissociation rates show to contribute to the difference in affinity, showing a decrease in *k*_on_ as well as an increase in *k*_off_ (shorter residence time) at higher concentrations of compound **4**. Furthermore, the calculated binding signal at XAC saturation (R_max_) remained at the same level, which indicates absence of any binding competition and therefore independent, allosteric interaction of compound **4** with A_2A_R. Together, these findings support a mode of action of compound **4** as a NAM.

**Figure 5.**
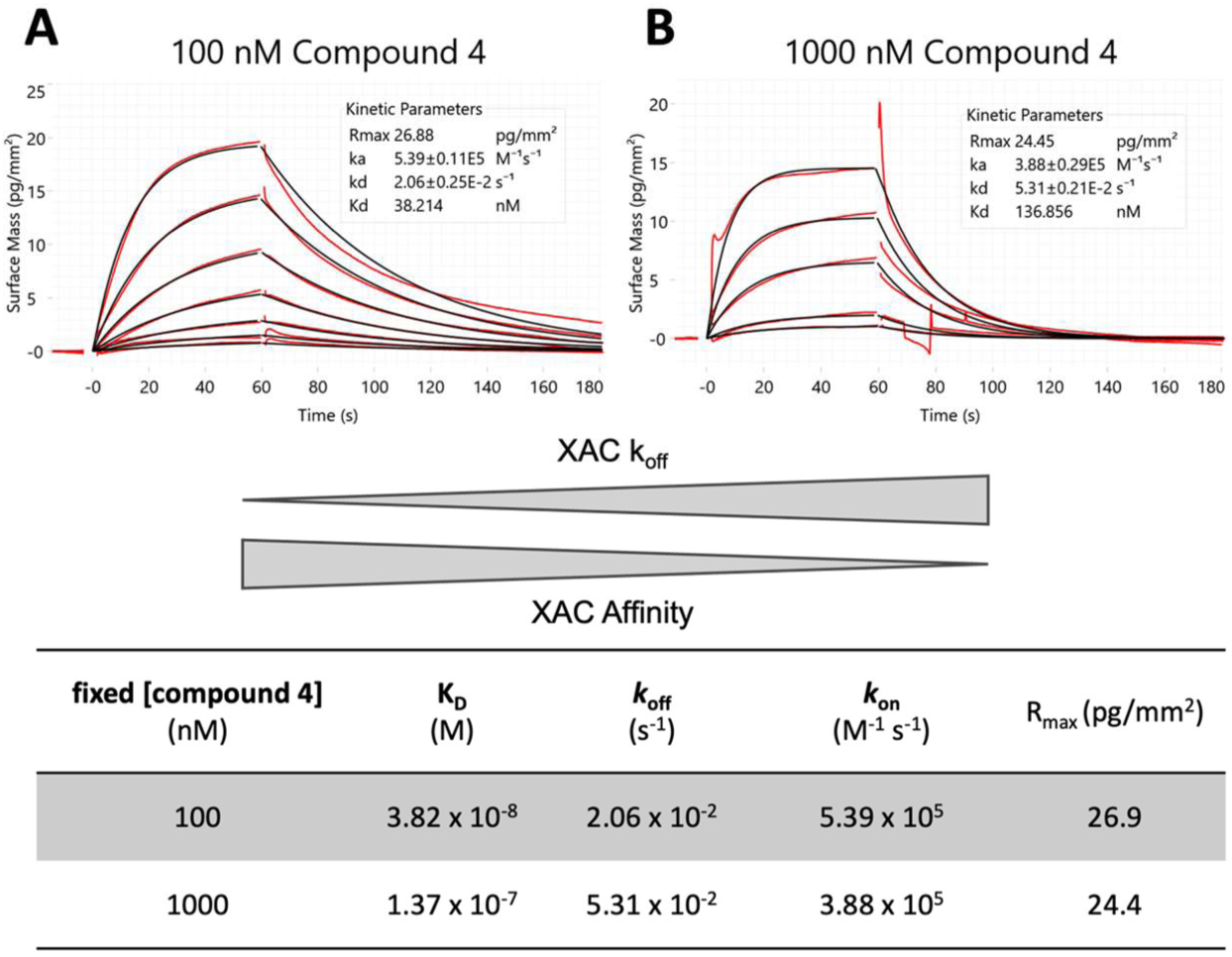
GCI kinetic characterization of XAC binding to A_2A_R at different concentrations of compound **4**: (A) 100 nM and (B) 1000 nM. Double-referenced binding signals are shown in red, 1:1 Langmuir interaction model fits are shown in black. Table: Kinetic parameters derived from fitting double-referenced binding signals to a 1:1 interaction model at two concentrations of compound **4**.

### The allosteric mode of action is conserved throughout the chemical family: from compound 5 to 28 to 45

Allosteric mode of action can further be supported by demonstrating the non-competitive binding of the NAMs and the saturable antagonism effect of the system through binding at an allosteric site. Saturation can be shown in a cellular assay by doing several dose response curves of an agonist with fixed concentrations of the tested compound, termed “shift assay”. Results for selected examples and Imaradenant (AZD4635), an orthosteric antagonist, are presented Figure. The curve stacking pattern observed for the compounds **5**, **28** and **45** Figure 6A-C (and not for Figure 6D) is characteristic of their allosteric mode of action^12,29^ and conserved across the chemical family, and more specifically compounds presented Table 4.

**Figure 6.**
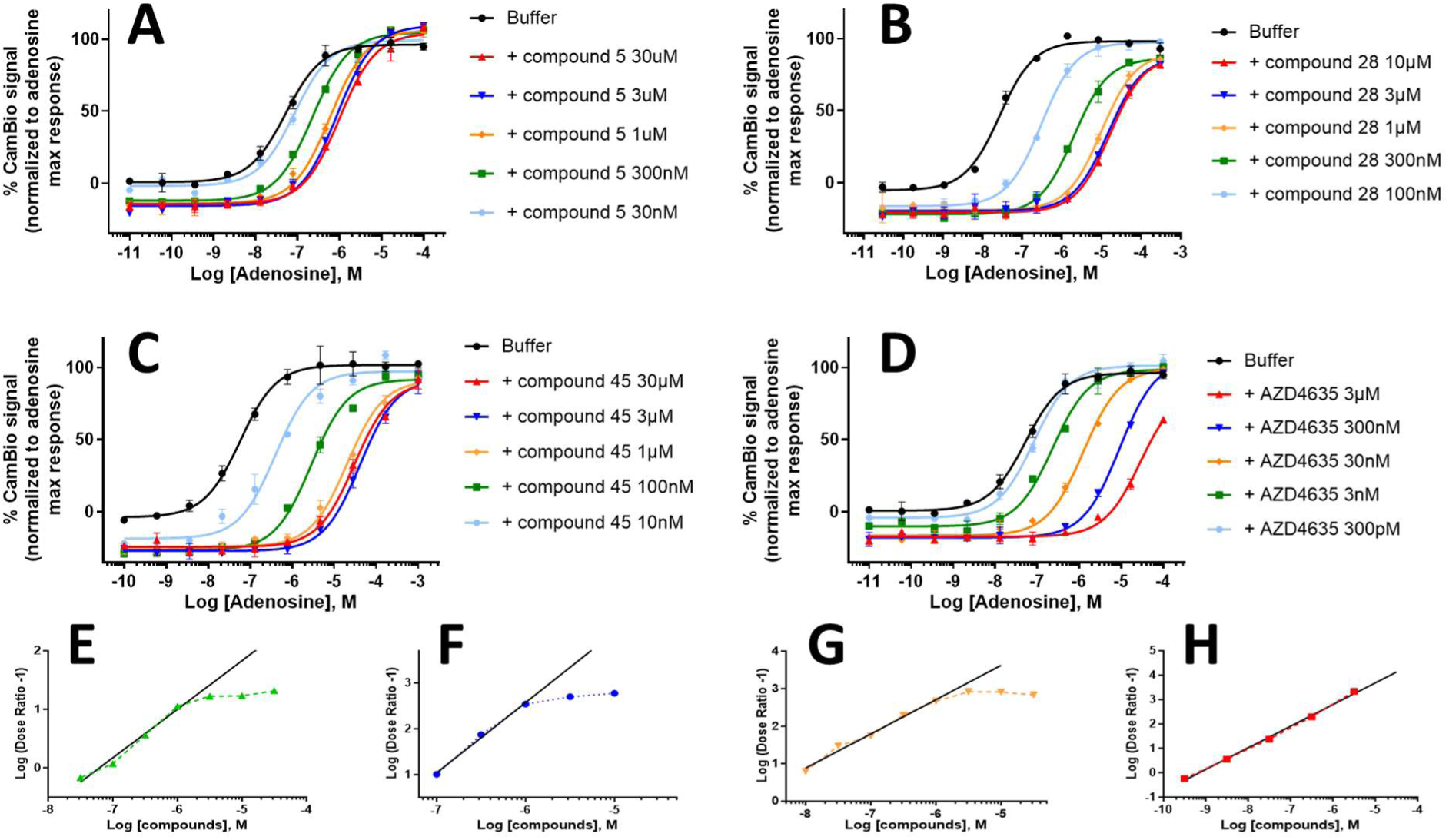
Respective shift assays and Schild plot analysis for compounds **5** (A&E), **28** (B&F) and **45** (C&G) and Imaradenant (D&H). The curve stacking in the shift assay confirms an allosteric compound mode of action.

These shift assay results can be exploited in a Schild plot analysis (regression of Log(Dose Ratio-1) versus Log[compound]) to quantify compounds’ allosteric effect.^12,29^ Maximum NAM antagonist activity is expected to occur when all allosteric sites are occupied by allosteric modulators in the presence of orthosteric agonists. Binding of both ligands being non-competitive, the corresponding Schild plot will present non-linear proportions. This is indeed observed for NAMs **5**, **28** and **45** (Figure 6E-G), highlighted in contrast to the linear dependency of the Log[compound] with the Log(Dose Ratio-1) for orthosteric ligands (Figure 6H).

Allosteric modulators can exert an independent effect on both the binding kinetic and the operational efficacy of a protein-ligand complex, which are reflected in the binding cooperativity factor α and the binding efficacy factor β. GCI measurements already showed that these NAMs acts on protein cooperativity, but the lack of variation in the maximum response of adenosine in the shift assays demonstrates that they act solely on the A_2A_R-ligand complex formation (Figure 6A-C). Hence, their Schild plots follow the equation (1), from which numeric values such as the binding cooperativity factor α and the equilibrium dissociation constant of the modulator-receptor complex K_B_ can be calculated (Table 5). This allowed us to quantify the improvement of the allosteric properties of the NAMs throughout the series. A difference of a thousand-fold was found between the α of the hit compound **5** and compound **45** demonstrating the strong participation of the additional substituents to the allosteric binding cooperativity. Additionally, the binding constant K_B_ was gradually lowered throughout the optimization process, supporting that the various substituents directly improved the NAMs’ affinity for A_2A_R.

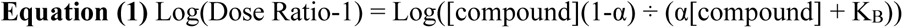

**Table 5.**
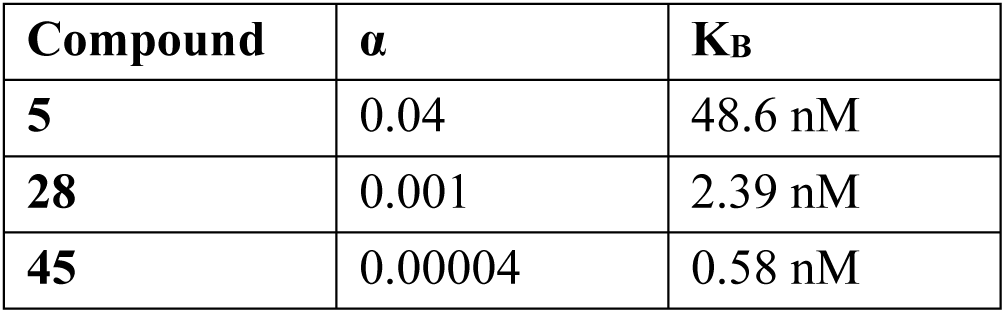
Determined binding cooperativity factor α and dissociation constant K_B_ for chosen NAM compounds.

| Compound | $\alpha$ | K<sub>B</sub> |
| --- | --- | --- |
| **5** | 0.04 | 48.6 nM |
| **28** | 0.001 | 2.39 nM |
| **45** | 0.00004 | 0.58 nM |

### NAMs retain their potency in high adenosine environments

Allosteric modulators being non-competitive ligands for protein binding, it is expected that their potency is retained even when endogenous ligand concentration is high. In the case of A_2A_R cancer immunotherapies, the drugs must be able to maintain potency in the TME which can accumulate up to µM concentrations of adenosine.^3^ Their fold loss of activity can be measured as a fold change in IC_50_ from low to highly immunosuppressive adenosine concentrations in cellular assays (20 nM – 3 µM, Figure 7).

**Figure 7.**
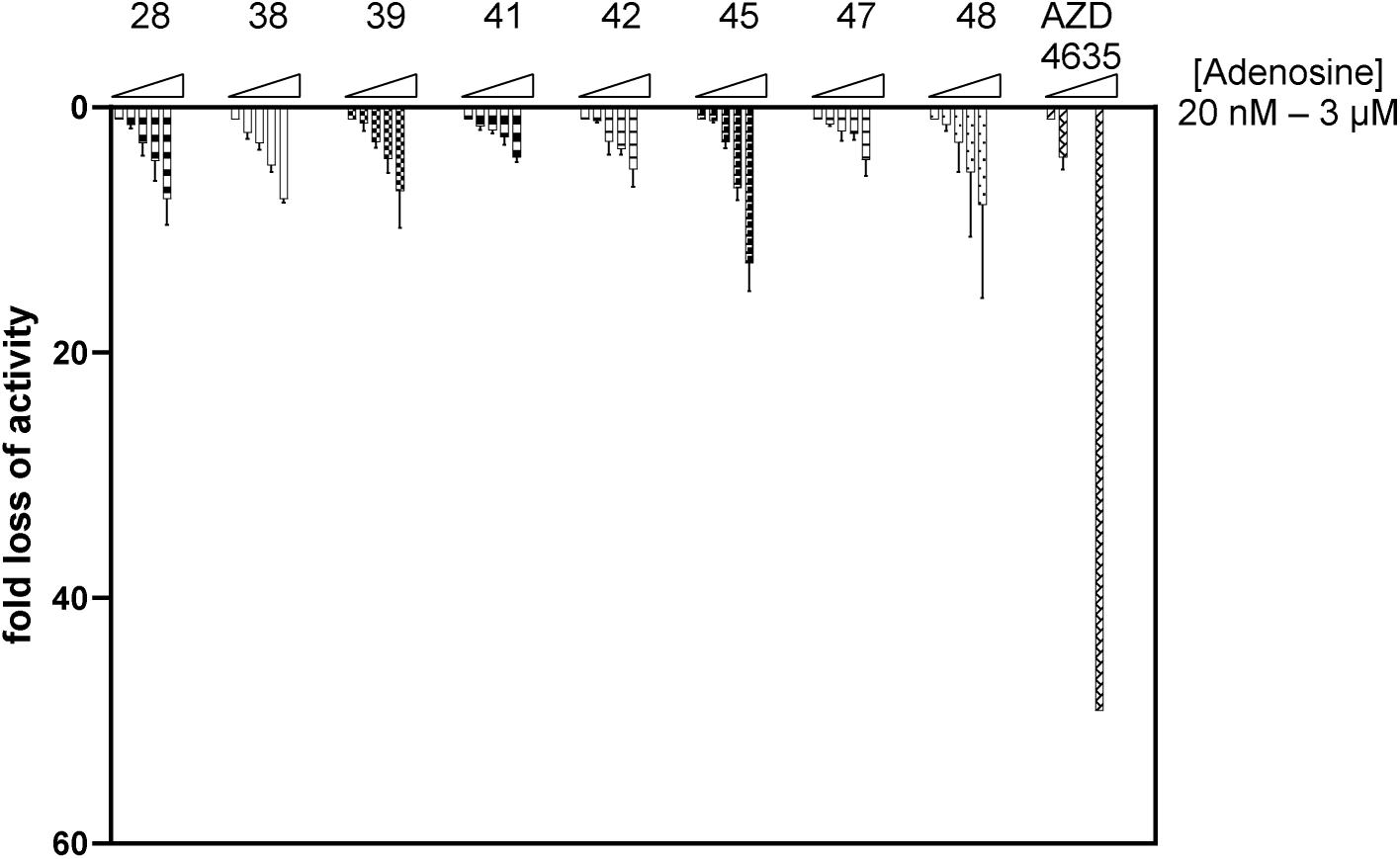
Fold loss of A_2A_R activity at increasing adenosine concentrations for selected NAMs and the orthosteric antagonist Imaradenant (AZD4635); given relative to their activity in presence of 20 nM of adenosine.

The NAMs exhibited various behaviors across the series with most compounds exhibiting a loss of activity smaller than 10-times than 10-times (e.g., compounds **28**, **38**, **41, 42** and **48**), which is expected to result in a retained potency in high adenosine environments. Other compounds, such as **45** and **48**, exhibited larger fold changes easily explained by their greater potency, resulting in wider shifts. Comparatively to orthosteric compounds, all NAMs showed significantly lower fold loss of activity inherent to their allosteric, non-competitive, binding mode.

### Proof of target engagement and downstream cascade activation for optimized compounds

The ability of selected NAMs to act on the A_2A_R-immunomodulatory pathway was analyzed by the measurement of downstream signaling effectors. As A_2A_R is known to act through the cAMP-PKA-CREB pathway,^1^ the downstream phosphorylation of CREB could be measured by phospho-flow cytometry in human blood immune cell populations obtained from healthy volunteers. Peripheral blood mononuclear cells (PBMCs) were cocultured in presence of NAMs then exposed to the highly immunosuppressive condition of 5 µM NECA (a stable adenosine analog), recapitulating an adenosine-rich TME. The results for pCREB intracellular staining in a CD4^+^ T lymphocytes (known to naturally express high levels of A_2A_R) are presented as a % restoration of a normal immune response (Figure 8). Early-development compound **5** demonstrated no immunomodulatory activity on CD4^+^ T cells. Interestingly, the orthosteric antagonist Imaradenant (AZD4635), which was dropped from clinical trials, demonstrated only partial restoration (54% ±10%). Meanwhile, optimized compounds such as **28**, **38**, and others showed the ability to fully restore the baseline levels of pCREB, indicating complete immune restoration at 1 µM of compound (Figure 8A). Titration of compound **28** revealed that immunological responsiveness of CD4^+^ T cells could be partially restored in translational settings using other concentrations of NAMs, including sub-micromolar concentrations (Figure 8B).

**Figure 8.**
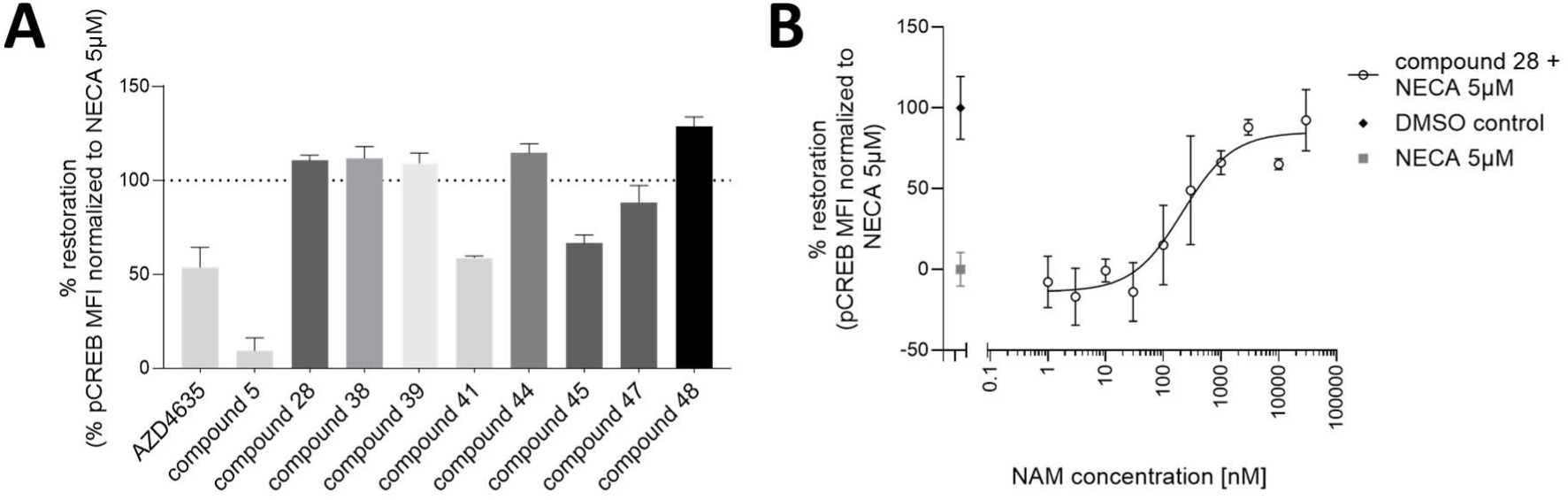
(A) Restoration of a normal immune response by representative compounds measured by the diminution of pCREB in CD4^+^ T cells immunosuppressed by 5µM NECA, mimicking an adenosine-rich TME. All compounds were tested at 1 µM, repetitions N = 2-15. (B) Compound **28** titration in presence of 5 µM NECA. Normal level of pCREB is given by the DMSO control, whereas maximal immunosuppression is shown in the “NECA 5 µM” contr

In a retrospective analysis for the NAM compounds, it is possible to correlate these encouraging translational results with these compounds’ full % inhibition of A_2A_R activity rather than based on their primary screen IC_50_ alone.

## Conclusion

In conclusion, we identified A_2A_R allosteric hits in a high-throughput screening campaign and confirmed their NAM mode of action. The following SAR campaign is the first expensively reported for A_2A_R allosteric modulators and led to the discovery of potent 2-amino-3,5-dicyanopyridine derivatives such as compounds **28** and **45**, with respective IC_50_ of 90 nM and 9 nM in the presence of 400 nM of adenosine and a full % inhibition of receptor activity at 10 µM. Both compounds allosteric mode of action could be demonstrated by shift assay and non-linearity of the Schild plot analysis and retained satisfactory potencies in high-adenosine concentrations. Correlating A_2A_R engagement and downstream signaling was possible thanks to the phosphorylation measurement of the downstream effector CREB for all the optimized NAMs (with IC_50_ < 200 nM and full % inhibition of receptor activity), clearly showcasing the potential of A_2A_R allosteric modulation as a novel approach for efficient and safer cancer immunotherapies.

## Material and methods

Unless otherwise noted, all products were obtained from commercial sources and used without further purification. Anhydrous solvents such as methanol, DMF or toluene were purchased over molecular sieve, enclosed by AcroSeal^®^ bottles. All preparative columns were performed by flash chromatography on a Büchi Pure C-815 Flash system with a UV detector. The corresponding PureFlash ID cartridges (4 g, 12 g, or 24 g, amorphous silica, 35–45 µm mesh) were purchased from Büchi and the flow rates were set according to the preset parameters (15, 30 & 32 mL/min respectively). Samples were loaded as solid deposit prepared with amorphous silica 40–60 µm mesh. All described yields are isolated unless stated otherwise. The ^1^H NMR spectra were recorded on a Bruker 600 MHz spectrometer equipped with a cryoprobe and are calibrated on the residual protonated solvent. The ^13^C NMR spectra were recorded at 125 MHz and the solvent resonance is used as internal standard. Both ^1^H NMR and ^13^C NMR chemical shifts are reported in parts per million downfield from tetramethylsilane. Low resolution mass spectra were recorded on an Advion PressionL, coupled to an ESI source, operating in positive and negative ion mode simultaneously. HRMS were recorded on a Xevo G2 TOF, coupled to an ESI source, operating either in positive or in negative acquisition mode.

### General procedures

#### General procedure A

Malononitrile (2 eq.), dry methanol (0.5 M), aldehyde (1 eq.), 4-DMAP (0.2 eq.) and piperidine (1.2 eq.) were sequentially added to a dry 10 mL round-bottom flask equipped with a magnetic stirrer. The flask was closed by a guard funnel filled with CaCl_2_ beads (to allow air exchange) and stirred at r.t. (r.t. range = 19-22°C) for 16 hours. Completion was assessed by TLC (cyclo:EtOAc 8:2). The medium was diluted with DCM (5 × V_MeOH_), silica was directly added on top of the mixture and the solvent was removed under reduced pressure. The crude was purified by flash chromatography on a pre-packed silica cartridge (12 g cartridge for 0.5 to 1 mmol of starting aldehyde) with a cyclo:EtOAc gradient (9:1→8:2, 6 CV; 8:2→1:1, 3 CV; 1:1, 3 CV; methanol wash).

#### General procedure B

Malononitrile (1 eq.), dry methanol (0.5 M), aldehyde (1 eq.) and 4-DMAP (0.2 eq.) were sequentially added to a dry 10 mL round-bottom flask equipped with a magnetic stirrer. The flask was closed by a guard funnel filled with CaCl_2_ beads (to allow air exchange) and stirred at r.t. for 15 min. The solution was cooled at 0°C before the addition of more malononitrile (1 eq.) and the dropwise addition of piperidine. The solution was slowly brought back to r.t. and stirred for 16 h overall. Completion was assessed by TLC (cyclo:EtOAc 8:2). The medium was diluted with DCM (5 × V_MeOH_), silica was directly added on top of the mixture and the solvent was removed under reduced pressure. The crude was purified by flash chromatography (12 g cartridge for 0.5 to 1 mmol of starting aldehyde) with a cyclo:EtOAc gradient (9:1→8:2, 6 CV; 8:2, 3 CV; 8:2→1:1, 3 CV; methanol wash).

#### General procedure C

In a small round-bottom flask equipped with a stirbar, the aldehyde (1 eq.) and the malononitrile (1 eq.) were solubilized in methanol (0.2 M) (solution A). The solution A was vigorously stirred right away. In case the aldehyde contains a basic site (e.g., pyridine), no catalyst is needed. Else, in a small vial, a solution of cyclic amine (1.2 eq.) in MeOH (0.2 M) was prepared (solution B). A drop or two of the solution B was added to the solution A. After stirring 30 min at r.t., more malononitrile (1 eq.) was added followed by dropwise addition of the remaining solution B. The reaction was stirred at r.t., open to air until completion, which was assessed by TLC (cyclo:EtOAc 8:2). The medium was diluted with DCM (5 × V_MeOH_), silica was directly added on top of the mixture and the solvent was removed under reduced pressure. The crude was purified by flash chromatography (12 g cartridge for 0.5 to 1 mmol of starting aldehyde) with a cyclo:EtOAc gradient (95:5, 3 CV; 95:5→8:2, 12 CV; 8:2, 6 CV; methanol wash).

#### General procedure D

In a 10 mL round-bottom flask equipped with a magnetic stirrer, 2-amino-4-(3-bromophenyl)-6-(piperidin-1-yl)pyridine-3,5-dicarbonitrile (1 eq.), boronic acid (3 eq.), potassium carbonate (3 eq.) and palladium chloride (0.05 eq.) were consecutively added to toluene (0.15 M). The mixture was stirred at 110°C for 3 h, open to air. Completion was assessed by TLC (cyclo:DCM:EtOAc 6:3:1). Palladium and salts were removed by filtration on a celite pad, washed with DCM (2 × 5 mL). Finally, the solvents were removed under reduced pressure and the crude product purified by flash chromatography on a silica cartridge (4 g cartridge for 0.15 mmol of starting aldehyde), solid deposit, with a cyclo:EtOAc gradient (10:0, 3 CV; 10:0→8:2, 9 CV; 8:2, 3 CV; methanol wash).

#### General procedure E

In a 2-5 mL microwave tube equipped with a magnetic stirrer, 2-amino-4-(3-bromophenyl)-6-(piperidin-1-yl)pyridine-3,5-dicarbonitrile (1 eq.), boronic acid (1.2 or 3 eq.) and potassium carbonate (3 eq.) were solubilized in a THF:H_2_O 2:1 mixture (0.05 M). The solution was bubbled with argon for 10 min before addition of tetrakis(triphenylphosphine)palladium(0) (0.05 eq.). The tube was sealed and heated at 80°C or 100°C for 15 to 30 min under microwave irradiation. Completion could not be assessed by TLC. The medium was diluted with water and the crude product was extracted by EtOAc × 3. solvent was removed under reduced pressure. The crude product was purified by flash chromatography on a silica cartridge (4 g cartridge for less than 0.5 mmol of starting bromoaryl), solid deposit, with a cyclo:EtOAc gradient (9:1, 6 CV; 9:1→6:4, 20 CV; 6:4, 6 CV; methanol wash).

#### General procedure F

In a 2-5 mL microwave tube equipped with a magnetic stirrer, (3-(2-amino-3,5-dicyano-6-(piperidin-1-yl)pyridin-4-yl)phenyl)boronic acid pinacol ester (1 eq.), bromoaryl (1.2 eq.) and potassium carbonate (3 eq.) were solubilized in a THF:H_2_O 2:1 mixture (0.05 M). The solution was bubbled with argon for 10 min before addition of tetrakis(triphenylphosphine)palladium(0) (0.05 eq.). The tube was sealed and heated at 80°C for 15 min under microwave irradiation. The medium was diluted with water and the crude product was extracted by EtOAc × 3. The organic phases were combined, washed with brine, dried over MgSO_4_, filtered and the solvent was removed under reduced pressure. The crude product was purified by flash chromatography on a silica cartridge (4 g cartridge for less than 0.5 mmol of starting bromoaryl), solid deposit, with a cyclo:(toluene:acetone 4:1) gradient (5:5, 6 CV; 5:5→0:10, 6 CV; 0:10, 6 CV; methanol wash).

#### General procedure G

In a microwave compatible tube equipped with a magnetic stirrer, 2-amino-6-chloro-4-phenylpyridine-3,5-dicarbonitrile (1 eq.) was solubilized in an anhydrous THF:EtOH 4:1 mixture (0.15 M). The desired amine (6 eq.) was added dropwise, and the tube was sealed and heated 30 min to 1 h at 120°C under microwave irradiation. The completion of the reaction was assessed by TLC (cyclo:EtOAc 6:4). Brine and a few drops of HCl 1 M (final pH ∼ 1) were added and the crude was extracted with EtOAc × 3. The organic phases were combined, dried over MgSO_4_, filtered and the solvent was removed under reduced pressure. The crude product was purified by flash chromatography on a silica cartridge (4 g cartridge for 50 mg of starting halogenoaryl), solid deposit, with either an isocratic cyclo:EtOAc gradient (8:2, 6 CV; methanol wash) or a DCM:MeOH gradient (10:0, 3 CV; 10:0→0:10, 9 CV; methanol wash).

### Compound descriptions

#### 2-amino-4-phenyl-6-(piperidin-1-yl)pyridine-3,5-dicarbonitrile (5)

The product was prepared following the general procedure C starting from benzaldehyde (0.05 mL, 0.5 mmol, 1 eq.) and was isolated as a white powder (38 mg, 0.13 mmol, 25% isolated yield). **^1^H NMR (600 MHz, CDCl_3_) δ** 7.56 – 7.42 (m, 4H), 5.36 (s, 2H), 3.84 – 3.78 (m, 4H), 1.74 – 1.67 (m, 6H). **^13^C NMR (151 MHz, CDCl_3_) δ** 162.5, 161.3, 159.5, 134.9, 130.6, 128.9, 128.8, 117.8, 116.7, 83.8, 81.8, 49.3, 26.1, 24.6. **MS (ESI^+^) calculated for C_18_H_17_N_5_:** [M+H]^+^ *m/z* = 304.2, found *m/z* = 304.0. **HRMS (ESI^+^) calculated for C_18_H_17_N_5_:** [M+H] ^+^ *m/z* = 304.1557, found *m/z* = 304.1559.

#### 2-amino-6-(azocan-1-yl)-4-phenylpyridine-3,5-dicarbonitrile (6)

The product was prepared following the general procedure G starting from 2-amino-6-chloro-4-phenylpyridine-3,5-dicarbonitrile (40 mg, 0.16 mmol, 1 eq.) and azocane (0.04 mL, 2 eq.) and was isolated as a white amorphous powder (7 mg, 0.02 mmol, 13% isolated yield). **^1^H NMR (600 MHz, CDCl_3_) δ** 7.53 – 7.49 (m, 3H), 7.49 – 7.44 (m, 2H), 5.64 (s, 2H), 3.92 (t, *J* = 5.8 Hz, 4H), 1.92 – 1.86 (m, 4H), 1.64 – 1.61 (m, 4H), 1.58 – 1.54 (m, 2H). **^13^C NMR (151 MHz, CDCl_3_) δ** 163.4, 158.7, 157.6, 135.1, 130.6, 128.9, 128.7, 117.9, 116.4, 81.9, 81.8, 52.5, 27.2, 27.1, 25.1. **MS (ESI^+^) calculated for C_20_H_21_N_5_:** [M+H]^+^ *m/z* = 332.2, found *m/z* = 332.1. **HRMS (ESI^+^) calculated for C_20_H_21_N_5_:** [M+H]^+^ *m/z* = 332.1870, found *m/z* = 332.1863.

#### 2-amino-6-(diethylamino)-4-phenylpyridine-3,5-dicarbonitrile (7)

The product was prepared following the general procedure G starting from 2-amino-6-chloro-4-phenylpyridine-3,5-dicarbonitrile (43 mg, 0.17 mmol, 1 eq.) and diethylamine (0.10 mL, 6 eq.) and was isolated as a white amorphous powder (6 mg, 0.02 mmol, 13% isolated yield). **^1^H NMR (600 MHz, CDCl_3_) δ** 7.50 (q, *J* = 3.7 Hz, 3H), 7.45 (dd, *J* = 6.7, 2.9 Hz, 2H), 5.37 (s, 2H), 3.74 (q, *J* = 7.0 Hz, 4H), 1.30 (t, *J* = 7.0 Hz, 6H). **^13^C NMR (151 MHz, CDCl_3_) δ** 163.0, 159.1, 158.6, 135.3, 130.4, 128.9, 128.7, 118.1, 116.7, 81.8, 81.5, 44.8, 13.7. **MS (ESI^+^) calculated for C_17_H_17_N_5_:** [M+H]^+^ *m/z* = 292.2, found *m/z* = 292.1. **HRMS (ESI^+^) calculated for C_17_H_17_N_5_:** [M+H]^+^ *m/z* = 292.1557, found *m/z* = 292.1555.

#### 2-amino-6-(bis(2-hydroxyethyl)amino)-4-phenylpyridine-3,5-dicarbonitrile (8)

The product was prepared following the general procedure G starting from 2-amino-6-chloro-4-phenylpyridine-3,5-dicarbonitrile (40 mg, 0.16 mmol, 1 eq.) and diethanolamine (0.03 mL, 2 eq.) and was isolated as a white amorphous powder (10 mg, 0.06 mmol, 19% isolated yield). **^1^H NMR (600 MHz, MeOD) δ** 7.54 – 7.48 (m, 3H), 7.48 – 7.41 (m, 2H), 3.97 (t, *J* = 5.7 Hz, 4H), 3.86 (t, *J* = 5.7 Hz, 4H). **^13^C NMR (151 MHz, MeOD) δ** 164.6, 161.1, 160.6, 137.3, 131.0, 129.7, 119.5, 117.1, 82.7, 81.7, 61.6, 54.9. **MS (ESI^+^) calculated for C_17_H_17_N_5_O_2_:** [M+Na]^+^ *m/z* = 346.1, found *m/z* = 346.0. **HRMS (ESI^+^) calculated for C_17_H_17_N_5_O_2_:** [M+H]^+^ *m/z* = 324.1456, found *m/z* = 324.1433.

#### 2-amino-4-methyl-6-(piperidin-1-yl)pyridine-3,5-dicarbonitrile (11)

The product was prepared following the general procedure C starting from excess acetaldehyde (1 mL), solventless, and a drop of piperidine. The intermediate was dried under an N_2_ flux and isolated as a translucent liquid. The second step was performed on 2-ethylidenemalononitrile and piperidine (0.05 mL, 1.2 eq.) and gave a white powder (9 mg, 0.04 mmol, 4% overall yield). **^1^H NMR (600 MHz, CDCl_3_) δ** 5.49 (s, 2H), 3.82 – 3.76 (m, 4H), 2.53 (s, 3H), 1.77 – 1.59 (m, 6H). **^13^C NMR (151 MHz, CDCl_3_) δ** 158.9, 117.2, 116.0, 49.4, 26.1, 24.5, 20.4. **MS (ESI^+^) calculated for C_13_H_15_N_5_:** [M+H]^+^ *m/z* = 242.1, found *m/z* = 242.2. **HRMS (ESI^+^) calculated for C_13_H_15_N_5_:** [M+H]^+^ *m/z* = 242.1401, found *m/z* = 242.1405.

#### 2-amino-4-(naphthalen-2-yl)-6-(piperidin-1-yl)pyridine-3,5-dicarbonitrile (12)

The product was prepared following the general procedure A starting from 2-naphthaldehyde (78 mg, 0.5 mmol, 1 eq.) and was isolated as a light orange paste (36 mg, 0.10 mmol, 20% isolated yield). **^1^H NMR (600 MHz, CDCl_3_) δ** 8.00 (d, *J* = 1.8 Hz, 1H), 7.98 (d, *J* = 8.5 Hz, 1H), 7.93 (dd, *J* = 7.7, 1.8 Hz, 1H), 7.56 (td, *J* = 7.6, 1.6 Hz, 3H), 5.38 (s, 2H), 3.85 – 3.80 (m, 4H), 1.73 – 1.69 (m, 6H). **^13^C NMR (151 MHz, CDCl_3_) δ** 162.4, 161.3, 159.6, 134.2, 133.0, 132.3, 129.1, 128.85, 128.83, 128.0, 127.6, 126.9, 125.7, 117.9, 116.8, 84.0, 82.0, 49.4, 26.2, 24.6. **MS (ESI^+^) calculated for C_22_H_19_N_5_:** [M+H]^+^ *m/z* = 354.2, found *m/z* = 354.4. **HRMS (ESI^+^) calculated for C_22_H_19_N_5_:** [M+Na]^+^ *m/z* = 376.1533, found *m/z* = 376.1524.

#### 2-amino-6-(piperidin-1-yl)-4-(o-tolyl)pyridine-3,5-dicarbonitrile (13)

The product was prepared following the general procedure A starting from 2-methylbenzaldehyde (0.06 mL, 0.5 mmol, 1 eq.) and was isolated as a white powder (41 mg, 0.13 mmol, 26% isolated yield). **^1^H NMR (600 MHz, CDCl_3_) δ** 7.38 (td, *J* = 7.5, 1.4 Hz, 1H), 7.34 – 7.27 (m, 2H), 7.17 (dd, *J* = 7.6, 1.4 Hz, 1H), 5.53 (s, 2H), 3.85 – 3.80 (m, 4H), 2.27 (s, 3H), 1.74 – 1.69 (m, 6H). **^13^C NMR (151 MHz, CDCl_3_) δ** 163.4, 159.7, 158.9, 135.1, 134.8, 130.9, 130.2, 128.1, 126.4, 116.8, 115.8, 84.6, 82.8, 49.5, 26.1, 24.4, 19.5. **MS (ESI^+^) calculated for C_19_H_19_N_5_:** [M+H]^+^ *m/z* = 318.2, found *m/z* = 318.2. **HRMS (ESI^+^) calculated for C_19_H_19_N_5_:** [M+H]^+^ *m/z* = 318.1714, found *m/z* = 318.1730.

#### 2-amino-6-(piperidin-1-yl)-4-(m-tolyl)pyridine-3,5-dicarbonitrile (14)

The product was prepared following the general procedure A starting from 3-methylbenzaldehyde (0.06 mL, 0.5 mmol, 1 eq.) and was isolated as a white powder (45 mg, 0.14 mmol, 28% isolated yield). **^1^H NMR (600 MHz, CDCl_3_) δ** 7.39 (t, *J* = 8.0 Hz, 1H), 7.31 (d, *J* = 7.2 Hz, 1H), 7.30 – 7.25 (m, 2H), 5.52 (s, 2H), 3.85 – 3.78 (m, 4H), 2.43 (s, 3H), 1.74 – 1.69 (m, 6H). **^13^C NMR (151 MHz, CDCl_3_) δ** 162.9, 160.7, 159.3, 138.7, 134.7, 131.5, 129.3, 128.9, 125.9, 117.6, 116.4, 83.9, 82.0, 49.7, 26.1, 24.5, 21.6. **MS (ESI^+^) calculated for C_19_H_19_N_5_:** [M+H]^+^ *m/z* = 318.2, found *m/z* = 318.2. **HRMS (ESI^+^) calculated for C_19_H_19_N_5_:** [M+H]^+^ *m/z* = 318.1714, found *m/z* = 318.1730.

#### 2-amino-6-(piperidin-1-yl)-4-(p-tolyl)pyridine-3,5-dicarbonitrile (15)

The product was prepared following the general procedure A starting from 4-methylbenzaldehyde (0.06 mL, 0.5 mmol, 1 eq.) and was isolated as a white powder (35 mg, 0.11 mmol, 22% isolated yield). **^1^H NMR (600 MHz, CDCl_3_) δ** 7.39 (d, *J* = 7.9 Hz, 2H), 7.31 (d, *J* = 7.8 Hz, 2H), 5.40 (s, 2H), 3.82 – 3.77 (m, 4H), 2.42 (s, 3H), 1.73 – 1.68 (m, 6H). **^13^C NMR (151 MHz, CDCl_3_) δ** 162.7, 161.1, 159.5, 140.9, 131.9, 129.7, 128.8, 117.9, 116.7, 83.8, 81.8, 49.5, 26.1, 24.5, 21.6. **MS (ESI^+^) calculated for C_19_H_19_N_5_:** [M+H]^+^ *m/z* = 318.2, found *m/z* = 318.2. **HRMS (ESI^+^) calculated for C_19_H_19_N_5_:** [M+H]^+^ *m/z* = 318.1714, found *m/z* = 318.1702.

#### 2-amino-4-(2-chlorophenyl)-6-(piperidin-1-yl)pyridine-3,5-dicarbonitrile (16)

The product was prepared following the general procedure A starting from 2-chlorobenzaldehyde (0.06 mL, 0.5 mmol, 1 eq.) and was isolated as a white powder (48 mg, 0.14 mmol, 28% isolated yield). **^1^H NMR (600 MHz, CDCl_3_) δ** 7.54 (dd, *J* = 7.9, 1.4 Hz, 1H), 7.44 (td, *J* = 7.7, 1.9 Hz, 1H), 7.40 (td, *J* = 7.5, 1.4 Hz, 1H), 7.31 (dd, *J* = 7.5, 1.9 Hz, 1H), 5.36 (s, 2H), 3.84 – 3. 79 (m, 4H), 1.75 – 1.66 (m, 7H). **^13^C NMR (151 MHz, CDCl_3_) δ** 160.3, 159.6, 158.9, 134.1, 132.3, 131.6, 130.5, 129.9, 127.5, 116.6, 115.5, 84.7, 82.7, 49.5, 26.1, 24.4. **MS (ESI^+^) calculated for C_18_H_16_N_5_Cl:** [M+H]^+^ *m/z* = 338.1, found *m/z* = 338.0. **HRMS (ESI^+^) calculated for C_18_H_16_N_5_Cl:** [M+H]^+^ *m/z* = 338.1167, found *m/z* = 338.1168.

#### 2-amino-4-(3-chlorophenyl)-6-(piperidin-1-yl)pyridine-3,5-dicarbonitrile (17)

The product was prepared following the general procedure A starting from 3-chlorobenzaldehyde (0.06 mL, 0.5 mmol, 1 eq.) and was isolated as a white powder (18 mg, 0.05 mmol, 11% isolated yield). **^1^H NMR (600 MHz, CDCl_3_) δ** 7.49 (ddd, *J* = 8.1, 2.3, 1.3 Hz, 1H), 7.47 – 7.42 (m, 2H), 7.36 (dt, *J* = 7.6, 1.5 Hz, 1H), 5.43 (s, 2H), 3.81 (dd, *J* = 6.4, 3.8 Hz, 4H), 1.73 – 1.67 (m, 6H). **^13^C NMR (151 MHz, CDCl_3_) δ** 161.0, 160.5, 159.2, 136.5, 134.9, 130.8, 130.4, 128.8, 127.0, 117.3, 116.1, 83.6, 81.8, 49.5, 26.1, 24.5. **MS (ESI^+^) calculated for C_18_H_16_N_5_Cl:** [M+H]^+^ *m/z* = 338.1, found *m/z* = 338.1. **HRMS (ESI^+^) calculated for C_18_H_16_N_5_Cl:** [M+H]^+^ *m/z* = 338.1167, found *m/z* = 338.1168.

#### 2-amino-4-(4-chlorophenyl)-6-(piperidin-1-yl)pyridine-3,5-dicarbonitrile (18)

The product was prepared following the general procedure A starting from 4-chlorobenzaldehyde (70 mg, 0.5 mmol, 1 eq.) and was isolated as a white powder (42 mg, 0.12 mmol, 25% isolated yield). **^1^H NMR (600 MHz, CDCl_3_) δ** 7.48 (d, *J* = 8.5 Hz, 1H), 7.43 (d, *J* = 8.5 Hz, 1H), 5.40 (s, 1H), 3.82 – 3.77 (m, 2H), 1.72 – 1.66 (m, 3H). **^13^C NMR (151 MHz, CDCl_3_) δ** 161.3, 160.7, 159.3, 137.0, 133.2, 130.2, 129.4, 117.5, 116.3, 83.5, 81.7, 49.5, 26.1, 24.5. **MS (ESI^+^) calculated for C_18_H_16_N_5_Cl:** [M+H]^+^ *m/z* = 338.1, found *m/z* = 338.0. **HRMS (ESI^+^) calculated for C_18_H_16_N_5_Cl:** [M+H]^+^ *m/z* = 338.1167, found *m/z* = 338.1168.

#### 2-amino-4-(2-methoxyphenyl)-6-(piperidin-1-yl)pyridine-3,5-dicarbonitrile (19)

The product was prepared following the general procedure B starting from 2-methoxybenzaldehyde (0.06 mL, 0.5 mmol, 1 eq.) and was isolated as a white powder (16 mg, 0.05 mmol, 10% isolated yield). **^1^H NMR (600 MHz, CDCl_3_) δ** 7.46 (ddd, *J* = 8.4, 7.5, 1.7 Hz, 1H), 7.25 (dd, *J* = 7.5, 1.8 Hz, 1H), 7.07 (td, *J* = 7.5, 1.0 Hz, 1H), 7.04 (dd, *J* = 8.4, 1.0 Hz, 1H), 5.55 (s, 2H), 3.87 (s, 3H), 3.86 – 3.78 (m, 4H), 1.73 – 1.70 (m, 6H). **^13^C NMR (151 MHz, CDCl_3_) δ** 160.3, 159.0, 156.3, 132.1, 130.1, 123.8, 121.0, 117.3, 116.3, 111.8, 85.3, 83.2, 55.9, 49.5, 26.1, 24.5. **MS (ESI^+^) calculated for C_19_H_19_N_5_O:** [M+H]^+^ *m/z* = 334.2, found *m/z* = 334.5. **HRMS (ESI^+^) calculated for C_19_H_19_N_5_O:** [M+H]^+^ *m/z* = 334.1663, found *m/z* = 334.1641.

#### 2-amino-4-(3-methoxyphenyl)-6-(piperidin-1-yl)pyridine-3,5-dicarbonitrile (20)

The product was prepared following the general procedure B starting from 3-methoxybenzaldehyde (0.06 mL, 0.5 mmol, 1 eq.) and was isolated as a white powder (18 mg, 0.05 mmol, 11% isolated yield). **^1^H NMR (600 MHz, CDCl_3_) δ** 7.42 (t, *J* = 8.0 Hz, 1H), 7.05 (tdd, *J* = 7.6, 2.1, 0.8 Hz, 2H), 6.99 (dd, *J* = 2.5, 1.6 Hz, 1H), 5.60 (s, 2H), 3.85 (s, 3H), 3.84 – 3.79 (m, 4H), 1.74 – 1.69 (m, 6H). **^13^C NMR (151 MHz, CDCl_3_) δ** 162.5, 160.6, 159.7, 159.2, 135.9, 130.1, 121.0, 117.4, 116.5, 116.3, 114.2, 83.9, 82.0, 55.6, 49.6, 26.1, 24.4. **MS (ESI^+^) calculated for C_19_H_19_N_5_O:** [M+H]^+^ *m/z* = 334.2, found *m/z* = 334.5. **HRMS (ESI^+^) calculated for C_19_H_19_N_5_O:** [M+H]^+^ *m/z* = 334.1663, found *m/z* = 334.1641.

#### 2-amino-4-(4-methoxyphenyl)-6-(piperidin-1-yl)pyridine-3,5-dicarbonitrile (21)

The product was prepared following the general procedure B starting from 4-methoxybenzaldehyde (0.06 mL, 0.5 mmol, 1 eq.) and was isolated as a white powder (24 mg, 0.07 mmol, 14% isolated yield). **^1^H NMR (600 MHz, CDCl_3_) δ** 7.47 (d, *J* = 9.4 Hz, 2H), 7.01 (d, *J* = 9.4 Hz, 2H), 5.46 (s, 2H), 3.86 (s, 3H), 3.85 – 3.77 (m, 4H), 1.75 – 1.70 (m, 6H). **^13^C NMR (151 MHz, CDCl_3_) δ** 162.2, 161.6, 161.3, 159.6, 130.6, 126.9, 118.1, 116.9, 114.4, 83.7, 81.7, 55.5, 49.5, 27.1, 26.1, 24.5. **MS (ESI^+^) calculated for C_19_H_19_N_5_O:** [M+H]^+^ *m/z* = 334.2, found *m/z* = 334.5. **HRMS (ESI^+^) calculated for C_19_H_19_N_5_O:** [M+H]^+^ *m/z* = 334.1663, found *m/z* = 334.1641.

#### 2-amino-4-(3-hydroxyphenyl)-6-(piperidin-1-yl)pyridine-3,5-dicarbonitrile (22)

The product was prepared following the general procedure A starting from 3-hydroxybenzaldehyde (61 mg, 0.5 mmol, 1 eq.) and was isolated as a white powder (38 mg, 0.12 mmol, 24% isolated yield). **^1^H NMR (600 MHz, (CD_3_)_2_SO) δ** 9.75 (s, 1H), 7.39 (s, 2H), 7.31 (t, *J* = 7.9 Hz, 1H), 6.91 (ddd, *J* = 8.2, 2.5, 1.0 Hz, 1H), 6.86 (dt, *J* = 7.5, 1.3 Hz, 1H), 6.82 (t, *J* = 2.0 Hz, 1H), 3.73 – 3.68 (m, 4H), 1.69 – 1.54 (m, 6H). **^13^C NMR (151 MHz, (CD_3_)_2_SO) δ** 161.8, 160.7, 159.7, 157.2, 136.5, 129.7, 119.2, 117.7, 116.8, 116.2, 115.3, 81.4, 80.7, 48.5, 25.6, 23.9. **MS (ESI^+^) calculated for C_18_H_17_N_5_O:** [M+H]^+^ *m/z* = 320.2, found *m/z* = 320.4. **HRMS (ESI^+^) calculated for C_18_H_17_N_5_O:** [M+H]^+^ *m/z* = 320.1506, found *m/z* = 320.1500.

#### 2-amino-4-(3-phenoxyphenyl)-6-(piperidin-1-yl)pyridine-3,5-dicarbonitrile (23)

The product was prepared following the general procedure A starting from 3-phenoxybenzaldehyde (0.09 mL, 0.5 mmol, 1 eq.) and was isolated as a white powder (36 mg, 0.09 mmol, 18% isolated yield). **^1^H NMR (600 MHz, CDCl_3_) δ** 7.47 (t, *J* = 8.0 Hz, 1H), 7.36 (tt, *J* = 7.4, 2.1 Hz, 2H), 7.19 (dt, *J* = 7.7, 1.4 Hz, 1H), 7.17 – 7.15 (m, 1H), 7.15 – 7.08 (m, 4H), 5.44 (s, 2H), 82 – 3.76 (m, 4H), 1.72 – 1.67 (m, 6H). **^13^C NMR (151 MHz, CDCl_3_) δ** 161.8, 160.8, 159.3, 157.6, 156.7, 136.4, 130.5, 130.1, 123.8, 123.4, 120.9, 119.4, 118.9, 117.5, 116.3, 83.7, 81.8, 49.4, 26.1, 24.5. **MS (ESI^+^) calculated for C_24_H_21_N_5_O:** [M+H]^+^ *m/z* = 396.2, found *m/z* = 396.1. **HRMS (ESI^+^) calculated for C_24_H_21_N_5_O:** [M+H]^+^ *m/z* = 396.1819, found *m/z* = 396.1806.

#### 2-amino-4-(3-(benzyloxy)phenyl)-6-(piperidin-1-yl)pyridine-3,5-dicarbonitrile (24)

The product was prepared following the general procedure A starting from 3-benzyloxybenzaldehyde (106 mg, 0.5 mmol, 1 eq.) and was isolated as a white powder (61 mg, 0.15 mmol, 30% isolated yield). **^1^H NMR (600 MHz, CDCl_3_) δ** 7.47 – 7.38 (m, 5H), 7.34 (t, *J* = 7.4 Hz, 1H), 7.13 – 7.07 (m, 3H), 5.47 (s, 2H), 5.10 (s, 2H), 3.83 – 3.78 (m, 4H), 1.73 – 1.68 (m, 6H). **^13^C NMR (151 MHz, CDCl_3_) δ** 162.3, 160.9, 159.4, 159.0, 136.7, 136.1, 130.1, 128.7, 128.2, 127.8, 121.4, 117.6, 117.3, 116.5, 115.2, 83.7, 81.9, 70.4, 49.4, 26.1, 24.5. **MS (ESI^+^) calculated for C_25_H_23_N_5_O:** [M+H]^+^ *m/z* = 410.2, found *m/z* = 410.1. **HRMS (ESI^+^) calculated for C_25_H_23_N_5_O:** [M+Na]^+^ *m/z* = 432.1795, found *m/z* = 432.1769.

#### methyl 3-(2-amino-3,5-dicyano-6-(piperidin-1-yl)pyridin-4-yl)benzoate (25)

The product was prepared following the general procedure C starting from methyl 3-formylbenzoate (492 mg, 3 mmol, 1 eq.) and piperidine (0.59 mL, 2 eq.) and was isolated as a white powder (179 mg, 0.5 mmol,16% isolated yield). **^1^H NMR (600 MHz, CDCl_3_) δ** 8.19 (dt, *J* = 7.8, 1.5 Hz, 2H), 3.94 (s, 3H), 3.85 – 3.79 (m, 4H), 1.74 – 1.69 (m, 6H). **^13^C NMR (151 MHz, CDCl_3_) δ** 166.2, 161.4, 160.5, 159.2, 135.1, 132.9, 131.5, 130.9, 129.9, 129.1, 117.2, 116.1, 83.6, 81.7, 52.4, 49.3, 26.0, 24.4. **MS (ESI^+^) calculated for C_20_H_19_N_5_O_2_:** [M+H]^+^ *m/z* = 362.2, found *m/z* = 362.5. **HRMS (ESI^+^) calculated for C_20_H_19_N_5_O_2_:** [M+H]^+^ *m/z* = 362.1612, found *m/z* = 362.1608.

#### 2-amino-4-(3-fluorophenyl)-6-(piperidin-1-yl)pyridine-3,5-dicarbonitrile (26)

The product was prepared following the general procedure A starting from 3-fluorobenzaldehyde (0.05 mL, 0.5 mmol, 1 eq.) and was isolated as a white powder (34 mg, 0.06 mmol, 11% isolated yield). **^1^H NMR (600 MHz, CDCl_3_) δ** 7.55 (t, *J* = 7.9 Hz, 1H), 7.43 (d, *J* = 7.7 Hz, 1H), 7.40 – 7.34 (m, 2H), 5.38 (s, 2H), 3.83 – 3.79 (m, 4H), 1.74 – 1.67 (m, 6H). **^13^C NMR (151 MHz, CDCl_3_) δ** 163.5, 161.0, 160.7 (d, *J* = 353.3 Hz), 159.4, 136.9 (d, *J* = 7.0 Hz), 130.8 (d, *J* = 8.3 Hz), 124.6 (d, *J* = 3.1 Hz), 117.6 (d, *J* = 20.9 Hz), 117.4, 116.3, 116.1 (d, *J* = 23.1 Hz), 83.6, 81.7, 49.3, 26.1, 24.5. **MS (ESI^+^) calculated for C_18_H_16_N_5_F:** [M+H]^+^ *m/z* = 322.2, found *m/z* = 322.7. **HRMS (ESI^+^) calculated for C_18_H_16_N_5_F:** [M+H]^+^ *m/z* = 322.1463, found *m/z* = 322.1478.

#### 2-amino-4-(3-bromophenyl)-6-(piperidin-1-yl)pyridine-3,5-dicarbonitrile (27)

**(small batch)** The product was prepared following the general procedure B starting from 3-bromobenzaldehyde (0.06 mL mg, 0.5 mmol, 1 eq.) and was isolated as a white powder (41 mg, 0.11 mmol, 21% isolated yield).

**(large batch)** The product was prepared following the general procedure C starting from 3-bromobenzaldehyde (5.83 g, 31.5 mmol, 1.05 eq.) at 3 M in MeOH and piperidine (0.15 mL, 0.05 eq.) and the intermediate was isolated as a white powder (6.76 g, 29 mmol, 97% isolated yield). The second step was performed twice on (3-bromobenylidene)malononitrile (3.50 g & 3.21 g, 15 mmol & 13.8 mmol, 1 eq.) and piperidine (1.55 mL & 1.34 mL, 1.05 eq.) and gave a white powder (1.05 g & 1.03 g, 2.7 mmol & 2.7 mmol, 18% & 20% isolated yield, 17% & 19% overall yield).

**^1^H NMR (600 MHz, CDCl_3_) δ** 7.65 (dt, *J* = 7.3, 1.9 Hz, 1H), 7.61 (t, *J* = 1.8 Hz, 1H), 7.43 – 7.36 (m, 2H), 5.50 (s, 2H), 3.84 – 3.79 (m, 4H), 1.76 – 1.68 (m, 6H). **^13^C NMR (151 MHz, CDCl_3_) δ** 160.9, 160.5, 159.3, 136.8, 133.7, 131.6, 130.6, 127.4, 122.9, 117.3, 116.1, 83.6, 81.8, 49.5, 26.1, 24.5. **MS (ESI^+^) calculated for C_18_H_16_N_5_Br:** [M+H]^+^ *m/z* = 382.1, 384.1, found *m/z* = 382.1, 384.1. **HRMS (ESI^+^) calculated for C_18_H_16_N_5_Br:** [M+H]^+^ *m/z* = 382.0662, found *m/z* = 382.0667.

#### 4-([1,1’-biphenyl]-3-yl)-2-amino-6-(piperidin-1-yl)pyridine-3,5-dicarbonitrile (28)

**(small batch)** The product was prepared following the general procedure A starting from [1,1’-biphenyl]-3-carbaldehyde (0.080 mL, 0.5 mmol, 1 eq.) and was isolated as a white powder (13 mg, 0.03 mmol, 7% isolated yield).

**(large batch)** The product was prepared following the general procedure E at 100°C for 30 min to 1 h (depending on the catalyst’s batch), starting from 2-amino-4-(3-bromophenyl)-6-(piperidin-1-yl)pyridine-3,5-dicarbonitrile (400 mg, 1.05 mmol, 1 eq.) and phenylboronic acid monohydrate (438 mg, 3 eq.). The crude’s treatment was performed as followed: (1) dilution of the medium by addition of H_2_O (20 mL); (2) extraction of the crude by EtOAc (3×15 mL); (3) organic phases were dried with MgSO_4_, filtered on a celite pad then concentrated under reduced pressure; (4) the crude was stirred at r.t. in a minimum of Et_2_O for 10 min then filtered; the step (4) was repeated until pure. The product was isolated as a white powder (290 mg, 0.15 mmol, 73% isolated yield).

**^1^H NMR (600 MHz, CDCl_3_) δ** 7.73 – 7.70 (m, 2H), 7.66 – 7.62 (m, 2H), 7.58 (t, *J* = 7.7 Hz, 1H), 7.49 (dt, *J* = 7.7, 1.5 Hz, 1H), 7.45 (t, *J* = 7.7 Hz, 2H), 7.37 (tt, *J* = 7.4, 1.3 Hz, 1H), 5.49 (s, 2H), 3.86 – 3.80 (m, 4H), 1.73 – 1.70 (m, 6H). **^13^C NMR (151 MHz, CDCl_3_) δ** 162.4, 161.1, 159.5, 142.0, 140.6, 135.2, 129.5, 129.4, 129.0, 127.9, 127.8, 127.7, 127.6, 117.8, 116.6, 83.8, 81.9, 49.5, 26.2, 24.5. **MS (ESI^+^) calculated for C_24_H_21_N_5_:** [M+H]^+^ *m/z* = 380.2, found *m/z* = 380.1. **HRMS (ESI^+^) calculated for C_24_H_21_N_5_:** [M+H]^+^ *m/z* = 380.1870, found *m/z* = 380.1859.

#### 2-amino-4-(3-nitrophenyl)-6-(piperidin-1-yl)pyridine-3,5-dicarbonitrile (29)

The product was prepared following the general procedure A starting from 3-nitrobenzaldehyde (75 mg, 0.5 mmol, 1 eq.) and was isolated as a white powder (28 mg, 0.08 mmol, 16% isolated yield). **^1^H NMR (600 MHz, CDCl_3_) δ** 8.39 (ddd, *J* = 8.2, 2.3, 1.1 Hz, 1H), 8.37 (t, *J* = 2.0 Hz, 1H), 7.81 (dt, *J* = 7.7, 1.4 Hz, 1H), 7.73 (t, *J* = 7.9 Hz, 1H), 5.48 (s, 2H), 3.86 – 3.82 (m, 4H), 1.77 – 1.68 (m, 6H). **^13^C NMR (151 MHz, CDCl_3_) δ** 160.4, 159.8, 159.3, 148.4, 136.5, 134.8, 130.3, 125.3, 124.2, 117.2, 115.9, 83.3, 81.6, 49.3, 26.2, 24.5. **MS (ESI^+^) calculated for C_18_H_17_N_5_O:** [M+H]^+^ *m/z* = 349.1, found *m/z* = 349.4. **HRMS (ESI^+^) calculated for C_18_H_17_N_5_O:** [M+H]^+^ *m/z* = 349.1408, found *m/z* = 349.1392.

#### 2-amino-4-(3-cyanophenyl)-6-(piperidin-1-yl)pyridine-3,5-dicarbonitrile (30)

The product was prepared following the general procedure A starting from 3-cyanobenzaldehyde (66 mg, 0.5 mmol, 1 eq.) and was isolated as a light orange solid (24 mg, 0.04 mmol, 7% isolated yield). **^1^H NMR (600 MHz, CDCl_3_) δ** 7.74 (dt, *J* = 7.7, 1.4 Hz, 1H), 7.69 (t, *J* = 1.7 Hz, 1H), 7.64 (dt, *J* = 7.8, 1.5 Hz, 2H), 7.58 (t, *J* = 7.7 Hz, 1H), 5.34 (s, 2H), 3.78 – 3.73 (m, 4H), 1.69 – 1.60 (m, 6H). **^13^C NMR (151 MHz, CDCl_3_) δ** 160.6, 159.9, 159.3, 136.3, 133.9, 133.1, 132.3, 130.0, 118.0, 117.3, 116.0, 113.5, 111.6, 83.2, 81.5, 49.3, 26.1, 24.5. **MS (ESI^+^) calculated for C_19_H_16_N_6_:** [M+H]^+^ *m/z* = 329.2, found *m/z* = 329.4. **HRMS (ESI^+^) calculated for C_19_H_16_N_6_:** [M+Na]^+^ *m/z* = 351.1329, found *m/z* = 351.1334.

#### 2-amino-4-(3-ethynylphenyl)-6-(piperidin-1-yl)pyridine-3,5-dicarbonitrile (31)

This synthesis is a two-step procedure starting by a Sonogashira cross-coupling and followed by a silyl deprotection. **(a)** In a 5 mL sealed tube purged with argon and equipped with a magnetic stirrer, anhydrous THF (1.5mL) was bubbled with argon for 5-10 min. 2-amino-4-(3-bromophenyl)-6-(piperidin-1-yl)pyridine-3,5-dicarbonitrile (57 mg, 0.15 mmol, 1 eq.), copper iodide (1.4 mg, 0.05 eq.) and Pd(dppf)Cl_2_ (5.5 mg, 0.05 eq.) were loaded and the vessel purged with argon. Dipropylamine (0.08 mL, 4 eq.) and triisopropylsilylacetylene (0.04 mL, 1.2 eq.) were added with a syringe. The reaction was sealed and heated to 80°C for 18 h. Completion was assessed by TLC (Rf = 0.25 in cyclo:EtOAc 8:2). The palladium and salts were removed by filtration on a celite pad, washed with EtOAc (3 × 5 mL) and all solvents were removed under reduced pressure. The crude intermediate was purified by flash chromatography on a 12 g silica cartridge, solid deposit, with a cyclo:EtOAc gradient (95:5→8:2, 12 CV) and isolated as a white powder (38 mg, 0.08 mmol, 52% isolated yield). **(b)** In a 5 mL round bottom flask with a magnetic stirrer, the intermediate (20 mg, 0.04 mmol, 1 eq.) was solubilized in anhydrous THF (1.5 mL). The mixture was cooled to 0°C then a TBAF solution (1 M in THF, 0.08 mL, 2 eq.) was slowly added. The reaction was left stirring for 15 min at 0°C. Completion was assessed by TLC (Rf = 0.15 in cyclo:EtOAc 8:2). The reaction was quenched by addition of water (10 mL) and brine (5 mL). The crude product was extracted by EtOAc (3 × 10 mL), dried over MgSO_4_, filtered, isolated under reduced pressure, and used as such (19 mg, 0.04 mmol, quantitative isolated yield).). **^1^H NMR (600 MHz, CDCl_3_) δ** 7.62 (dt, *J* = 7.4, 1.7 Hz, 1H), 7.59 (d, *J* = 1.7 Hz, 1H), 7.48 (t, *J* = 7.6 Hz, 1H), 7.45 (dt, *J* = 7.8, 1.8 Hz, 1H), 5.36 (s, 2H), 3.86 – 3.76 (m, 4H), 3.12 (s, 1H), 1.75 – 1.68 (m, 6H). **^13^C NMR (151 MHz, CDCl_3_) δ** 161.3, 161.0, 159.4, 135.3, 134.1, 132.3, 129.07, 129.05, 123.2, 117.5, 116.3, 83.6, 81.7, 78.6, 49.3, 26.2, 24.5. **MS (ESI^+^) calculated for C_20_H_17_N_5_:** [M+H]^+^ *m/z* = 328.2, found *m/z* = 328.1. **HRMS (ESI^+^) calculated for C_20_H_17_N_5_:** [M+H]^+^ *m/z* = 328.1557, found *m/z* = 328.1543.

#### 2-amino-4-(2,6-dimethylphenyl)-6-(piperidin-1-yl)pyridine-3,5-dicarbonitrile (32)

The dimethylbenzaldehyde (67 mg, 0.5 mmol, 1 eq.) and was isolated as a white powder (35 mg, 11 mmol, 21% isolated yield). **^1^H NMR (600 MHz, CDCl_3_) δ** 7.26 (t, *J* = 7.6 Hz, 1H), 7.14 (d, *J* = 7.6 Hz, 2H), 5.69 (s, 2H), 3.87 – 3.82 (m, 4H), 2.16 (s, 6H), 1.74 – 1.71 (m, 6H). **^13^C NMR (151 MHz, CDCl_3_) δ** 163.6, 159.1, 134.7, 134.4, 129.7, 128.2, 116.4, 115.3, 84.3, 82.5, 49.5, 26.1, 24.4, 19.8. **MS (ESI^+^) calculated for C_20_H_21_N_5_:** [M+H]^+^ *m/z* = 332.2, found *m/z* = 332.1. **HRMS (ESI^+^) calculated for C_20_H_21_N_5_:** [M+Na]^+^ *m/z* = 354.1690, found *m/z* = 354.1679.

#### 2-amino-4-(3,5-dimethylphenyl)-6-(piperidin-1-yl)pyridine-3,5-dicarbonitrile (33)

The product was prepared following the general procedure A starting from 3,5-dimethylbenzaldehyde (0.07 mL, 0.5 mmol, 1 eq.) and was isolated as a white powder (64 mg, 0.19 mmol, 39% isolated yield). **^1^H NMR (600 MHz, CDCl_3_) δ** 7.13 (s, 1H), 7.07 (s, 2H), 5.62 (s, 2H), 3.83 – 3.79 (m, 4H), 2.38 (s, 6H), 1.76 – 1.67 (m, 6H). **^13^C NMR (151 MHz, CDCl_3_) δ** 163.2, 159.2, 138.5, 134.7, 132.5, 126.4, 117.6, 116.4, 83.9, 82.0, 49.7, 26.1, 24.4, 21.5. **MS (ESI^+^) calculated for C_20_H_21_N_5_:** [M+H]^+^ *m/z* = 332.2, found *m/z* = 332.1. **HRMS (ESI^+^) calculated for C_20_H_21_N_5_:** [M+Na]^+^ *m/z* = 354.1690, found *m/z* = 354.1679.

#### 2-amino-4-(3-chloro-2-fluorophenyl)-6-(piperidin-1-yl)pyridine-3,5-dicarbonitrile (34)

The product was prepared following the general procedure C starting from 3-chloro-2-fluorobenzaldehyde (79 mg, 0.5 mmol, 1 eq.) and piperidine (0.06 mL, 1.2 eq.) and isolated as a white powder (39 mg, 0.11 mmol, 22% isolated yield). **^1^H NMR (600 MHz, CDCl_3_) δ** 7.58 – 7.52 (m, 1H), 7.30 – 7.26 (m, 1H), 7.24 (t, *J* = 7.8 Hz, 1H), 5.37 (s, 2H), 3.86 – 3.77 (m, 4H), 1.76 – 1.65 (m, 6H). **^13^C NMR (151 MHz, CDCl_3_) δ** 160.3, 159.2, 155.6, 153.9 (d, *J* = 252.2 Hz), 133.0, 128.8 (d, *J* = 1.4 Hz), 125.2 (d, *J* = 4.7 Hz), 124.4 (d, *J* = 15.3 Hz), 122.5 (d, *J* = 17.8 Hz), 116.9, 115.8, 84.4, 82.3, 49.2, 26.1, 24.5. **MS (ESI^+^) calculated for C_18_H_15_N_5_ClF:** [M+H]^+^ *m/z* = 356.1, found *m/z* = 356.1. **HRMS (ESI^+^) calculated for C_18_H_15_N_5_ClF:** [M+H]^+^ *m/z* = 356.1073, found *m/z* = 356.1070.

#### 2-amino-4-(3-chloro-4-fluorophenyl)-6-(piperidin-1-yl)pyridine-3,5-dicarbonitrile (35)

The product was prepared following the general procedure C starting from 3-chloro-4-fluorobenzaldehyde (317 mg, 2 mmol, 1 eq.) and piperidine (0.24 mL, 1.2 eq.) and isolated as a white powder (125 mg, 0.35 mmol, 18% isolated yield). **^1^H NMR (600 MHz, CDCl_3_) δ** 7.55 (dd, *J* = 6.8, 2.2 Hz, 1H), 7.37 (ddd, *J* = 8.4, 4.4, 2.2 Hz, 1H), 7.29 (t, *J* = 8.6 Hz, 1H), 5.50 (s, 2H), 3.84 – 3.79 (m, 4H), 1.75 – 1.67 (m, 6H). **^13^C NMR (151 MHz, CDCl_3_) δ** 160.5, 160.1, 159.5 (d, *J* = 253.6 Hz), 159.3, 131.9 (d, *J* = 4.2 Hz), 131.3, 129.0 (d, *J* = 7.8 Hz), 122.1 (d, *J* **calculated for C_18_H_15_N_5_FCl:** [M+H]^+^ *m/z* = 356.1, found *m/z* = 356.3. **HRMS (ESI^+^) calculated for C_18_H_15_N_5_ClF:** [M+Na]^+^ *m/z* = 378.0893, found *m/z* = 378.0892.

#### 2-amino-4-(3-chloro-5-fluorophenyl)-6-(piperidin-1-yl)pyridine-3,5-dicarbonitrile (36)

The product was prepared following the general procedure C starting from 3-chloro-5-fluorobenzaldehyde (317 mg, 2 mmol, 1 eq.) and piperidine (0.24 mL, 1.2 eq.) and isolated as a white powder (136 mg, 0.38 mmol, 19% isolated yield). **^1^H NMR (600 MHz, CDCl_3_) δ** 7.26 – 7.22 (m, 2H), 7.09 (dt, *J* = 8.1, 1.7 Hz, 1H), 5.51 (s, 2H), 3.84 – 3.80 (m, 4H), 1.73 – 1.67 (m, 6H). **^13^C NMR (151 MHz, CDCl_3_) δ** 162.6 (d, *J* = 252.3 Hz), 160.3, 159.7 (d, *J* = 2.3 Hz), 159.2, 137.7 (d, *J* = 8.9 Hz), 136.0 (d, *J* = 10.7 Hz), 124.9 (d, *J* = 3.1 Hz), 118.3, 117.0, 115.8, 114.7 (d, *J* = 23.0 Hz), 83.3, 81.6, 49.4, 26.1, 24.4. **MS (ESI^+^) calculated for C_18_H_15_N_5_FCl:** [M+H]^+^ *m/z* = 356.1, found *m/z* = 356.4. **HRMS (ESI^+^) calculated for C_18_H_15_N_5_ClF:** [M+H]^+^ *m/z* = 356.1073, found *m/z* = 356.1070.

#### 2-amino-4-(5-chloro-2-fluorophenyl)-6-(piperidin-1-yl)pyridine-3,5-dicarbonitrile (37)

The product was prepared following the general procedure C starting from 5-chloro-2-fluorobenzaldehyde (79 mg, 0.5 mmol, 1 eq.) and piperidine (0.06 mL, 1.2 eq.) and isolated as a white powder (35 mg, 0.10 mmol, 20% isolated yield). **^1^H NMR (600 MHz, CDCl_3_) δ** 7.45 (ddd, *J* = 8.8, 4.4, 2.7 Hz, 1H), 7.35 (dd, *J* = 6.0, 2.6 Hz, 1H), 7.19 (t, *J* = 8.9 Hz, 1H), 5.36 (s, 2H), 3.85 – 3.79 (m, 4H), 1.75 – 1.66 (m, 6H). **^13^C NMR (151 MHz, CDCl_3_) δ** 160.3, 159.2, 157.7 (d, *J* = 250.3 Hz), 155.2, 132.4 (d, *J* = 8.4 Hz), 130.3 (d, *J* = 2.5 Hz), 129.9 (d, *J* = 3.6 Hz), 124.4 (d, *J* = 17.2 Hz), 118.1 (d, *J* = 23.2 Hz), 116.9, 115.7, 84.3, 82.3, 49.2, 26.1, 24.5. **MS (ESI^+^) calculated for C_18_H_15_N_5_ClF:** [M+H]^+^ *m/z* = 356.1, found *m/z* = 356.1. **HMRS (ESI^+^) calculated for C_18_H_15_N_5_ClF:** [M+H]^+^ *m/z* = 356.1073, found *m/z* = 356.1070.

#### 2-amino-4-(2’-fluoro-[1,1’-biphenyl]-3-yl)-6-(piperidin-1-yl)pyridine-3,5-dicarbonitrile (38)

The product was prepared following the general procedure A starting from 2’-fluoro-[1,1’-biphenyl]-3-carbaldehyde (100 mg, 0.5 mmol, 1 eq.) and was isolated as a white powder (45 mg, 0.11 mmol, 23% isolated yield). **^1^H NMR (600 MHz, CDCl_3_) δ** 7.72 – 7.67 (m, 2H), 7.59 (t, *J* = 7.6 Hz, 1H), 7.51 (dt, *J* = 7.7, 1.5 Hz, 1H), 7.44 – 7.39 (m, 2H), 7.35 – 7.31 (m, 1H), 7.09 – 7.03 (m, 1H), 5.38 (s, 2H), 3.88 – 3.80 (m, 4H), 1.78 – 1.63 (m, 6H). **^13^C NMR (151 MHz, CDCl_3_) δ** 163.3 (d, *J* = 246.0 Hz), 162.1, 161.3, 159.6, 142.8 (d, *J* = 7.5 Hz), 140.7 (d, *J* = 2.1 Hz), 135.5, 130.5 (d, *J* = 8.4 Hz), 129.5 (d, *J* = 26.6 Hz), 128.2, 127.9, 123.3, 117.9, 116.7, 114.6 (d, *J* = 21.0 Hz), 114.4 (d, *J* = 22.0 Hz), 83.7, 81.8, 49.4, 26.2, 24.6. **MS (ESI^+^) calculated for C_24_H_20_N_5_F:** [M+H]^+^ *m/z* = 398.2, found *m/z* = 398.8. **HRMS (ESI^+^) calculated**

#### 2-amino-4-(3’-fluoro-[1,1’-biphenyl]-3-yl)-6-(piperidin-1-yl)pyridine-3,5-dicarbonitrile (39)

The product was prepared following the general procedure A starting from 3’-fluoro-[1,1’-biphenyl]-3-carbaldehyde (100 mg, 0.5 mmol, 1 eq.) and was isolated as a white powder (34 mg, 0.08 mmol, 17% isolated yield). **^1^H NMR (600 MHz, CDCl_3_) δ** 7.71 (dq, *J* = 7.7, 1.6 Hz, 1H), 7.68 (q, *J* = 1.6 Hz, 1H), 7.58 (d, *J* = 7.7 Hz, 1H), 7.55 – 7.49 (m, 2H), 7.37 – 7.30 (m, 1H), 7.23 (td, *J* = 7.5, 1.2 Hz, 1H), 7.17 (ddd, *J* = 10.9, 8.2, 1.2 Hz, 1H), 5.37 (s, 2H), 3.83 – 3.79 (m, 4H), 1.71 (t, *J* = 3.1 Hz, 6H). **^13^C NMR (151 MHz, CDCl_3_) δ** 162.0, 161.4, 159.9 (d, *J* = 248.7 Hz), 159.6, 136.5, 135.1, 131.3 (d, *J* = 3.3 Hz), 131.1 (d, *J* = 3.2 Hz), 129.59 (d, *J* = 1.1 Hz), 129.55 (d, *J* = 4.0 Hz), 129.1, 128.1, 124.7 (d, *J* = 3.6 Hz), 117.8, 116.7, 116.3 (d, *J* = 22.5 Hz), 83.8, 81.8, 49.4, 26.1, 24.6. **MS (ESI^+^) calculated for C_24_H_20_N_5_F:** [M+H]^+^ *m/z* = 398.2, found *m/z* = 398.8. **HRMS (ESI^+^) calculated for C_24_H_20_N_5_F:** [M+Na]^+^ *m/z* = 420.1595, found *m/z* = 420.1594.

#### 2-amino-4-(4’-fluoro-[1,1’-biphenyl]-3-yl)-6-(piperidin-1-yl)pyridine-3,5-dicarbonitrile (40)

The product was prepared following the general procedure A starting from 4’-fluoro-[1,1’-biphenyl]-3-carbaldehyde (100 mg, 0.5 mmol, 1 eq.) and was isolated as a white powder (34 mg, 0.08 mmol, 17% isolated yield). **^1^H NMR (600 MHz, CDCl_3_) δ** 7.68 – 7.64 (m, 2H), 7.61 – 7.55 (m, 3H), 7.48 (dt, *J* = 7.7, 1.5 Hz, 1H), 7.18 – 7.11 (m, 2H), 5.37 (s, 2H), 3.84 – 3.79 (m, 4H), 1.76 – 1.70 (m, 6H). **^13^C NMR (151 MHz, CDCl_3_) δ** 162.9 (d, *J* = 246.7 Hz), 162.2, 161.3, 159.6, 141.0, 136.8 (d, *J* = 3.4 Hz), 135.4, 129.5, 129.3, 129.2 (d, *J* = 8.2 Hz), 127.8 (d, *J* = 19.2 Hz), 117.9, 116.8, 115.9 (d, *J* = 21.5 Hz), 83.7, 81.8, 49.4, 26.2, 24.6. **MS (ESI^+^) calculated for C_24_H_20_N_5_F:** [M+H]^+^ *m/z* = 398.2, found *m/z* = 398.8. **HRMS (ESI^+^) calculated for C_24_H_20_N_5_F:** [M+Na]^+^ *m/z* = 420.1595, found *m/z* = 420.1594.

#### 2-amino-4-(2’-methyl-[1,1’-biphenyl]-3-yl)-6-(piperidin-1-yl)pyridine-3,5-dicarbonitrile (41)

The product was prepared following the general procedure D starting from 2-amino-4-(3-bromophenyl)-6-(piperidin-1-yl)pyridine-3,5-dicarbonitrile (60 mg, 0.16 mmol, 1 eq.) and 2-methylphenylboronic acid (63 mg, 3 eq.) and was isolated as a white powder (30 mg, 0.08 mmol, 49% isolated yield). **^1^H NMR (600 MHz, CDCl_3_) δ** 7.55 (t, *J* = 7.7 Hz, 1H), 7.47 (ddt, *J* = 7.4, 5.8, 1.4 Hz, 2H), 7.43 (t, *J* = 1.8 Hz, 1H), 7.32 – 7.25 (m, 2H), 5.35 (s, 2H), 3.83 – 3.76 (m, 4H), 2.32 (s, 3H), 1.86 – 1.66 (m, 6H). **^13^C NMR (151 MHz, CDCl_3_) δ** 162.3, 161.3, 159.6, 142.7, 141.1, 135.8, 134.8, 131.4, 130.4, 130.1, 129.7, 128.7, 127.7, 127.3, 126.0, 117.8, 116.7, 83.9, 81.9, 49.4, 26.1, 24.6, 20.7. **MS (ESI^+^) calculated for C_24_H_21_N_5_:** [M+H]^+^ *m/z* = 394.2, found *m/z* = 394.9. **HRMS (ESI^+^) calculated for C_24_H_21_N_5_:** [M+Na]^+^ *m/z* = 416.1846, found

#### 2-amino-4-(3’-methyl-[1,1’-biphenyl]-3-yl)-6-(piperidin-1-yl)pyridine-3,5-dicarbonitrile (42)

The product was prepared following the general procedure D starting from 2-amino-4-(3-bromophenyl)-6-(piperidin-1-yl)pyridine-3,5-dicarbonitrile (60 mg, 0.16 mmol, 1 eq.) and 3-methylphenylboronic acid (63 mg, 3 eq.) and was isolated as a white powder (47 mg, 0.12 mmol, 76% isolated yield). **^1^H NMR (600 MHz, CDCl_3_) δ** 7.70 (ddd, *J* = 7.7, 1.9, 1.2 Hz, 1H), 7.69 (t, *J* = 1.8 Hz, 1H), 7.57 (t, *J* = 7.7 Hz, 1H), 7.47 – 7.46 (m, 1H), 7.43 (d, *J* = 8.2 Hz, 1H), 7.34 (t, *J* = 7.9 Hz, 1H), 7.18 (d, *J* = 7.2 Hz, 1H), 5.36 (s, 2H), 3.82 (m, 4H), 2.43 (s, 3H), 1.72 (m, 6H). **^13^C NMR (151 MHz, CDCl_3_) δ** 162.4, 161.4, 159.6, 142.2, 140.6, 138.5, 135.2, 129.3, 128.9, 128.5, 128.3, 127.9, 127.5, 124.7, 117.9, 116.8, 83.8, 81.9, 49.4, 26.2, 24.6, 21.7. **MS (ESI^+^) calculated for C_24_H_21_N_5_:** [M+H]^+^ *m/z* = 394.2, found *m/z* = 394.9. **HRMS (ESI^+^) calculated for C_24_H_21_N_5_:** [M+Na]^+^ *m/z* = 416.1846, found *m/z* = 416.1857.

#### 2-amino-4-(4’-methyl-[1,1’-biphenyl]-3-yl)-6-(piperidin-1-yl)pyridine-3,5-dicarbonitrile (43)

The product was prepared following the general procedure A starting from 4’-methyl-[1,1’-biphenyl]-3-carbaldehyde (98 mg, 0.5 mmol, 1 eq.) and was isolated as a white powder (56 mg, 0.14 mmol, 28% isolated yield). **^1^H NMR (600 MHz, CDCl_3_) δ** 7.71 – 7.67 (m, 2H), 7.56 (t, *J* = 7.6 Hz, 1H), 7.53 (d, *J* = 7.9 Hz, 2H), 7.45 (dd, *J* = 7.7, 1.5 Hz, 1H), 7.26 (d, *J* = 8.1 Hz, 2H), 5.37 (s, 2H), 3.84 – 3.79 (m, 4H), 1.74 – 1.69 (m, 6H). **^13^C NMR (151 MHz, CDCl_3_) δ** 162.4, 161.4, 159.6, 141.9, 137.8, 137.6, 135.2, 129.7, 129.31, 129.28, 127.7, 127.40, 127.36, 117.9, 116.8, 83.8, 81.8, 49.4, 26.2, 24.6, 21.3. **MS (ESI^+^) calculated for C_24_H_21_N_5_:** [M+H]^+^ *m/z* = 394.2, found *m/z* = 394.8. **HRMS (ESI^+^) calculated for C_24_H_21_N_5_:** [M+Na]^+^ *m/z* = 416.1846, found *m/z* = 416.1857.

#### 2-amino-6-(piperidin-1-yl)-4-(3-(pyridin-2-yl)phenyl)pyridine-3,5-dicarbonitrile (44)

The product was prepared following the general procedure F starting from (3-(2-amino-3,5-dicyano-6-(piperidin-1-yl)pyridin-4-yl)phenyl)boronic acid pinacol ester (64 mg, 0.15 mmol, 1 eq.) and 2-bromopyridine (28 mg, 1.2 eq.) and was isolated as a white powder (17 mg, 0.04 mmol, 30% isolated yield). **^1^H NMR (600 MHz, CDCl_3_) δ** 8.92 (s, 1H), 8.65 (s, 1H), 7.99 (dt, *J* = 8.0, 1.8 Hz, 1H), 7.75 – 7.68 (m, 2H), 7.64 (t, *J* = 7.7 Hz, 1H), 7.56 (dt, *J* = 7.7, 1.4 Hz, 1H), 7.44 (t, *J* = 5.9 Hz, 1H), 5.40 (s, 2H), 3.86 – 3.80 (m, 4H), 1.76 – 1.70 (m, 6H). **^13^C NMR (151 MHz, CDCl_3_) δ** 162.0, 161.1, 159.4, 135.4, 129.39, 129.36, 129.0, 127.5, 122.7, 121.1, 117.6, 116.5, 83.7, 81.7, 49.2, 26.0, 24.4. **MS (ESI^+^) calculated for C_23_H_20_N_6_:** [M+H]^+^ *m/z* = 381.2, found *m/z* = 381.0. **HRMS (ESI^+^) calculated for C_23_H_20_N_6_:** [M+H]^+^ *m/z* = 381.1823, found *m/z* = 381.1803.

#### 2-amino-6-(piperidin-1-yl)-4-(3-(pyridin-3-yl)phenyl)pyridine-3,5-dicarbonitrile (45)

The product was prepared following the general procedure E at 100°C for 30 min, starting from 2-amino-4-(3-bromophenyl)-6-(piperidin-1-yl)pyridine-3,5-dicarbonitrile (58 mg, 0.15 mmol, 1 eq.) and 3-pyridineboronic acid (55 mg, 3 eq.) and was isolated as a white powder (79 mg, 0.15 mmol, quantitative isolated yield). **^1^H NMR (600 MHz, CDCl_3_) δ** 8.92 (s, 1H), 8.65 (s, 1H), 7.99 (dt, *J* = 8.0, 1.8 Hz, 1H), 7.75 – 7.68 (m, 2H), 7.64 (t, *J* = 7.7 Hz, 1H), 7.56 (dt, *J* = 7.7, 1.4 Hz, 1H), 7.44 (t, *J* = 5.9 Hz, 1H), 5.40 (s, 2H), 3.86 – 3.80 (m, 4H), 1.76 – 1.70 (m, 6H). **^13^C NMR (151 MHz, CDCl_3_) δ** 161.9, 161.2, 159.6, 149.1, 148.6, 138.7, 136.1, 135.7, 134.9, 129.8, 129.4, 128.5, 127.9, 123.8, 117.9, 116.7, 83.6, 81.8, 49.4, 26.2, 24.6. **MS (ESI^+^) calculated for C_23_H_20_N_6_:** [M+H]^+^ *m/z* = 381.2, found *m/z* = 381.1. **HRMS (ESI^+^) calculated for C_23_H_20_N_6_:** [M+H]^+^ *m/z* = 381.1823, found *m/z* = 381.1836.

#### 2-amino-6-(piperidin-1-yl)-4-(3-(pyridin-4-yl)phenyl)pyridine-3,5-dicarbonitrile (46)

The product was prepared following the general procedure E at 80°C for 15 min, starting from 2-amino-4-(3-bromophenyl)-6-(piperidin-1-yl)pyridine-3,5-dicarbonitrile (58 mg, 0.15 mmol, 1 eq.) and 4-pyridineboronic acid (55 mg, 3 eq.) and was isolated as a white powder (5 mg, 0.013 mmol, 9% isolated yield). **^1^H NMR (600 MHz, CDCl_3_) δ** 8.71 – 8.67 (m, 2H), 7.79 – 7.74 (m, 2H), 7.65 (t, *J* = 8.1 Hz, 1H), 7.61 – 7.58 (m, 3H), 5.41 (s, 2H), 3.86 – 3.80 (m, 4H), 1.75 – 1.69 (m, 6H). **^13^C NMR (151 MHz, CDCl_3_) δ** 161.7, 161.2, 159.6, 149.3, 138.6, 136.0, 132.4, 132.3, 130.1, 129.4, 128.8, 128.7, 128.1, 122.6, 117.9, 116.8, 83.6, 81.8, 49.5, 26.3, 24.6. **MS (ESI^+^) calculated for C_23_H_20_N_6_:** [M+H]^+^ *m/z* = 381.2, found *m/z* = 381.1. **HRMS (ESI^+^) calculated for C_23_H_20_N_6_:** [M+H]^+^ *m/z* = 381.1823, found *m/z* = 381.1836.

#### 2-amino-6-(piperidin-1-yl)-4-(3-(thiophen-2-yl)phenyl)pyridine-3,5-dicarbonitrile (47)

The product was prepared following the general procedure E at 100°C for 30 min, starting from 2-amino-4-(3-bromophenyl)-6-(piperidin-1-yl)pyridine-3,5-dicarbonitrile (58 mg, 0.16 mmol, 1 eq.) and 2-thiopheneboronic acid (23 mg, 3 eq.) and was isolated as a white powder (58 mg, 0.16 mmol, quantitative isolated yield). **^1^H NMR (600 MHz, CDCl_3_) δ** 7.74 (ddd, *J* = 7.8, 1.9, 1.1 Hz, 1H), 7.71 (t, *J* = 1.8 Hz, 1H), 7.52 (t, *J* = 7.8 Hz, 1H), 7.39 (ddd, *J* = 7.7, 1.8, 1.1 Hz, 1H), 7.37 (dd, *J* = 3.6, 1.2 Hz, 1H), 7.31 (dd, *J* = 5.1, 1.2 Hz, 1H), 7.09 (dd, *J* = 5.1, 3.6 Hz, 1H), 5.44 (s, 2H), 3.85 – 3.80 (m, 4H), 1.75 – 1.69 (m, 6H). **^13^C NMR (151 MHz, CDCl_3_) δ** 162.0, 161.0, 159.4, 143.4, 135.5, 135.2, 129.6, 128.25, 128.16, 127.7, 126.3, 125.6, 124.1, 117.6, 116.5, 83.8, 81.9, 49.5, 26.2, 24.5. **MS (ESI^+^) calculated for C_22_H_19_N_5_S:** [M+H]^+^ *m/z* = 386.1, found *m/z* = 386.1. **HRMS (ESI^+^) calculated for C_22_H_19_N_5_S:** [M+H]^+^ *m/z* =

#### 2-amino-6-(piperidin-1-yl)-4-(3-(thiophen-3-yl)phenyl)pyridine-3,5-dicarbonitrile (48)

The product was prepared following the general procedure E at 80°C for 15 min, starting from 2-amino-4-(3-bromophenyl)-6-(piperidin-1-yl)pyridine-3,5-dicarbonitrile (58 mg, 0.16 mmol, 1 eq.) and 3-thiopheneboronic acid (23 mg, 1.2 eq.) and was isolated as a white powder (42 mg, 0.11 mmol, 73% isolated yield). **^1^H NMR (600 MHz, CDCl_3_) δ** 7.66 – 7.62 (m, 2H), 7.46 (t, *J* = 7.7 Hz, 1H), 7.44 (dd, *J* = 2.9, 1.4 Hz, 1H), 7.36 – 7.33 (m, 2H), 7.32 (dd, *J* = 5.0, 2.9 Hz, 1H), 5.33 (s, 2H), 3.76 – 3.72 (m, 4H), 1.69 – 1.62 (m, 6H). **^13^C NMR (151 MHz, CDCl_3_) δ** 162.2, 161.3, 159.6, 141.7, 136.6, 135.4, 129.4, 128.7, 127.5, 127.1, 126.59, 126.55, 121.4, 117.8, 116.7, 83.7, 81.8, 49.4, 26.1, 24.6. **MS (ESI^+^) calculated for C_22_H_19_N_5_S:** [M+Na]^+^ *m/z* = 408.1, found *m/z* = 408.0. **HRMS (ESI^+^) calculated for C_22_H_19_N_5_S:** [M+H]^+^ *m/z* = 386.1435, found *m/z* = 386.1438.

#### 4-(3-(1H-pyrazol-5-yl)phenyl)-2-amino-6-(piperidin-1-yl)pyridine-3,5-dicarbonitrile (49)

A Suzuki cross-coupling step was performed following the general procedure F starting from (3-(2-amino-3,5-dicyano-6-(piperidin-1-yl)pyridin-4-yl)phenyl)boronic acid pinacol ester (64 mg, 0.15 mmol, 1 eq.) and 5-bromo-1-(tetrahydro-2*H*-pyran-2-yl)-1*H*-pyrazole (42 mg, 1.2 eq.). The THP deprotection of the pyrazole was done in a dry 10 mL round-bottom flask flushed with argon. The intermediate (1 eq.) was solubilized in a 1:1 DCM:MeOH mixture (3 mL). *p*-toluenesulfonic acid (39 mg, 1.5 eq.) was added and the solution was stirred at r.t. for 20 h. Completion was assessed by TLC (Rf = 0.18 in toluene:acetone 8:2). Two successive purifications were performed by flash chromatography on a pre-packed 4 g silica cartridge with a (toluene:acetone 4:1):MeOH isocratic gradient (9:1, 12 CV) then a cyclo:DCM gradient (95:5, 3 CV; 95:5→0:10, 18 CV; methanol wash) (42 mg, 0.11 mmol, 76% overall isolated yield). **^1^H NMR (600 MHz, CDCl_3_) δ** 7.88 (dt, *J* = 7.8, 1.4 Hz, 1H), 7.86 (t, *J* = 1.8 Hz, 1H), 7.63 (s, 1H), 7.53 (t, *J* = 7.7 Hz, 1H), 7.44 (dt, *J* = 7.8, 1.3 Hz, 1H), 6.65 (s, 1H), 5.33 (s, 2H), 3.77 – 3.73 (m, 4H), 1.67 – 1.62 (m, 6H). **^13^C NMR (151 MHz, CDCl_3_) δ** 161.8, 161.1, 159.5, 135.8, 129.7, 129.2, 128.3, 126.7, 117.7, 116.6, 83.7, 81.8, 49.4, 26.2, 24.6. **MS (ESI^+^) calculated for C_21_H_19_N_7_:** [M+H]^+^ *m/z* = 370.2, found *m/z* = 370.3. **HRMS (ESI^+^) calculated for C_21_H_19_N_7_:** [M+H]^+^ *m/z* = 370.1775, found *m/z* = 370.1779.

#### 4-(3-(1,3,4-oxadiazol-2-yl)phenyl)-2-amino-6-(piperidin-1-yl)pyridine-3,5-dicarbonitrile (50)

In a 25 mL round-bottom flask equipped with a magnetic stirrer and a condenser, methyl 3-(2-amino-3,5-dicyano-6-(piperidin-1-yl)pyridin-4-yl)benzoate (54 mg, 0.15 mmol, 1 eq.) was suspended in anhydrous methanol (1.5 mL). Hydrazine monohydrate (37 mg, 5 eq.) was at 80°C for 16 h. Solvents were removed under reduced pressure and the crude product was rinsed with DCM:MeOH 7:3 (10 mL) and dried under reduced pressure. In a 25 mL round-bottom flask equipped with a magnetic stirrer and a condenser, the intermediate was suspended in an excess of triethyl orthoformate (1 mL). The flask was flushed with argon and heated at 100°C for 20 h. The reaction was quenched by addition of water (20 mL) and the crude product was extracted by EtOAc (3 × 15 mL). The organic phases were combined, washed with brine, dried over MgSO_4_, filtered and the solvent was removed under reduced pressure. The crude product was purified by flash chromatography on a 12 g silica cartridge, solid deposit, with a cyclo:EtOAc gradient (8:2, 3 CV; 8:2→6:4, 12 CV; 6:4, 6 CV) and isolated as a white powder (32 mg, 0.09 mmol, 58% overall isolated yield). **^1^H NMR (600 MHz, CDCl_3_) δ** 8.50 (s, 1H), 8.26 (dt, *J* = 7.6, 1.6 Hz, 1H), 8.21 (t, *J* = 1.8 Hz, 1H), 7.70 (t, *J* = 7.8 Hz, 1H), 7.67 (dt, *J* = 7.8, 1.6 Hz, 1H), 5.66 (s, 2H), 3.87 – 3.82 (m, 4H), 1.76 – 1.70 (m, 6H). **^13^C NMR (151 MHz, CDCl_3_) δ** 164.1, 161.2, 160.2, 159.2, 152.9, 136.0, 132.2, 130.1, 129.1, 127.5, 124.5, 117.2, 115.9, 83.6, 81.6, 49.6, 26.2, 24.4. **MS (ESI^+^) calculated for C_20_H_17_N_7_O:** [M+H]^+^ *m/z* = 372.2, found *m/z* = 372.4. **HRMS (ESI^+^) calculated for C_20_H_17_N_7_O:** [M+H]^+^ *m/z* = 372.1568, found *m/z* = 372.1565.

#### 4-(3-(1H-tetrazol-5-yl)phenyl)-2-amino-6-(piperidin-1-yl)pyridine-3,5-dicarbonitrile (51)

The product was prepared following the general procedure C starting from 3-(2-(tetrahydro-2*H*-pyran-2-yl)-2*H*-tetrazol-5-yl)benzaldehyde (60 mg, 0.23 mmol, 1 eq.) and purified by flash chromatography on a pre-packed 4 g silica cartridge with a cyclo:DCM gradient (95:5, 3 CV; 95:5→0:10, 18 CV; methanol wash). The THP deprotection of the tetrazole was performed in the presence of Dowex 50WX8 H^+^ (1 g of resin for 100 mg of compound), in a 98:2 EtOH:H_2_O mixture (1 mL) at 80°C for 16 h. The resin was filtered off and rinsed with warm EtOH (3 × 2 mL) and, after solvent evaporation, the product was isolated as a white powder (10 mg, 0.03 mmol, 11% overall isolated yield). **^1^H NMR (600 MHz, CD_3_OD) δ** 8.22 (dt, *J* = 7.8, 1.4 Hz, 1H), 8.19 (t, *J* = 1.8 Hz, 1H), 7.63 (t, *J* = 7.7 Hz, 1H), 7.51 (ddd, *J* = 7.6, 1.9, 1.2 Hz, 1H), 3.82 (dd, *J* = 6.4, 4.1 Hz, 3H), 1.78 – 1.67 (m, 7H). **^13^C NMR (151 MHz, CD_3_OD) δ** 163.6, 162.9, 162.0, 161.8, 137.6, 131.7, 130.2, 130.0, 129.1, 128.0, 118.9, 117.2, 83.4, 82.5, 50.2, 27.2, 25.6. **MS (ESI^-^) calculated for C_19_H_17_N_9_:** [M-H]^-^ *m/z* = 370.2, found *m/z* = 370.0. **HRMS (ESI^-^) calculated for C_19_H_17_N_9_:** [M-H]^-^ *m/z* = 370.1524, found *m/z* = 370.1500.

#### 2-amino-6-chloro-4-phenylpyridine-3,5-dicarbonitrile (52)

In a dry 25 mL round bottom malononitrile (0.66 g, 2 eq.) were solubilized in pyridine (0.40 mL, 1 eq.). The reaction was stirred under argon at 100°C for 1 h and monitored by TLC until complete consumption of starting orthoformate. The resulting dark red mixture was cooled in an ice bath and HCl 37% was added dropwise under vigorous stirring (/!\ fumes). The mixture was heated to 80°C for 2 h. The crude product was cooled in an ice bath, diluted with water (10 mL) and extracted by EtOAc (3 × 15 mL). The organic phases were combined, washed with brine, dried over MgSO_4_, filtered and the solvent was removed under reduced pressure. The crude product was stirred with cold Et_2_O, and the resulting solid was filtered, rinsed with cold Et_2_O (2 × 2 mL) and dried under reduced pressure for 24 h. The crude powder was used without further purification (224 mg, 0.88 mmol, 18% isolated yield). **^1^H NMR (600 MHz, (CD_3_)_2_SO) δ** 7.62 – 7.56 (m, 2H), 7.49 – 7.43 (m, 1H), 7.44 – 7.38 (m, 2H), 5.68 (s, 2H). **^13^C NMR (151 MHz, (CD_3_)_2_SO) δ** 186.9, 137.9, 130.6, 128.0, 127.5, 120.6, 119.1.

#### 2’-fluoro-[1,1’-biphenyl]-3-carbaldehyde (53)

The product was prepared following the general procedure C starting from 3-bromobenzaldehyde (0.12 mL, 1.0 mmol, 1 eq.) and 2-fluorophenylboronic acid (276 mg, 2 eq.) and was isolated as a translucent oil (135 mg, 0.68 mmol, 68% isolated yield). **^1^H NMR (600 MHz, CDCl_3_) δ** 10.09 (s, 1H), 8.06 (q, *J* = 1.9 Hz, 1H), 7.90 (dt, *J* = 7.7, 1.4 Hz, 1H), 7.83 (dtd, *J* = 7.7, 1.7, 1.2 Hz, 1H), 7.62 (t, *J* = 7.7 Hz, 1H), 7.48 (td, *J* = 7.7, 1.8 Hz, 1H), 7.38 (dddd, *J* = 8.3, 7.4, 5.0, 1.8 Hz, 1H), 7.25 (td, *J* = 7.5, 1.2 Hz, 1H), 7.19 (ddd, *J* = 10.8, 8.3, 1.2 Hz, 1H). **^13^C NMR (151 MHz, CDCl_3_) δ** 192.3 (d, *J* = 1.5 Hz), 159.9 (d, *J* = 248.6 Hz), 137.0, 136.8, 135.1 (d, *J* = 3.3 Hz), 130.8 (d, *J* = 3.2 Hz), 130.5 (d, *J* = 2.8 Hz), 129.9 (d, *J* = 8.2 Hz), 129.3, 128.9, 127.8 (d, *J* = 13.1 Hz), 124.8 (d, *J* = 3.7 Hz), 116.4 (d, *J* = 22.5 Hz).

#### 3’-fluoro-[1,1’-biphenyl]-3-carbaldehyde (54)

The product was prepared following the general procedure C starting from 3-bromobenzaldehyde (0.12 mL, 1.0 mmol, 1 eq.) and 3-fluorophenylboronic acid (276 mg, 2 eq.) and was isolated as a translucent oil (154 mg, 0.77 mmol, 77% isolated yield). **^1^H NMR (600 MHz, CDCl_3_) δ** 10.10 (s, 1H), 8.11 – 8.06 (m, 1H), 7.89 (dt, *J* = 7.6, 1.4 Hz, 1H), 7.84 (ddd, *J* = 7.7, 2.0, 1.2 Hz, 1H), 7.63 (t, *J* = 7.6 Hz, 1H), 7.46 – 7.40 (m, 2H), 7.33 (ddd, *J* = 9.9, 2.6, 1.7 Hz, 1H), 7.13 – 7.08 (m, 1H). **^13^C NMR (151 MHz, CDCl_3_) δ** 192.2 (d, *J* = 1.8 Hz), 163.4 (d, *J* = 246.2 Hz), 142.1 (d, *J* = 7.7 Hz), 141.1 (d, *J* = 2.3 Hz), 137.2, 133.1, 130.7 (d, *J* = 8.4 Hz), 129.8, 129.4, 128.2, 123.0 (d, *J* = 3.0 Hz), 115.0 (d, *J* = 21.1 Hz), 114.3 (d, *J* = 22.2 Hz).

#### 4’-fluoro-[1,1’-biphenyl]-3-carbaldehyde (55)

The product was prepared following the fluorophenylboronic acid (276 mg, 2 eq.) and was isolated as a translucent oil (115 mg, 0.58 mmol, 58% isolated yield). **^1^H NMR (600 MHz, CDCl_3_) δ** 10.09 (s, 1H), 8.06 (t, *J* = 1.8 Hz, 1H), 7.86 (dt, *J* = 7.6, 1.4 Hz, 1H), 7.82 (ddd, *J* = 7.7, 2.0, 1.2 Hz, 1H), 7.63 – 7.57 (m, 3H), 7.19 – 7.14 (m, 2H). **^13^C NMR (151 MHz, CDCl_3_) δ** 192.4 (d, *J* = 2.0 Hz), 163.0 (d, *J* = 247.7 Hz), 141.4, 137.1, 136.0 (d, *J* = 3.2 Hz), 133.0, 129.7, 129.0 (d, *J* = 8.5 Hz), 128.9, 128.0, 116.1 (d, *J* = 21.7 Hz).

#### 2’-methyl-[1,1’-biphenyl]-3-carbaldehyde (56)

The product was prepared following the general procedure C starting from 3-bromobenzaldehyde (0.12 mL, 1.0 mmol, 1 eq.) and *o*-tolylboronic acid (202 mg, 1.5 eq.) and was isolated as a translucent oil (29 mg, 0.15 mmol, 15% isolated yield). **^1^H NMR (600 MHz, CDCl_3_) δ** 10.09 (s, 1H), 8.10 (t, *J* = 1.8 Hz, 1H), 7.87 – 7.83 (m, 2H), 7.60 (t, *J* = 7.6 Hz, 1H), 7.47 – 7.41 (m, 2H), 7.37 (t, *J* = 7.6 Hz, 1H), 7.22 (ddt, *J* = 7.5, 1.9, 1.0 Hz, 1H), 2.44 (s, 3H). **^13^C NMR (151 MHz, CDCl_3_) δ** 192.5, 142.5, 139.8, 138.8, 137.0, 133.2, 129.6, 129.1, 128.9, 128.7, 128.4, 128.1, 124.4, 21.7.

#### 3’-methyl-[1,1’-biphenyl]-3-carbaldehyde (57)

The product was prepared following the general procedure C starting from 3-bromobenzaldehyde (0.12 mL, 1.0 mmol, 1 eq.) and *m*-tolylboronic acid (202 mg, 1.5 eq.) and was isolated as a translucent oil (69 mg, 0.35 mmol, 35% isolated yield). **^1^H NMR (600 MHz, CDCl_3_) δ** 10.07 (s, 1H), 7.87 (dt, *J* = 6.8, 1.9 Hz, 1H), 7.85 (td, *J* = 1.6, 0.8 Hz, 1H), 7.63 – 7.56 (m, 2H), 7.32 – 7.27 (m, 3H), 7.23 (dt, *J* = 7.3, 1.1 Hz, 1H), 2.27 (s, 3H). **^13^C NMR (151 MHz, CDCl_3_) δ** 192.5, 192.5, 143.1, 140.5, 136.5, 135.4, 130.7, 130.6, 129.8, 129.0, 128.3, 128.1, 126.2, 20.5.

#### 4’-methyl-[1,1’-biphenyl]-3-carbaldehyde (58)

The product was prepared following the general procedure C starting from 3-bromobenzaldehyde (0.12 mL, 1.0 mmol, 1 eq.) and *p*-tolylboronic acid (202 mg, 1.5 eq.) and was isolated as a translucent oil (113 mg, 0.58 mmol, 58% isolated yield). **^1^H NMR (600 MHz, CDCl_3_) δ** 10.09 (s, 1H), 8.09 (t, *J* = 1.8 Hz, 1H), 7.86 – 7.83 (m, 2H), 7.60 (t, *J* = 7.6 Hz, 1H), 7.56 – 7.51 (m, 2H), 7.32 – 7.27 (m, 2H), 2.42 (s, 3H). **^13^C NMR (151 MHz, CDCl_3_) δ** 192.6, 142.3, 138.1, 136.9, 133.0, 129.9, 129.6, 128.5, 128.1, 127.1, 21.3.

#### 3-(tetrazol-5-yl)benzaldehyde (59)

In a 100 mL round bottom flask equipped with a magnetic stirrer and a condenser, the 3-cyanobenzaldehyde (393 mg, 3.0 mmol, 1 eq.) was solubilized in DMF (1 mL) then diluted in water (75 mL). Both the sodium azide (292 mg, 1.5 eq.) and the zinc chloride (408 mg, 3 eq.) were added. The suspension was vigorously stirred and heated at 110°C for 48 h. The reaction completion was assessed by TLC (Rf = 0.08 in DCM:MeOH 98:2) before being cooled it in an ice bath. The precipitate was filtered, rinsed with ice-cold water (2 × 10 mL) then with diethylether (2 × 5 mL) and dried under reduced pressure. The crude salts were then solubilized in aqueous HCl 1 M (10 mL) and extracted by EtOAc (3 × 15 mL) to remove the zinc salts. The crude powder could be used without further purification (220 mg, 1.3 mmol, 42% isolated yield). **^1^H NMR (600 MHz, (CD_3_)_2_SO) δ** 10.13 (s, 1H), 8.57 (t, *J* = 1.7 Hz, 1H), 8.36 (dt, *J* = 7.8, 1.5 Hz, 1H), 8.12 (dt, *J* = 7.7, 1.4 Hz, 1H), 7.86 (t, *J* = 7.7 Hz, 1H). **^13^C NMR (151 MHz, (CD_3_)_2_SO) δ** 192.7, 136.9, 132.4, 132.1, 130.4, 127.3. **MS (ESI^+^) calculated for C_8_H_6_N_4_O:** [M-H]^-^ *m/z* = 173.0, found *m/z* = 173.0.

#### 3-(2-(tetrahydro-2H-pyran-2-yl)-2H-tetrazol-5-yl)benzaldehyde (60)

In a dry 25 mL round bottom flask equipped with a magnetic stirrer and a condenser, 3,4-dihydro-2*H*-pyrane (0.18 mL, 2 eq.) was solubilized in dry toluene (5 mL) and mixed with a solution of 3-(tetrazol-5-yl)benzaldehyde (174 mg, 1.0 mmol, 1 eq.) in dry DMF (0.79 mL). The TFA (0.008 mL, 0.1 eq.) was added dropwise. The flask was purged with argon and heated at 110°C for 24 h. The completion of the reaction was assessed by TLC (Rf = 0.57 in DCM:MeOH 98:2) before being quenched by addition of a 10% Na_2_CO_3_ solution (10 mL) and water (20 mL). The crude product was extracted by EtOAc (3 × 15 mL) and the organic phases were combined, washed with brine (20 mL), dried over MgSO_4_, filtered and the solvents were removed under reduced pressure. The purification was done by flash chromatography on a 12 g silica cartridge with a cyclo:DCM gradient (8:2, 3 CV; 8:2→0:10, 6 CV; methanol wash) and isolated as an oil (66 mg, 0.26 mmol, 26% isolated yield). **^1^H NMR (600 MHz, CDCl_3_) δ** 10.11 (s, 1H), 8.70 (t, *J* = 1.8 Hz, 1H), 8.47 (dt, *J* = 7.7, 1.5 Hz, 1H), 8.01 (dt, *J* = 7.7, 1.4 Hz, 1H), 7.68 (t, *J* = 7.7 Hz, 1H), 6.10 (dd, *J* = 7.6, 2.9 Hz, 1H), 4.08 – 4.02 (m, 1H), 3.88 – 3.80 (m, 1H), 2.57 – 2.46 (m, 1H), 2.25 – 2.15 (m, 2H), 1.88 – 1.71 (m, 3H). **^13^C NMR (151 MHz, CDCl_3_) δ** 191.81, 164.1, 137.1, 132.7, 130.8, 129.9, 129.0, 128.6, 88.2, 67.1, 29.2, 24.7, 20.9. **MS (ESI^-^) calculated for C_13_H_14_N_4_O_2_:** [M+Cl]^-^ *m/z* = 293.1, found *m/z* = 293.1.

### cAMP measurement assay (CamBio) protocol

CHO cells expressing A_2A_R and the biosensor were seeded in a black 384-well plate (Nunc) and grown at 37°C and 5% CO_2_ overnight. They were pre-incubated for 6 minutes with compounds at varied concentrations up to 30 µM, before adding the corresponding EC_80_ of A_2A_R agonist, 400 nM of adenosine (Sigma). The cAMP biosensor allowed real-time monitoring of signals using an FDSS/µCELL (Hamamatsu). The assay was conducted in 1X Hanks Balanced Salt Solution (HBSS). The total volume of the reaction was 80 µL (45 µL of cells, 15 µL of antagonist and 20 µL of agonist). Data analysis was performed using GraphPad Prism.

### NAM MoA confirmation (“shift assay”) protocol

CHO cells expressing A_2A_R and the biosensor were seeded in a black 384-well plate (Nunc) and grown at 37°C and 5% CO_2_ overnight. They were pre-incubated for 6 minutes with compounds at varied concentrations up to 30 µM, before adding adenosine at increasing concentrations up to 1 mM (CRC). The cAMP biosensor readings and assay volumes were as described in the CamBio assay. Data analysis was performed using GraphPad Prism.

A Schild regression plot was calculated for the compounds by using the shift assay data. The Log(DoseRatio-1) was plotted on the y axis against the Log of the antagonist concentration on the x axis.

### Binding cooperativity factor α and dissociation constant K_B_ determination

Using the data from the shift assay, α and K_B_ were calculated directly in GraphPad Prism using the non-linear regression equation “allosteric EC_50_ shift, X is log(concentration)” as per instructions.

### Interaction analysis by Grating-coupled Interferometry (GCI)

Kinetic binding measurements were performed using GCI on a Creoptix WAVEdelta system (Creoptix – a Malvern Panalytical brand). A PCP-NTA Creoptix WAVEchip (sensor chip) was conditioned according to the manufacturer’s specifications. The surface was activated by a 120 second injection of 0.5 mM NiCl_2_ at a flow rate of 10 ml/min on 2 parallel flow channels. Purified A_2_AR was injected at 100 mg/mL in running buffer (20 mM Tris-HCl pH 7.5, 350 mM NaCl, 0.1% (w/v) n-Dodecyl-β-D-maltopyranoside (DDM), 2% (v/v) Dimethyl Sulfoxide (DMSO) at a flow rate of 1 mL/min to capture the receptor *via* its c-terminal His-tag to a level of 5500 pg/mm^2^. The surface of both channels was stabilized with several injections of running buffer. The kinetic measurements of XAC binding to A_2_AR were performed using A-B-A type injections, ensuring tested compounds being present at 100 nM, 1 µM or 10 µM, respectively, during the full measurement from baseline throughout XAC dissociation. XAC was injected in a dilution series from 1.6 nM to 200 nM for 60 seconds at a flow rate of 50 mL/min, followed by dissociation in running buffer for 120 seconds.

Data analysis was performed using the Creoptix WAVEcontrol software (Creoptix – a Malvern Panalytical brand). The response signals were double referenced, subtracting the signal recorded on the reference channel w/o protein as well as the signal of blank injections with running buffer. Double-referenced signals were fit to a 1:1 Langmuir binding model, determining association rate (k_a_), dissociation rate (k_d_), maximum response (R_max_) and the dissociation constant (K_D_) for each interaction. Single injection cycles showing significant distortions, or no binding response were excluded from the analysis.

### pCREB measurement protocol

PBMCs were isolated from healthy human whole blood. Thawed PBMCs were incubated with A_2A_R antagonists in the presence of NECA and were subsequently fixed and permeabilized using ice cold paraformaldehyde and methanol-based kit reagents (ThermoFisher and Miltenyi, respectively). Cells were washed with FACS buffer and stained with fluorochrome-conjugated antibodies directed against the following antigens: pCREB (Miltenyi), CD4 and CD8 (Life Technologies), and CD3 (BD Biosciences). Stained cells were acquired using a Fortessa flow cytometer (BD). Gating of the desired cell subsets was performed using FSC-A vs FSC-H to remove doublet cells, FSC vs SSC parameters to identify cells with lymphocyte-like morphology, followed by CD3^+^ T cell and CD4^+^ or CD8^+^ sub-gating. The mean fluorescence intensity (MFI) fold change in of intracellular pCREB in gated CD4^+^ or CD8^+^ T cells was calculated using DMSO-only condition and then normalized with NECA-only condition. The results shown are an average of several experiments done on different donors, with a minimum of two repetitions for each donor.

## Supporting information

Medicinal chemistry table

## ASSOCIATED CONTENT

Molecular formula strings with biological data

Supporting figures; Shift assay plots, ^1^H NMR, ^13^C NMR, and HPLC analysis results of target compounds.

## AUTHOR INFORMATION

### Author Contributions

M.B., M.H., A.G., C.S., S.T., D.P., H.H. and L.S. designed the experiments and analyzed the data. M.B., A.S. and S.T. designed, synthesized, and analyzed the compounds. A.G., C.S., M.G. and D.P. performed the biological experiments. M.H. and L.S. performed the docking studies. M.B. wrote the manuscript, with edits from M.H., A.G., C.S., S.T., H.H., D.P. and L.S. T.D.S., A.G., D.P. and L.S. contributed to the idea of allosterically targeting A_2a_R D.P. and L.S. acquired funding for the project.

### Notes

Conflict of interest: Authors of the School of Pharmaceutical Sciences of the University of Geneva and the Institute of Pharmaceutical Sciences of Western Switzerland (ISPSO) are minor or major shareholders in the company Adoram Therapeutics.

## ACKNOWLEDGMENT

The authors thank Pr. Walter Reith from the University of Geneva for the extensive discussions, as well as Dr Michael Hennig, Dr Matilde Trabuco, Dr Fabio Andres, and their colleagues from LeadXPro for their help in elaborating results of biophysical measurements and this manuscript. NMR spectra acquisitions were possible thanks to Dr Laurence Marcourt and Frédéric Borlat at the NMR core facility of the School of Pharmaceutical Sciences (ISPSO). MS and HRMS measurements were performed with the assistance of Dr Emmanuel Varesio, respectively at the School of Pharmaceutical Sciences (ISPSO) and at the Chemical Biology Mass Spectrometry core facility (Faculty of Science - University of Geneva). Authors would like to acknowledge the work and support of the team of the technology transfer office of the University of Geneva UNITEC. The present research received the financial support of Innosuisse (project grant 46357.1 IP-LS), the Swiss National Fund (project grant 186405 to L.S), the University of Geneva, and the Eclosion Fundation for which the authors are grateful. For the purpose of Open Access, a CC BY public copyright license is applied to any Author Accepted Manuscript version arising from this submission.

## ABBREVIATIONS

A_2A_R: Adenosine 2A Receptor
CHO: Chinese Hamster Ovary
CV: Column Volume
DCM: Dichloromethane
4-DMAP: 4-Dimethylaminopyridine
DMF: Dimethylformamide
DMSO: Dimethylsulfoxide
ESI: Electrospray Ionization
GCI: Grating-Coupled Interferometry
GPCR: G-Protein Coupled Receptor
(HR)MR: (High Resolution) Mass Spectrometry
MFI: Mean Fluorescence Intensity
NAM: Negative Allosteric Modulator
NMR: Nuclear Magnetic Resonance
PBMC: Peripheral Blood Mononuclear Cells
SAR: Structure-Activity Relationship
TFA: Trifluoroacetic acid
THF: Tetrahydrofuran
THP: Tetrahydropyran
TLC: Thin Layer Chromatography
TME: Tumor Microenvironment
TOF: Time Of Flight

## REFERENCES

(1) Jacobson, K. A.; Gao, Z.-G. Adenosine Receptors as Therapeutic Targets. Nat Rev Drug Discov 2006, 5 (3), 247–264. 10.1038/nrd1983.

(2) de Lera Ruiz, M.; Lim, Y.-H.; Zheng, J. Adenosine A2A Receptor as a Drug Discovery Target. J. Med. Chem. 2014, 57 (9), 3623–3650. 10.1021/jm4011669.

(3) Borea, P. A.; Gessi, S.; Merighi, S.; Vincenzi, F.; Varani, K. Pharmacology of Adenosine Receptors: The State of the Art. Physiol Rev 2018, 98 (3), 1591–1625. 10.1152/physrev.00049.2017.

(4) Allard, B.; Allard, D.; Buisseret, L.; Stagg, J. The Adenosine Pathway in Immuno-Oncology. Nat Rev Clin Oncol 2020, 17 (10), 611–629. 10.1038/s41571-020-0382-2.

(5) Fredholm, B. B.; IJzerman, A. P.; Jacobson, K. A.; Linden, J.; Müller, C. E. International Union of Basic and Clinical Pharmacology. LXXXI. Nomenclature and Classification of Adenosine Receptors—An Update. Pharmacol Rev 2011, 63 (1), 1–34. 10.1124/pr.110.003285.

(6) Allard, B.; Beavis, P. A.; Darcy, P. K.; Stagg, J. Immunosuppressive Activities of Adenosine in Cancer. Curr Opin Pharmacol 2016, 29, 7–16. 10.1016/j.coph.2016.04.001.

(7) Immunopharmacology and Inflammation; Riccardi, C., Levi-Schaffer, F., Tiligada, E., Eds.; Springer: Cham, 2018.

(8) Sidders, B.; Zhang, P.; Goodwin, K.; O’Connor, G.; Russell, D. L.; Borodovsky, A.; Armenia, J.; McEwen, R.; Linghu, B.; Bendell, J. C.; Bauer, T. M.; Patel, M. R.; Falchook, G. S.; Merchant, M.; Pouliot, G.; Barrett, J. C.; Dry, J. R.; Woessner, R.; Sachsenmeier, K. Adenosine Signaling Is Prognostic for Cancer Outcome and Has Predictive Utility for Immunotherapeutic Response. Clin Cancer Res 2020, 26 (9), 2176–2187. 10.1158/1078-0432.CCR-19-2183.

(9) Pinna, A. Adenosine A2A Receptor Antagonists in Parkinson’s Disease: Progress in Clinical Trials from the Newly Approved Istradefylline to Drugs in Early Development and Those Already Discontinued. CNS drugs 2014, 28, 455–474. 10.1007/s40263-014-0161-7.

(10) Augustin, R. C.; Leone, R. D.; Naing, A.; Fong, L.; Bao, R.; Luke, J. J. Next Steps for Clinical Translation of Adenosine Pathway Inhibition in Cancer Immunotherapy. J Immunother Cancer 2022, 10 (2), e004089. 10.1136/jitc-2021-004089.

(11) Korkutata, M.; Agrawal, L.; Lazarus, M. Allosteric Modulation of Adenosine A2A Receptors as a New Therapeutic Avenue. Int J Mol Sci 2022, 23 (4), 2101. 10.3390/ijms23042101.

(12) Kenakin, T. The Quantitative Characterization of Functional Allosteric Effects. Curr Protoc Pharmacol 2017, 76, 9.22.1–9.22.10. 10.1002/cpph.18.

(13) Lane, J. R.; Jaakola, V.-P.; IJzerman, A. P. Chapter 1 - The Structure of the Adenosine Receptors: Implications for Drug Discovery. In Advances in Pharmacology; Jacobson, K. A., Linden, J., Eds.; Pharmacology of Purine and Pyrimidine Receptors; Academic Press, 2011; Vol. 61, pp 1–40. 10.1016/B978-0-12-385526-8.00001-1.

(14) Carpenter, B.; Lebon, G. Human Adenosine A2A Receptor: Molecular Mechanism of Ligand Binding and Activation. Front. Pharmacol. 2017, 8. 10.3389/fphar.2017.00898.

(15) Liu, W.; Chun, E.; Thompson, A. A.; Chubukov, P.; Xu, F.; Katritch, V.; Han, G. W.; Roth, C. B.; Heitman, L. H.; IJzerman, A. P.; Cherezov, V.; Stevens, R. C. Structural Basis for Allosteric Regulation of GPCRs by Sodium Ions. Science 2012, 337 (6091), 232–236. 10.1126/science.1219218.

(16) Massink, A.; Amelia, T.; Karamychev, A.; IJzerman, A. P. Allosteric Modulation of G Protein-coupled Receptors by Amiloride and Its Derivatives. Perspectives for Drug Discovery? Med Res Rev 2020, 40 (2), 683–708. 10.1002/med.21633.

(17) Caliman, A. D.; Miao, Y.; McCammon, J. A. Mapping the Allosteric Sites of the A _2A_ Adenosine Receptor. Chem Biol Drug Des 2018, 91 (1), 5–16. 10.1111/cbdd.13053.

(18) Giorgi, I.; Biagi, G.; Bianucci, A. M.; Borghini, A.; Livi, O.; Leonardi, M.; Pietra, D.; Calderone, V.; Martelli, A. N6-1,3-Diphenylurea Derivatives of 2-Phenyl-9-Benzyladenines and 8-Azaadenines: Synthesis and Biological Evaluation as Allosteric Modulators of A2A Adenosine Receptors. European Journal of Medicinal Chemistry 2008, 43 (8), 1639–1647. 10.1016/j.ejmech.2007.10.021.

(19) Welihinda, A. A.; Amento, E. P. Positive Allosteric Modulation of the Adenosine A2a Receptor Attenuates Inflammation. J Inflamm (Lond) 2014, 11 (1), 37. 10.1186/s12950-014-0037-0.

(20) Welihinda, A.; Ravikumar, P.; Kaur, M.; Mechanic, J.; Yadav, S.; Kang, G. J.; Amento, E. Positive Allosteric Modulation of A2AR Alters Immune Cell Responses and Ameliorates Psoriasis-Like Dermatitis in Mice. Journal of Investigative Dermatology 2022, 142 (3, Part A), 624–632.e6. 10.1016/j.jid.2021.07.174.

(21) Korkutata, M.; Saitoh, T.; Cherasse, Y.; Ioka, S.; Duo, F.; Qin, R.; Murakoshi, N.; Fujii, S.; Zhou, X.; Sugiyama, F.; Chen, J.-F.; Kumagai, H.; Nagase, H.; Lazarus, M. Enhancing Endogenous Adenosine A2A Receptor Signaling Induces Slow-Wave Sleep without Affecting Body Temperature and Cardiovascular Function. Neuropharmacology 2019, 144, 122–132. 10.1016/j.neuropharm.2018.10.022.

(22) Lin, Y.; Roy, K.; Ioka, S.; Otani, R.; Amezawa, M.; Ishikawa, Y.; Cherasse, Y.; Kaushik, M. K.; Klewe-Nebenius, D.; Zhou, L.; Yanagisawa, M.; Oishi, Y.; Saitoh, T.; Lazarus, M. Positive Allosteric Adenosine A2A Receptor Modulation Suppresses Insomnia Associated with Mania- and Schizophrenia-like Behaviors in Mice. Front. Pharmacol. 2023, 14, 1138666. 10.3389/fphar.2023.1138666.

(23) Gao, Z.-G.; Ijzerman, A. P. Allosteric Modulation of A2A Adenosine Receptors by Amiloride Analogues and Sodium Ions. Biochemical Pharmacology 2000, 60 (5), 669–676. 10.1016/S0006-2952(00)00360-9.

(24) Lu, Y.; Liu, H.; Yang, D.; Zhong, L.; Xin, Y.; Zhao, S.; Wang, M.-W.; Zhou, Q.; Shui, W. Affinity Mass Spectrometry-Based Fragment Screening Identified a New Negative Allosteric Modulator of the Adenosine A2A Receptor Targeting the Sodium Ion Pocket. ACS Chem. Biol. 2021, 16 (6), 991–1002. 10.1021/acschembio.0c00899.

(25) Murray, T. J.; Zimmerman, S. C.; Kolotuchin, S. V. Synthesis of Heterocyclic Compounds Containing Three Contiguous Hydrogen Bonding Sites in All Possible Arrangements. Tetrahedron 1995, 51 (2), 635–648. 10.1016/0040-4020(94)00922-H.

(26) Sarkar, S.; Das, D. K.; Khan, A. T. Synthesis of Fully-Substituted Pyridines and Dihydropyridines in a Highly Chemoselective Manner Utilizing a Multicomponent Reaction (MCR) Strategy. RSC Adv. 2014, 4 (96), 53752–53760. 10.1039/C4RA08237K.

(27) Hanna, N.; Kicka, S.; Chiriano, G.; Harrison, C.; Sakouhi, H. O.; Trofimov, V.; Kranjc, A.; Nitschke, J.; Pagni, M.; Cosson, P.; Hilbi, H.; Scapozza, L.; Soldati, T. Identification of Anti-Mycobacterium and Anti-Legionella Compounds With Potential Distinctive Structural Scaffolds From an HD-PBL Using Phenotypic Screens in Amoebae Host Models. Front. Microbiol. 2020, 11. 10.3389/fmicb.2020.00266.

(28) Lindsley, C. W. 2013 Philip S. Portoghese Medicinal Chemistry Lectureship: Drug Discovery Targeting Allosteric Sites. J. Med. Chem. 2014, 57 (18), 7485–7498. 10.1021/jm5011786.

(29) Jakubík, J.; Randáková, A.; Chetverikov, N.; El-Fakahany, E. E.; Doležal, V. The Operational Model of Allosteric Modulation of Pharmacological Agonism. Sci Rep 2020, 10 (1), 14421. 10.1038/s41598-020-71228-y.

