## Supplementary information for "Discovery of the first-efficacious A_2A_R negative allosteric modulators for high adenosine cancer immunotherapies"

Supporting Figures	2

Shift assay plots for selected examples	3

HPLC traces for selected examples	6

^1^H & ^13^C NMR spectra for target compounds	20

References	67

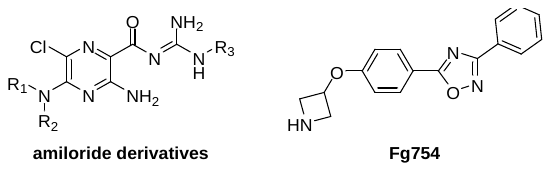

### Supporting Figures

*Figure S1. Structures of the known A_2A_R NAMs.*^[1,2]^

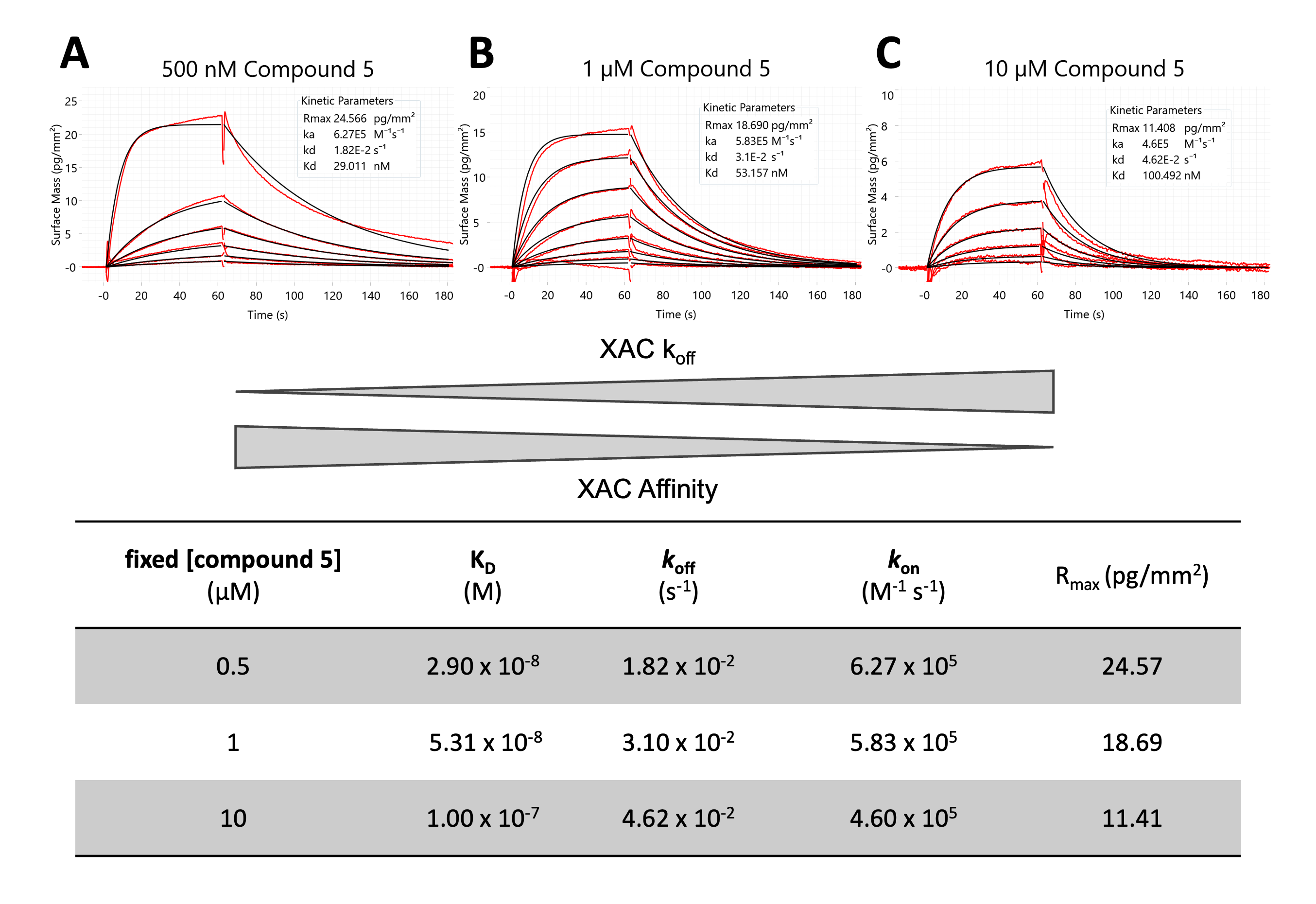
*Figure S2. GCI kinetic characterization of XAC binding to A_2A_R at different concentrations of compound* ***5****: (A) 500 nM, (B) 1 µM and (C) 10 µM. Double-referenced binding signals are shown in red, 1:1 Langmuir interaction model fits are shown in black. Table: Kinetic parameters derived from fitting double-referenced binding signals to a 1:1 interaction model at two concentrations of compound* ***5****.*

### Shift assay plots for selected examples

| Compound 4 | 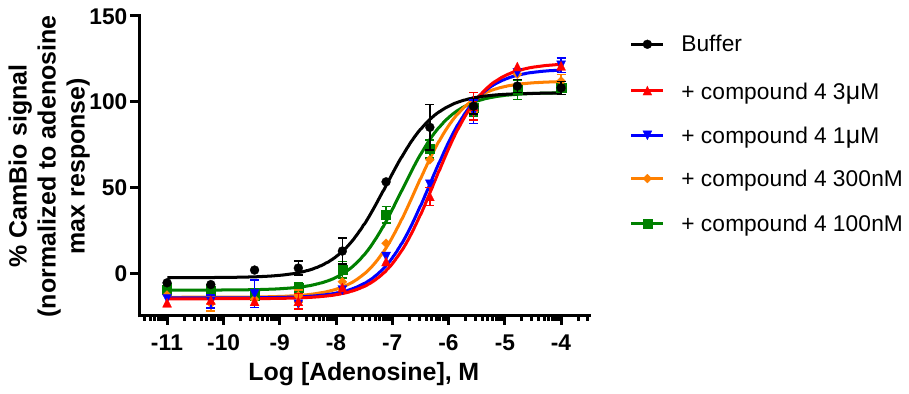 |
| --- | --- |
| Compound 38 | 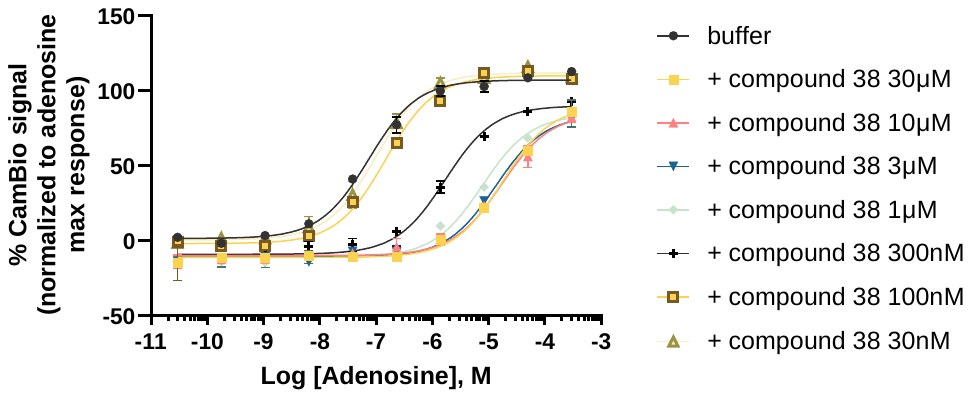 |
| Compound 39 | 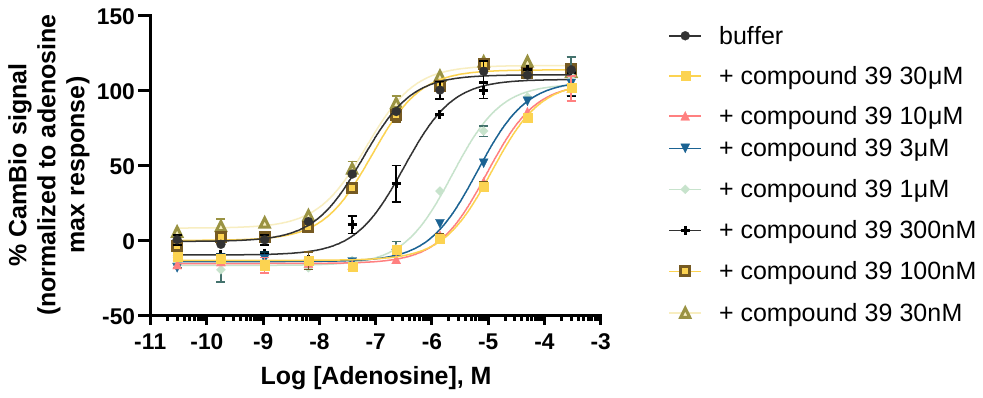 |
| Compound 41 | 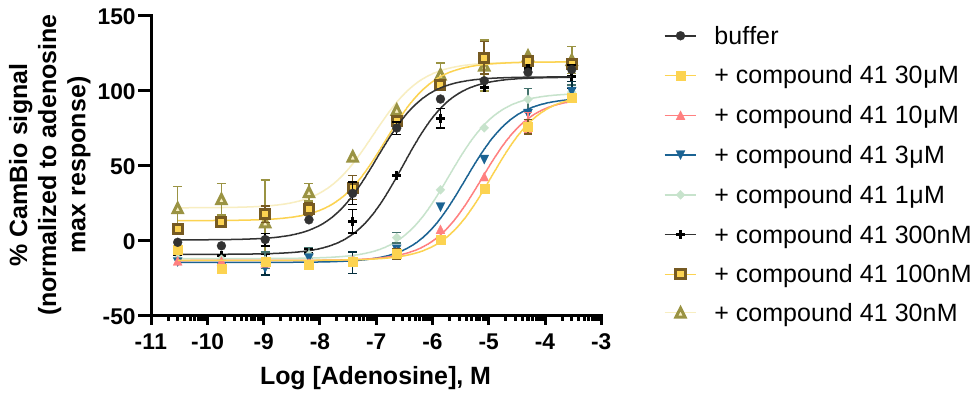 |
| Compound 42 | 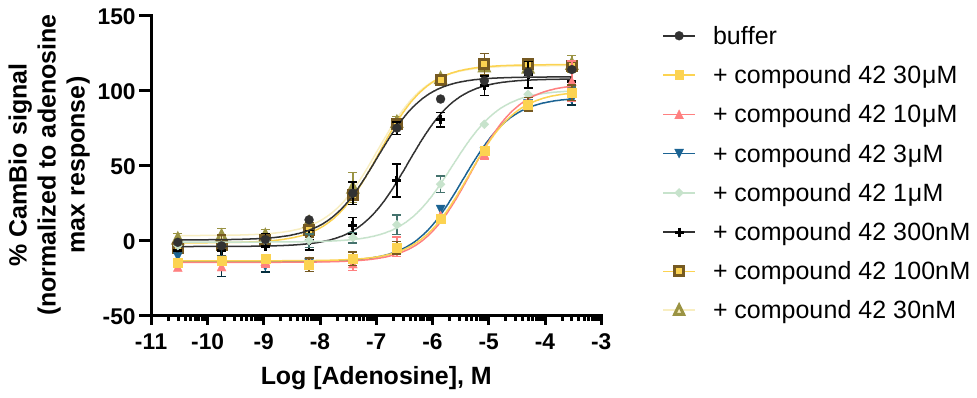 |
| Compound 44 | 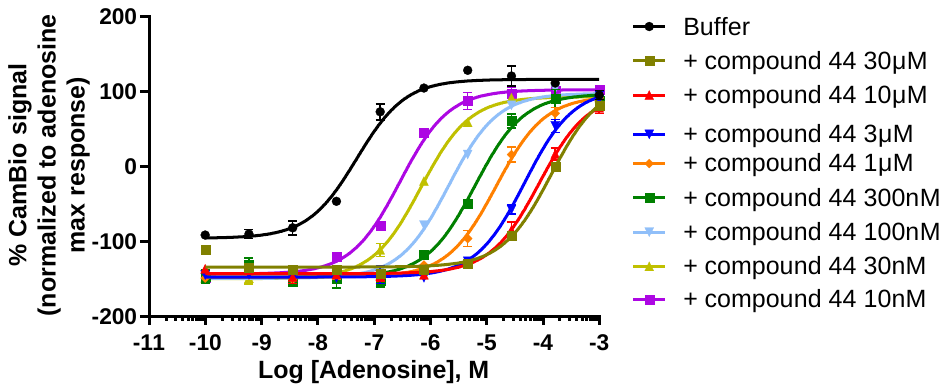 |
| Compound 46 | 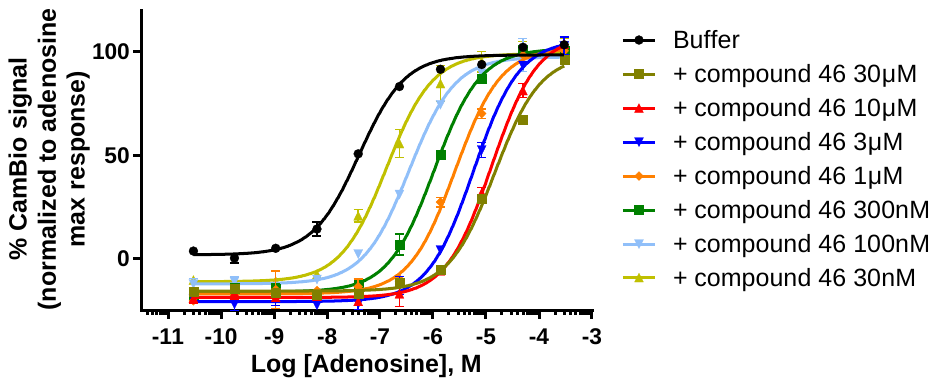 |
| Compound 47 | 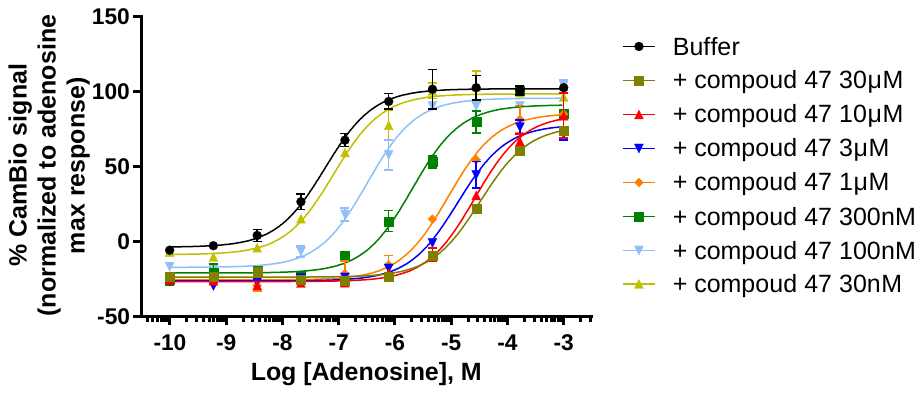 |
| Compound 48 | 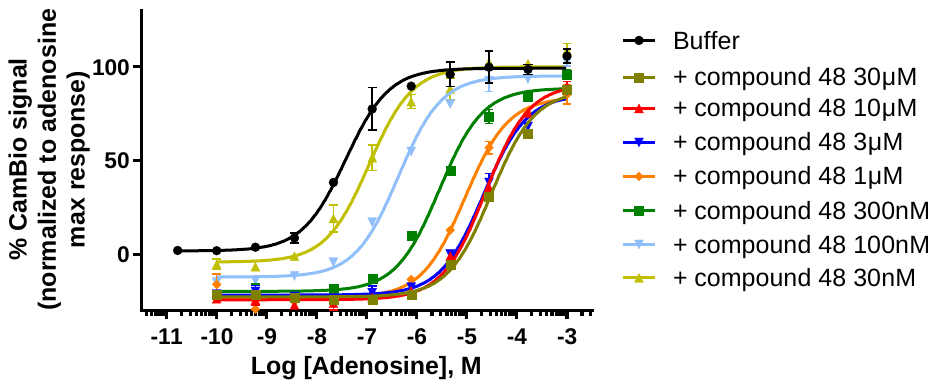 |

### HPLC traces for selected examples

**Compound 5:**

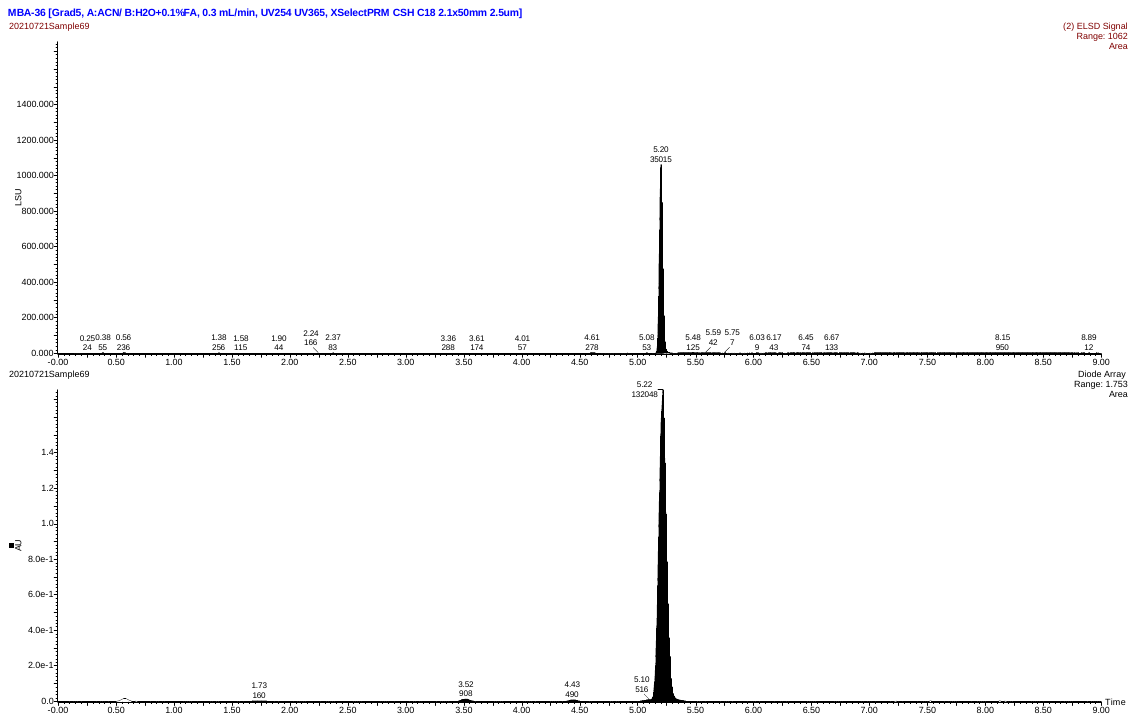
UV retention time: 5.22 min

UV purity: 98.4%

| UV rt (min) | UV peak height | UV peak area |
| --- | --- | --- |
| 1.733 | 1402 | 160.017 |
| 3.517 | 12478 | 907.966 |
| 4.433 | 7463 | 490.216 |
| 5.100 | 8535 | 516.309 |
| 5.217 | 1752576 | 132047.500 |

**Compound 11:**

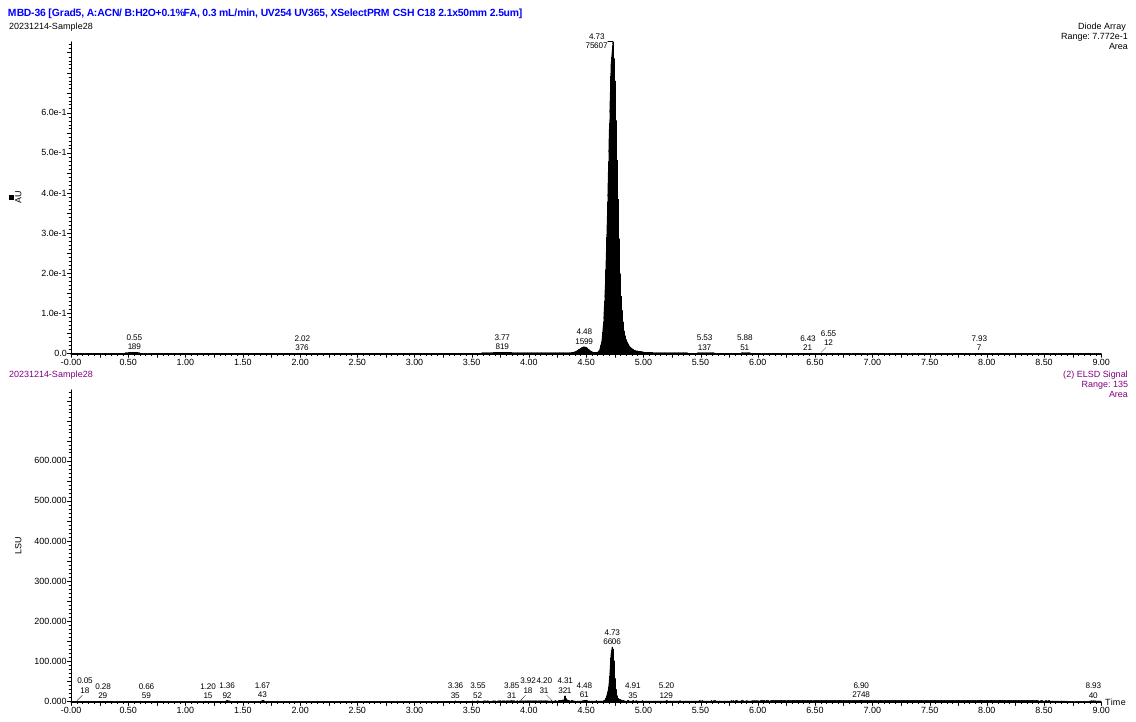
UV retention time (rt): 4.73 min

UV purity: 96.2%

| UV rt (min) | UV peak height | UV peak area |
| --- | --- | --- |
| 2.017 | 296 | 376.017 |
| 3.767 | 2594 | 819.133 |
| 4.483 | 15429 | 1598.768 |
| 4.733 | 777216 | 75606.875 |
| 5.533 | 798 | 137.008 |
| 5.883 | 605 | 51.275 |
| 6.433 | 336 | 21.383 |
| 6.550 | 223 | 12.300 |

**Compound 13:**

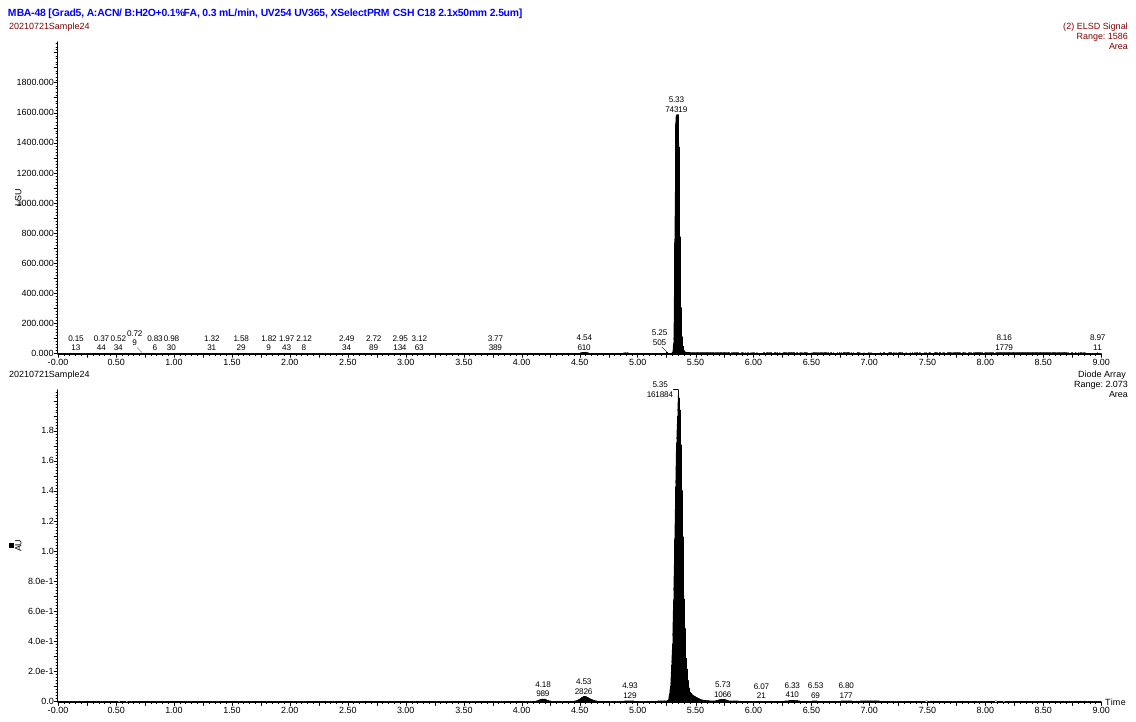
UV retention time: 5.35 min

UV purity: 96.6%

| UV rt (min) | UV peak height | UV peak area |
| --- | --- | --- |
| 4.183 | 14805 | 989.066 |
| 4.533 | 31161 | 2826.334 |
| 4.933 | 2113 | 129.183 |
| 5.350 | 2072896 | 161883.547 |
| 5.733 | 11682 | 1066.234 |
| 5.950 | 92 | 3.683 |
| 6.067 | 261 | 20.508 |
| 6.333 | 5530 | 409.616 |
| 6.533 | 950 | 68.850 |
| 6.800 | 1434 | 176.613 |

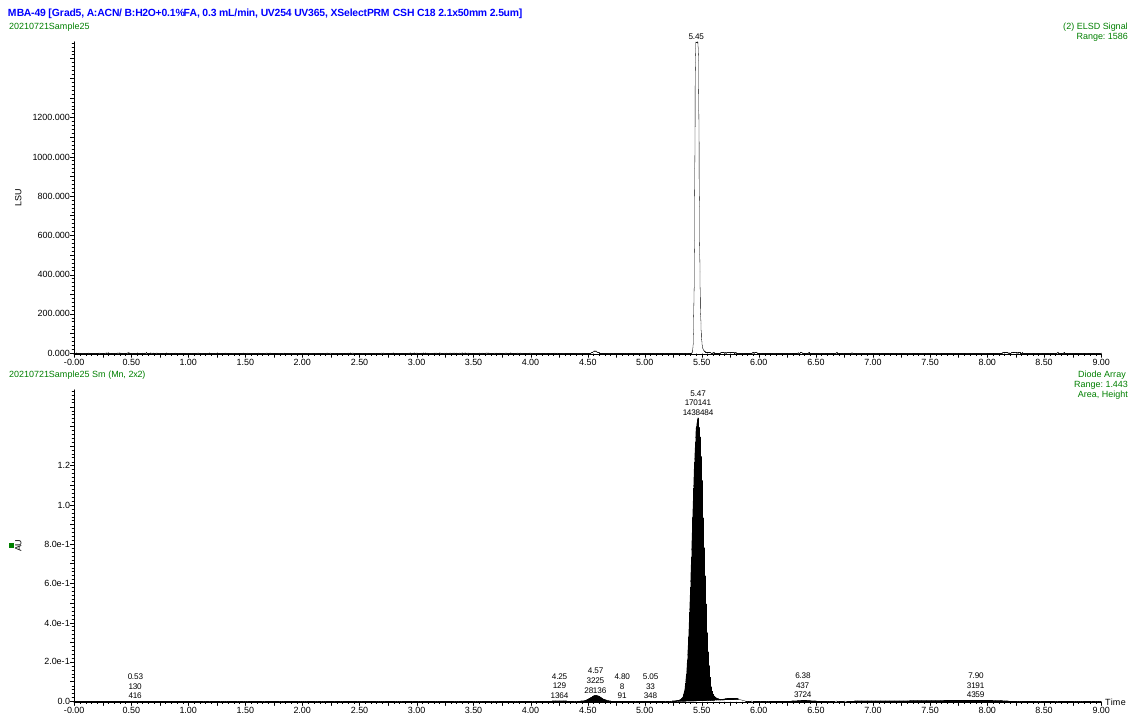
**Compound 14:**

UV retention time: 5.47 min

UV purity: 95.0%

| **#** | **Name** | **RT** | **Area** |
| --- | --- | --- | --- |
| **1** | compound | 5.47 | 171011344 |
| **2** | compound impurity 1 | 4.57 | 3226215 |
| **3** | compound impurity 2 | 5.77 | 2259913 |
| **4** | compound impurity 3 | 7.90 | 4010548 |

**Compound 15:**

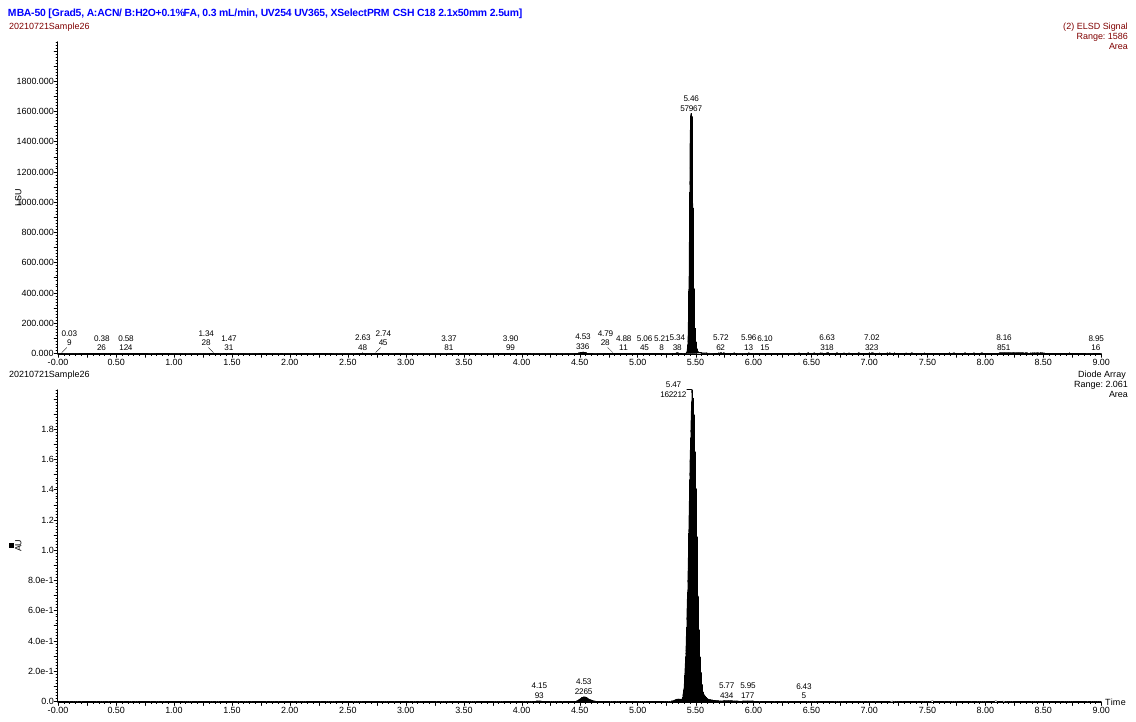
UV retention time: 5.47 min

UV purity: 98.2%

| UV rt (min) | UV peak height | UV peak area |
| --- | --- | --- |
| 4.150 | 2246 | 93.150 |
| 4.533 | 29396 | 2265.468 |
| 5.467 | 2060672 | 162212.297 |
| 5.767 | 4618 | 434.258 |
| 5.950 | 2422 | 177.092 |
| 6.433 | 156 | 5.167 |

**Compound 28:**

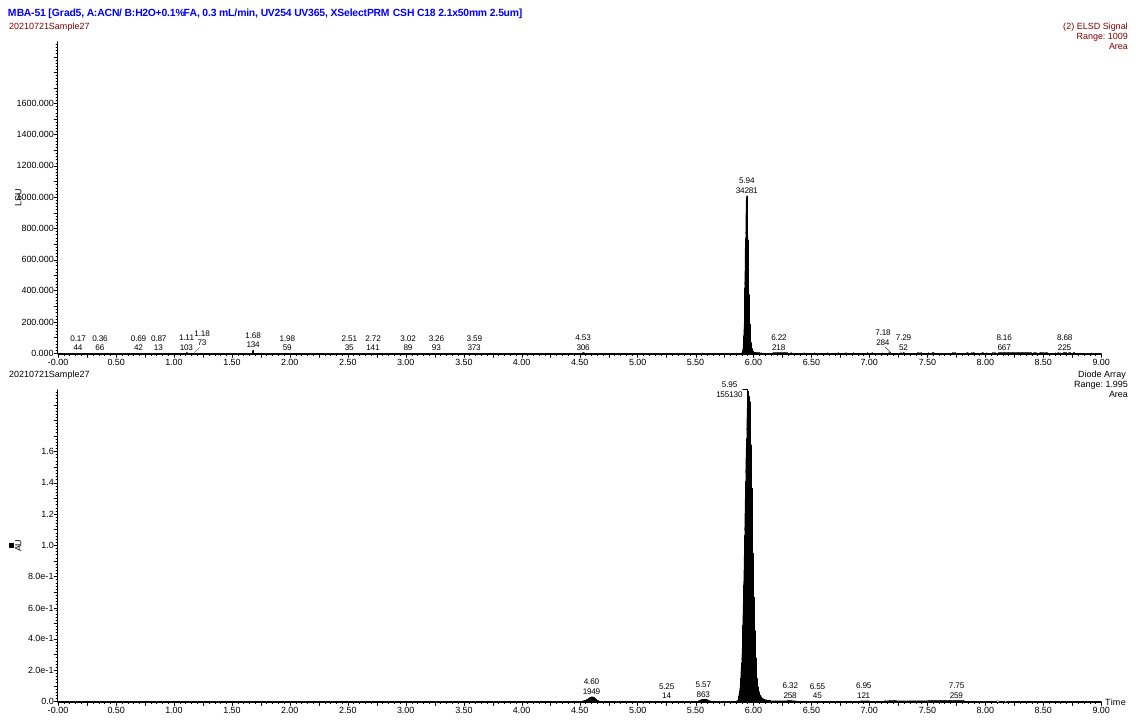
UV retention time: 5.95 min

UV purity: 97.8%

| UV rt (min) | UV peak height | UV peak area |
| --- | --- | --- |
| 4.600 | 27780 | 1949.298 |
| 5.250 | 411 | 13.650 |
| 5.567 | 13004 | 862.633 |
| 5.950 | 1995328 | 155129.531 |
| 6.317 | 3430 | 258.200 |
| 6.550 | 903 | 45.100 |
| 6.950 | 1147 | 121.399 |
| 7.750 | 694 | 258.875 |

**Compound 33:**

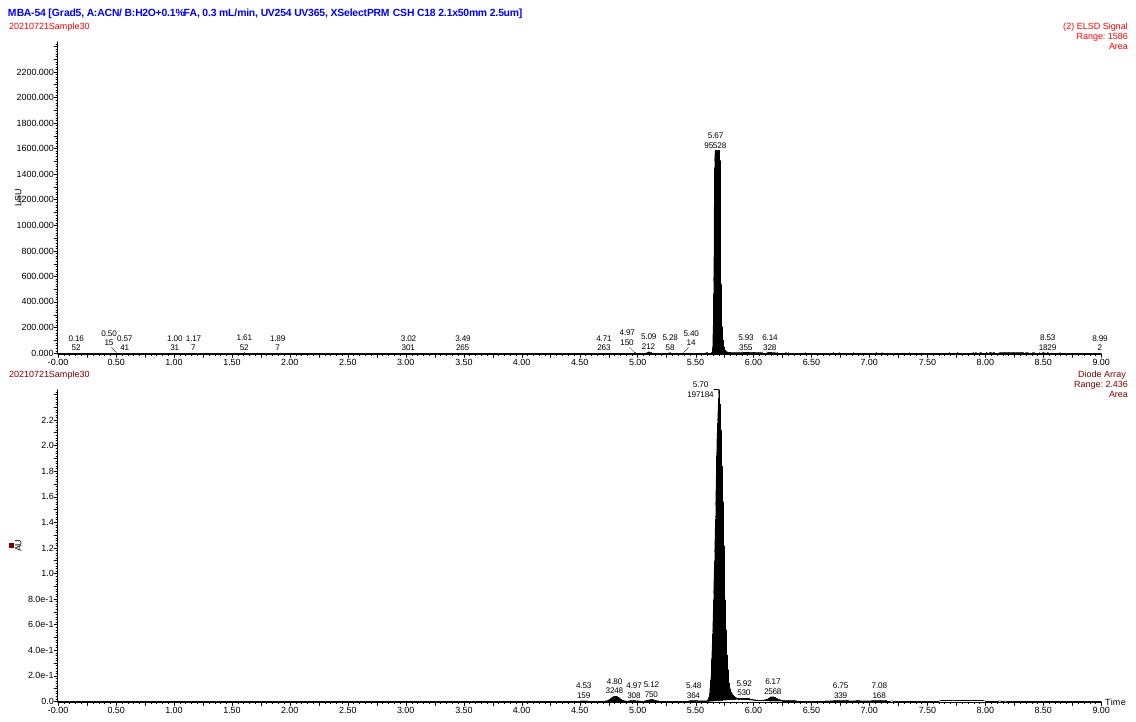
UV retention time: 5.70 min

UV purity: 96.1%

| UV rt (min) | UV peak height | UV peak area |
| --- | --- | --- |
| 4.533 | 3517 | 159.183 |
| 4.800 | 38700 | 3247.774 |
| 4.967 | 4771 | 308.142 |
| 5.117 | 11409 | 750.383 |
| 5.483 | 4194 | 364.134 |
| 5.700 | 2427022 | 197183.859 |
| 5.917 | 7687 | 529.880 |
| 6.167 | 29927 | 2567.550 |

**Compound 44:**

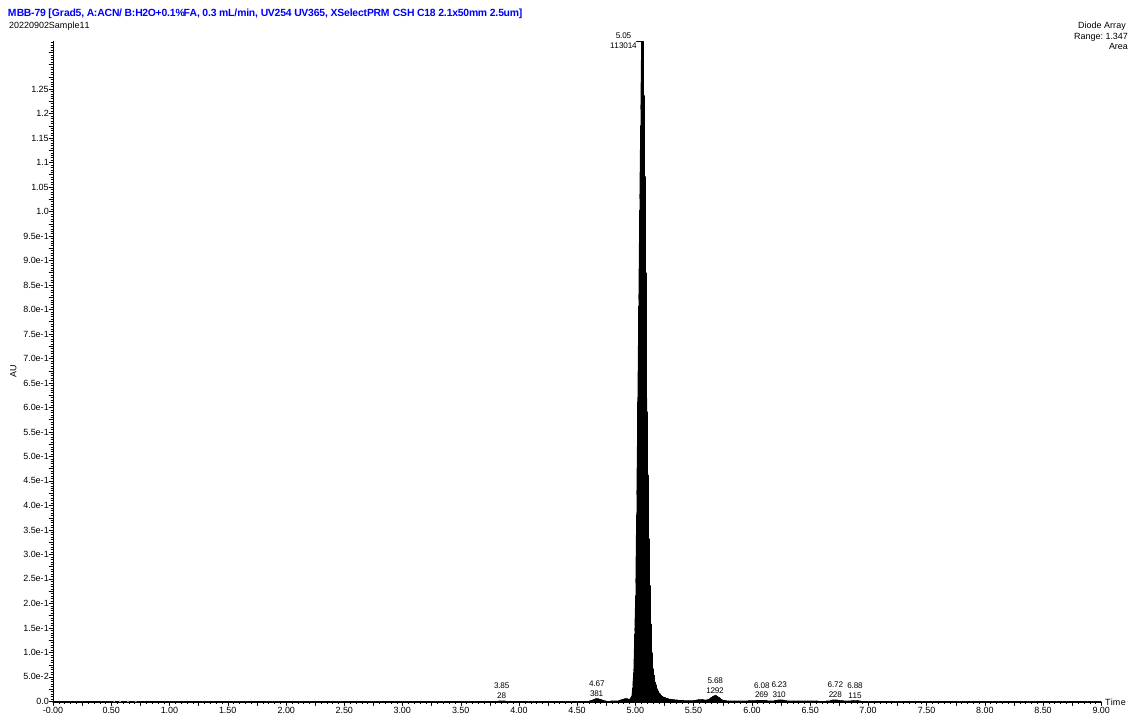
UV retention time: 5.05 min

UV purity: 97.7%

| UV rt (min) | UV peak height | UV peak area |
| --- | --- | --- |
| 3.850 | 410 | 27.900 |
| 4.667 | 5091 | 381.200 |
| 5.050 | 1346926 | 113014.359 |
| 5.684 | 11398 | 1291.777 |
| 6.084 | 1863 | 269.141 |
| 6.234 | 2277 | 309.534 |
| 6.717 | 2457 | 227.516 |
| 6.884 | 1507 | 114.600 |

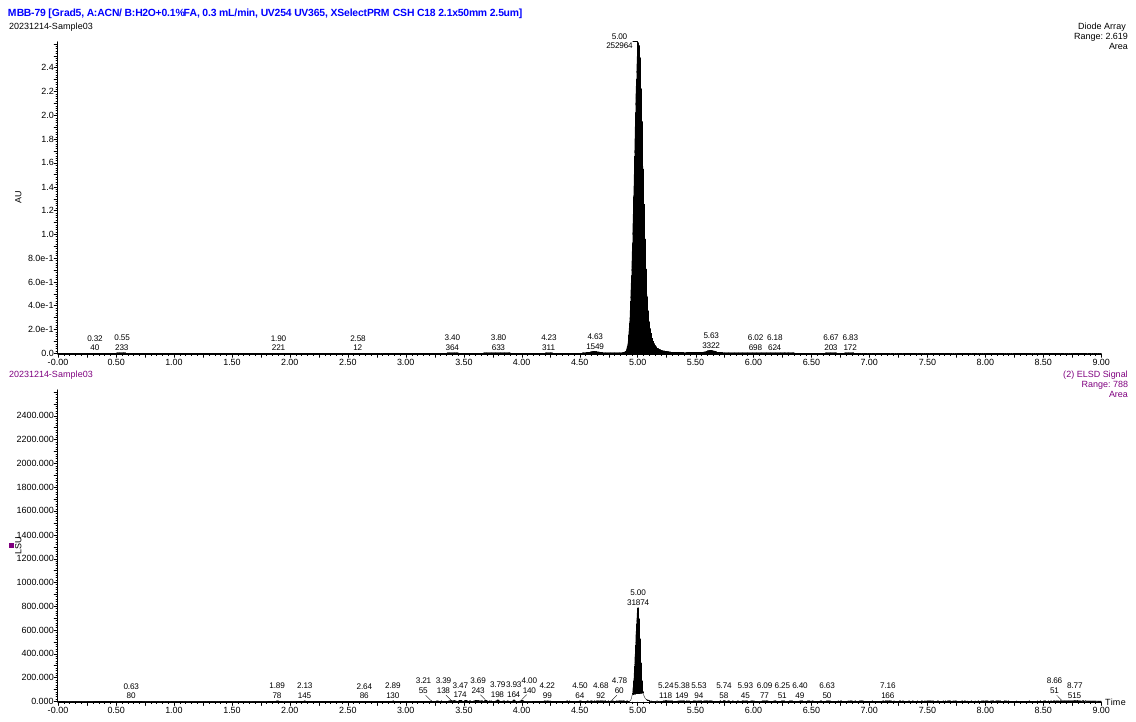
**Compound 45:**

UV retention time (rt): 5.00 min

UV purity: 97.0%

| UV rt (min) | UV peak height | UV peak area |
| --- | --- | --- |
| 1.900 | 219 | 220.883 |
| 2.583 | 150 | 12.117 |
| 3.400 | 1772 | 364.408 |
| 3.800 | 2446 | 633.387 |
| 4.233 | 1445 | 310.883 |
| 4.633 | 12797 | 1548.844 |
| 5.000 | 2618577 | 252963.563 |
| 5.633 | 22441 | 3321.590 |
| 6.017 | 3703 | 697.804 |
| 6.183 | 3475 | 624.450 |
| 6.667 | 2059 | 203.033 |

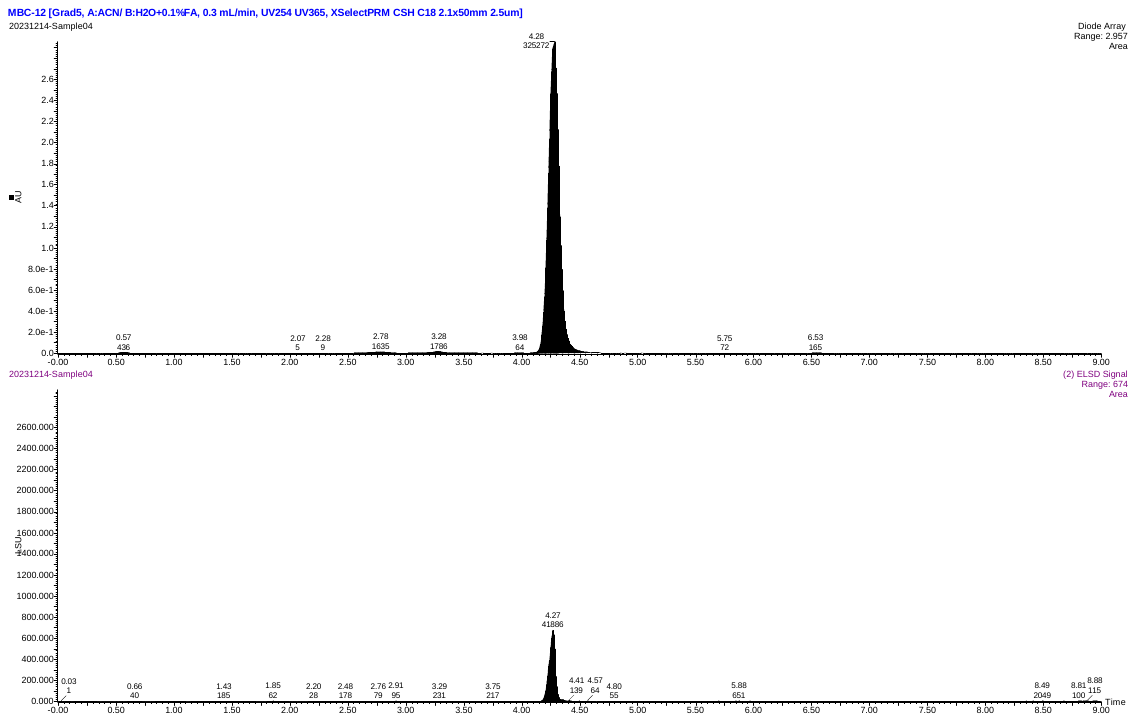
**Compound 46:**

UV retention time: 4.28 min

UV purity: 98.8%

| UV rt (min) | UV peak height | UV peak area |
| --- | --- | --- |
| 0.750 | 41 | 2.684 |
| 1.383 | 28 | 12.042 |
| 2.067 | 59 | 4.524 |
| 2.283 | 51 | 9.226 |
| 2.783 | 11288 | 1634.858 |
| 3.283 | 13041 | 1786.324 |
| 3.983 | 891 | 63.900 |
| 4.283 | 2953111 | 325271.969 |
| 5.750 | 833 | 72.367 |
| 6.533 | 2597 | 164.633 |

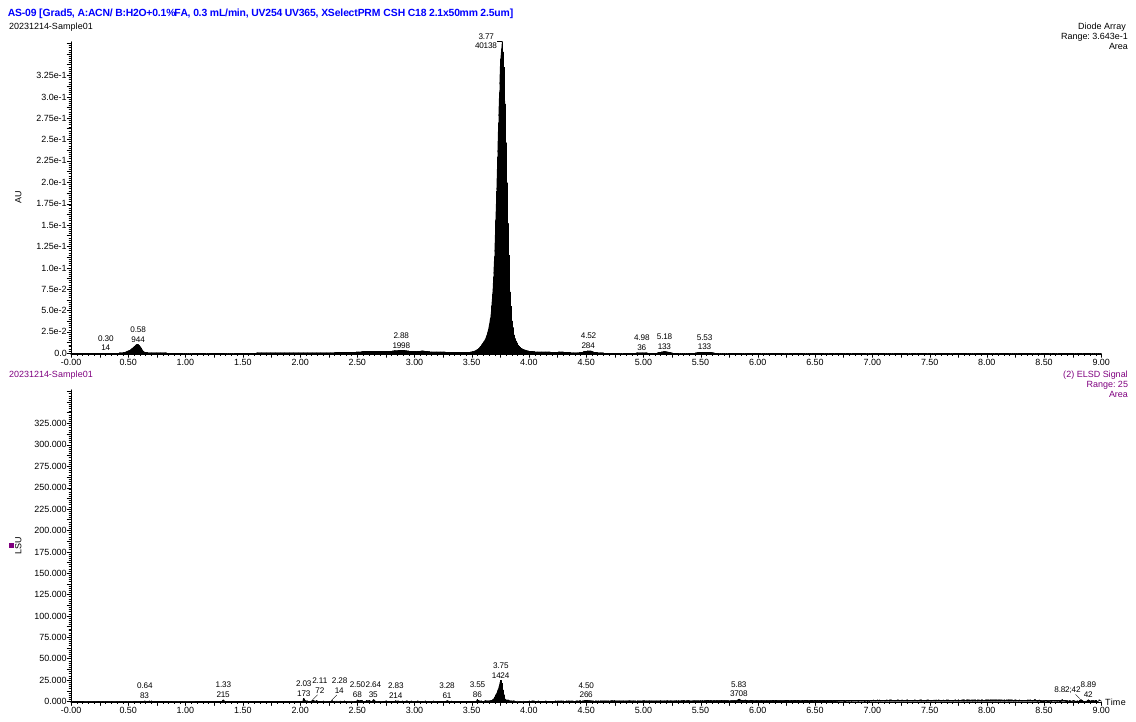
**Compound 47:**

UV retention time: 3.77 min

UV purity: 96.0%

| UV rt (min) | UV peak height | UV peak area |
| --- | --- | --- |
| 2.883 | 2385 | 1093.150 |
| 3.767 | 363191 | 39098.125 |
| 4.517 | 2237 | 218.458 |
| 4.983 | 544 | 36.233 |
| 5.183 | 1747 | 133.017 |
| 5.533 | 1232 | 132.617 |

**Compound 48:**

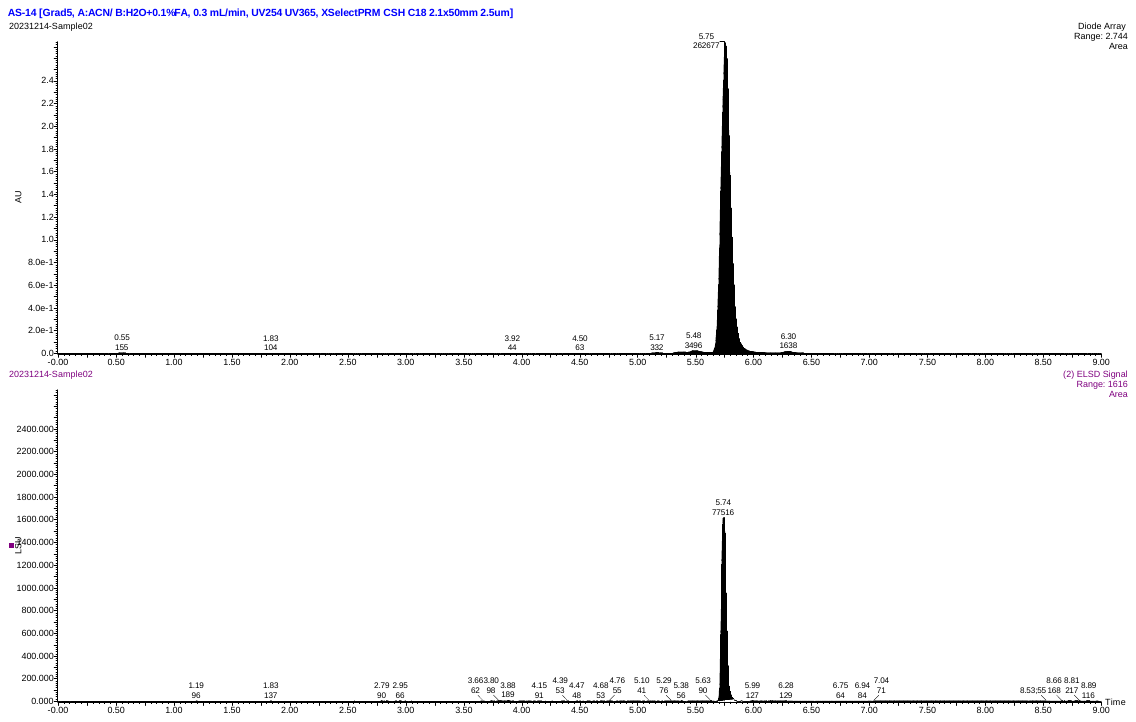
UV retention time (rt): 5.75 min

UV purity: 97.8%

| UV rt (min) | UV peak height | UV peak area |
| --- | --- | --- |
| 0.650 | 64 | 2.783 |
| 1.833 | 146 | 103.950 |
| 3.917 | 647 | 44.100 |
| 4.167 | 13 | 0.217 |
| 4.500 | 778 | 63.250 |
| 5.167 | 4258 | 331.550 |
| 5.483 | 21699 | 3496.273 |
| 5.750 | 2743930 | 262677.031 |
| 6.300 | 15876 | 1637.712 |

**Compound 49:**

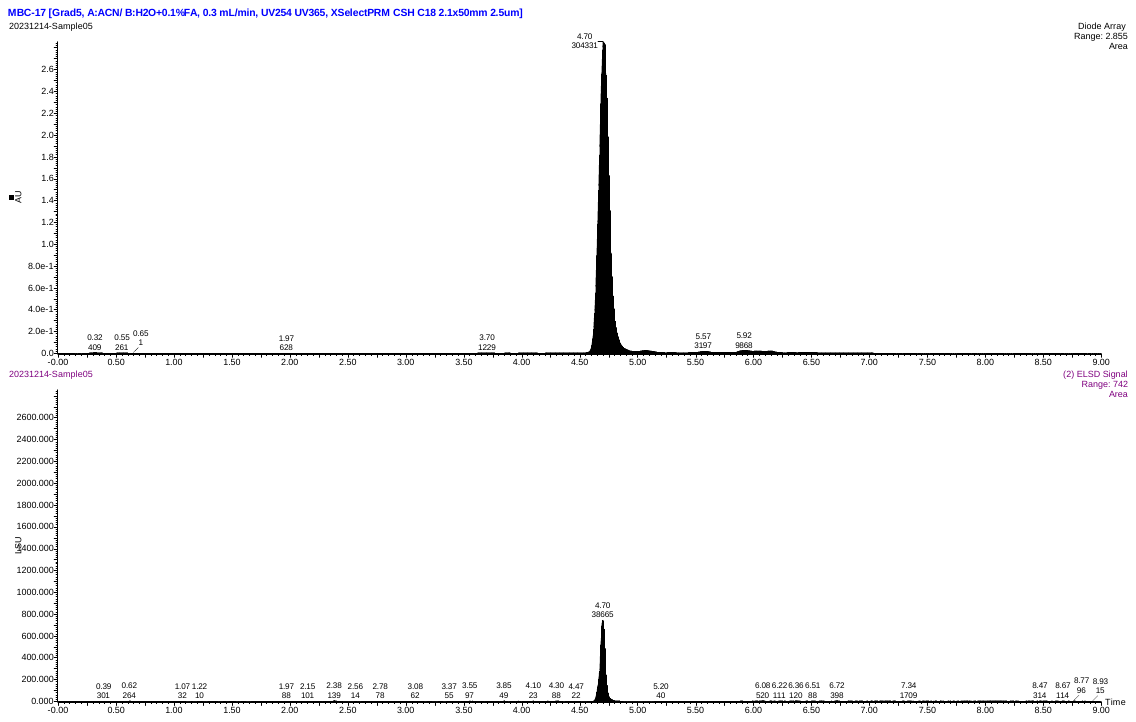
UV retention time (rt): 4.70 min

UV purity: 95.3%

| UV rt (min) | UV peak height | UV peak area |
| --- | --- | --- |
| 0.650 | 29 | 0.783 |
| 1.967 | 441 | 627.643 |
| 3.700 | 2723 | 1229.253 |
| 4.700 | 2855391 | 304331.125 |
| 5.567 | 16487 | 3196.939 |
| 5.917 | 25647 | 9868.385 |

**Compound 50:**

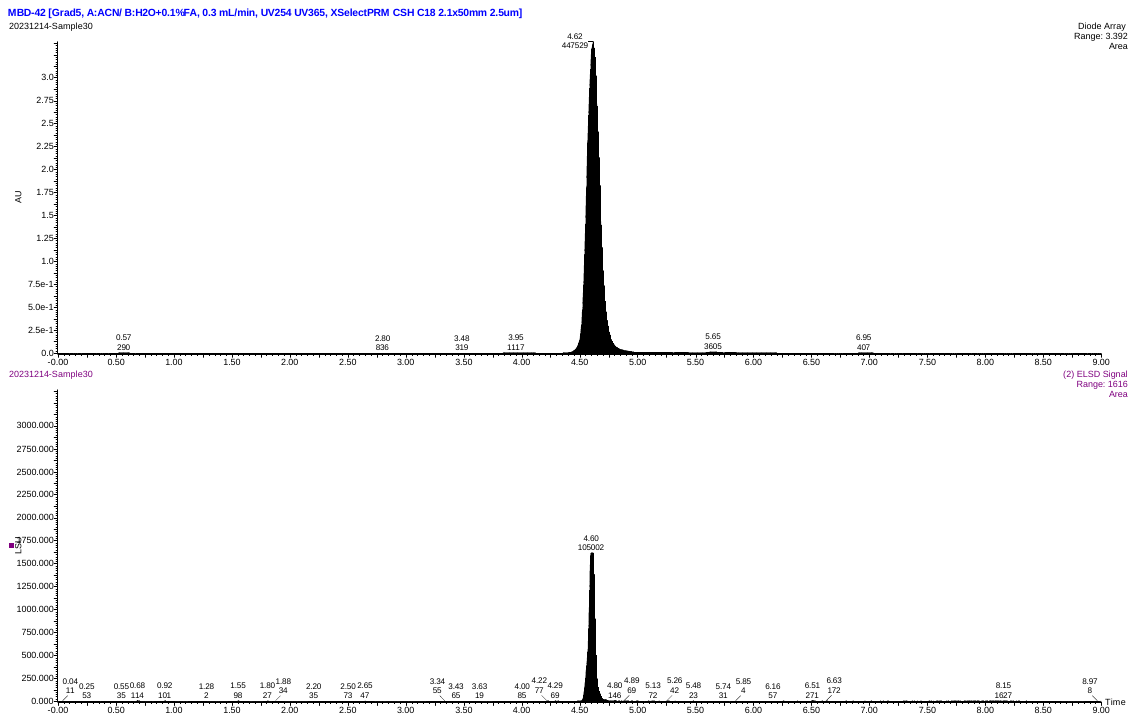
UV retention time (rt): 4.62 min

UV purity: 98.7%

| UV rt (min) | UV peak height | UV peak area |
| --- | --- | --- |
| 2.800 | 1369 | 836.411 |
| 3.483 | 1256 | 319.470 |
| 3.950 | 3687 | 1117.057 |
| 4.617 | 3392246 | 447528.813 |
| 5.650 | 14945 | 3604.956 |

###  ^1^H & ^13^C NMR spectra for target compounds

Compound 5:

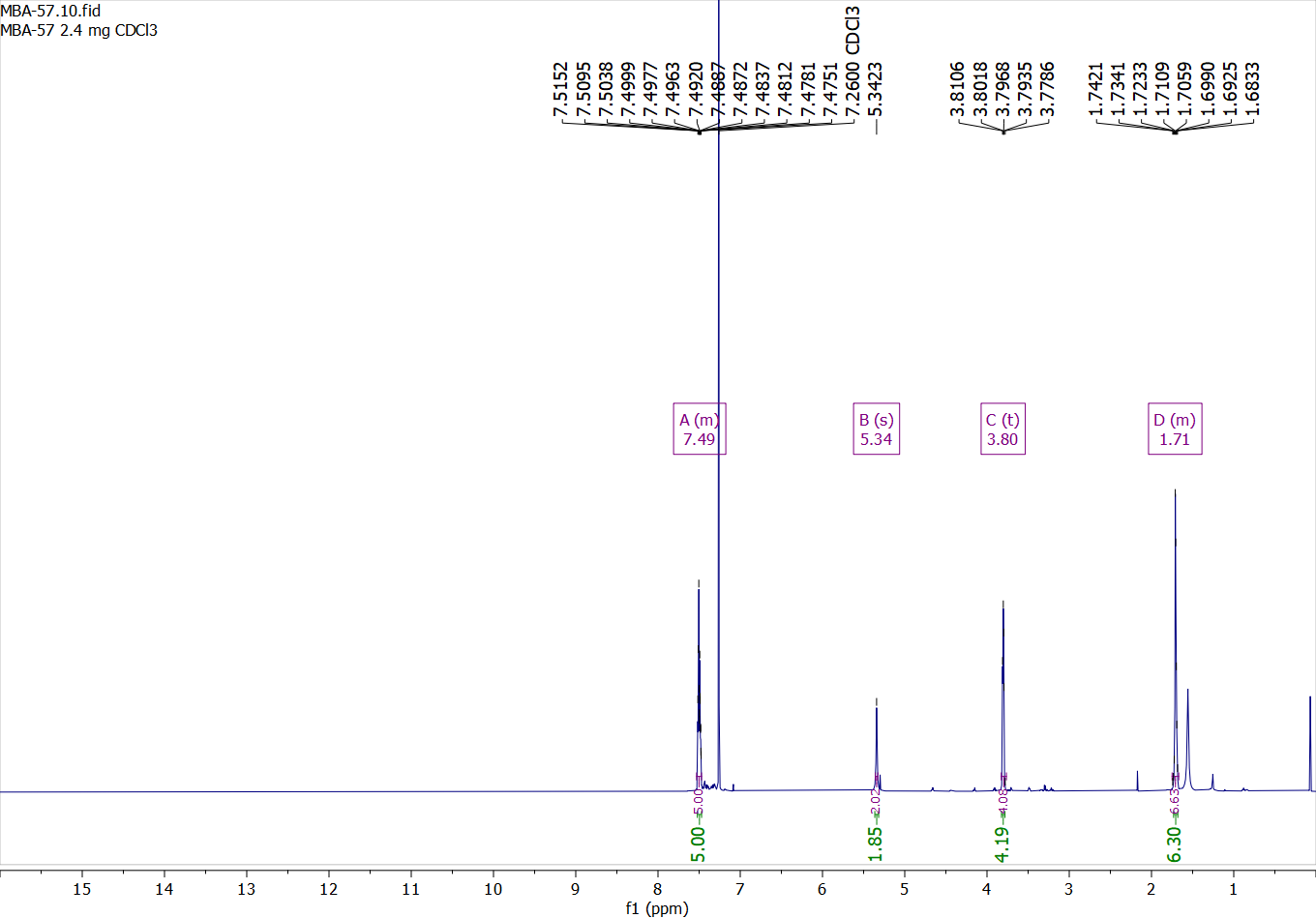

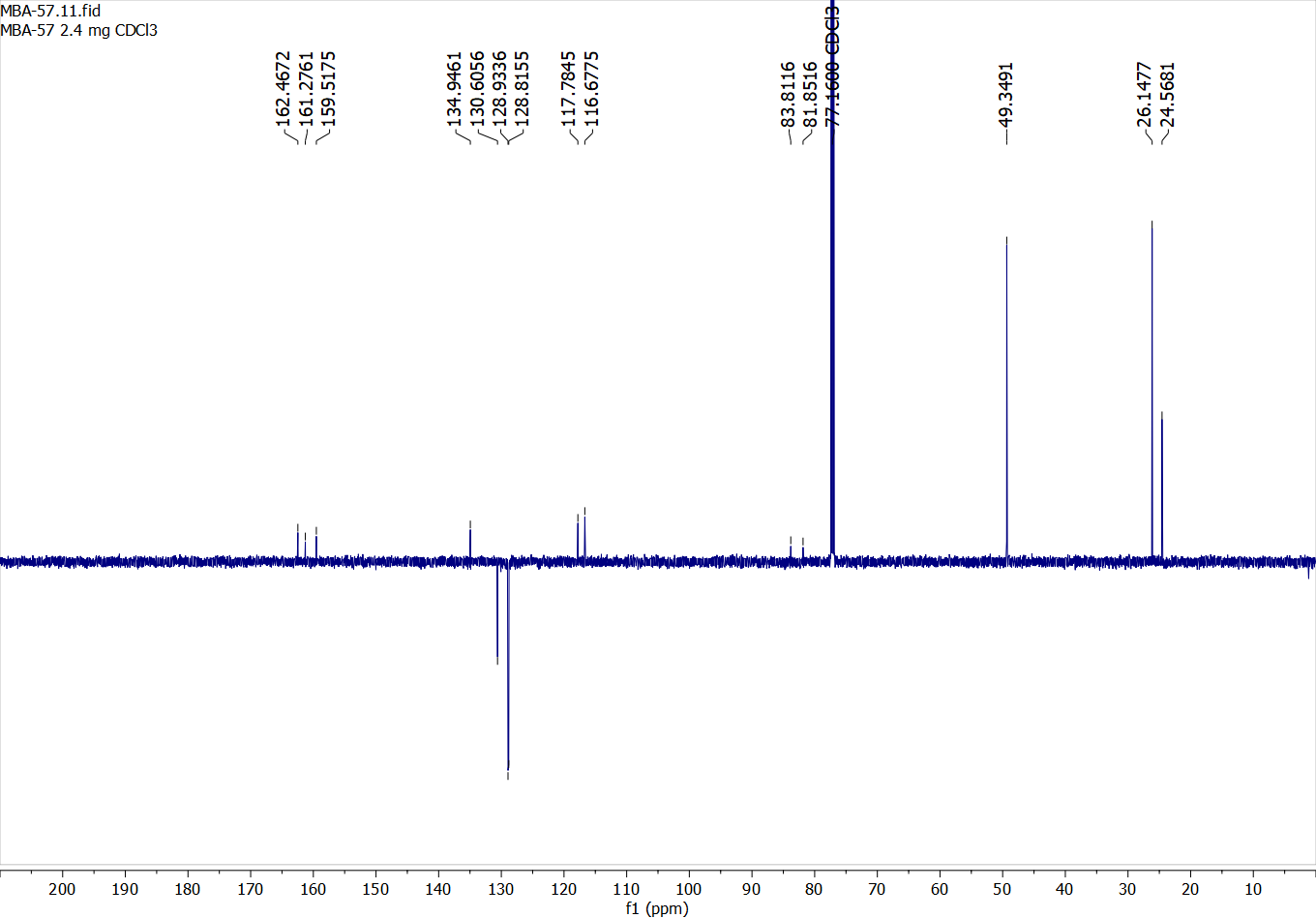

Compound 6:

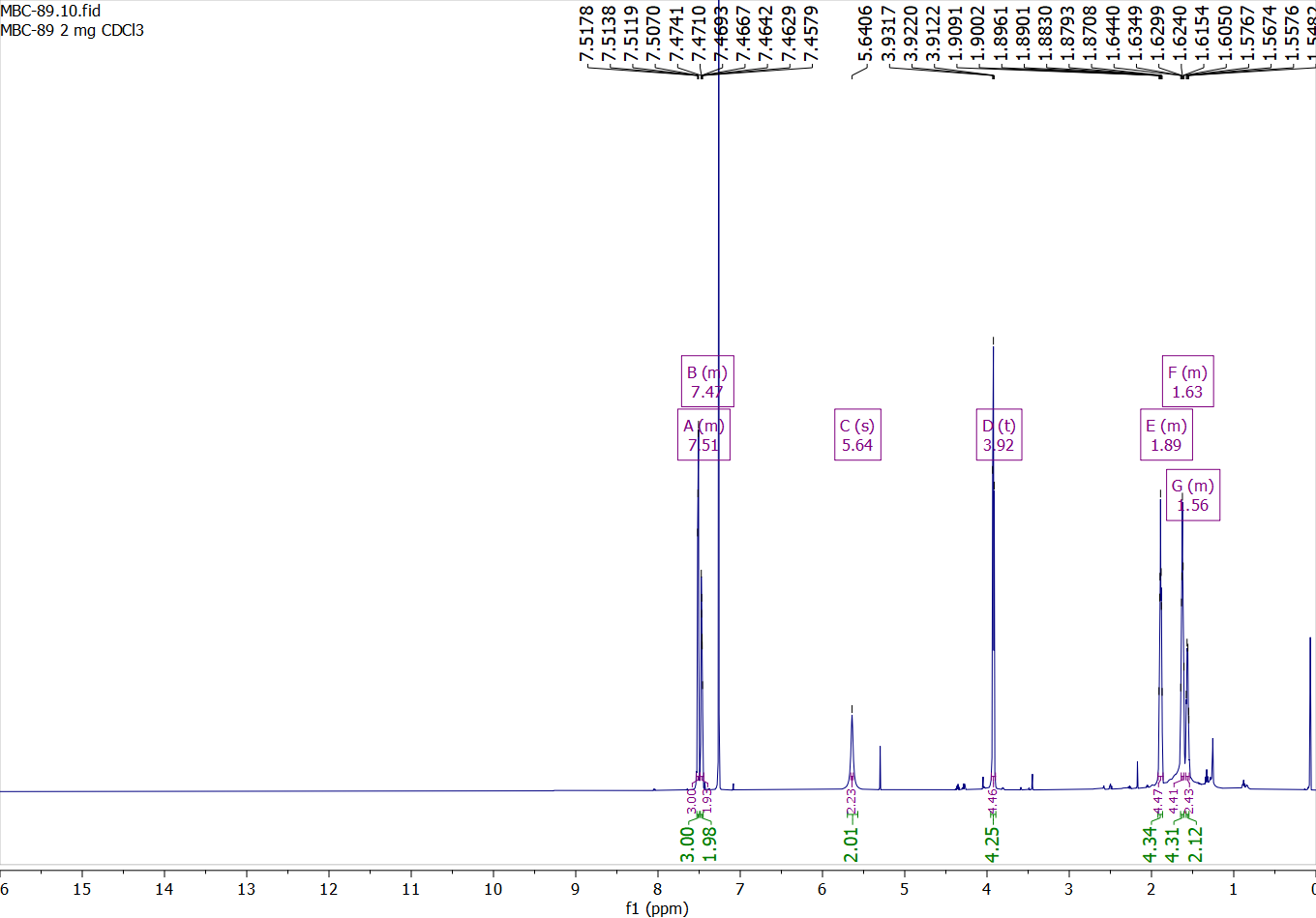

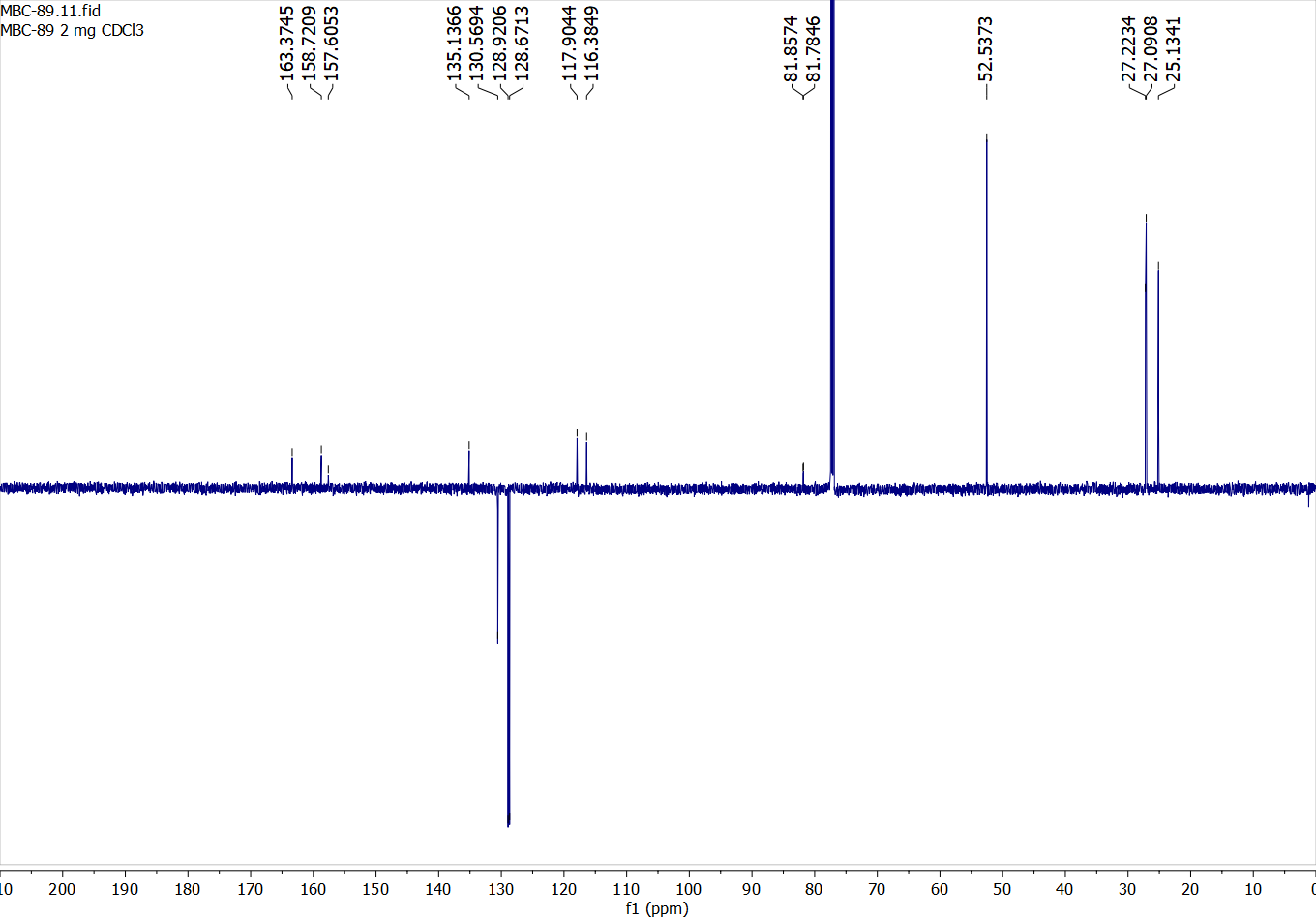

Compound 7:

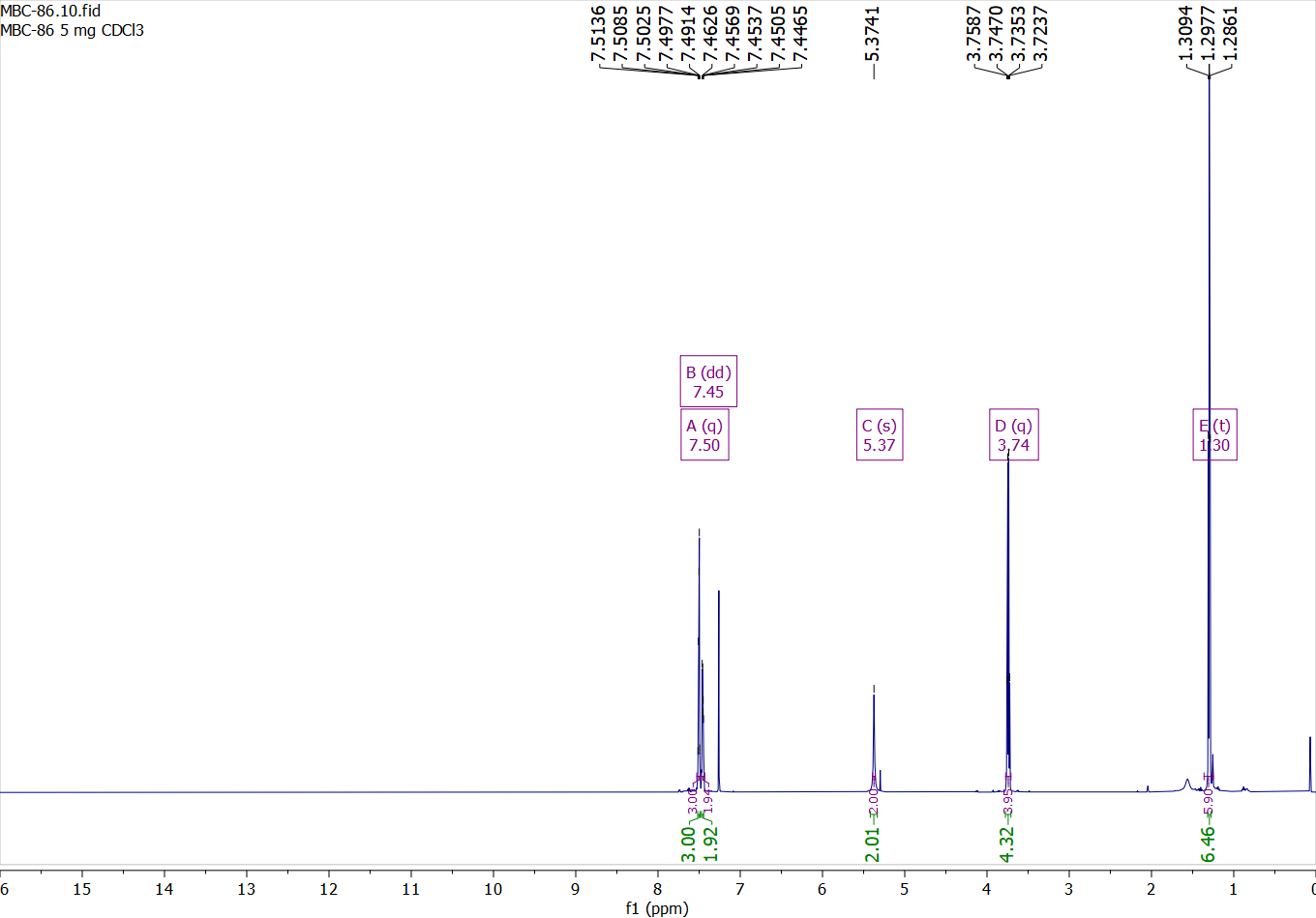

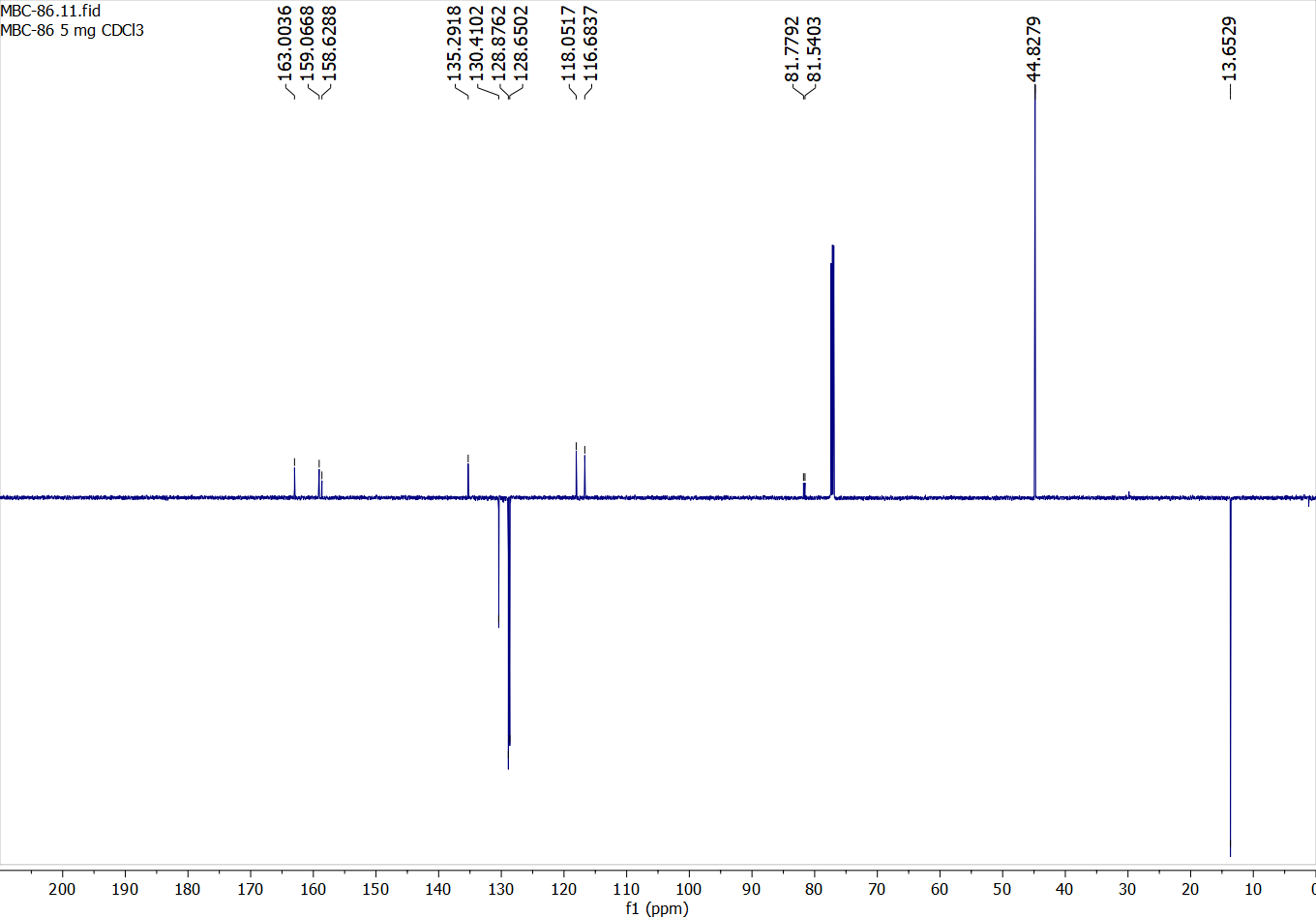

Compound 8:

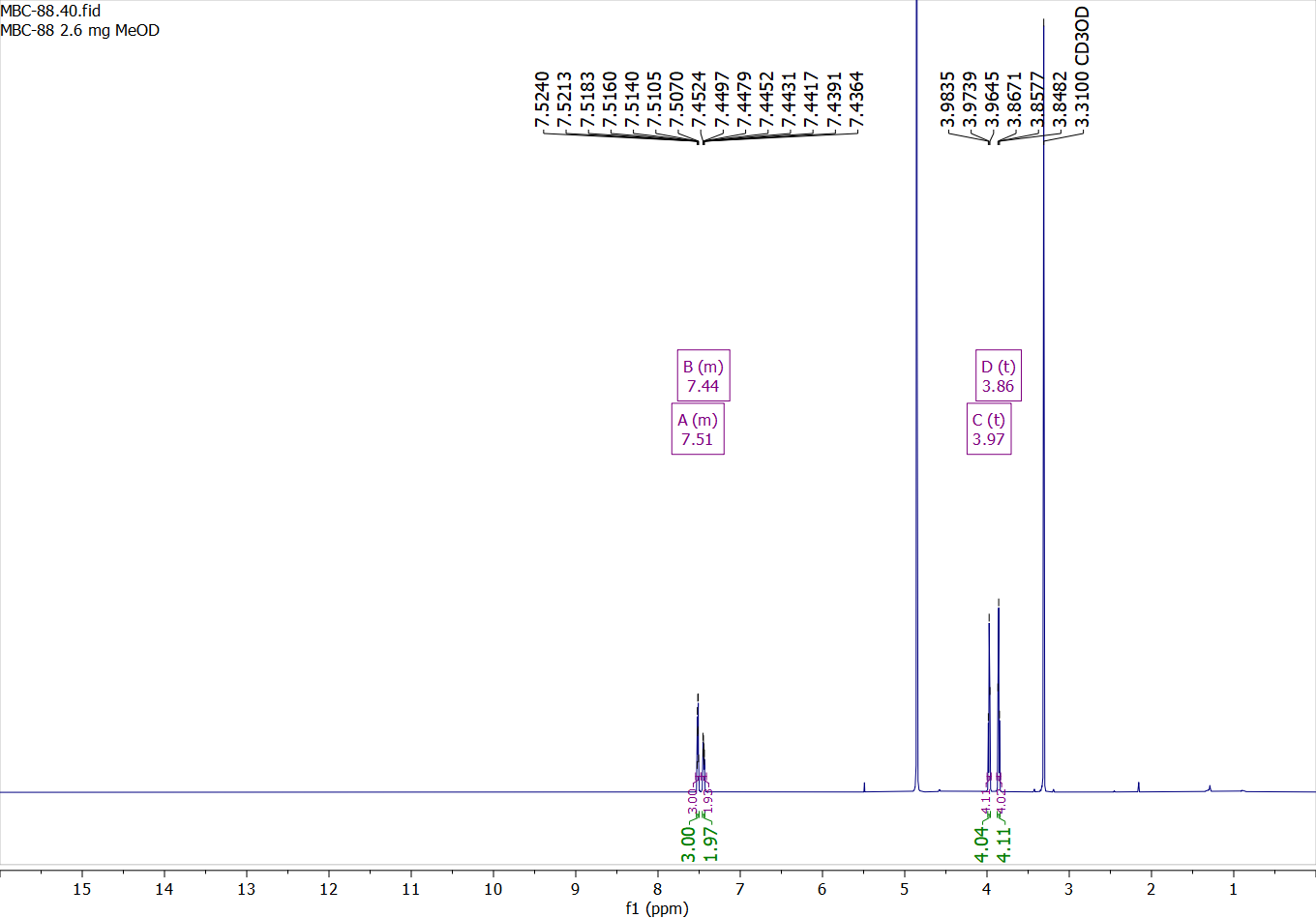

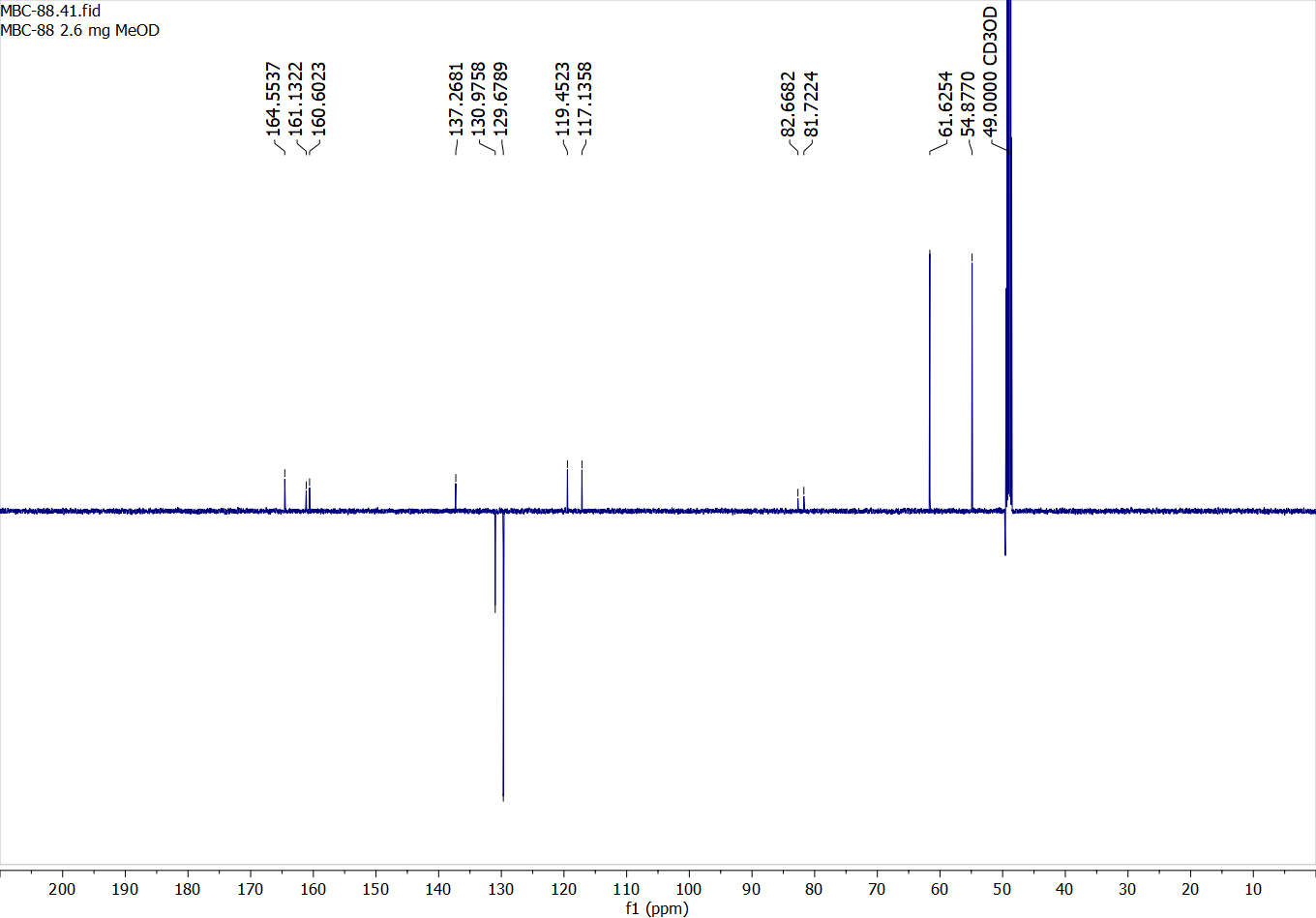

Compound 11:

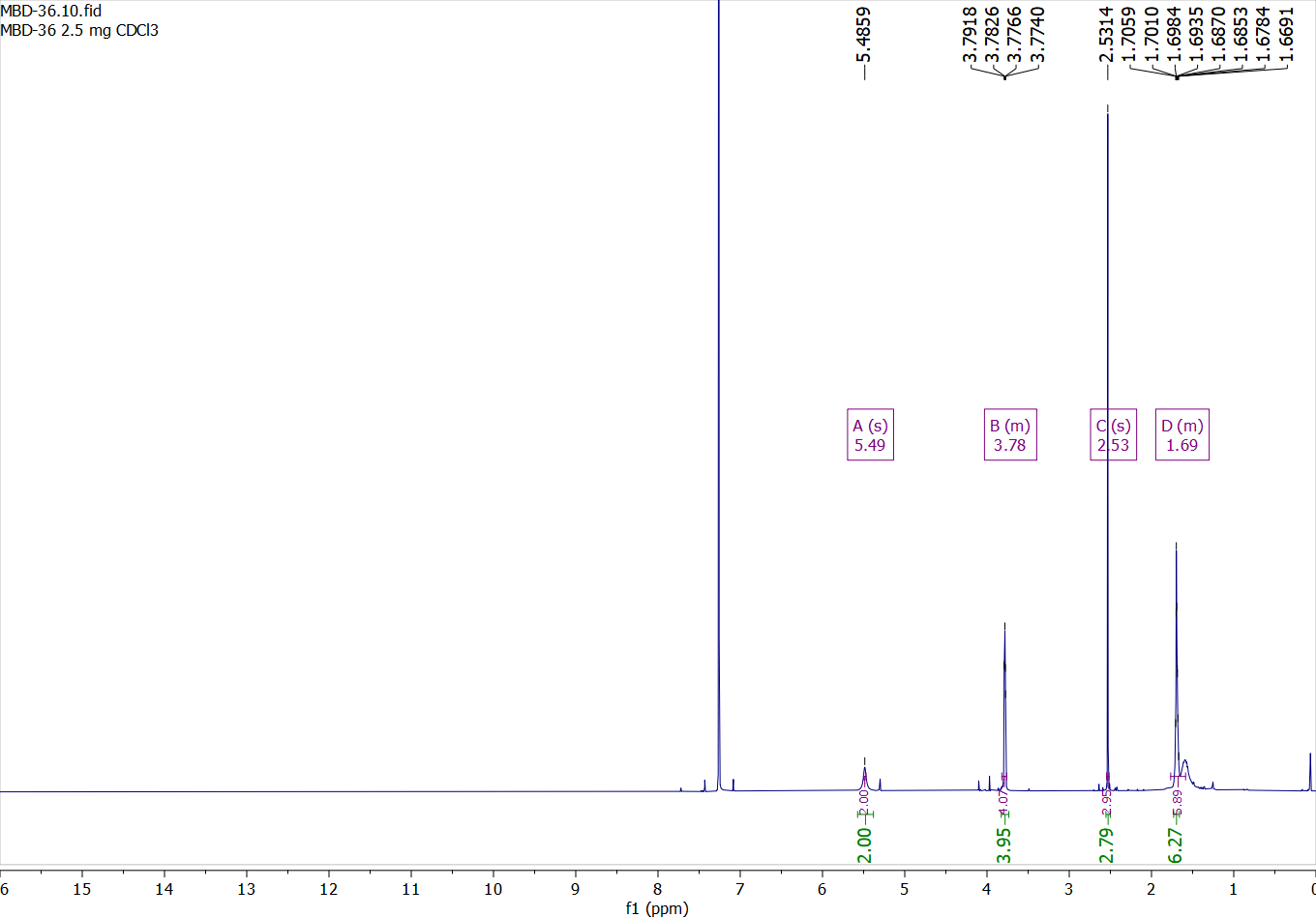

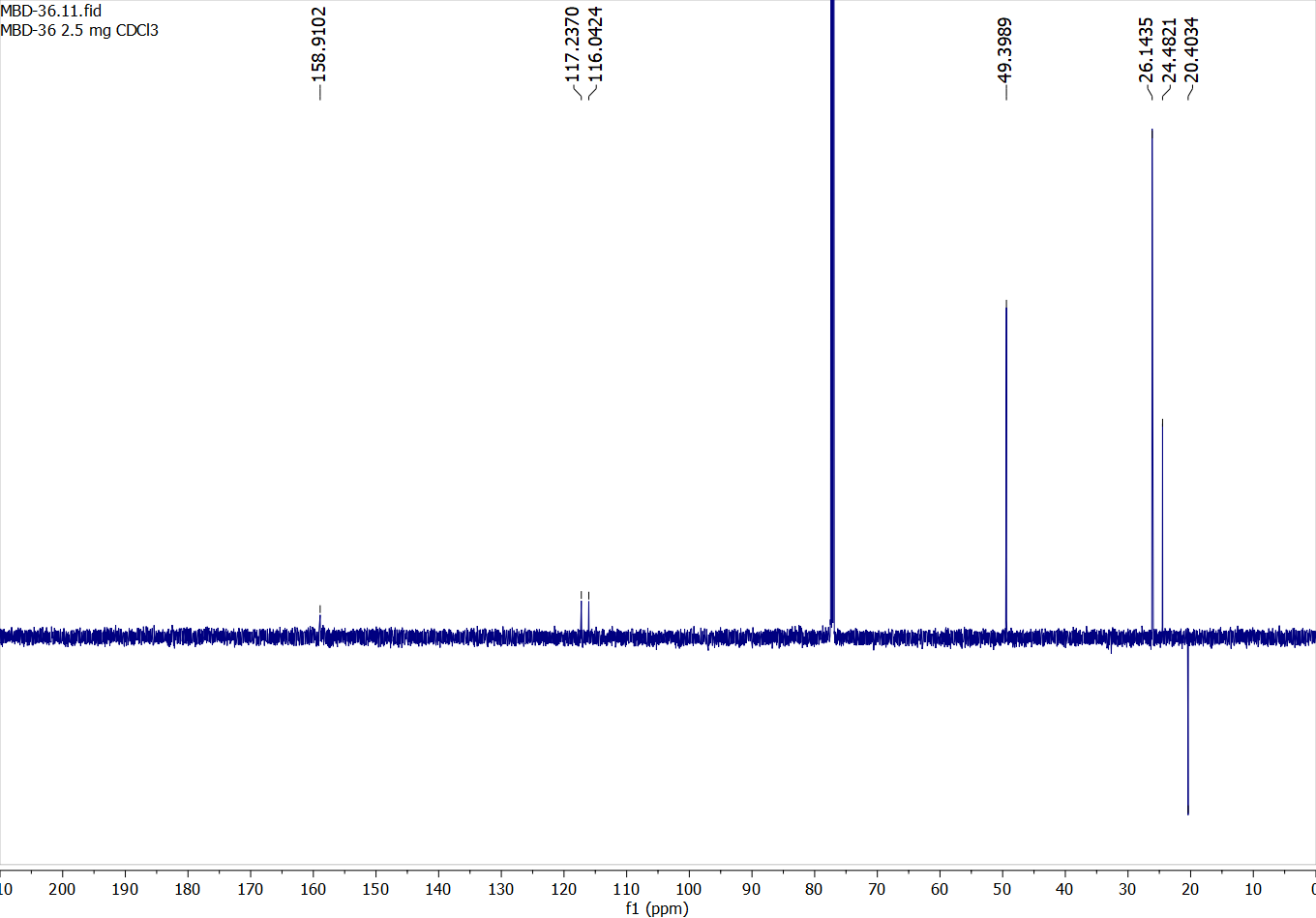

Compound 12:

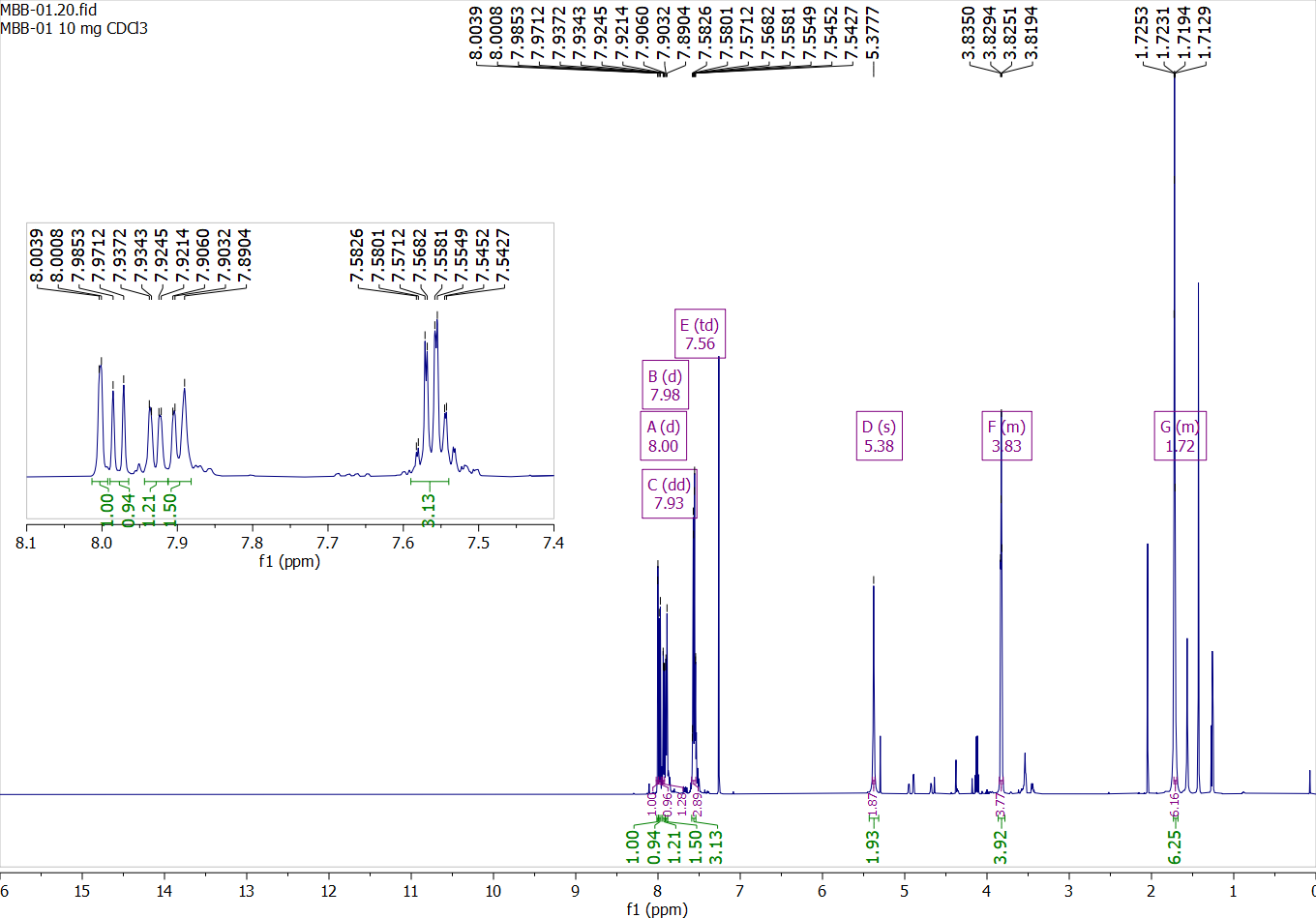

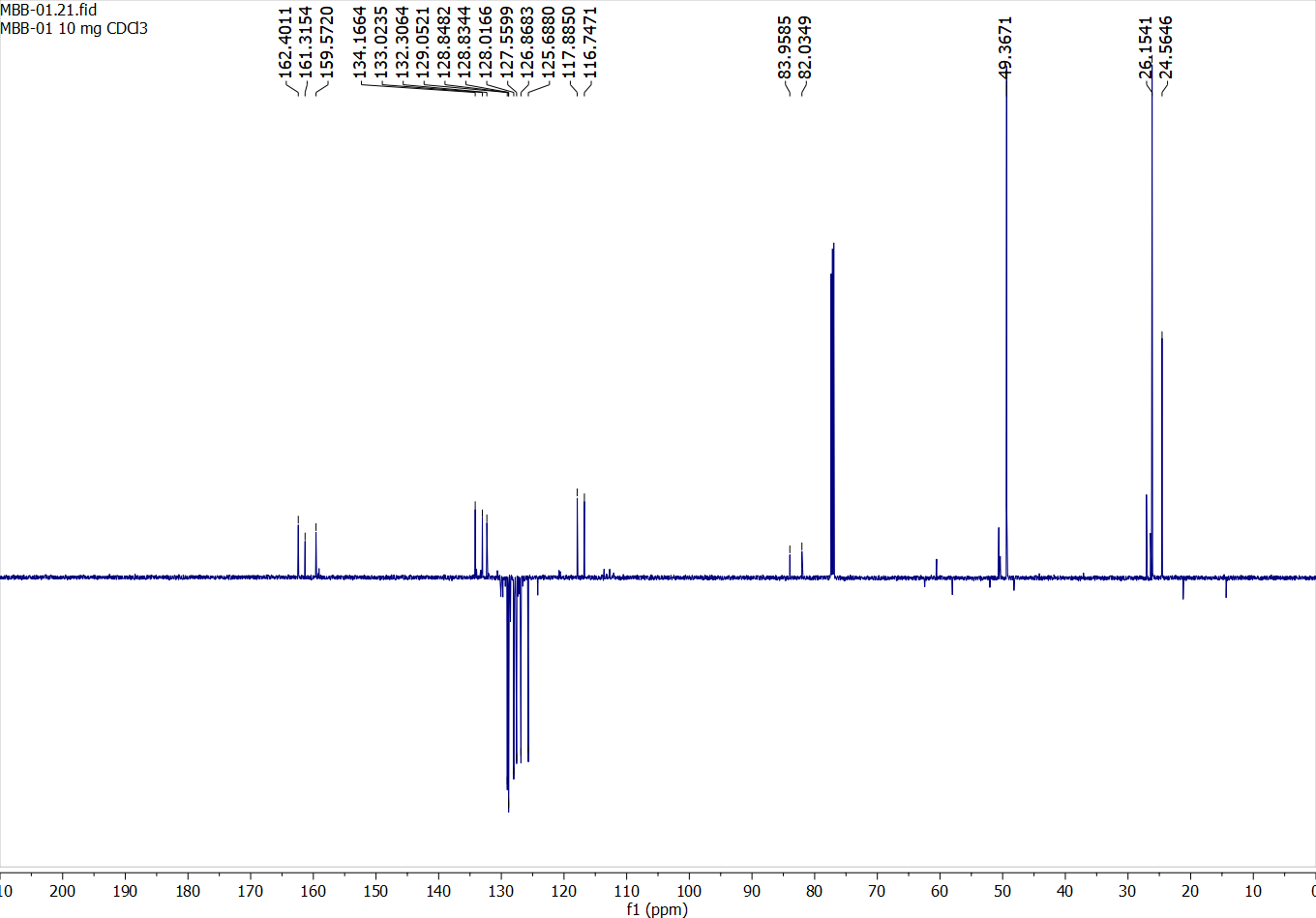

Compound 13:

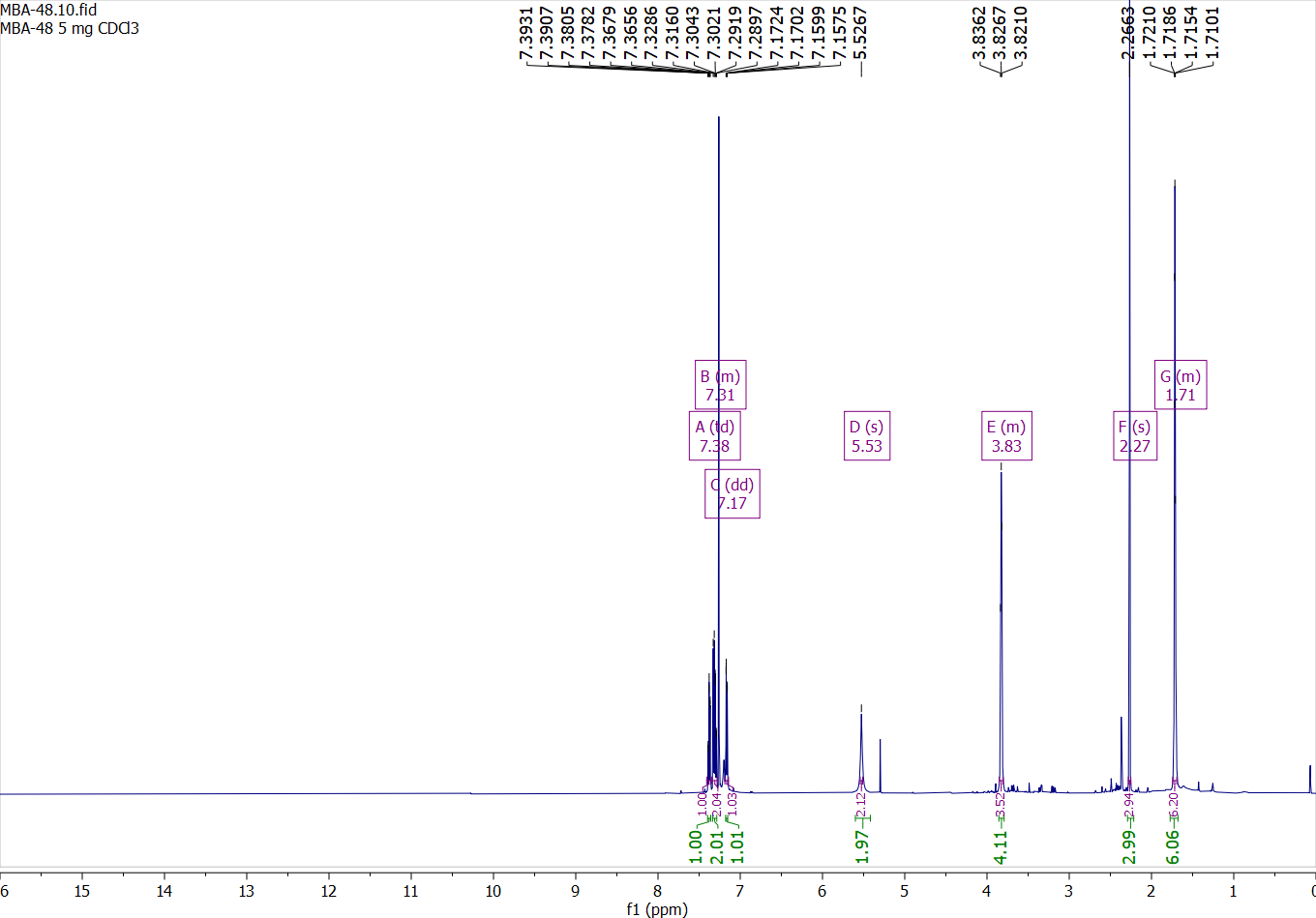

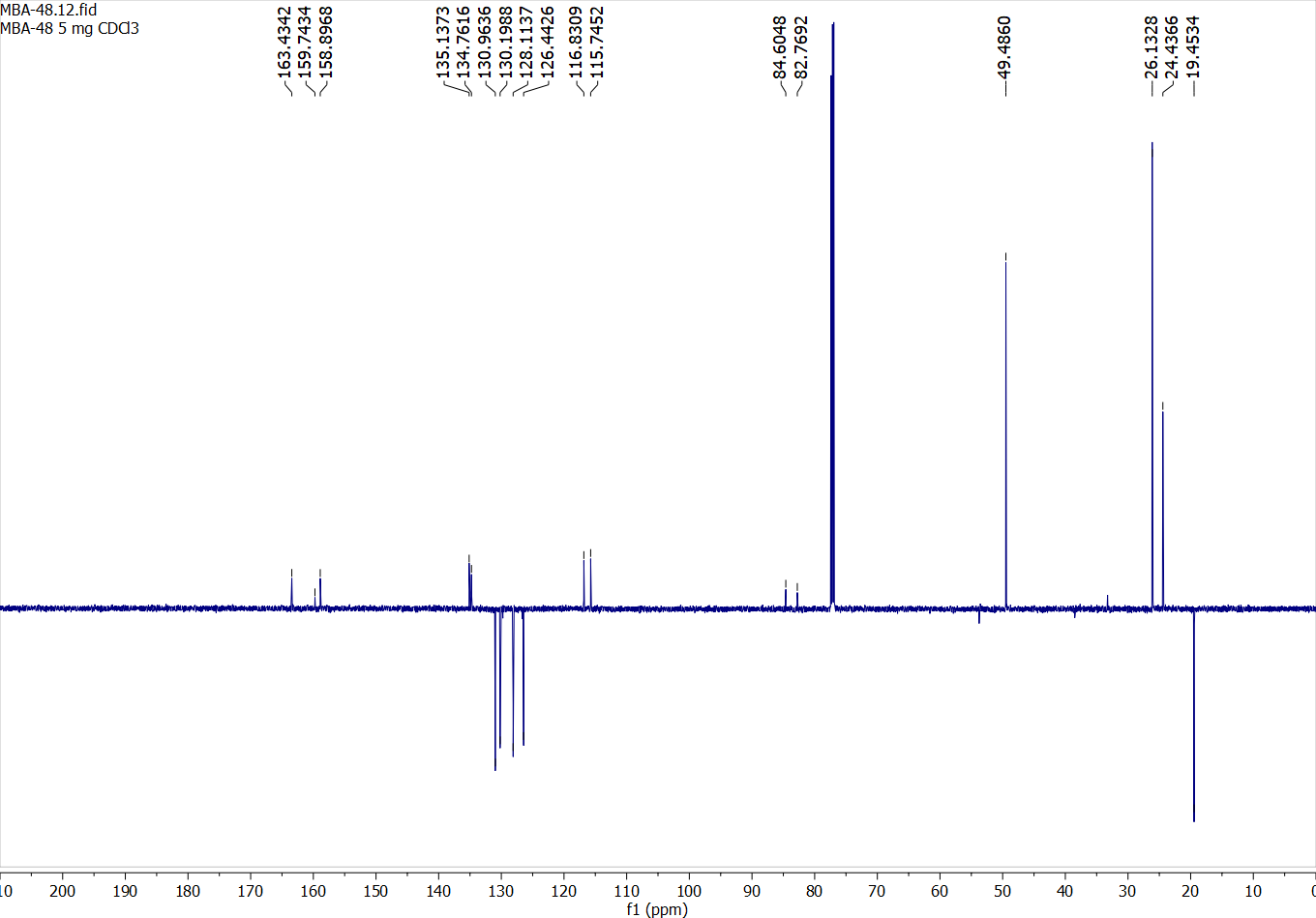

Compound 14:

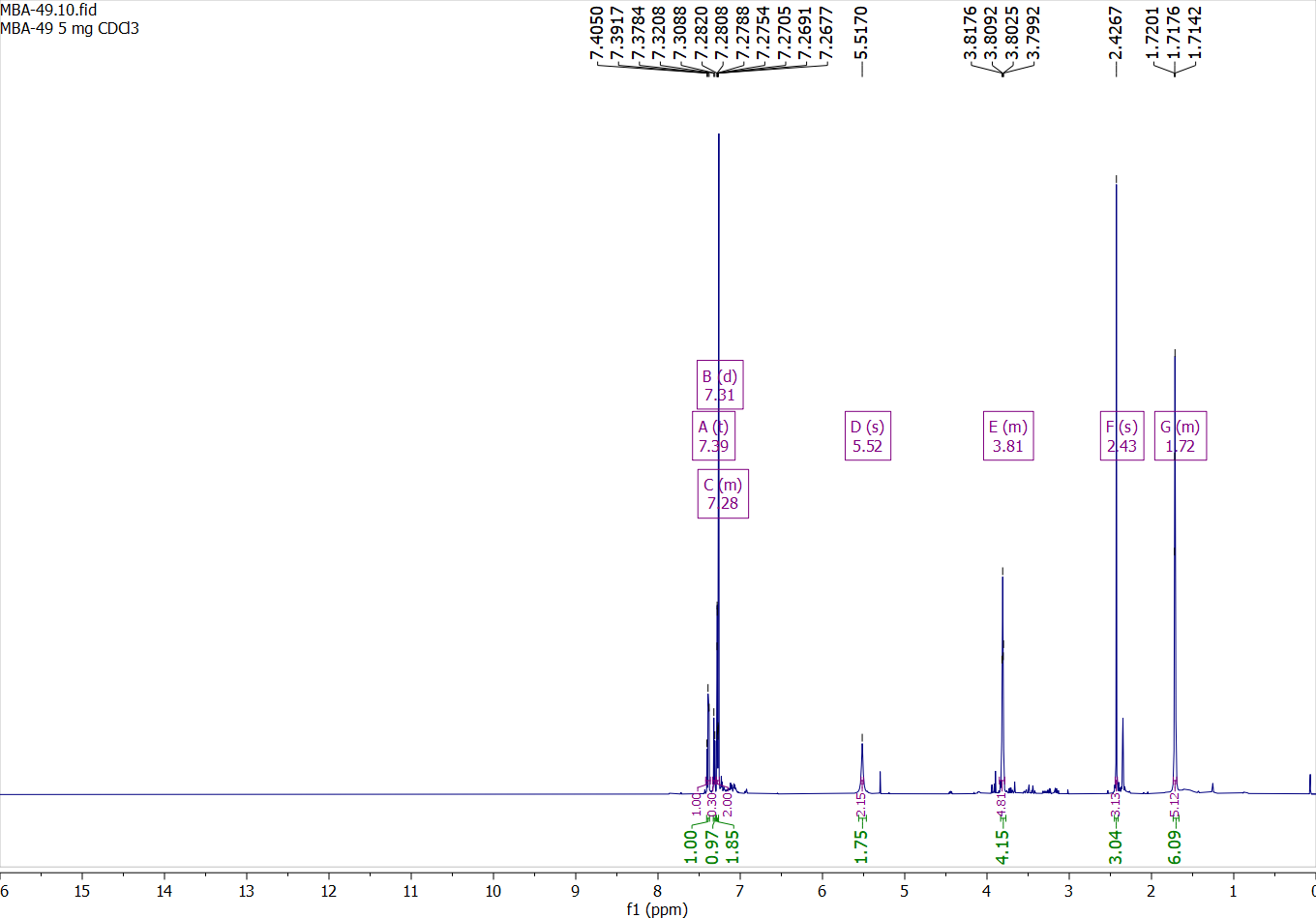

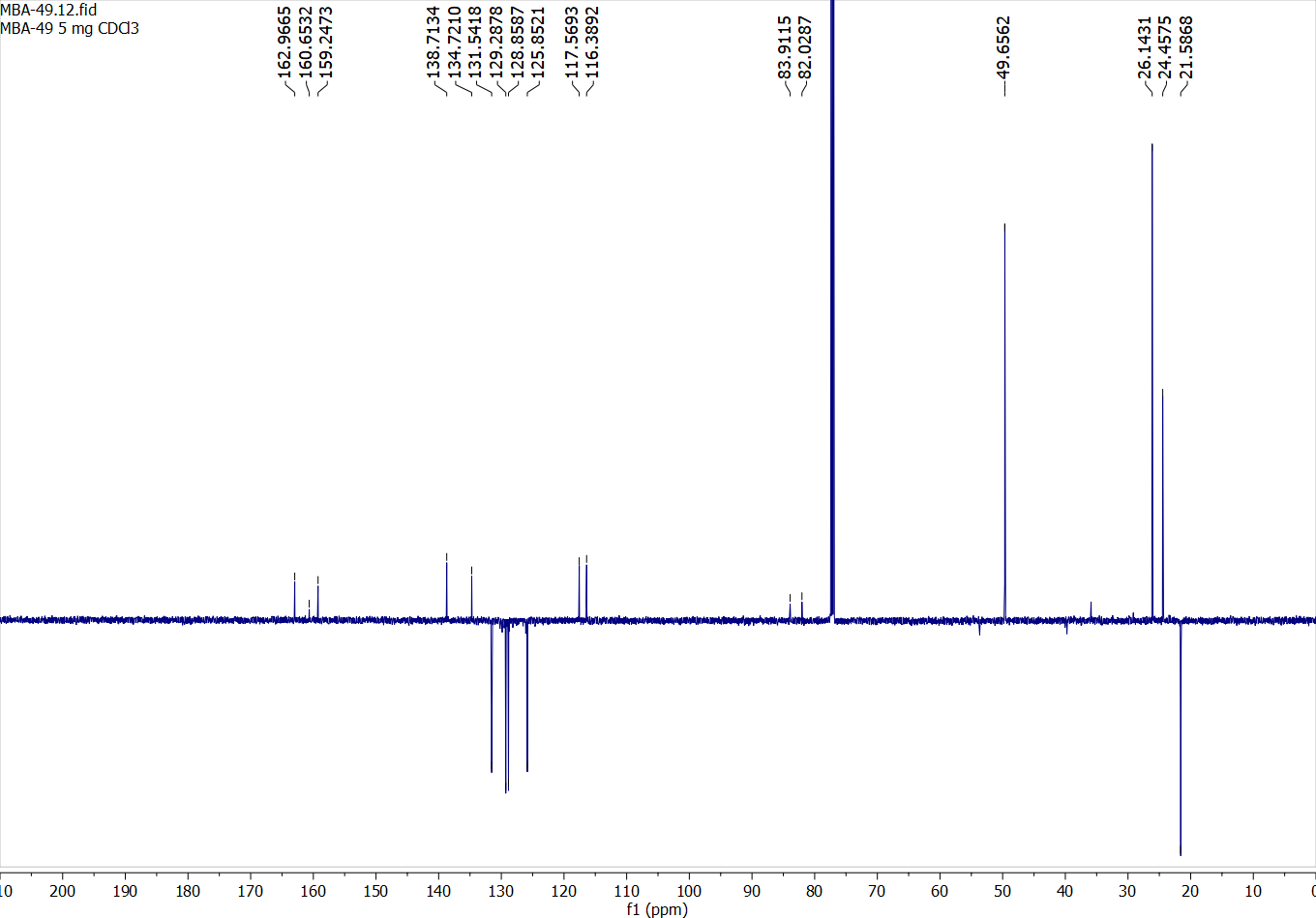

Compound 15:

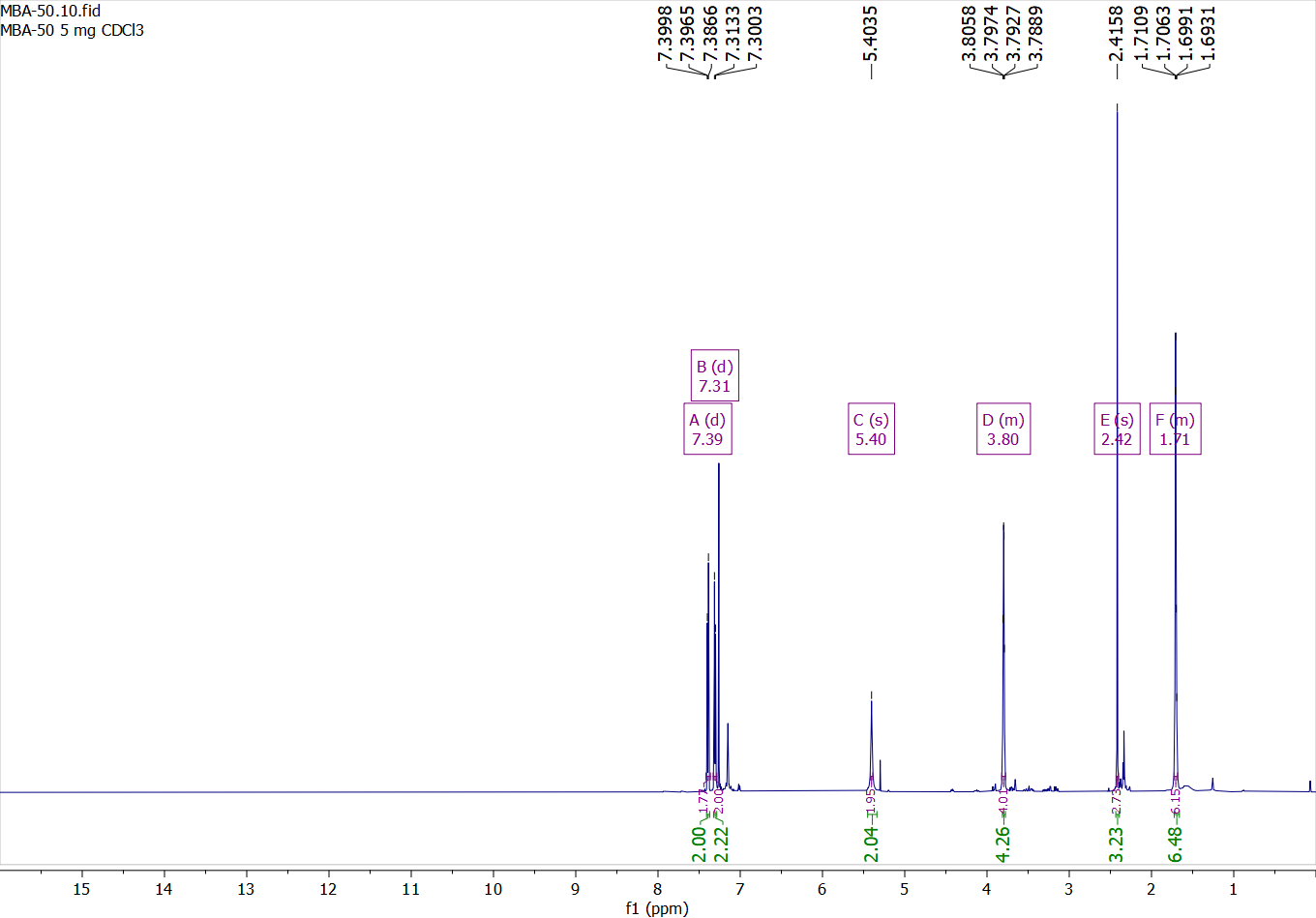

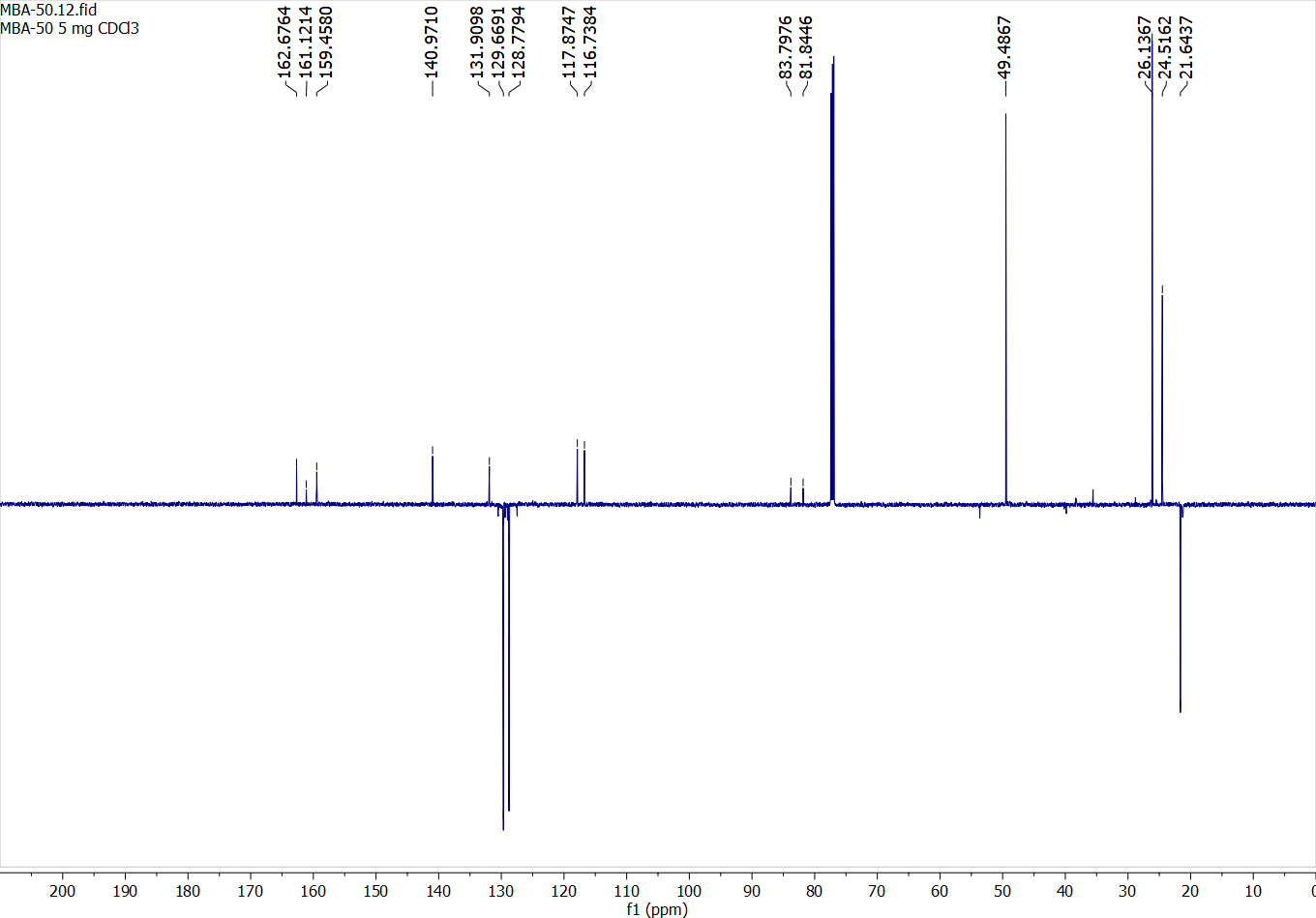

Compound 16:

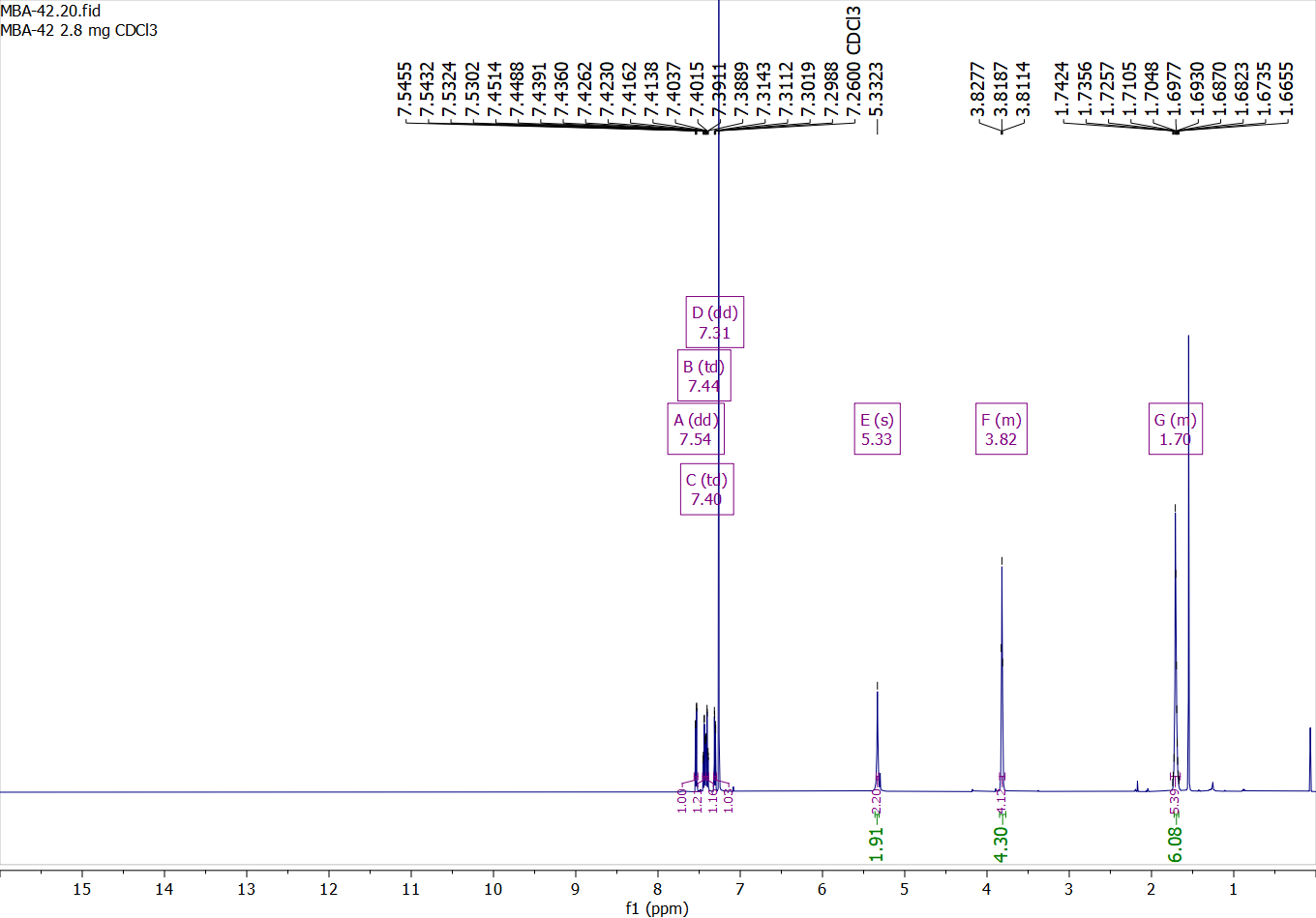

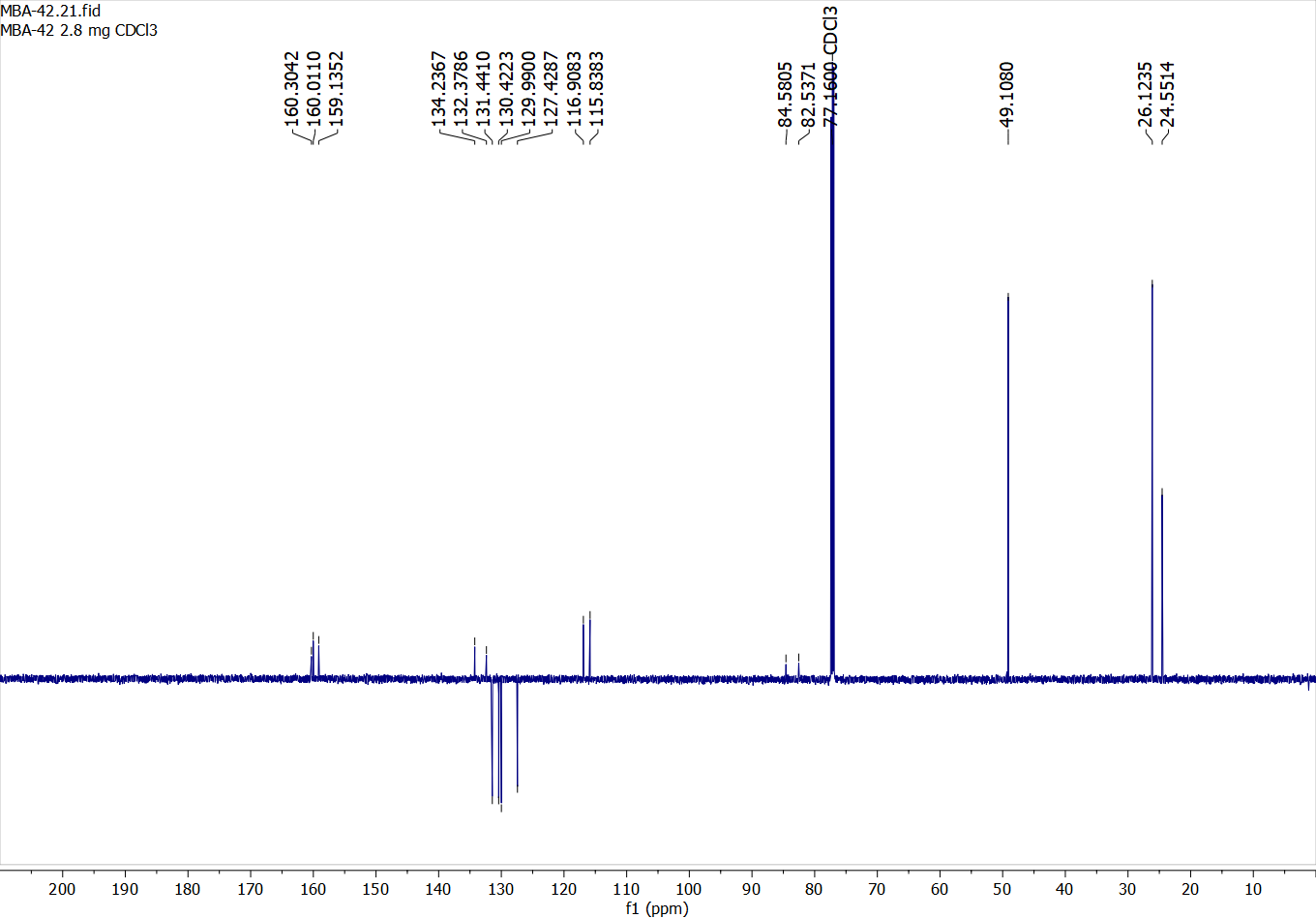

Compound 17:

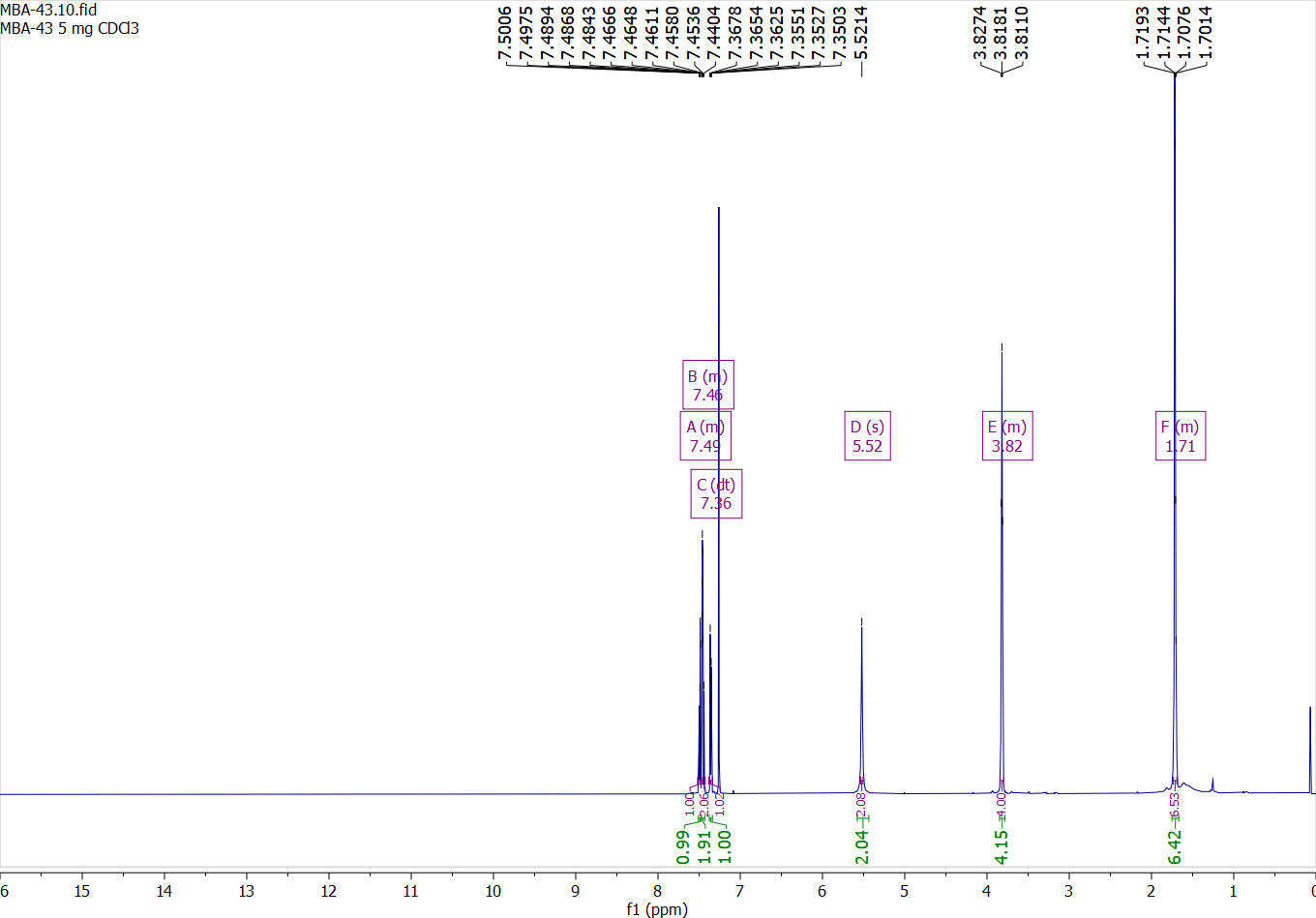

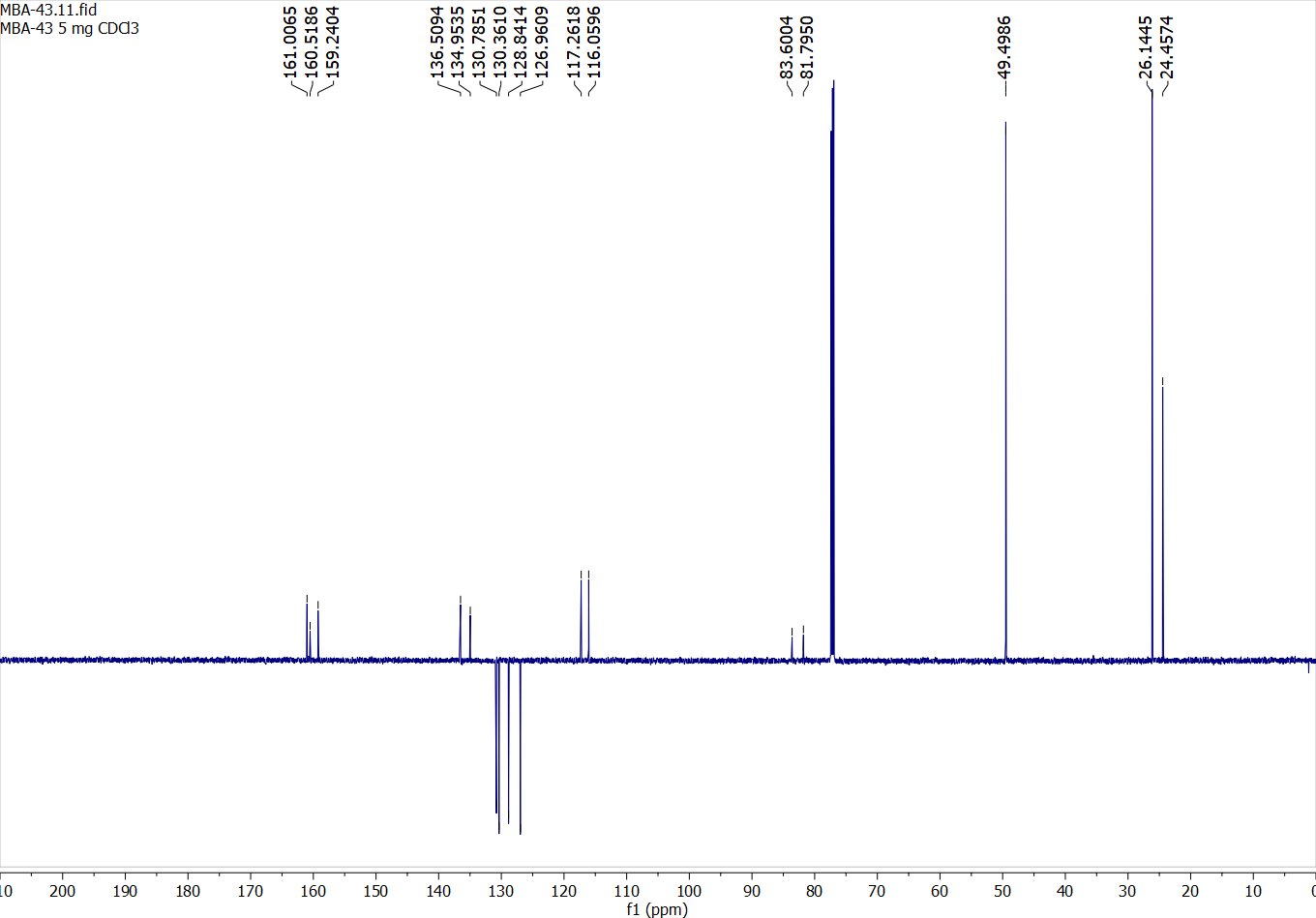

Compound 18:

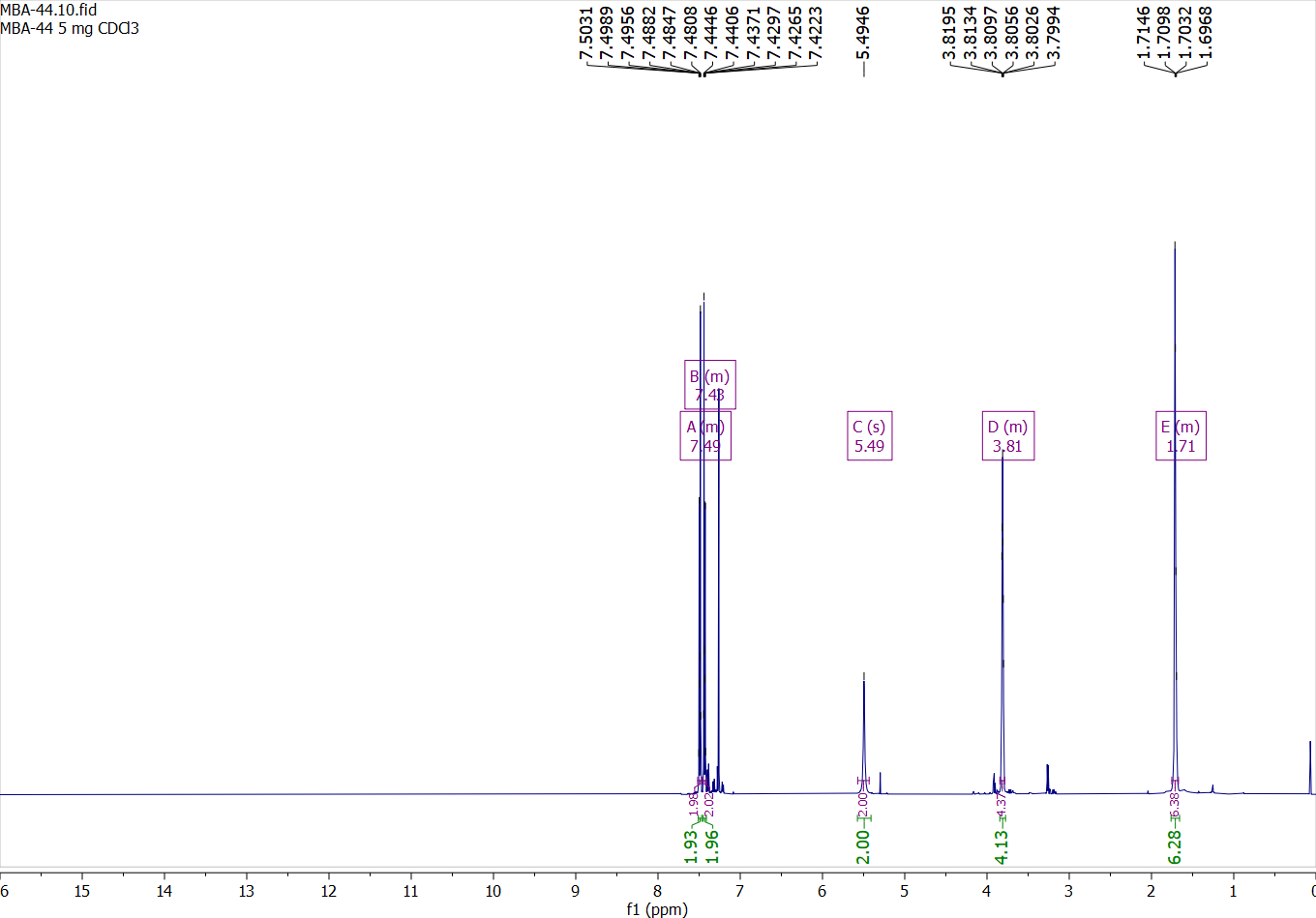

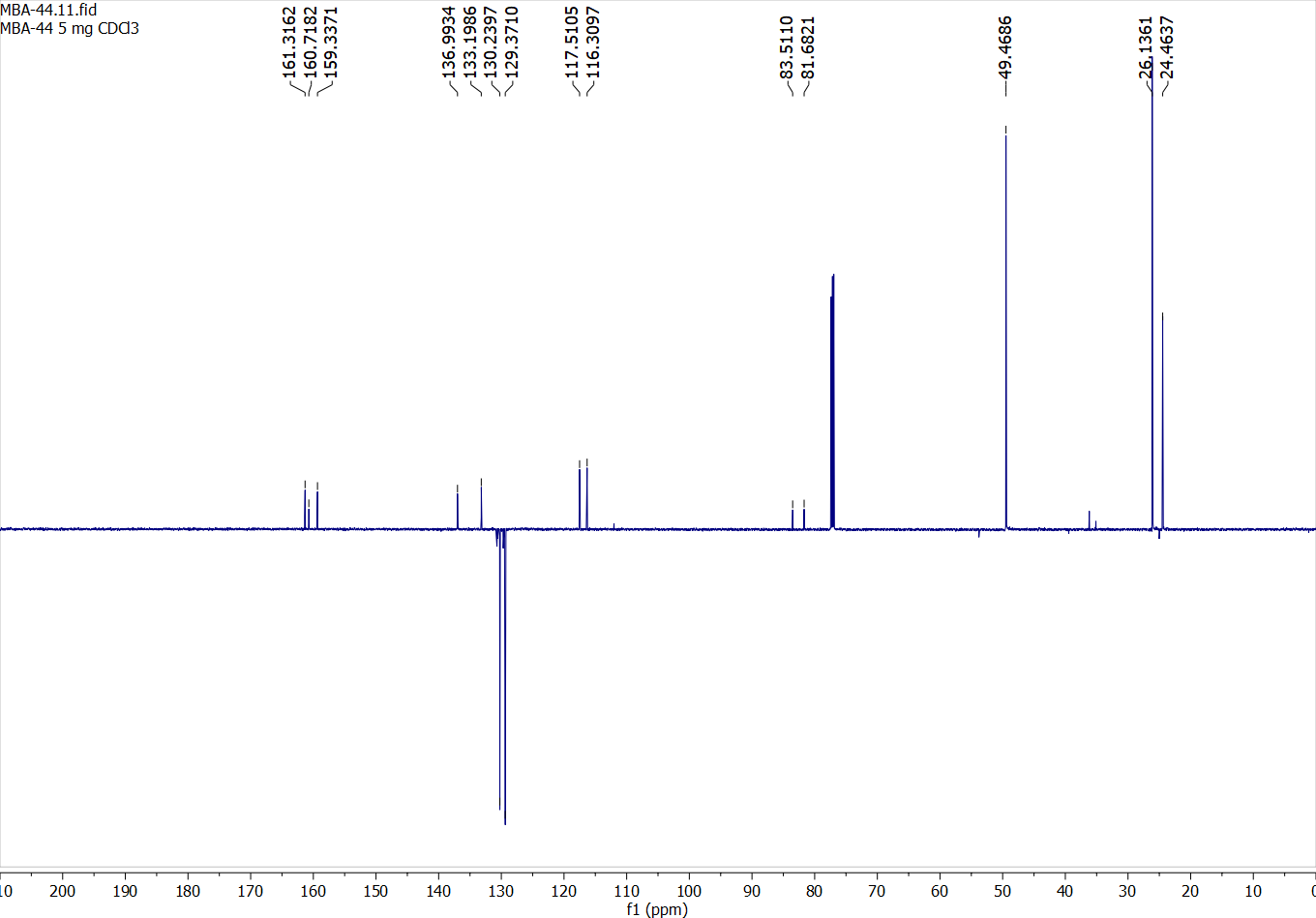

Compound 19:

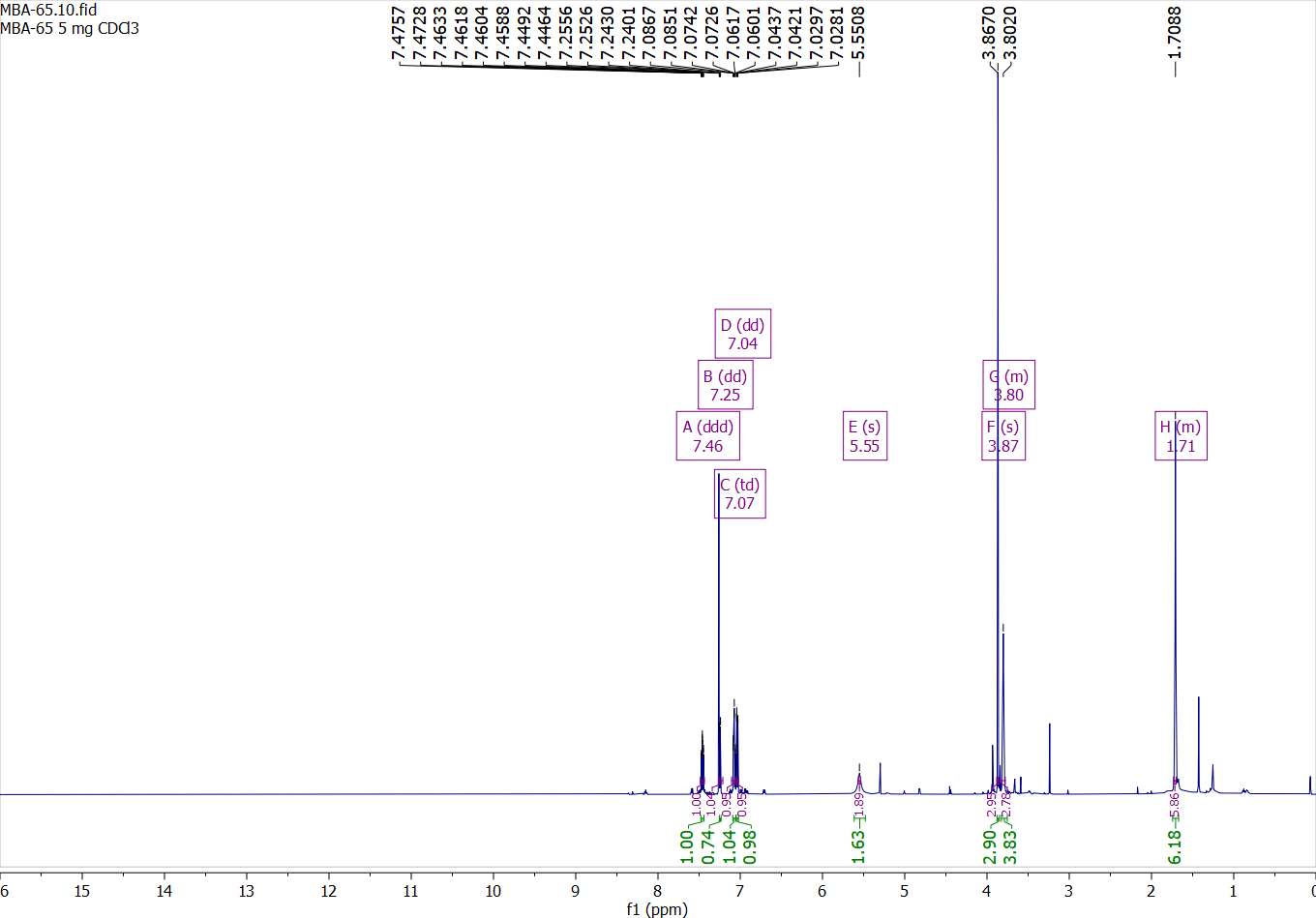

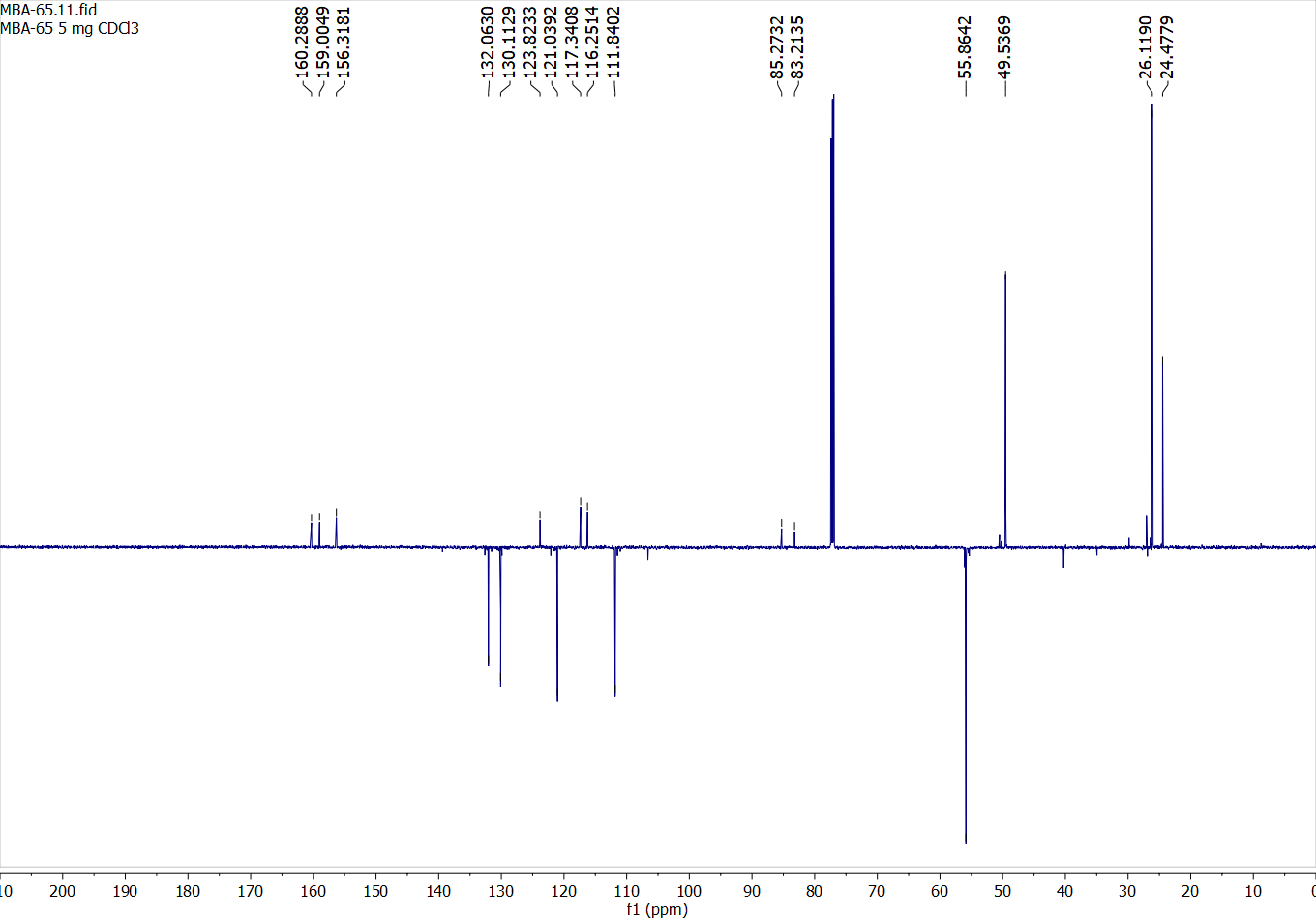

Compound 20:

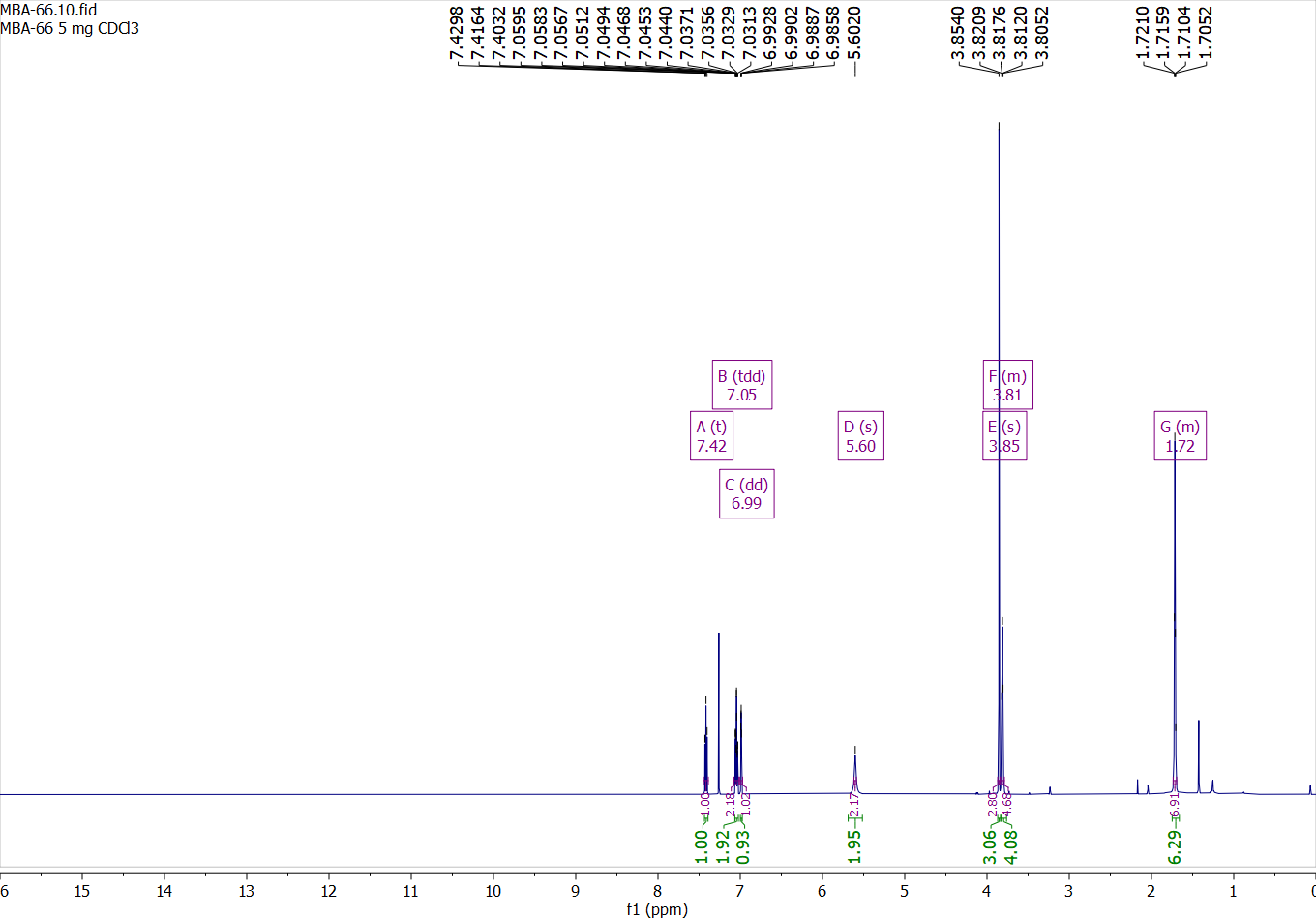

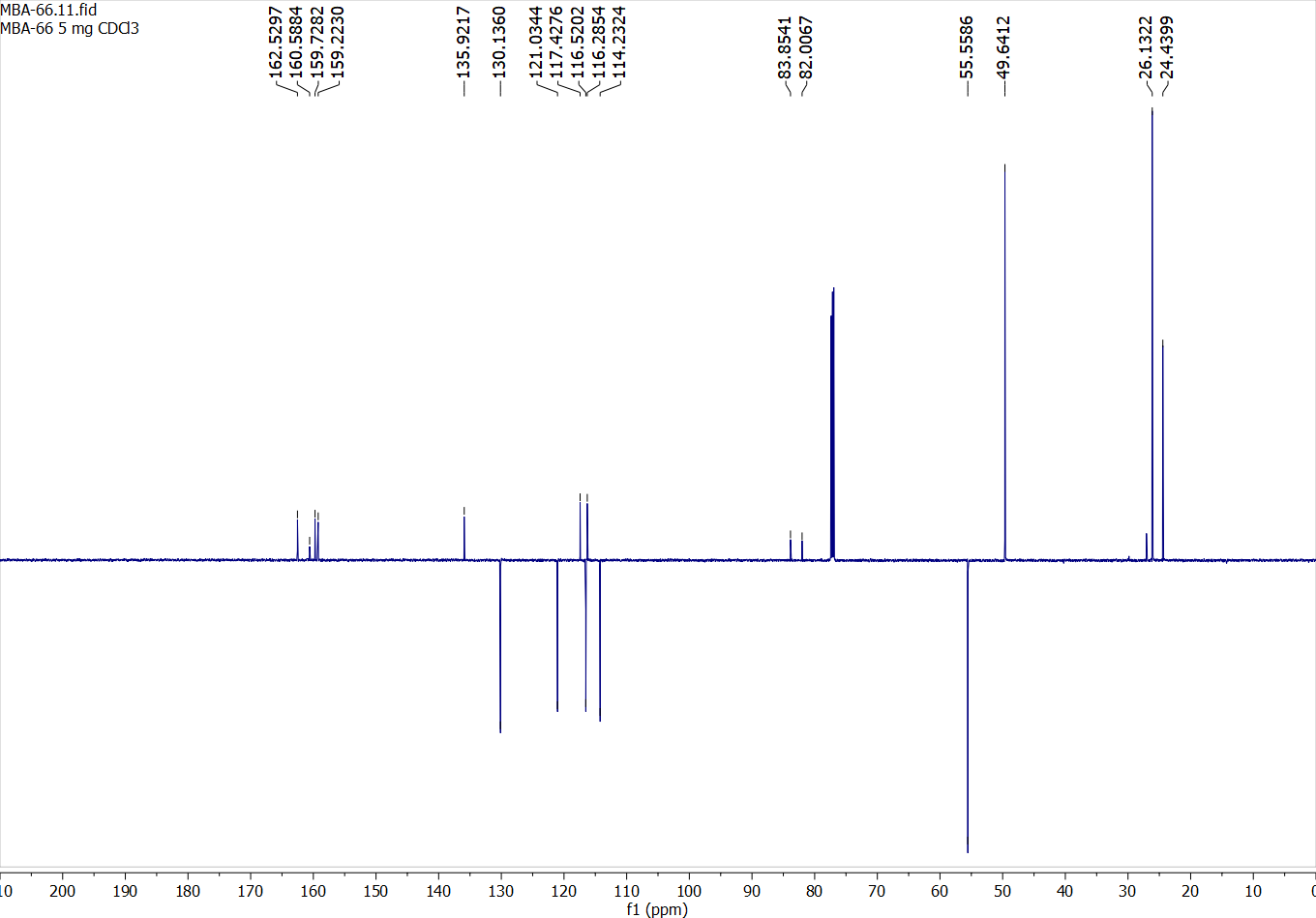

Compound 21:

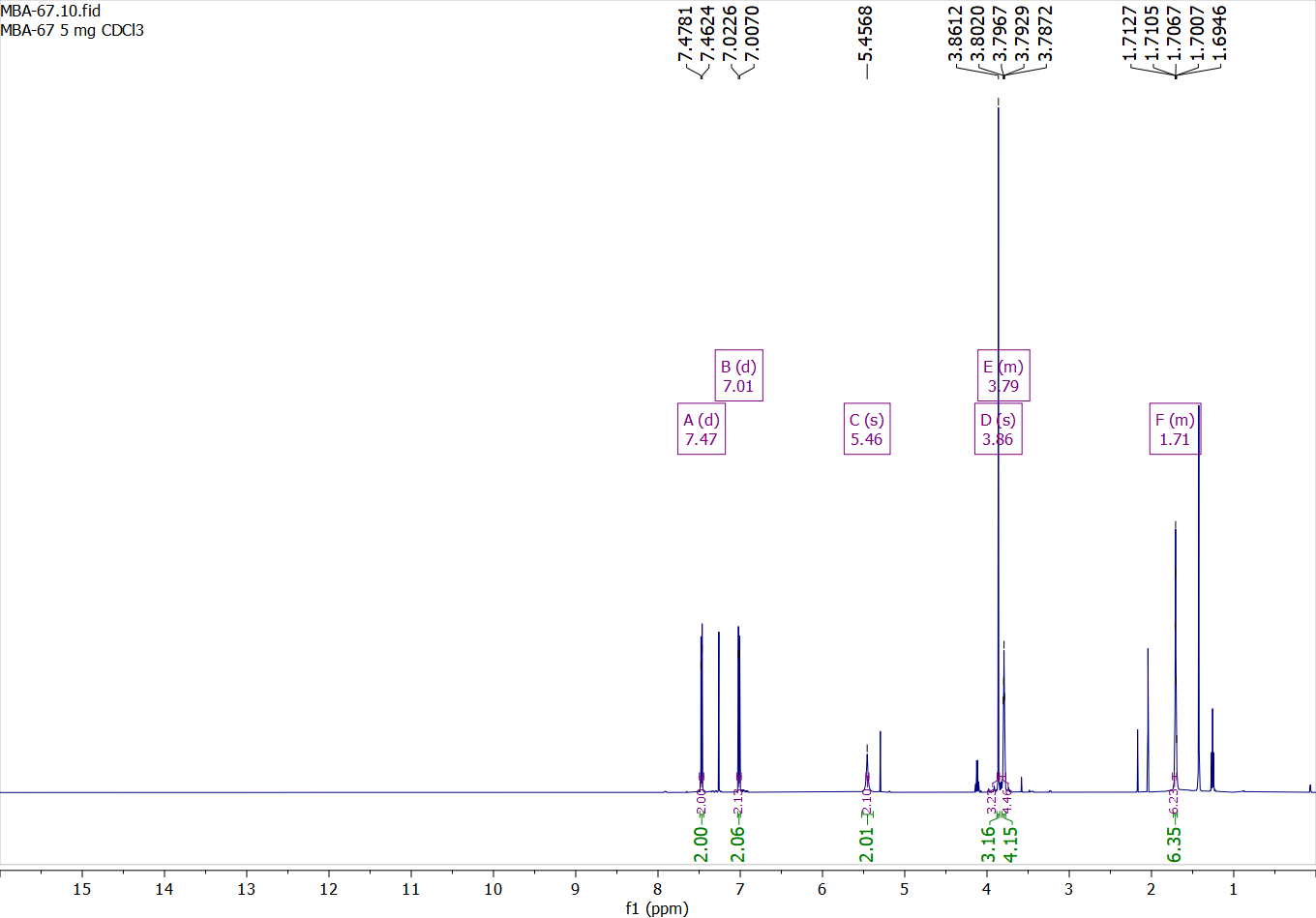

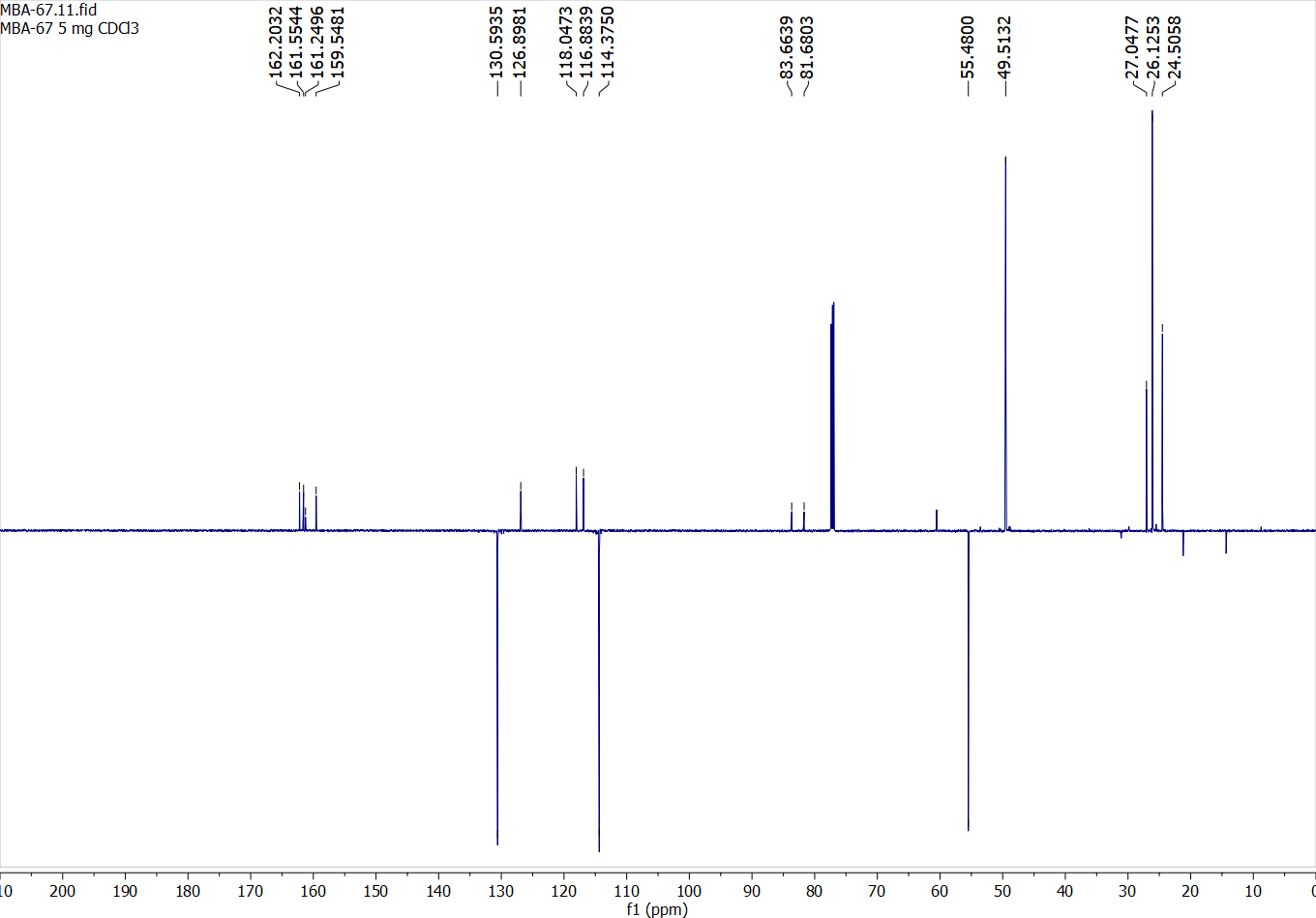

Compound 22:

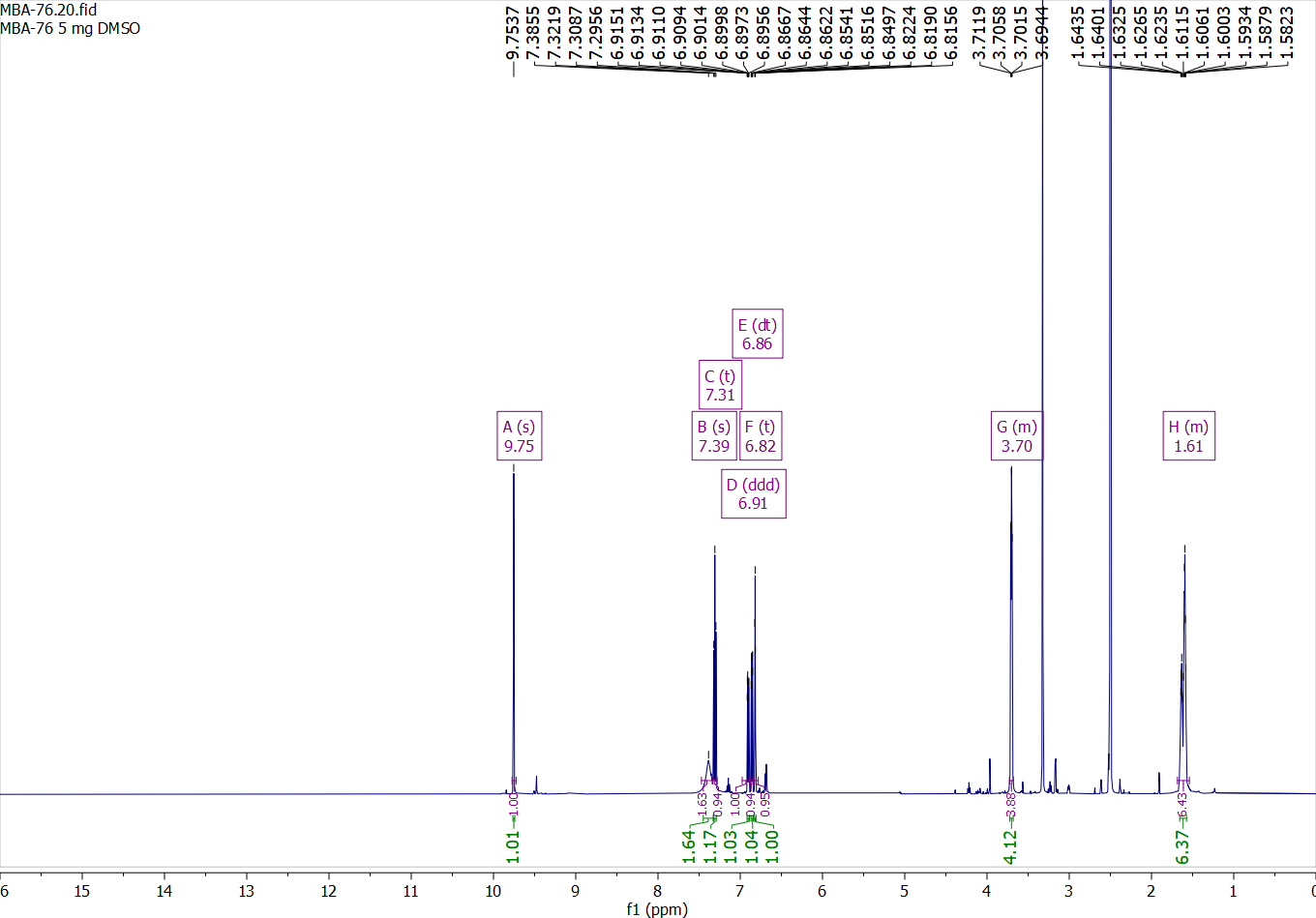

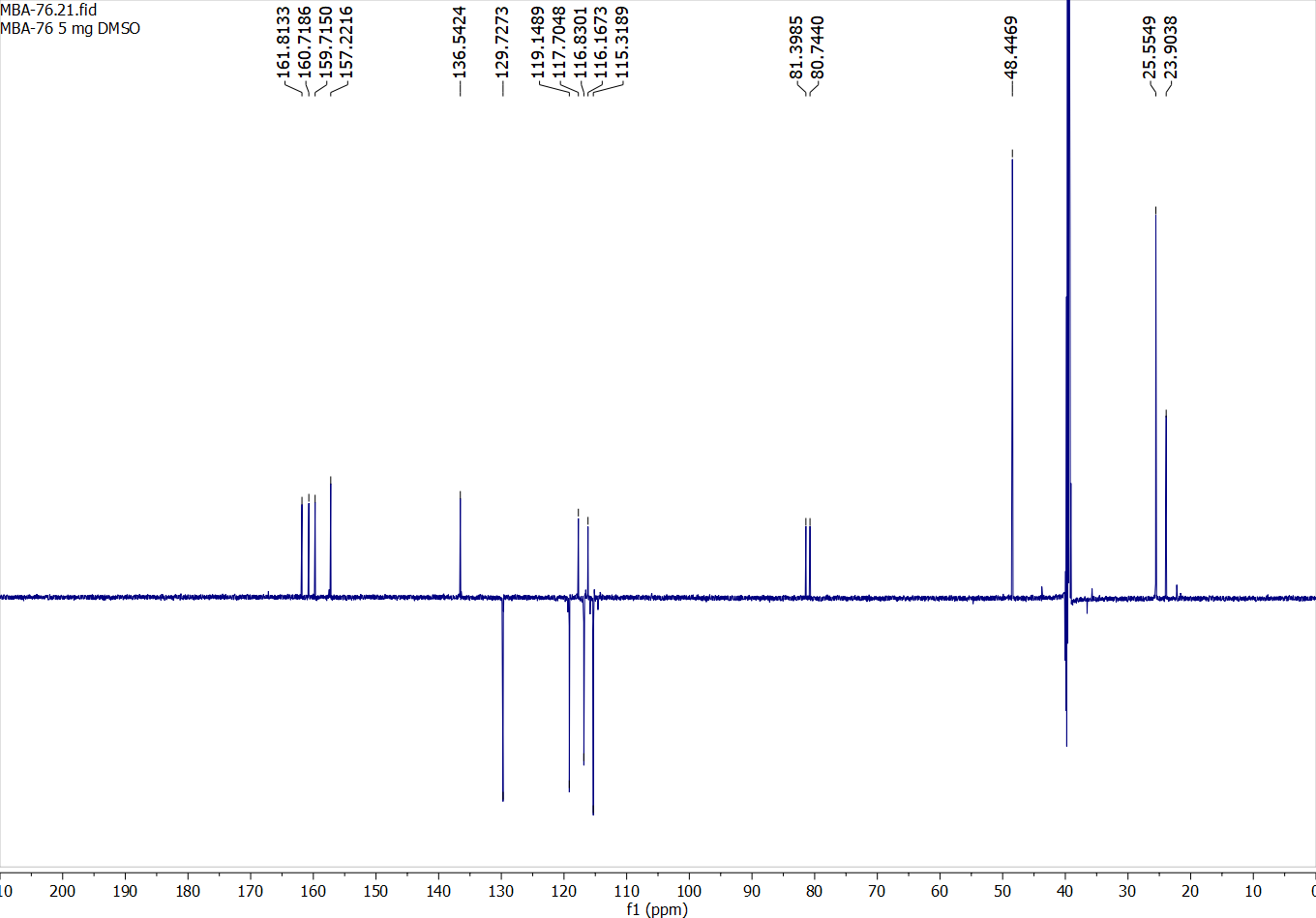

Compound 23:

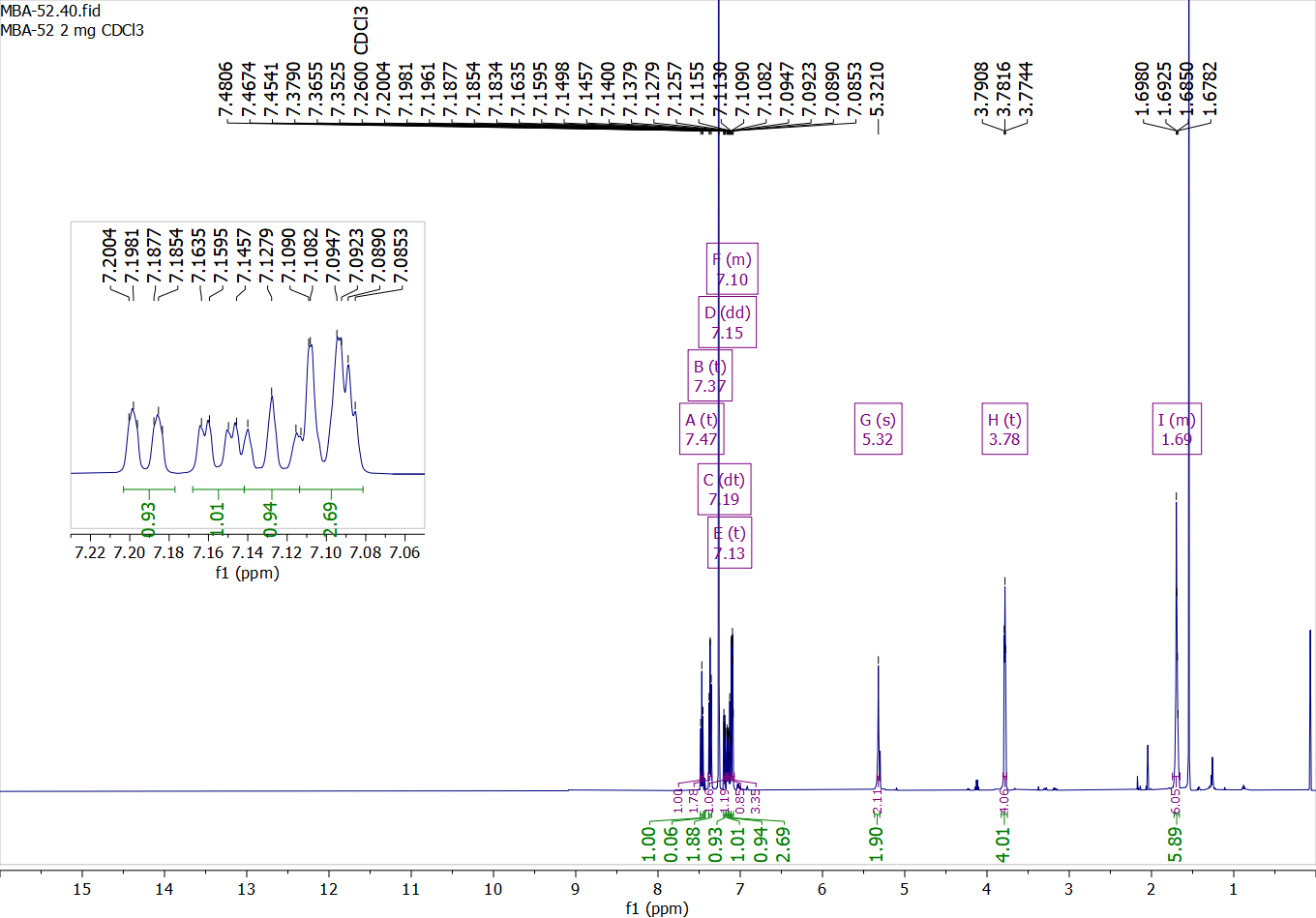

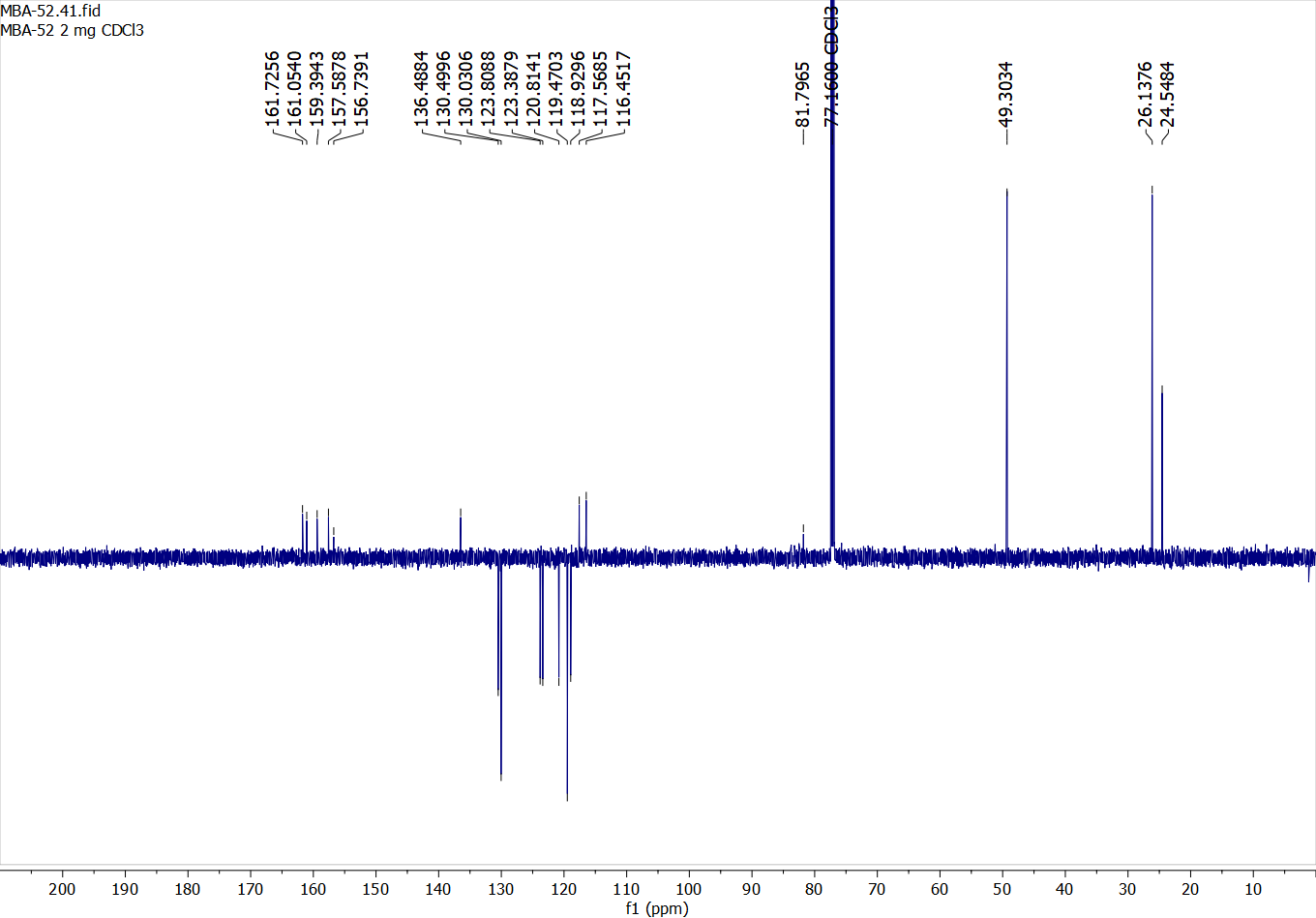

Compound 24:

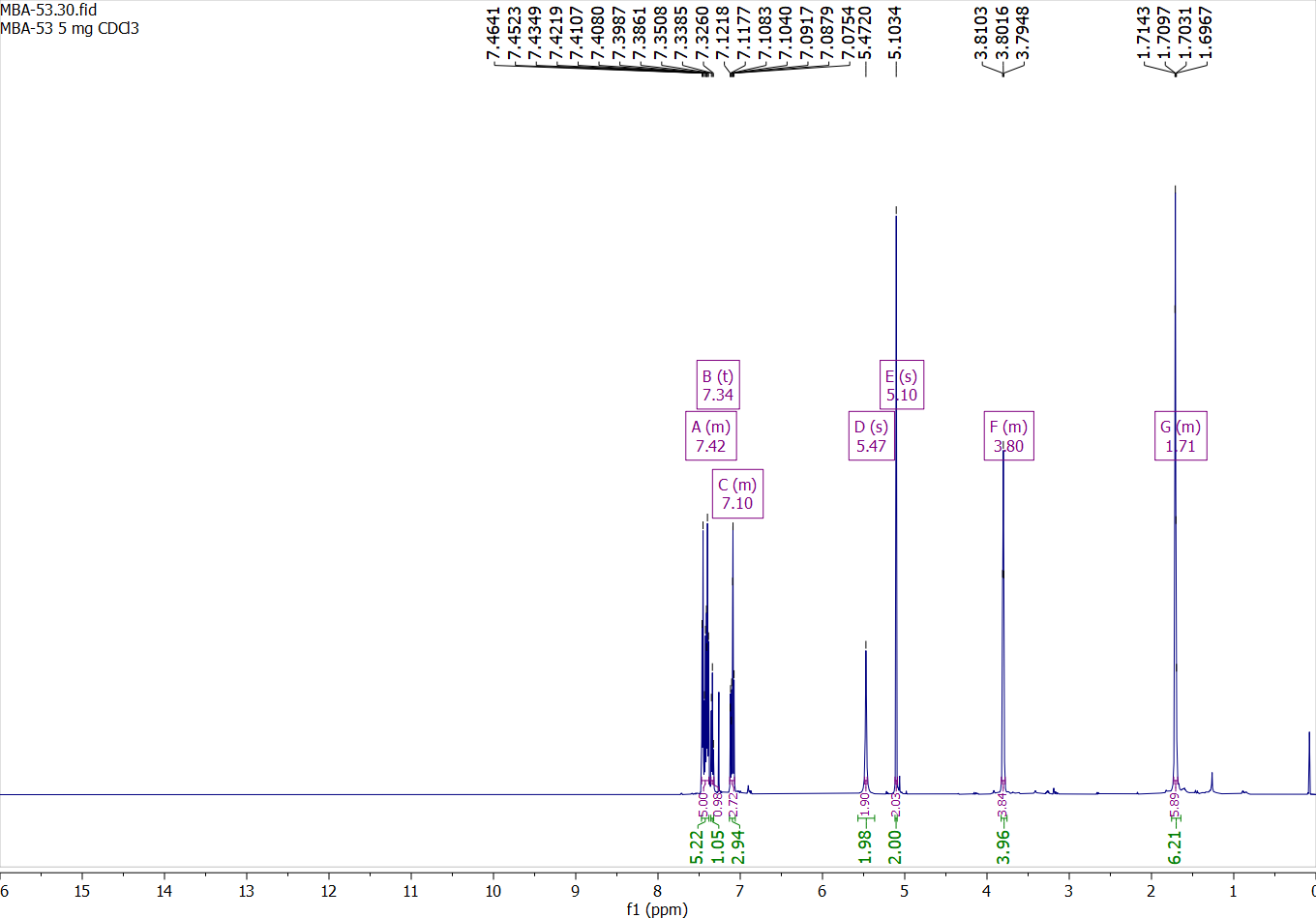

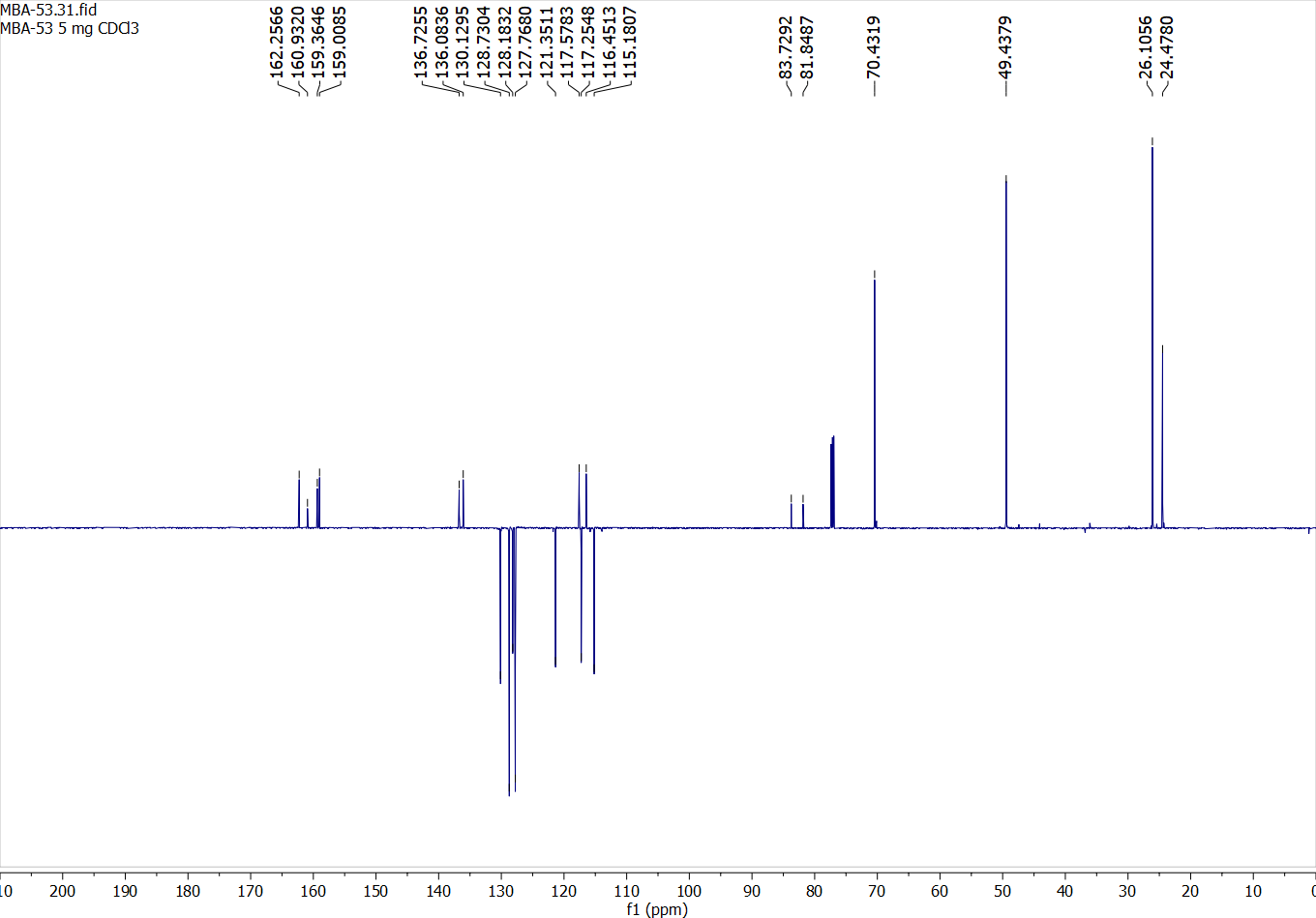

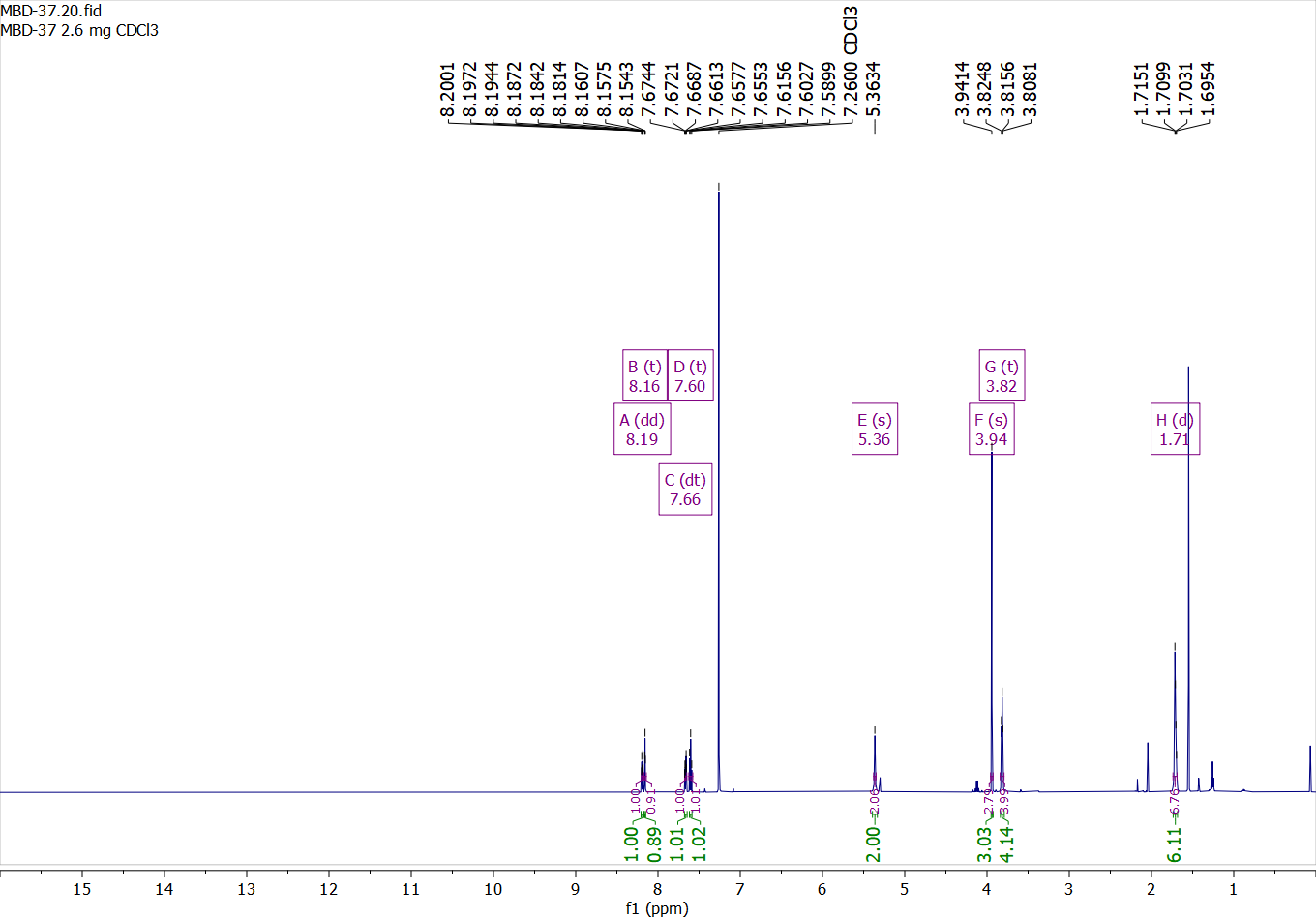
Compound 25:

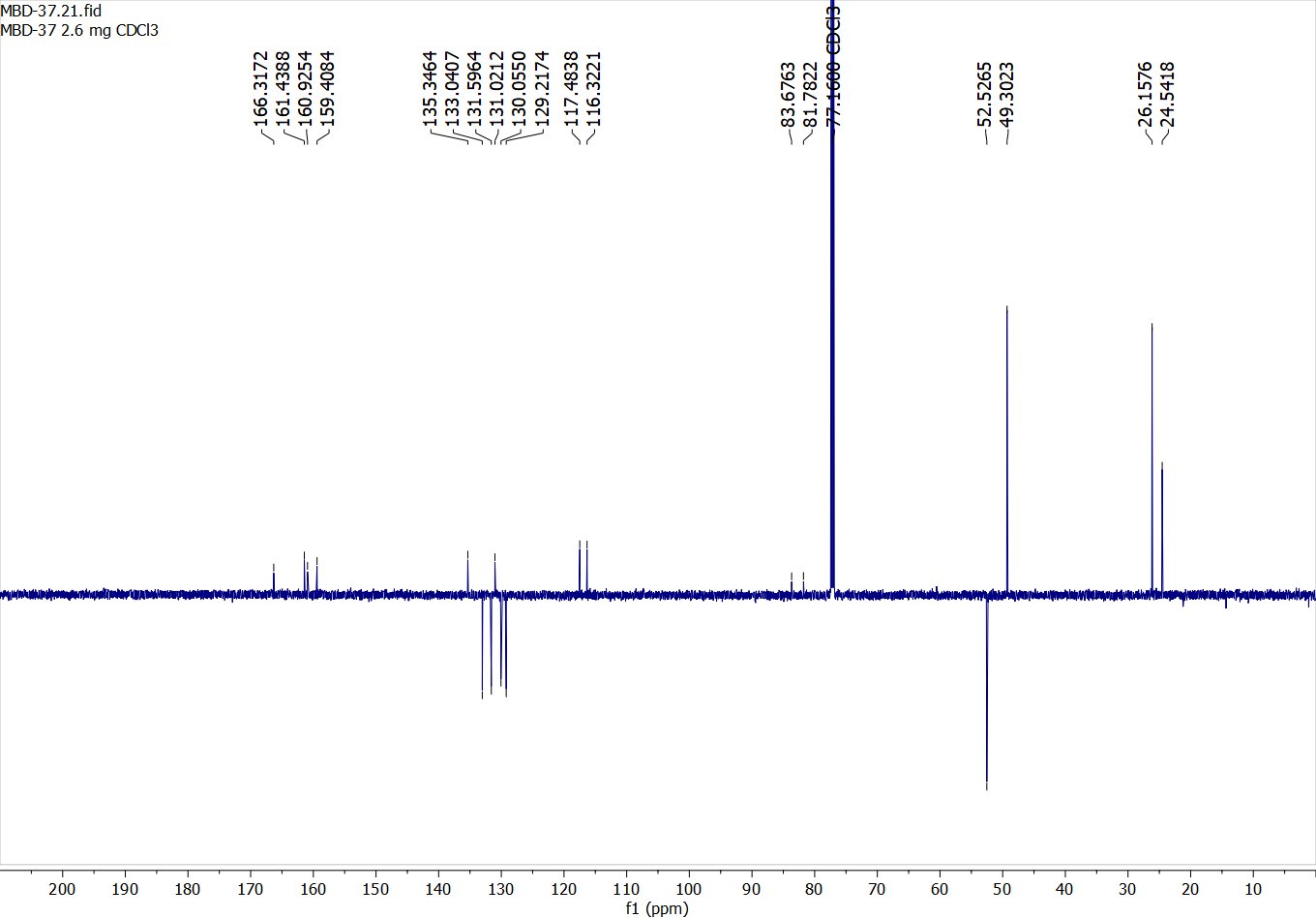

Compound 26:

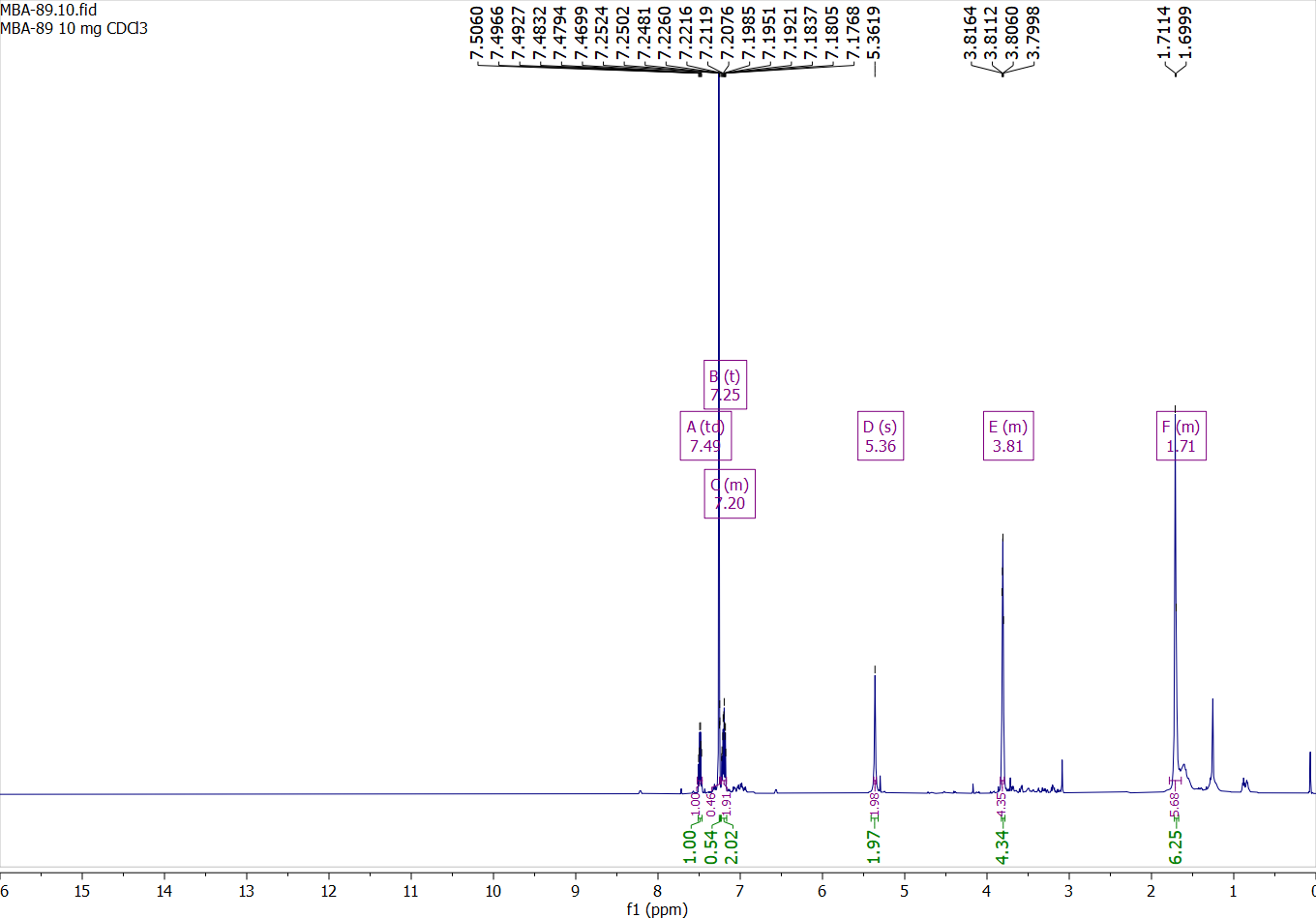

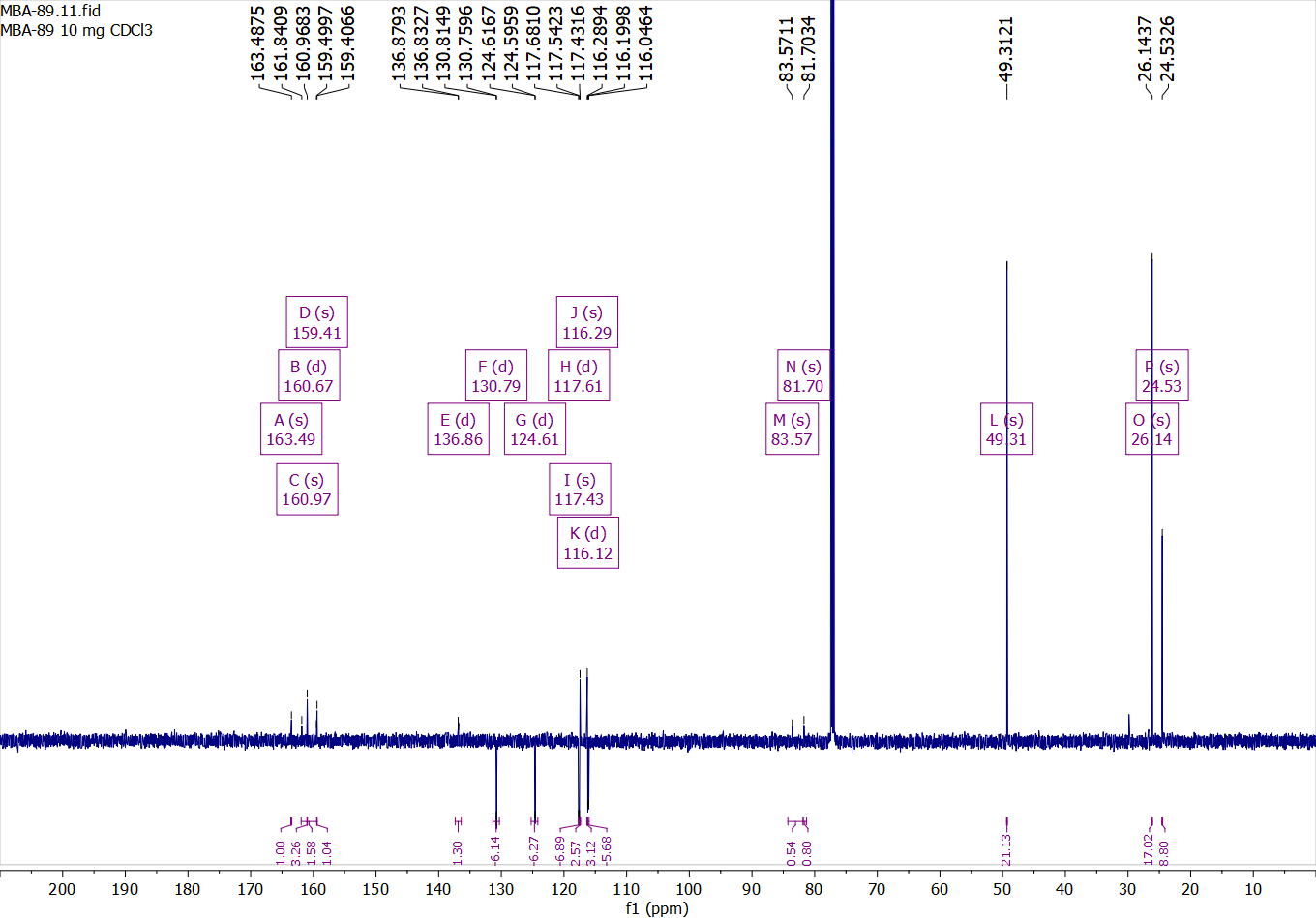

Compound 27:

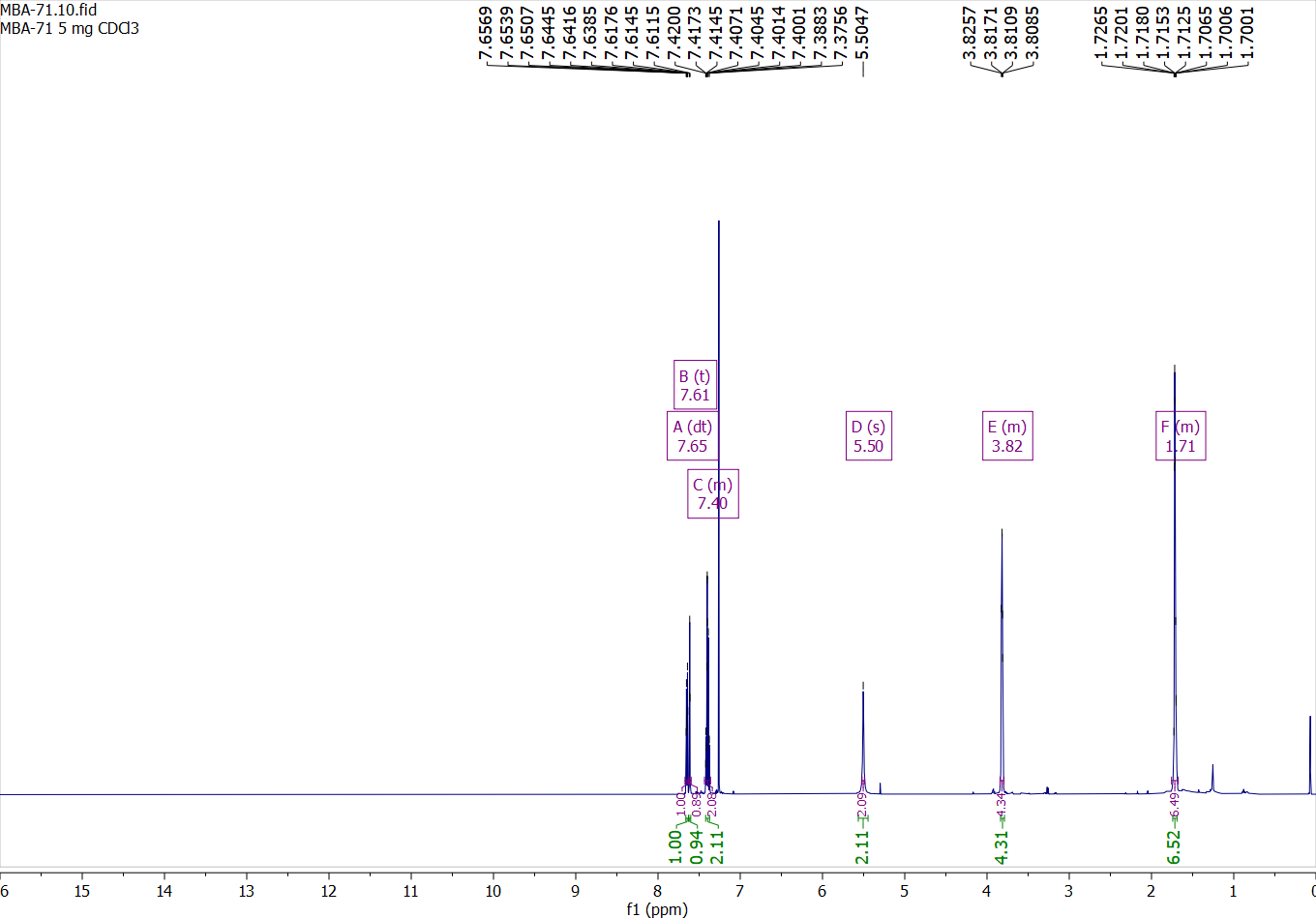

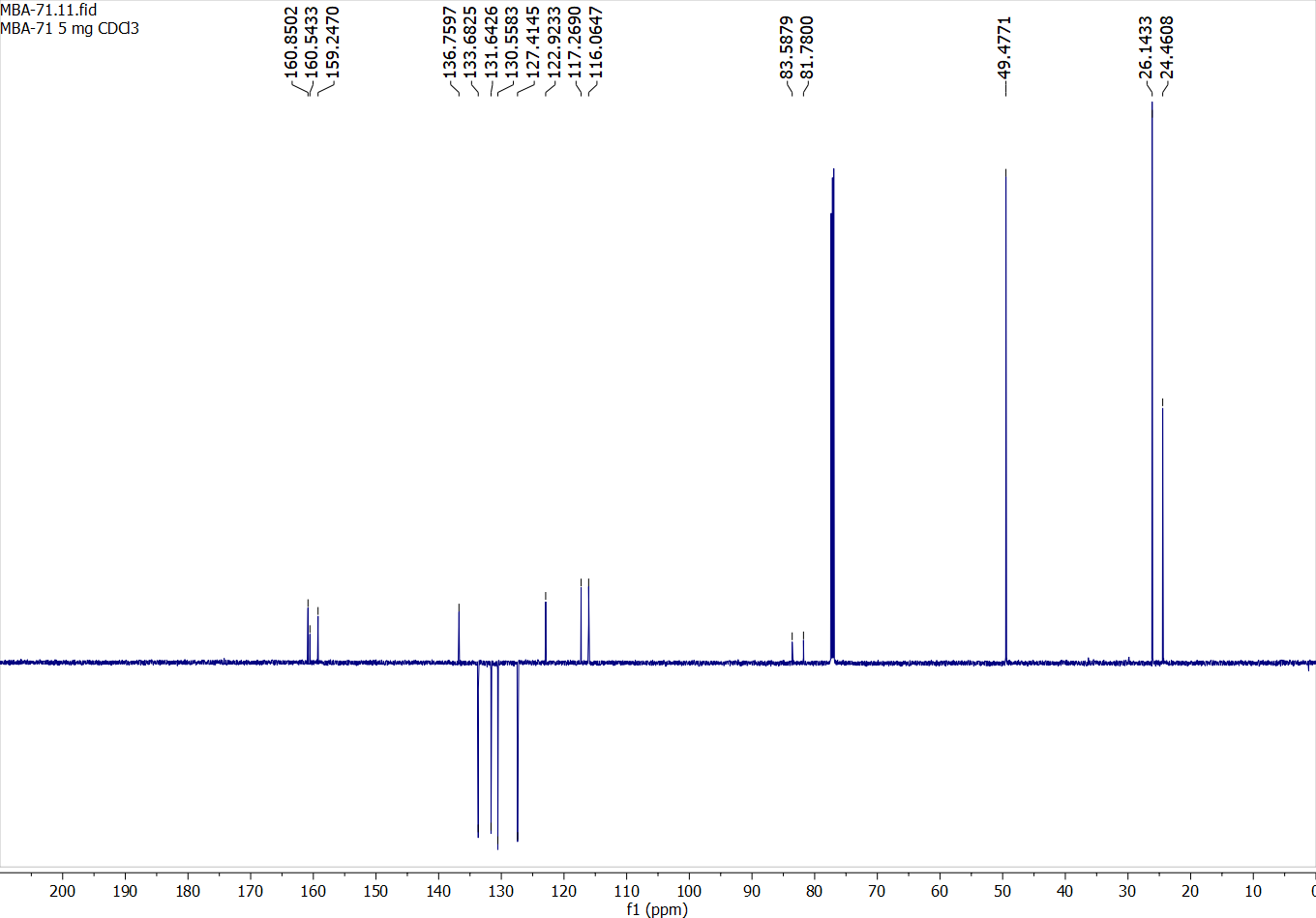

Compound 28:

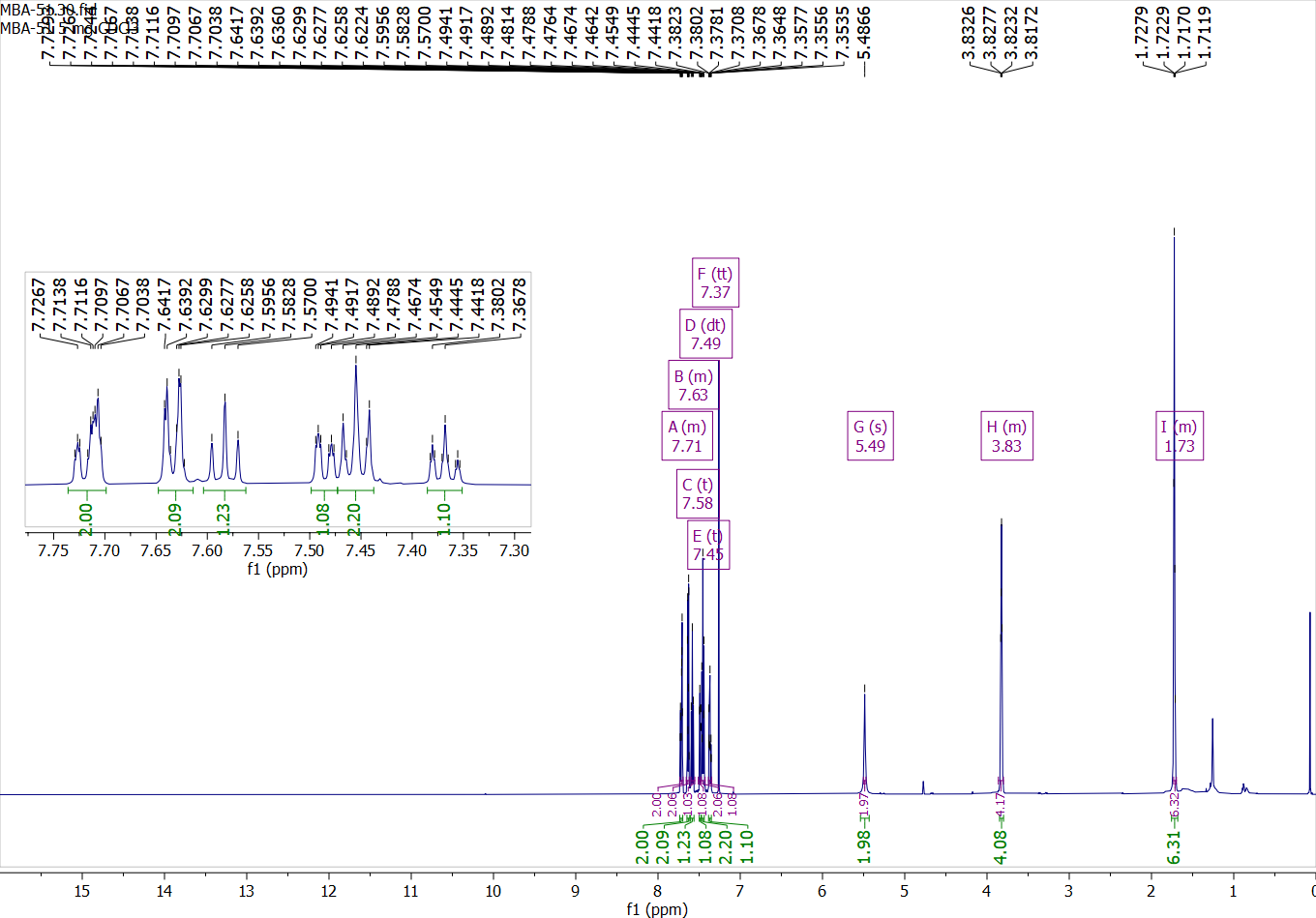

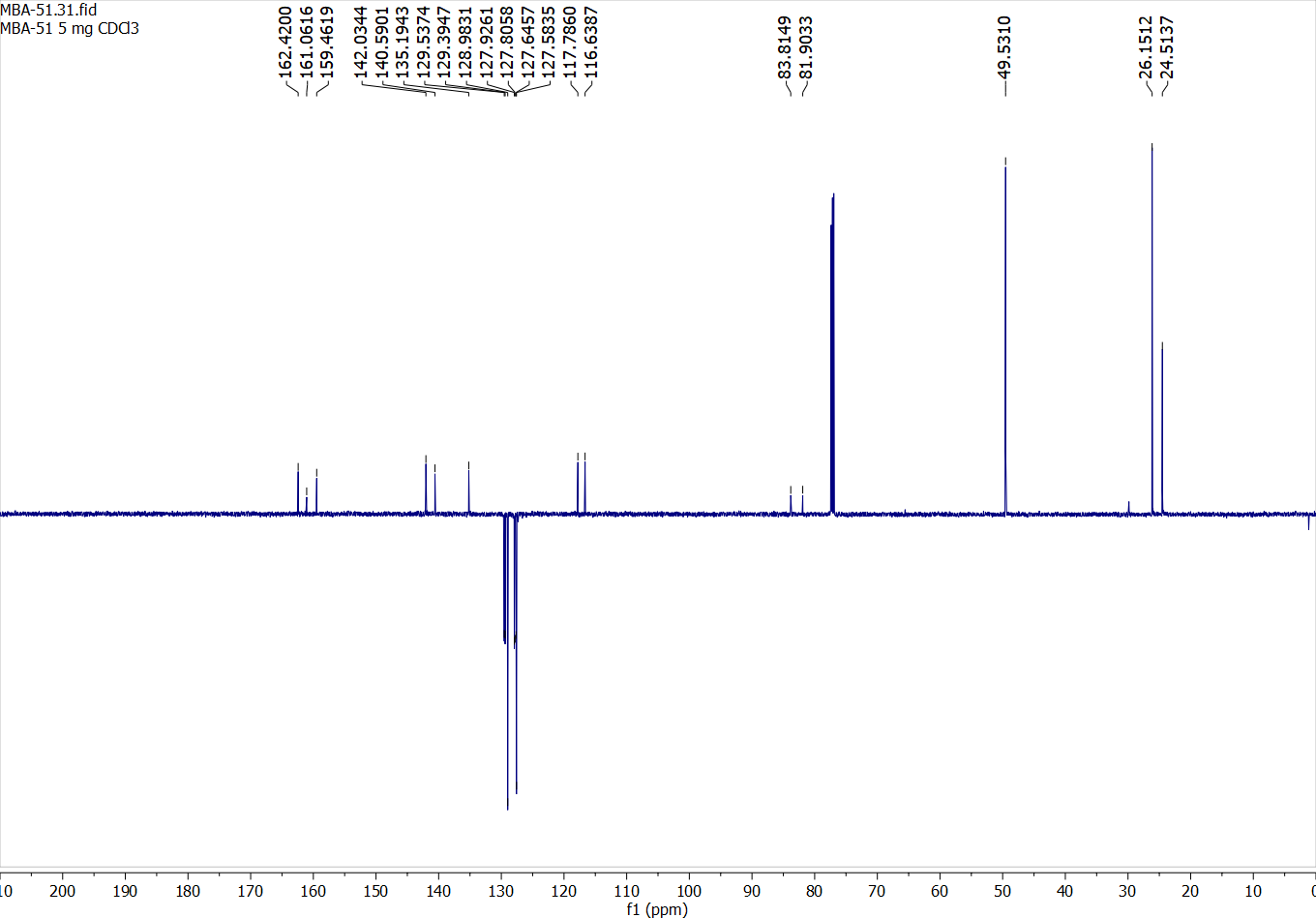

Compound 29:

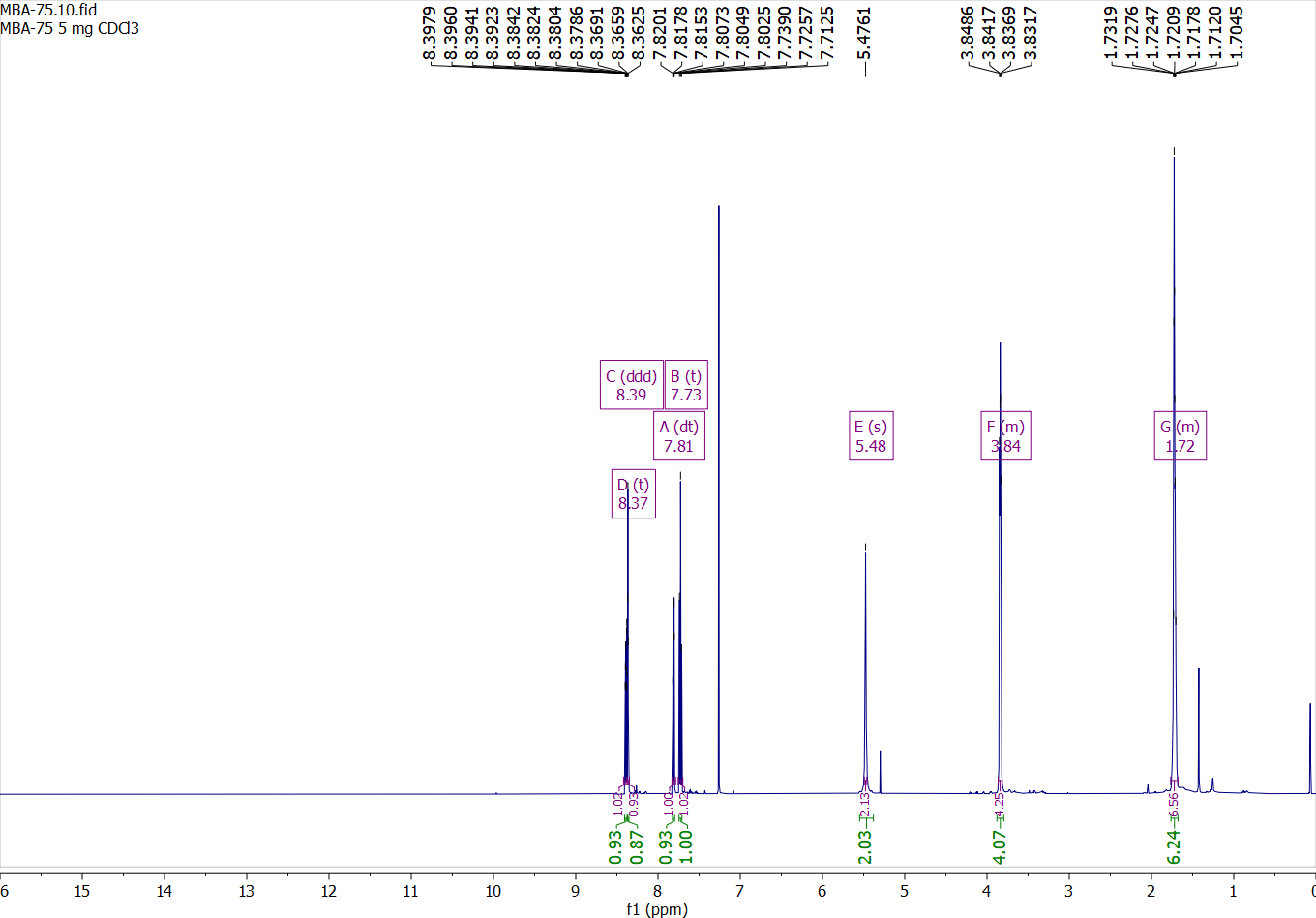

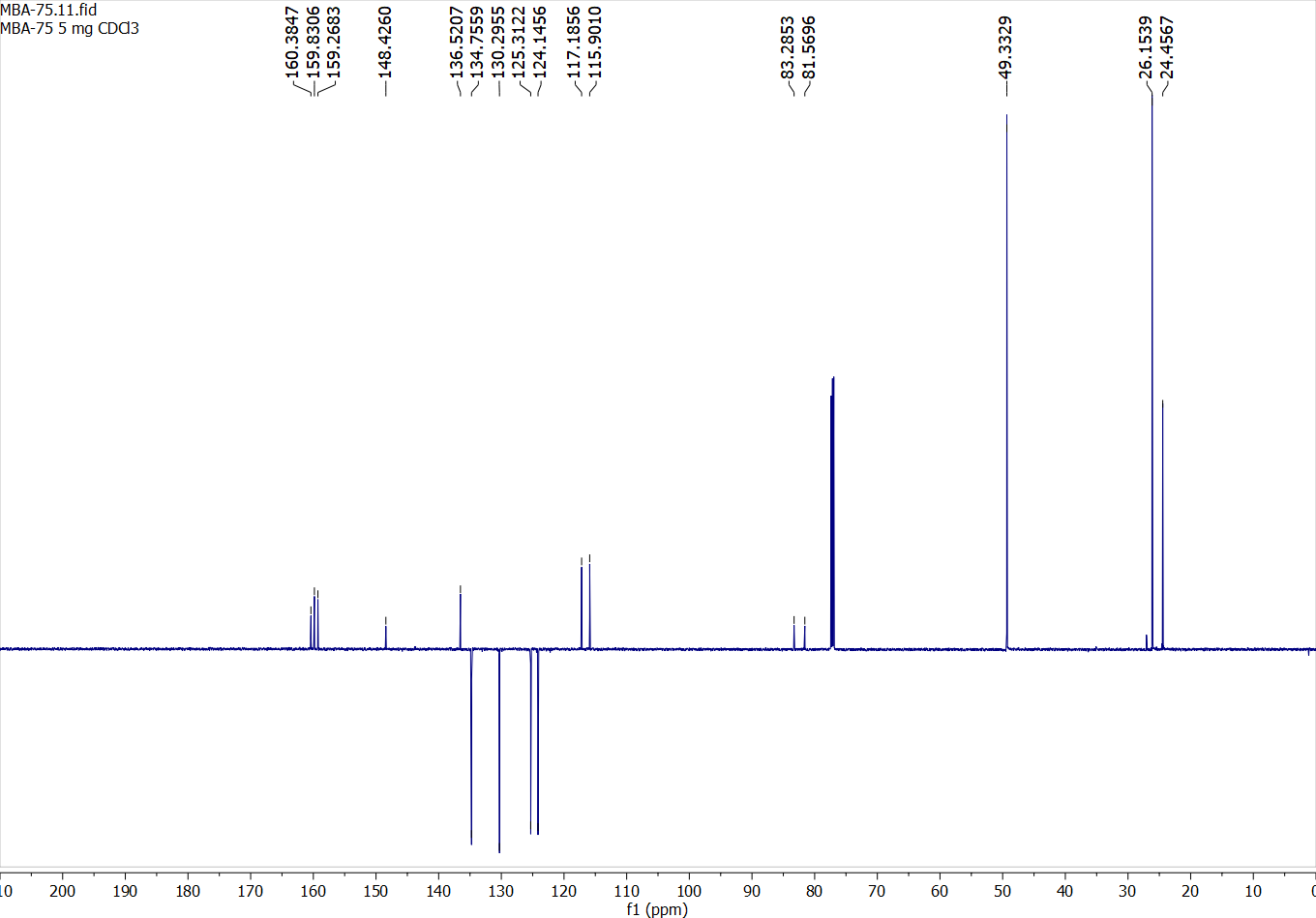

Compound 30:

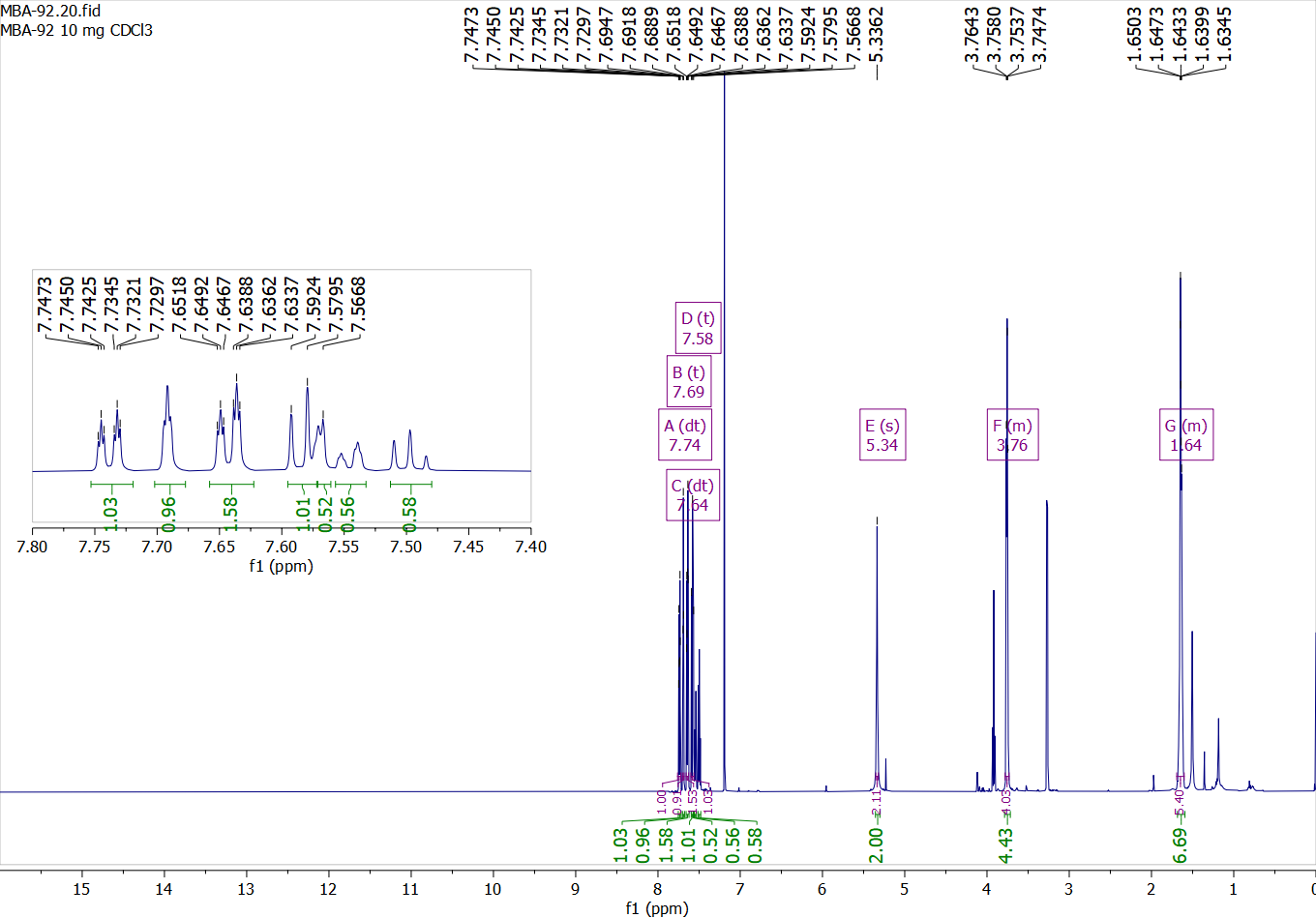

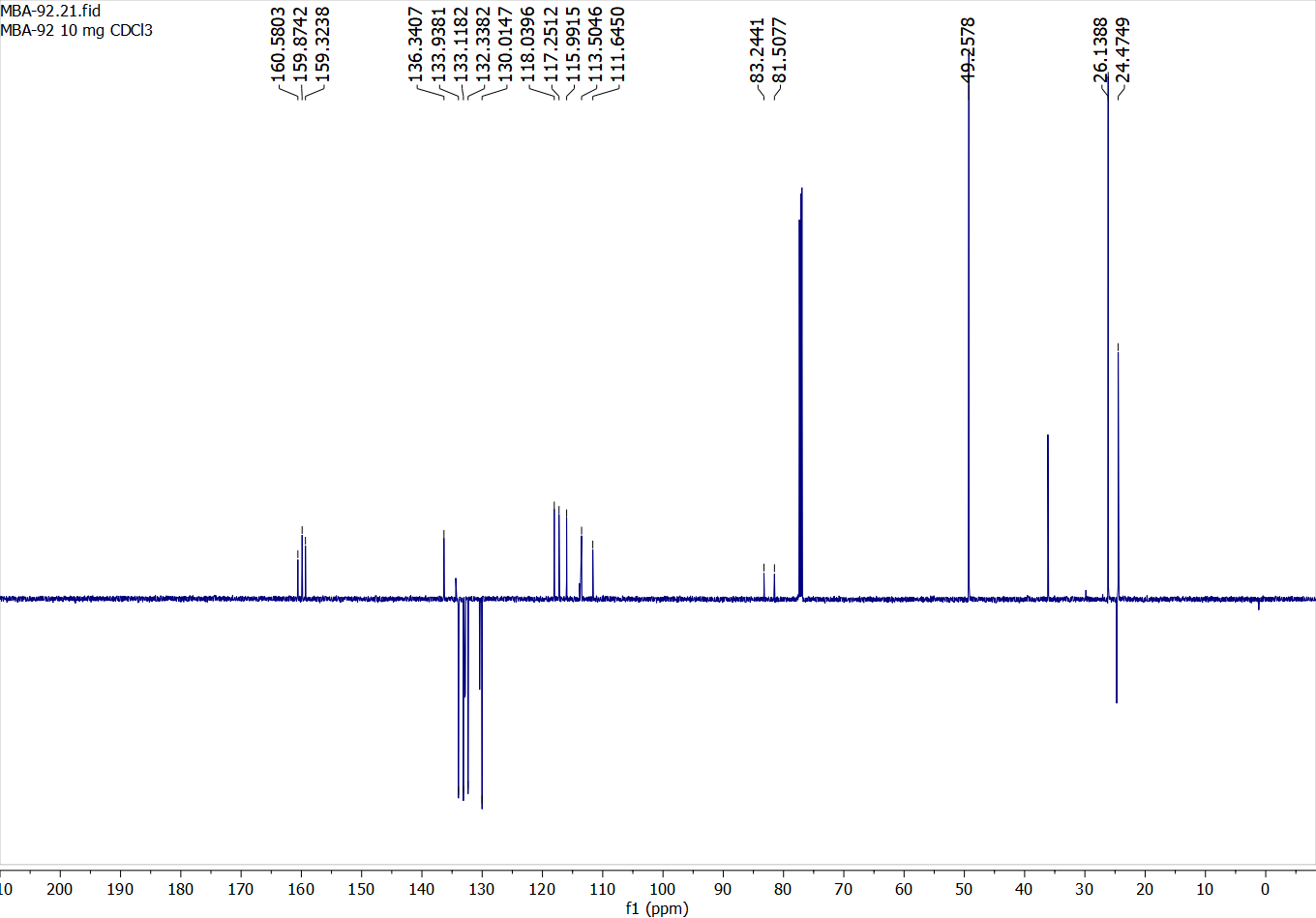

Compound 31:

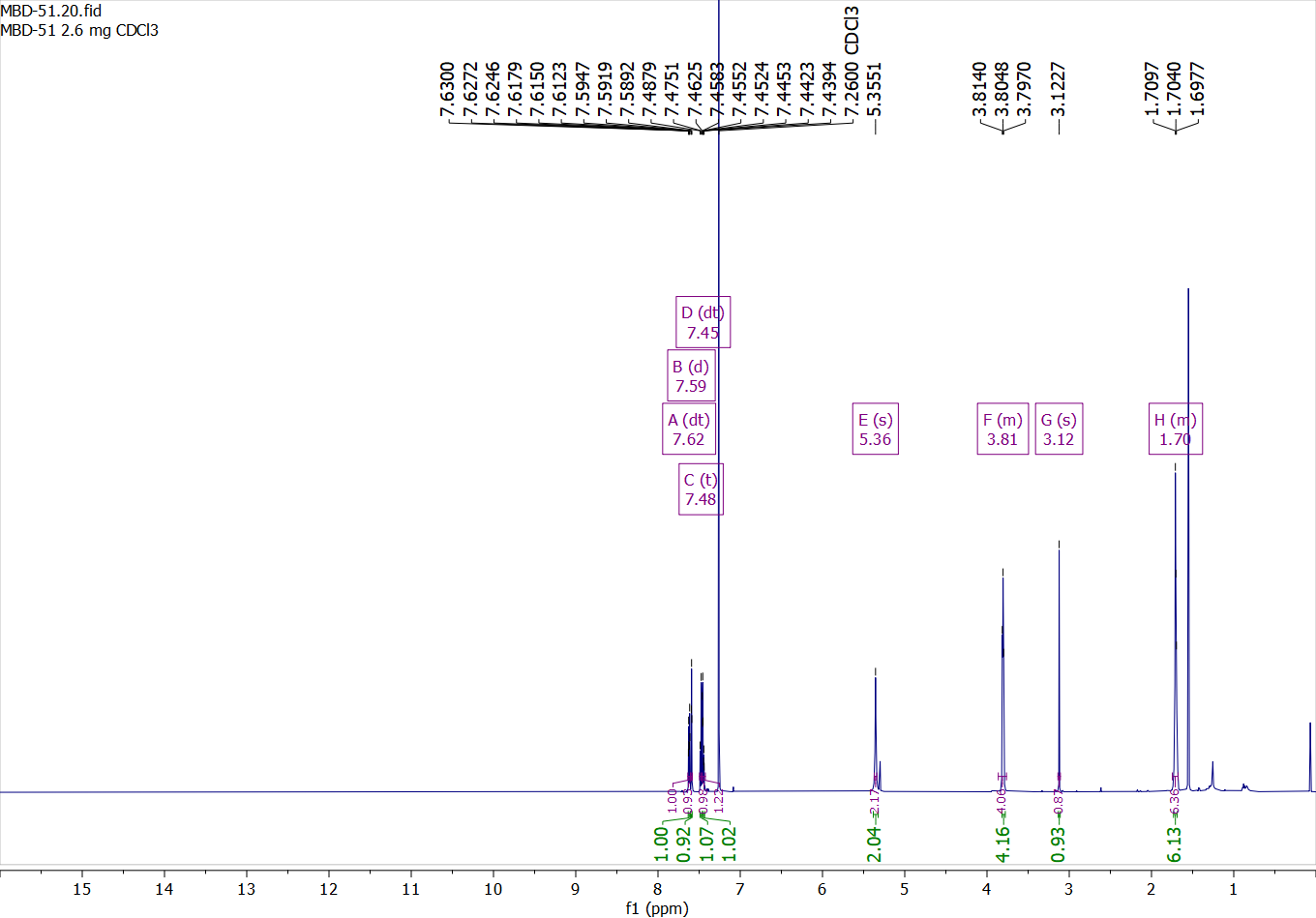

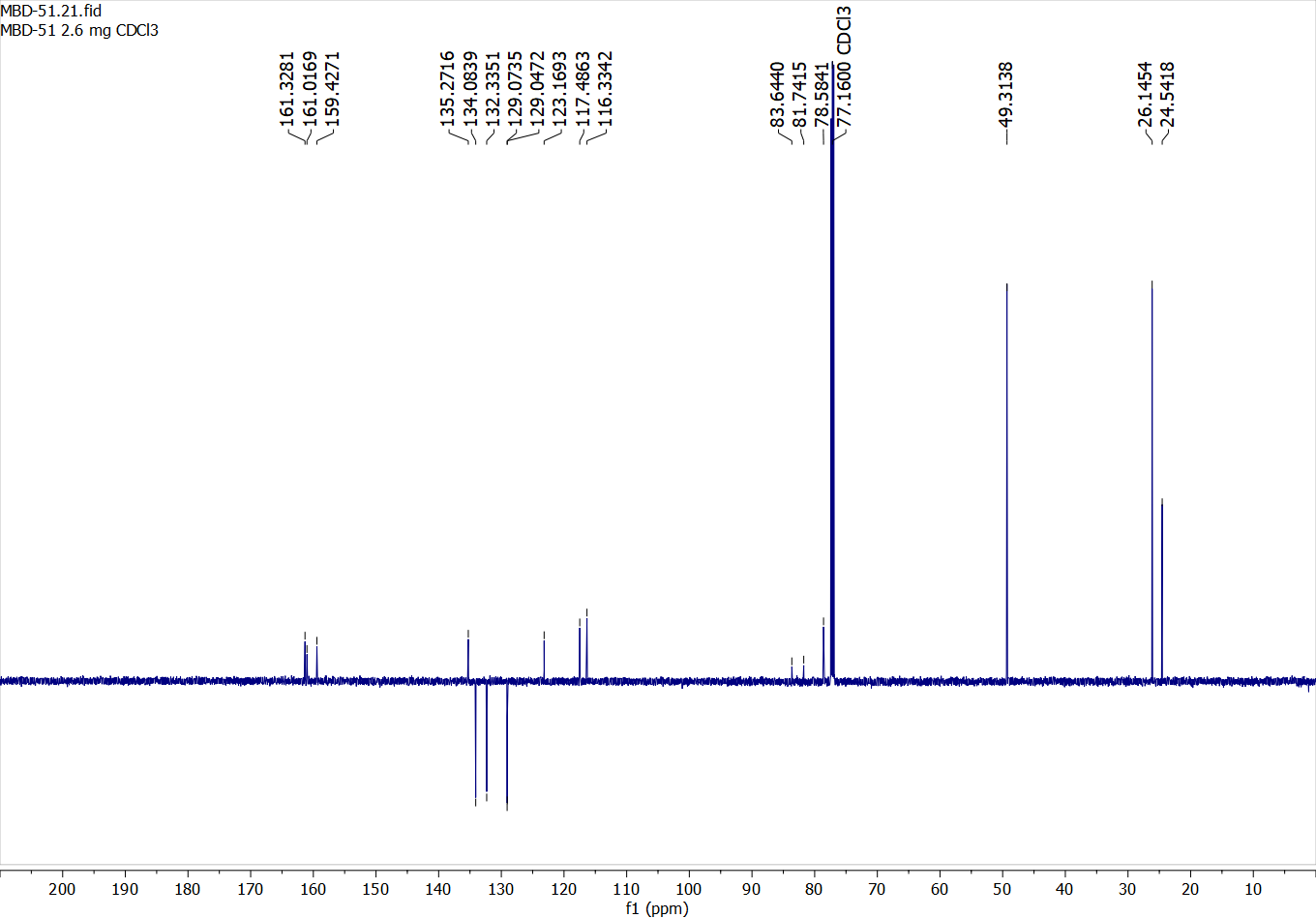

Compound 32:

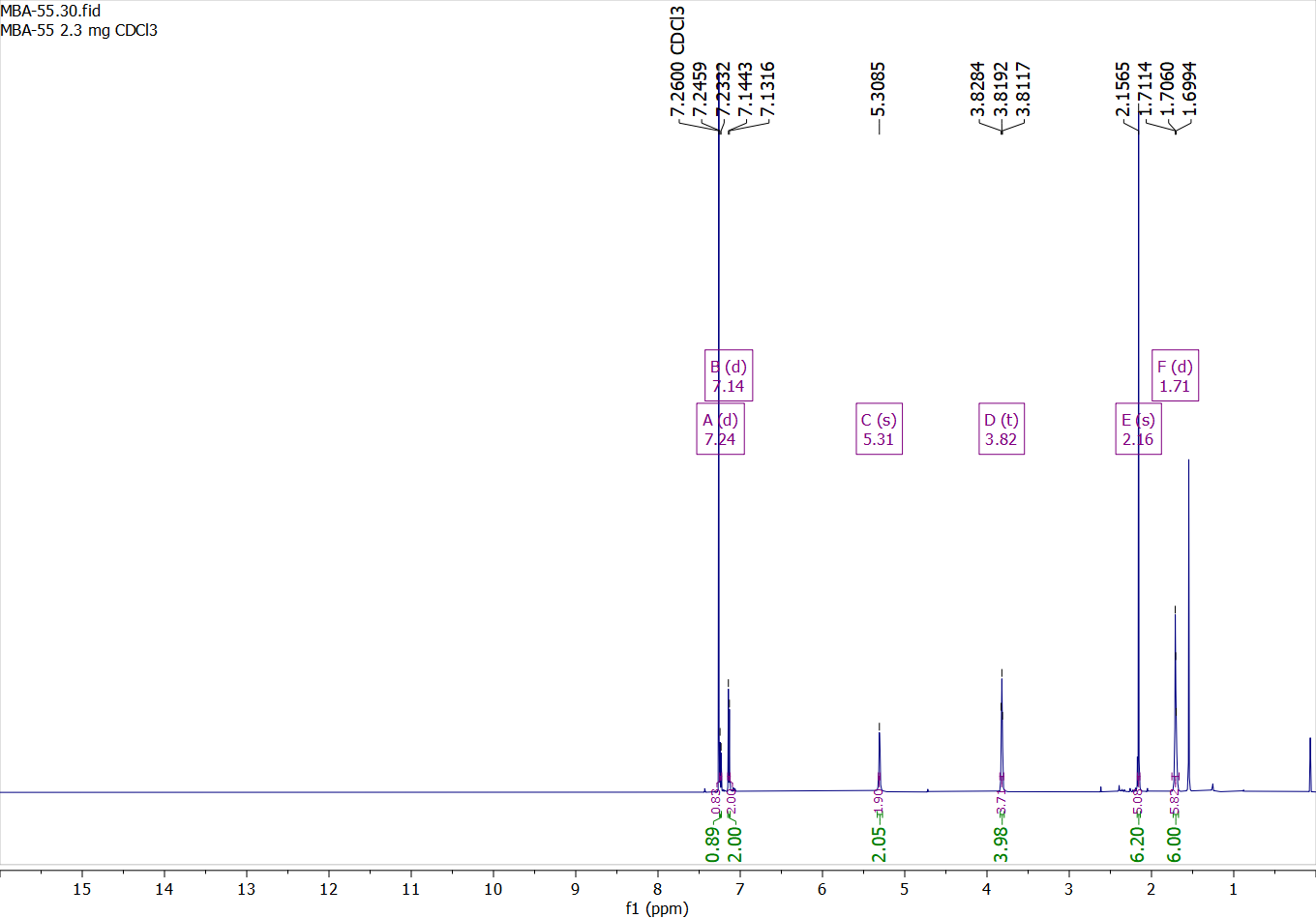

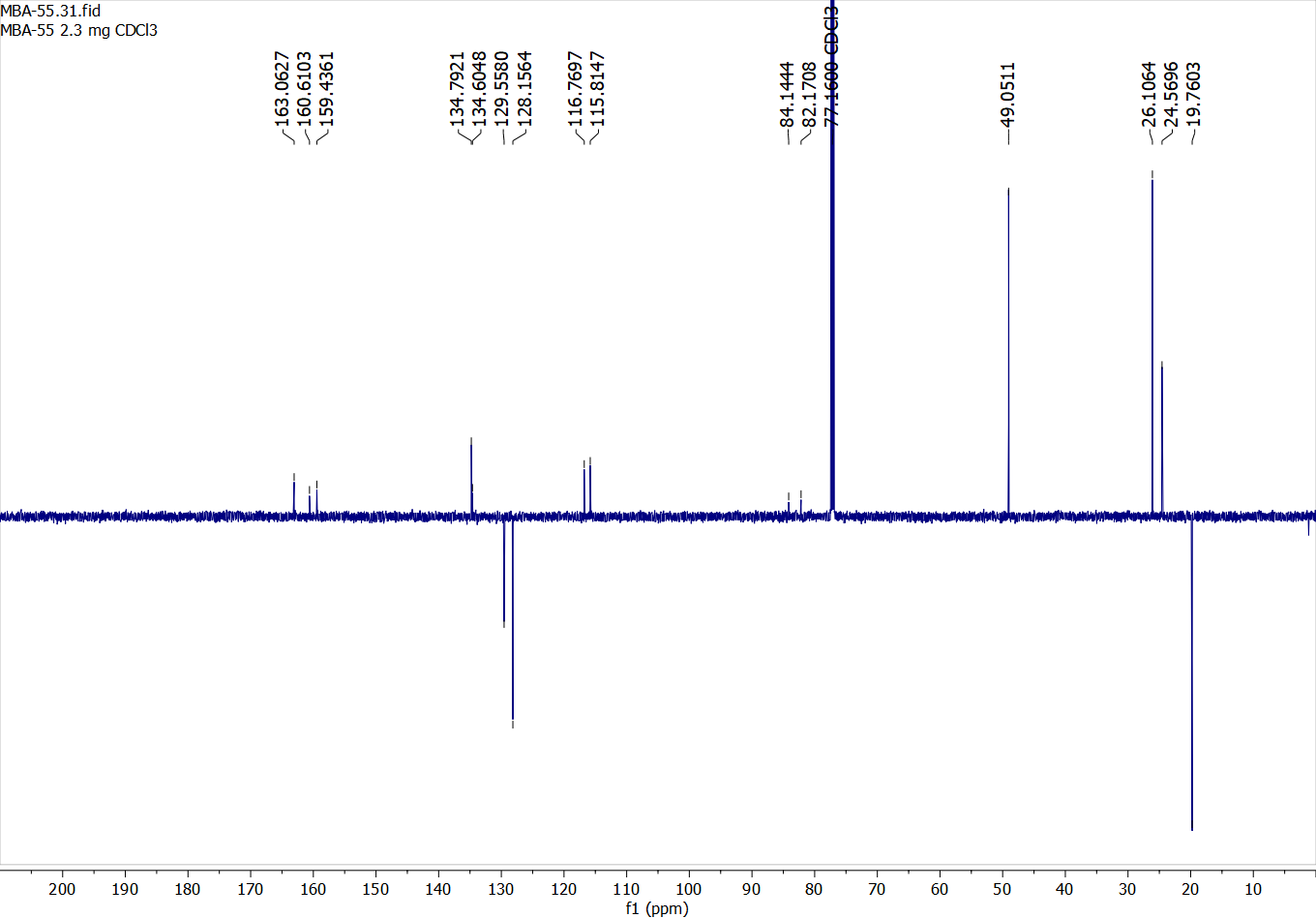

Compound 33:

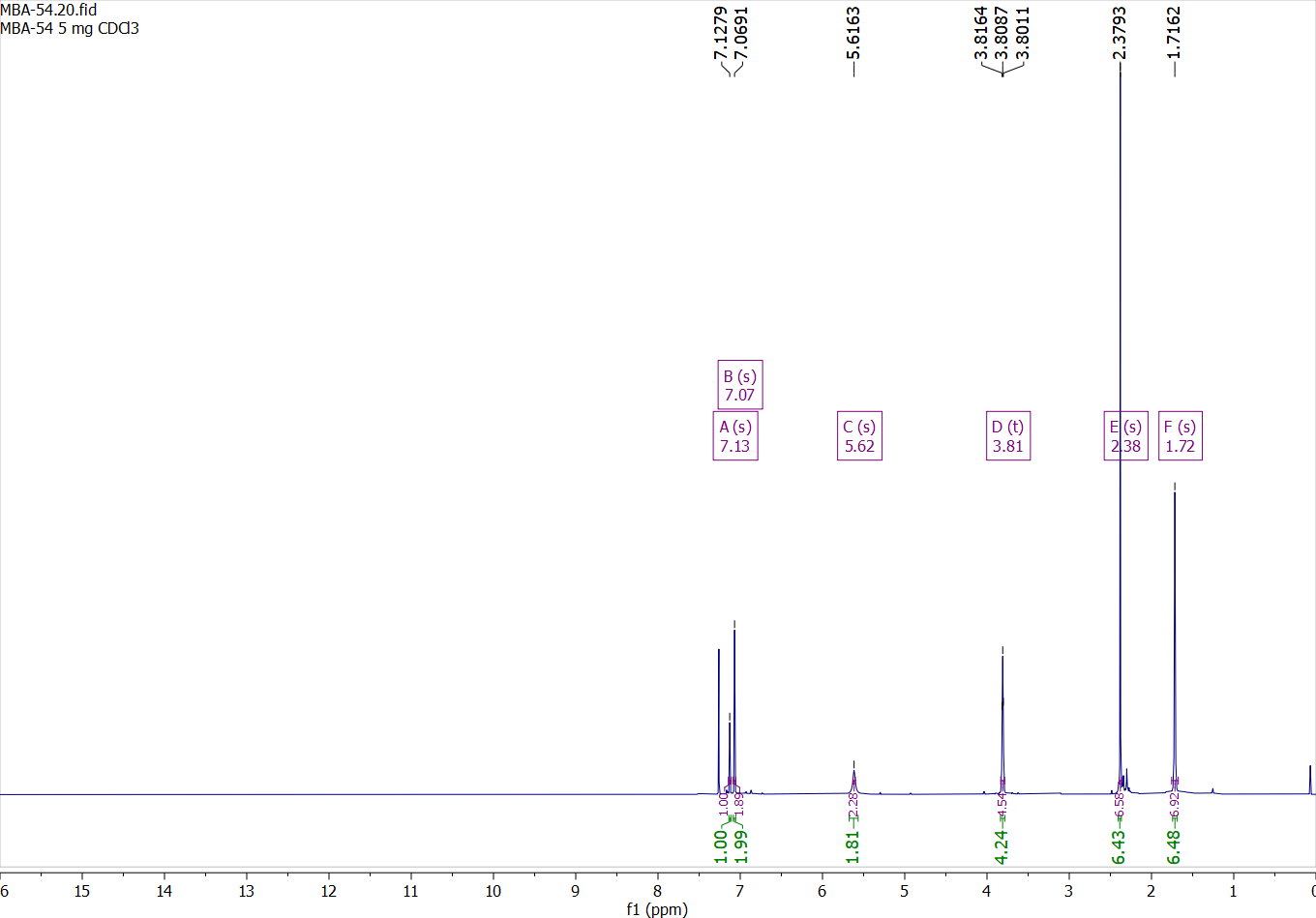

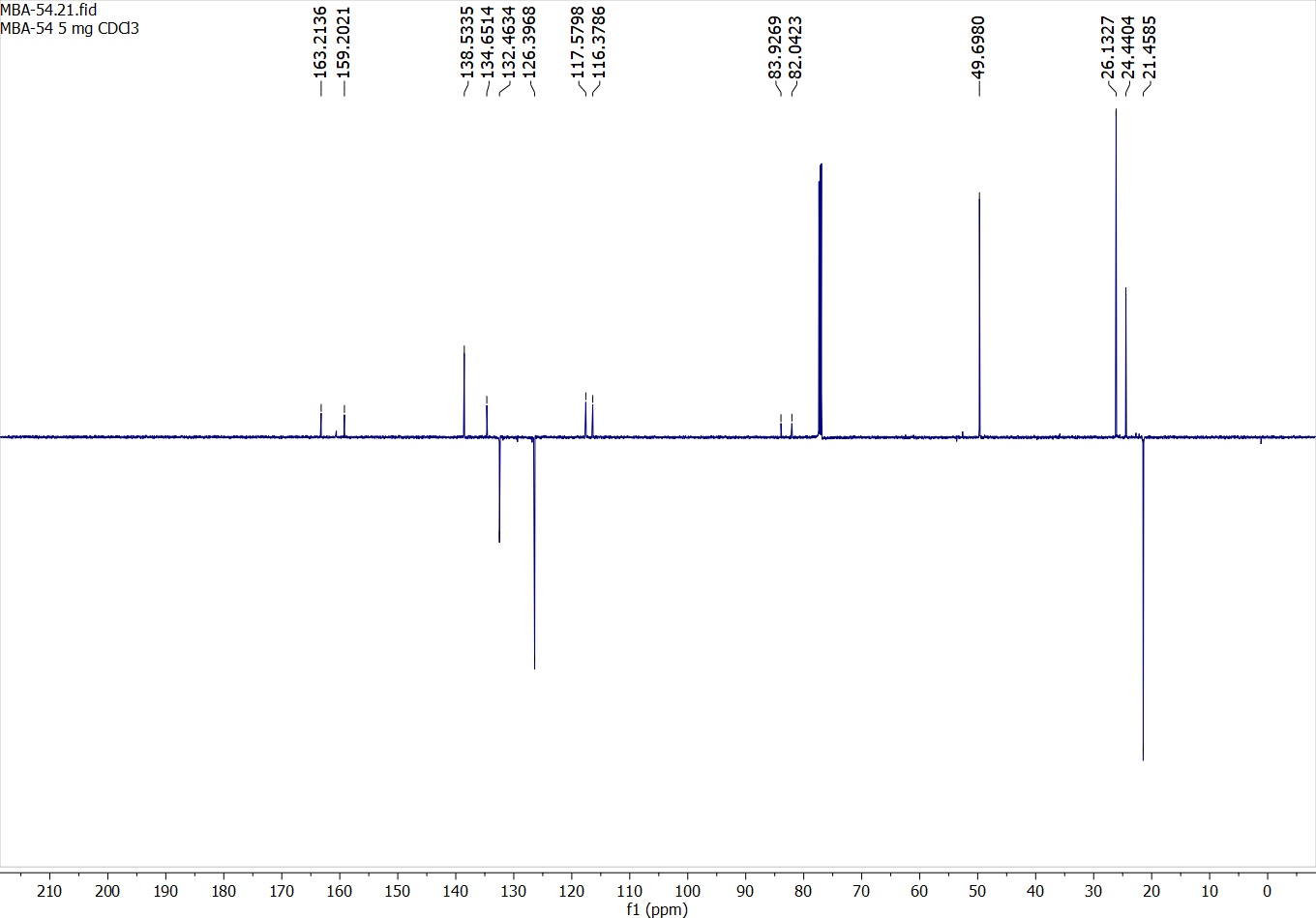

Compound 34:

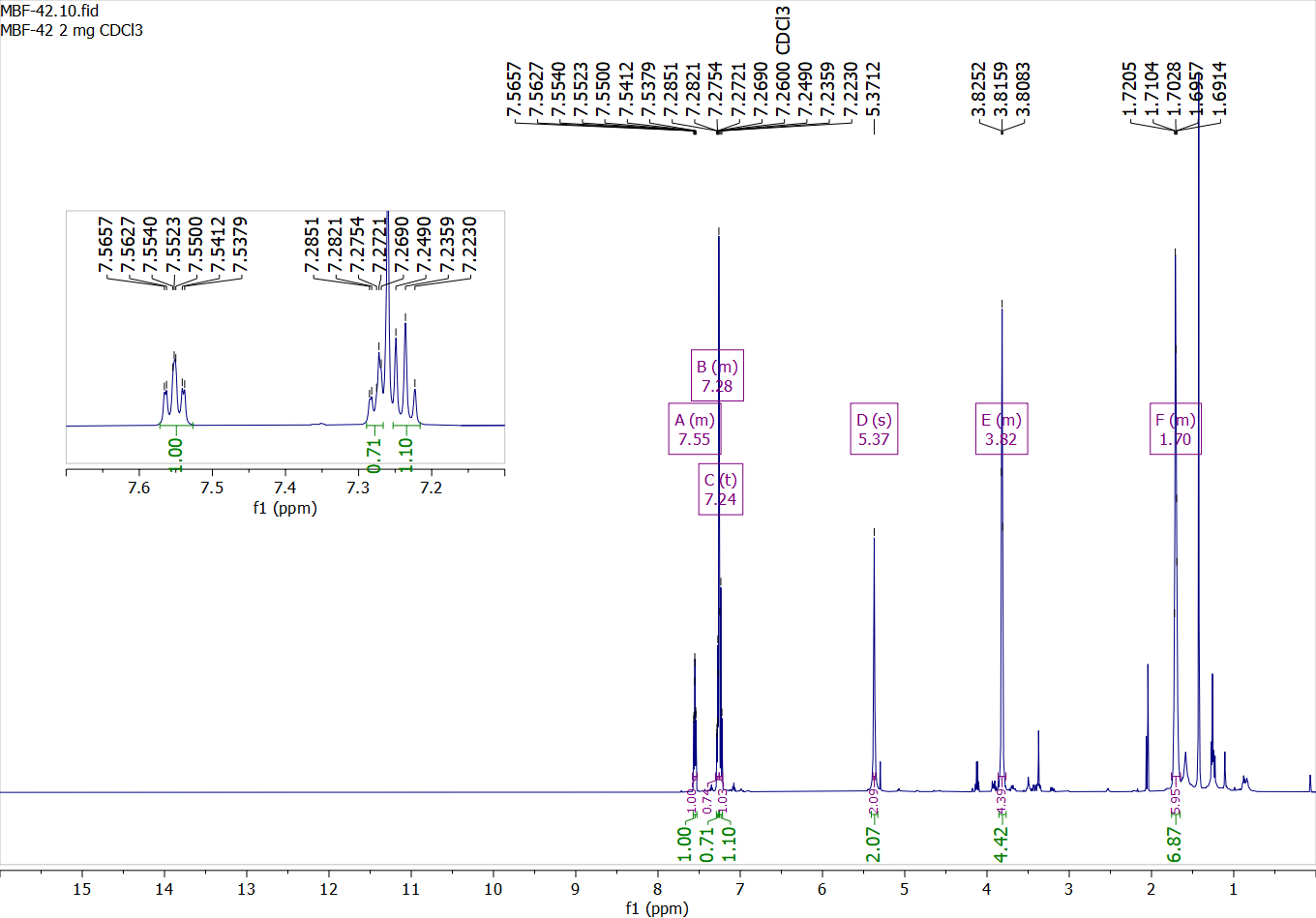

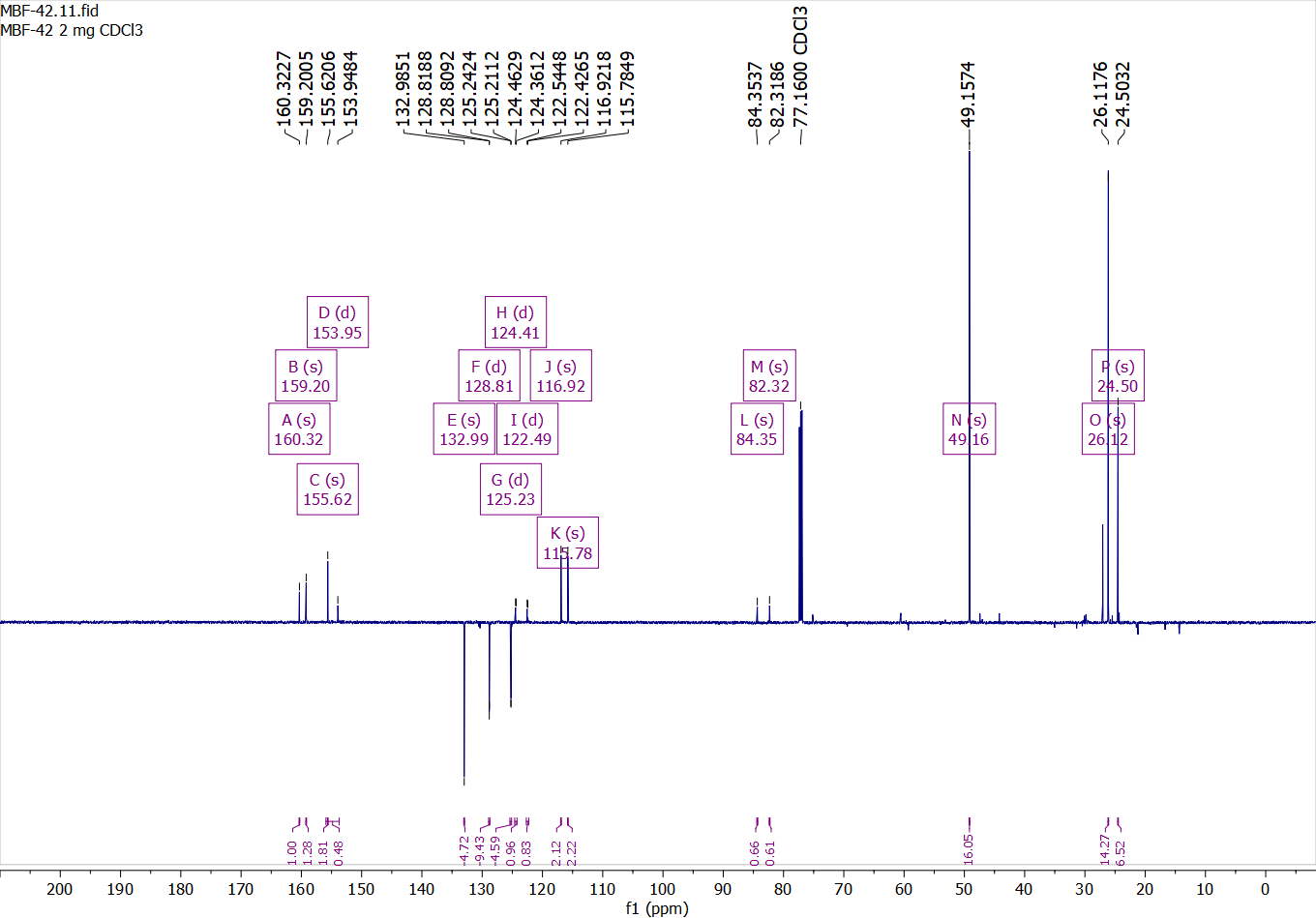

Compound 35:

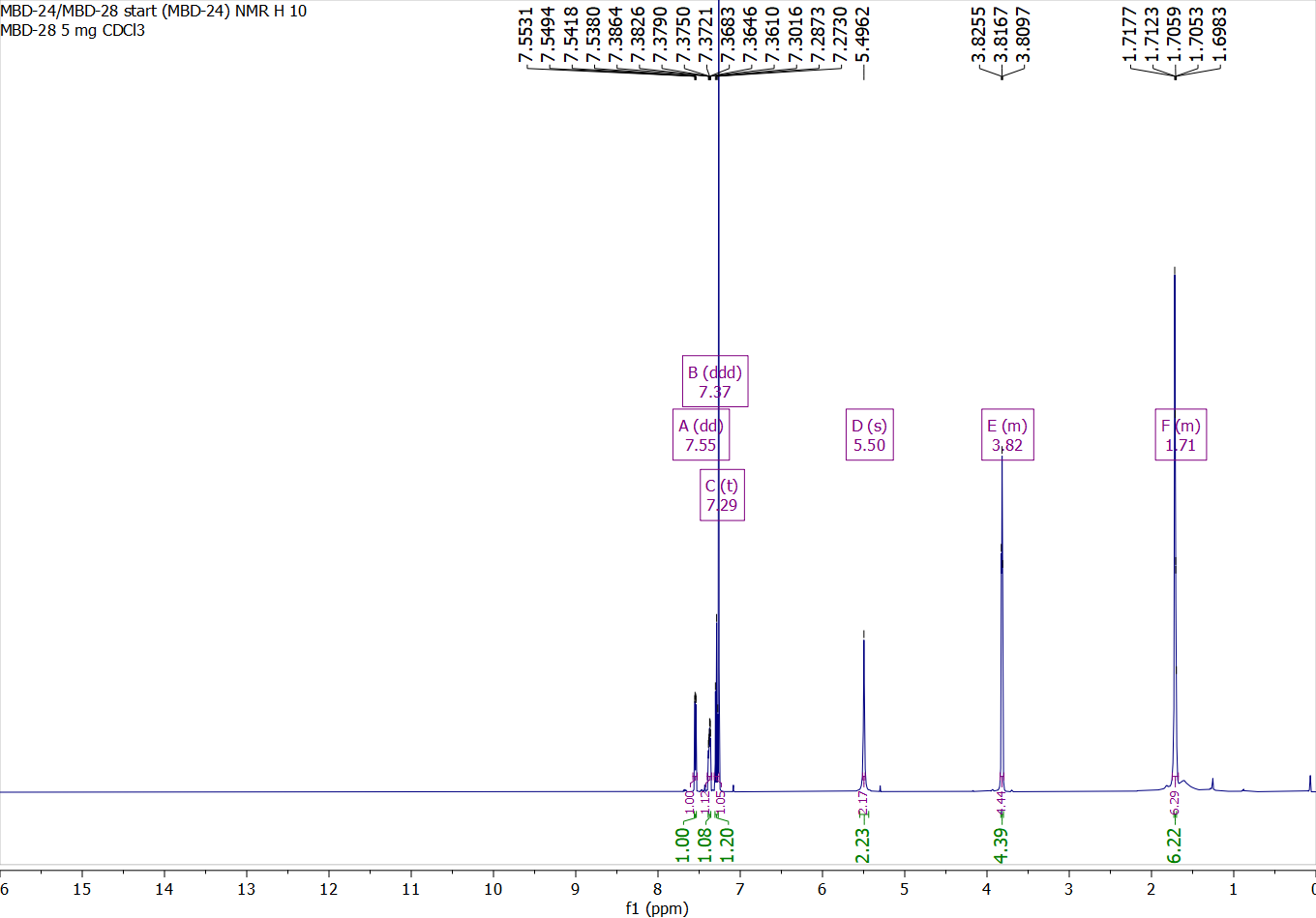

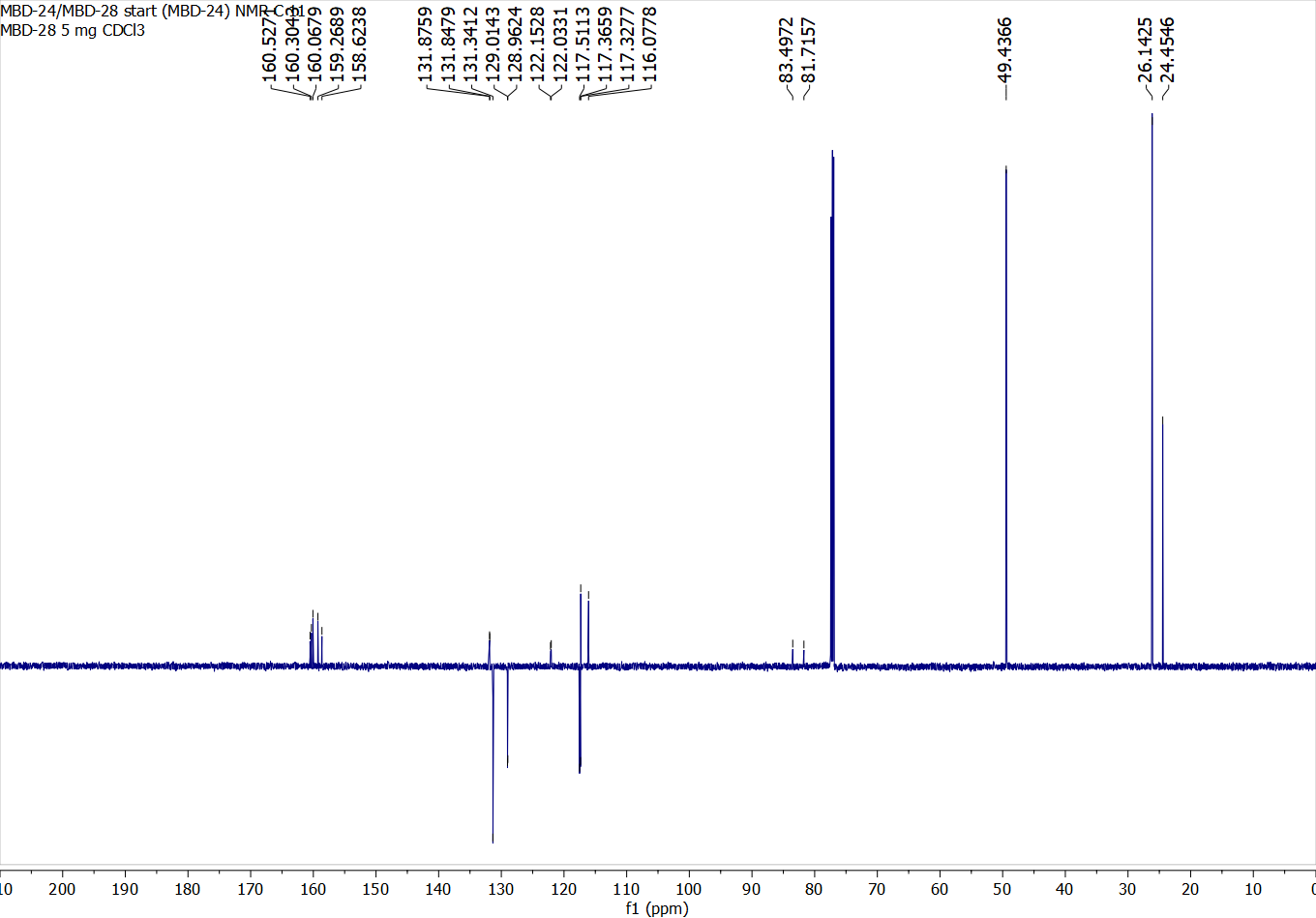

Compound 36:

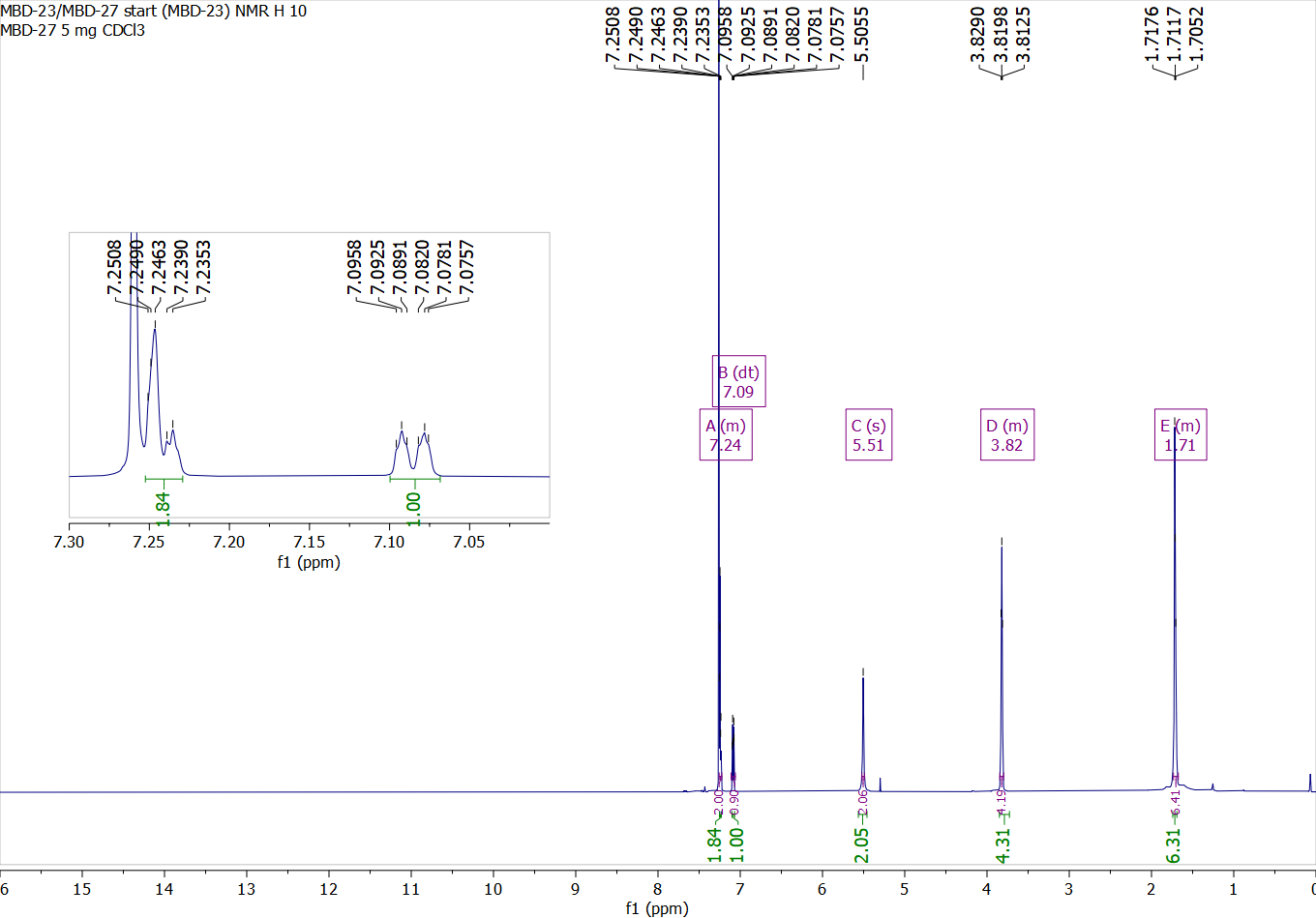

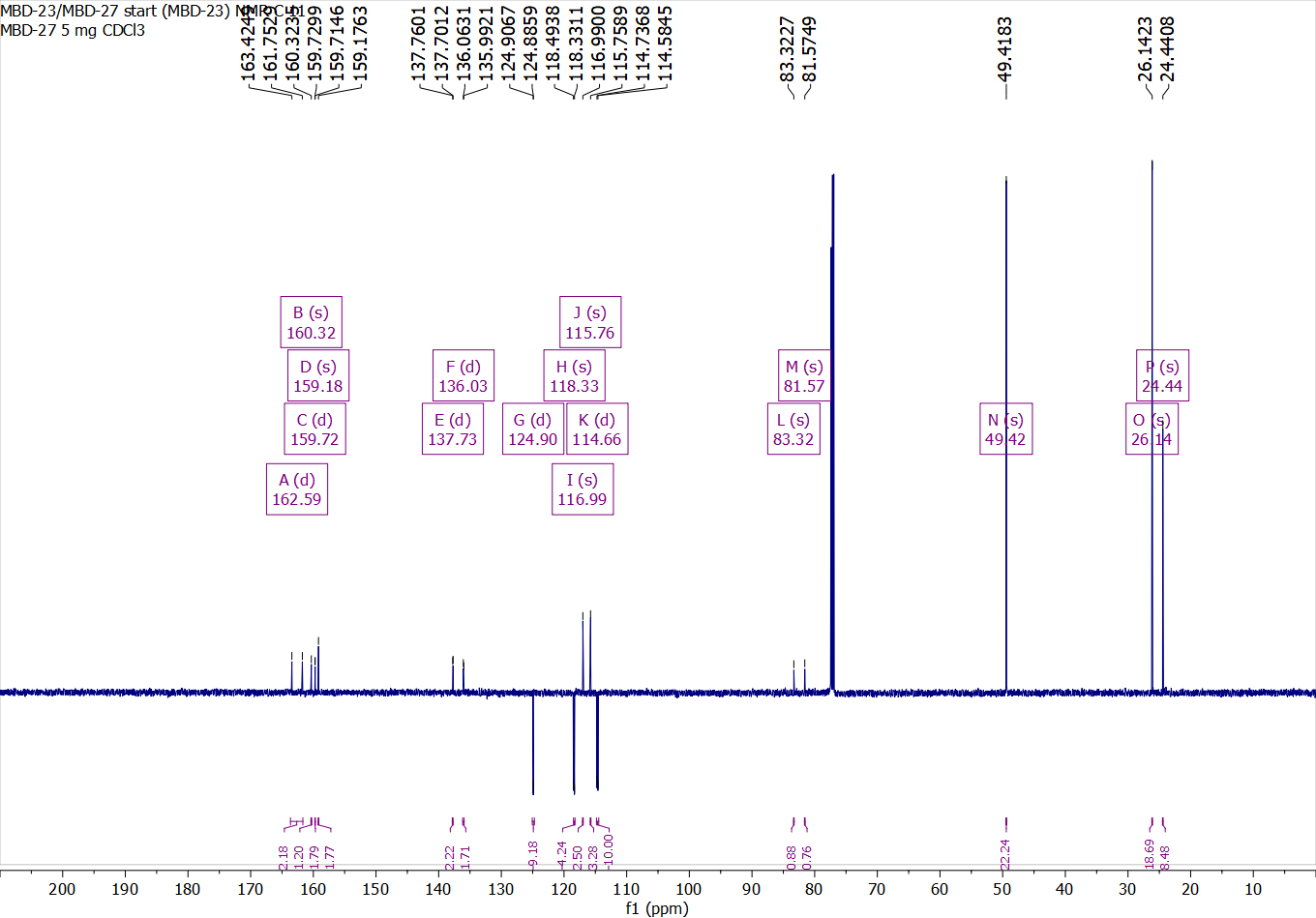

Compound 37:

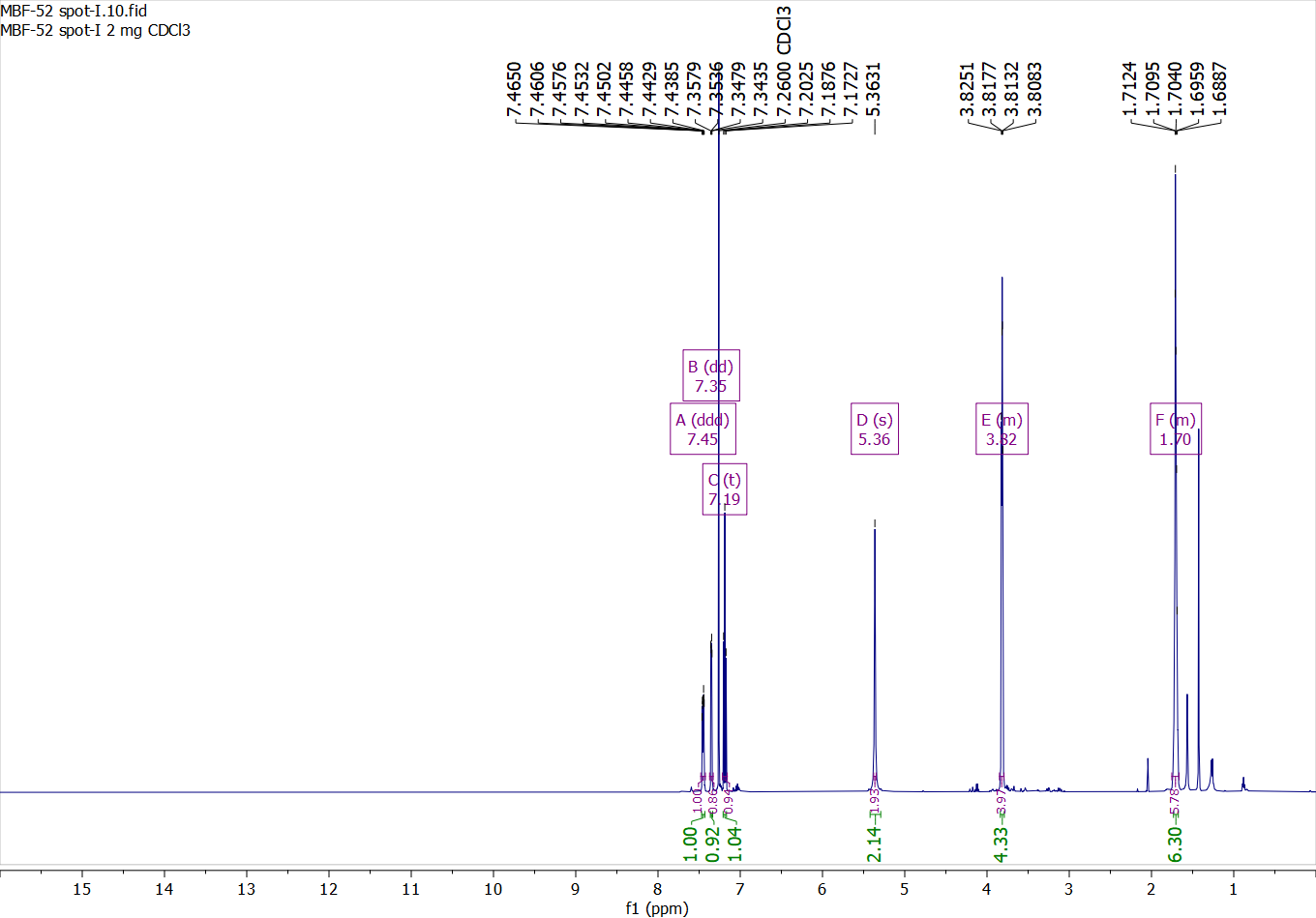

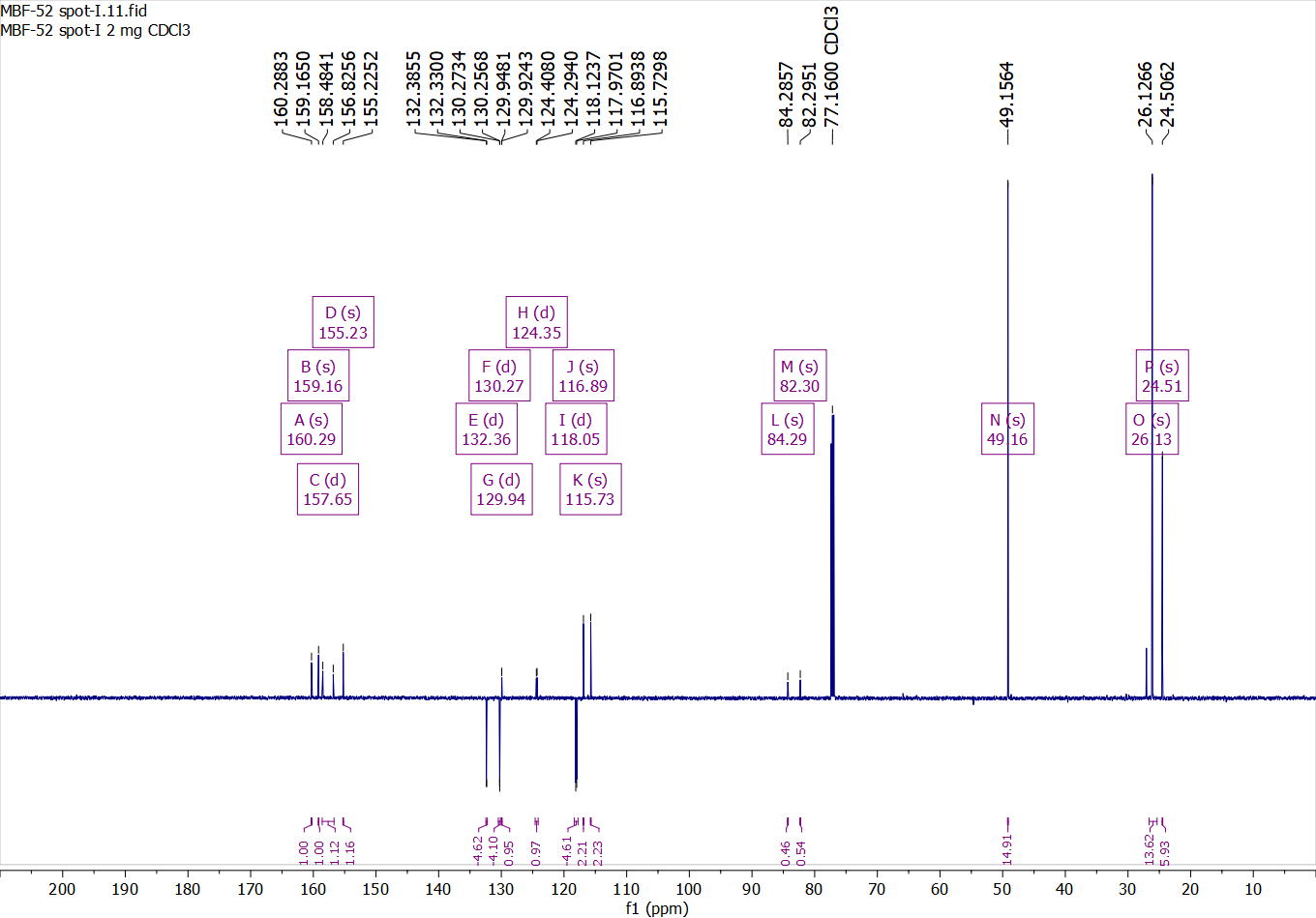

Compound 38:

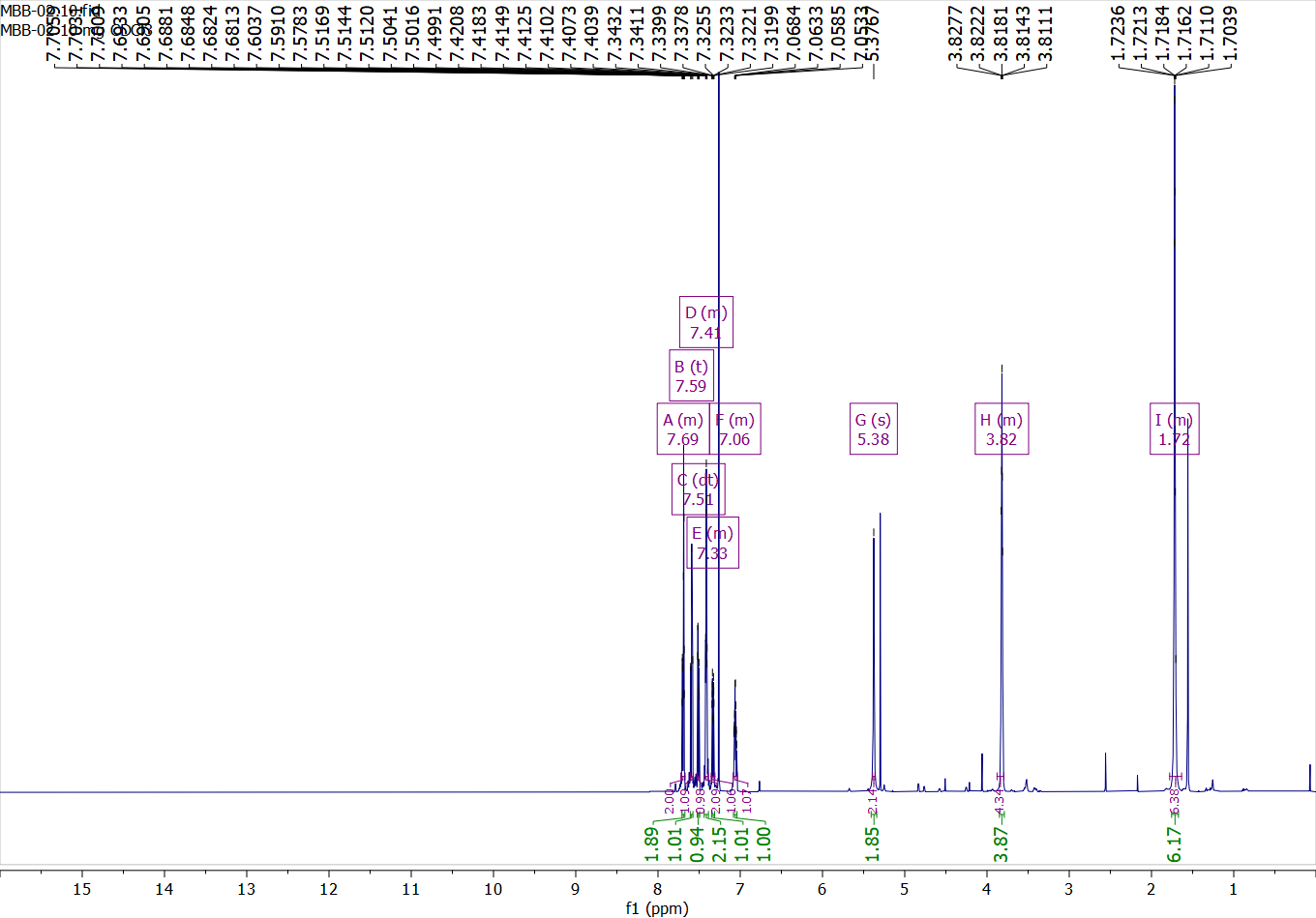

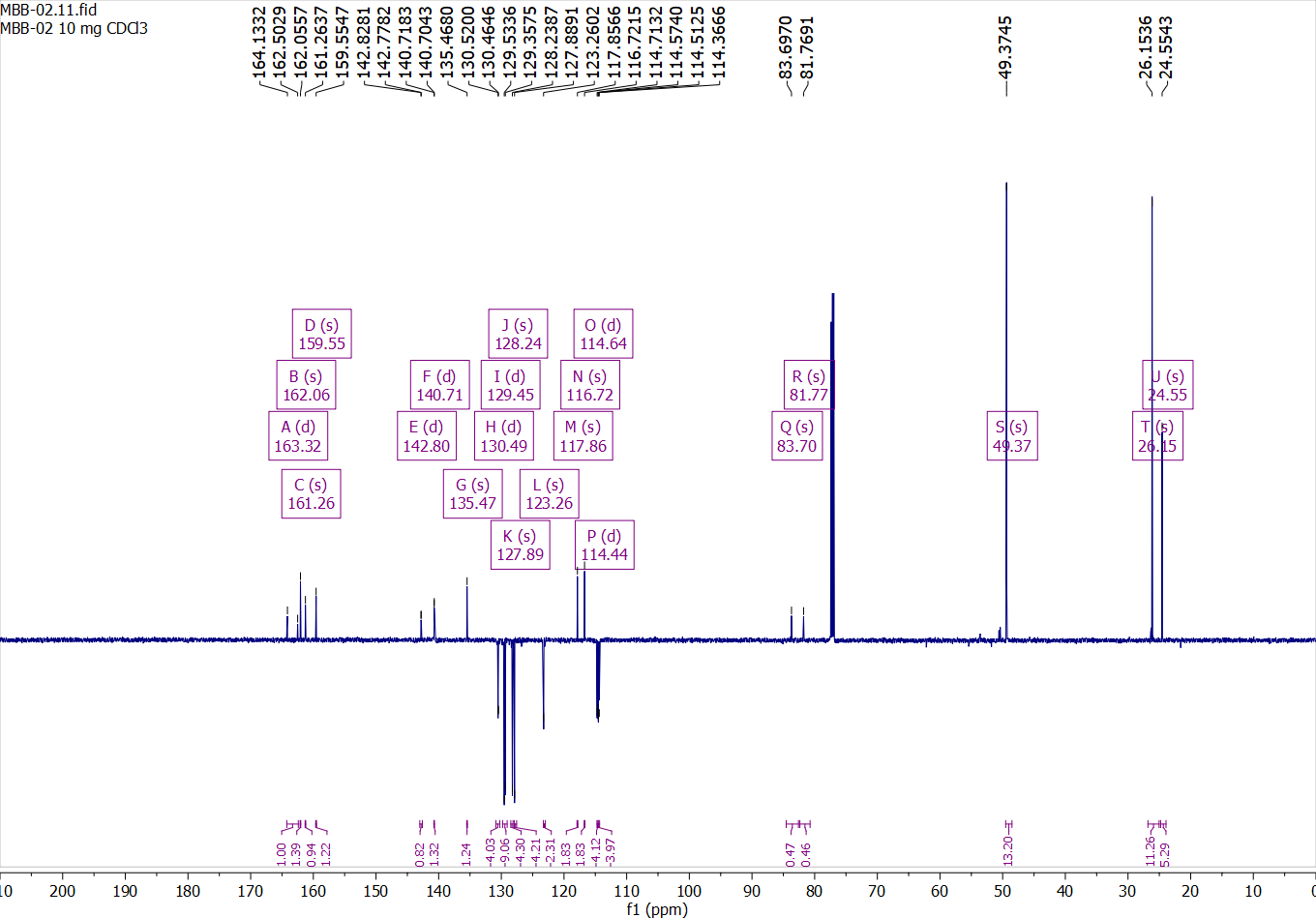

Compound 39:

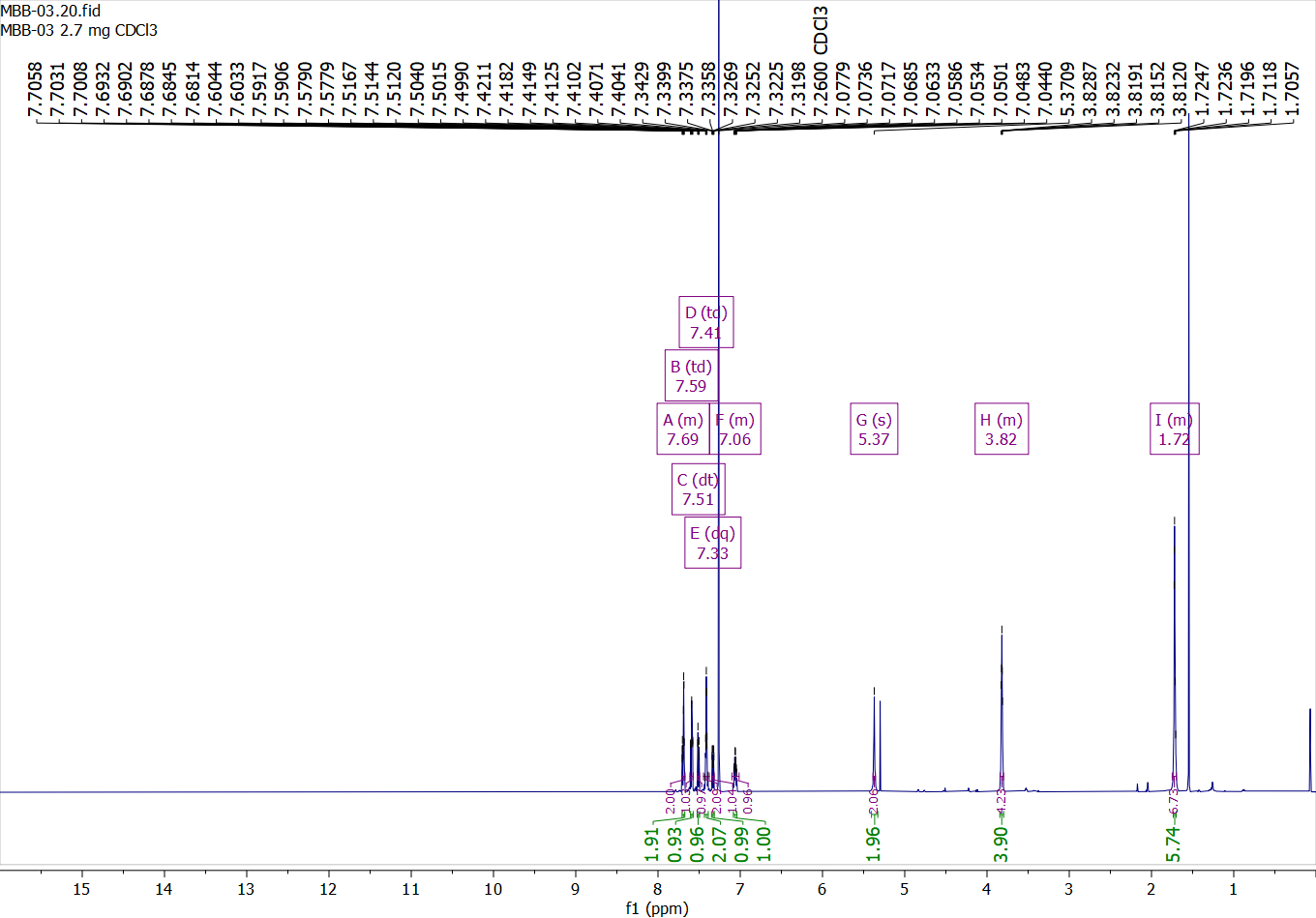

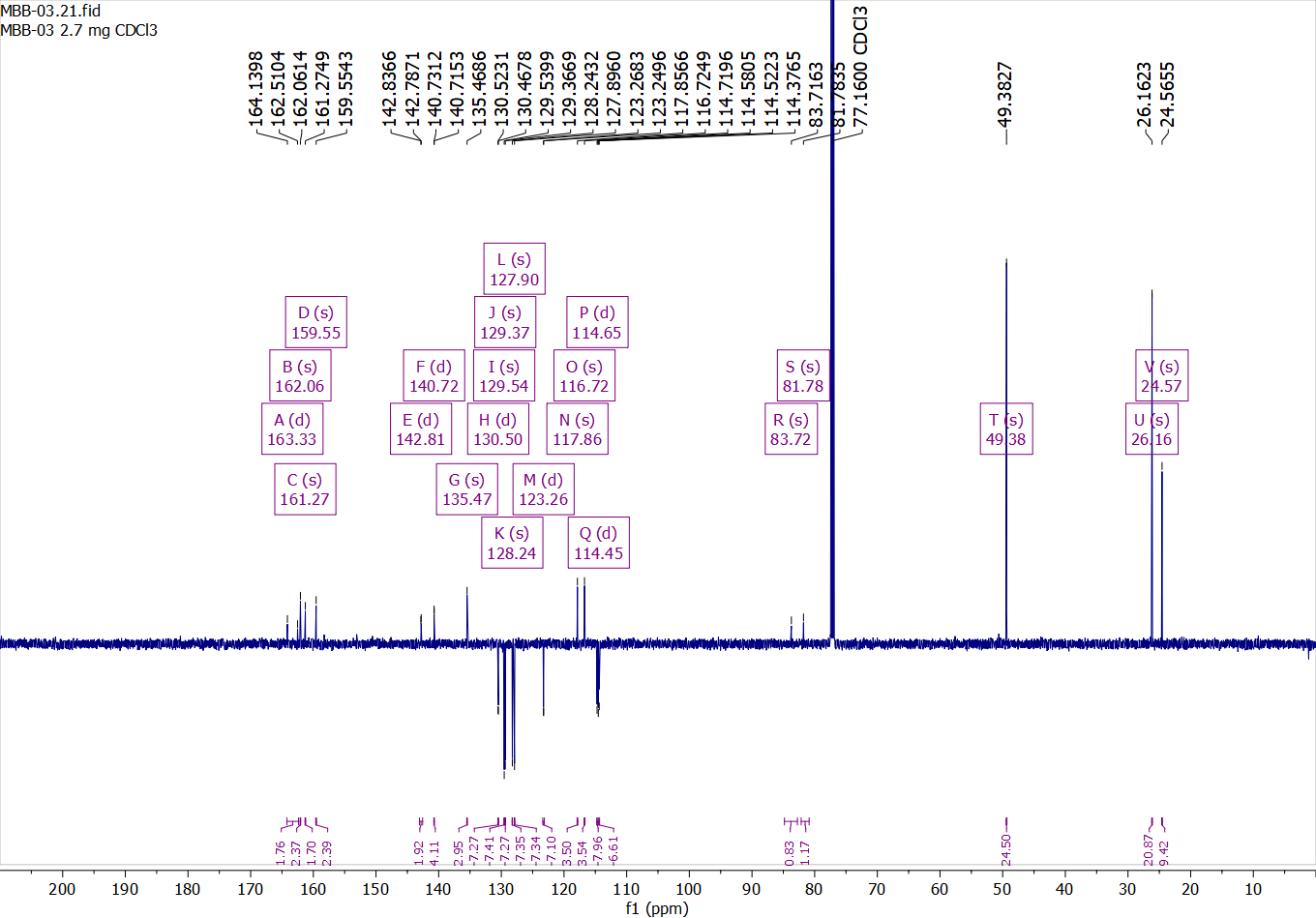

Compound 40:

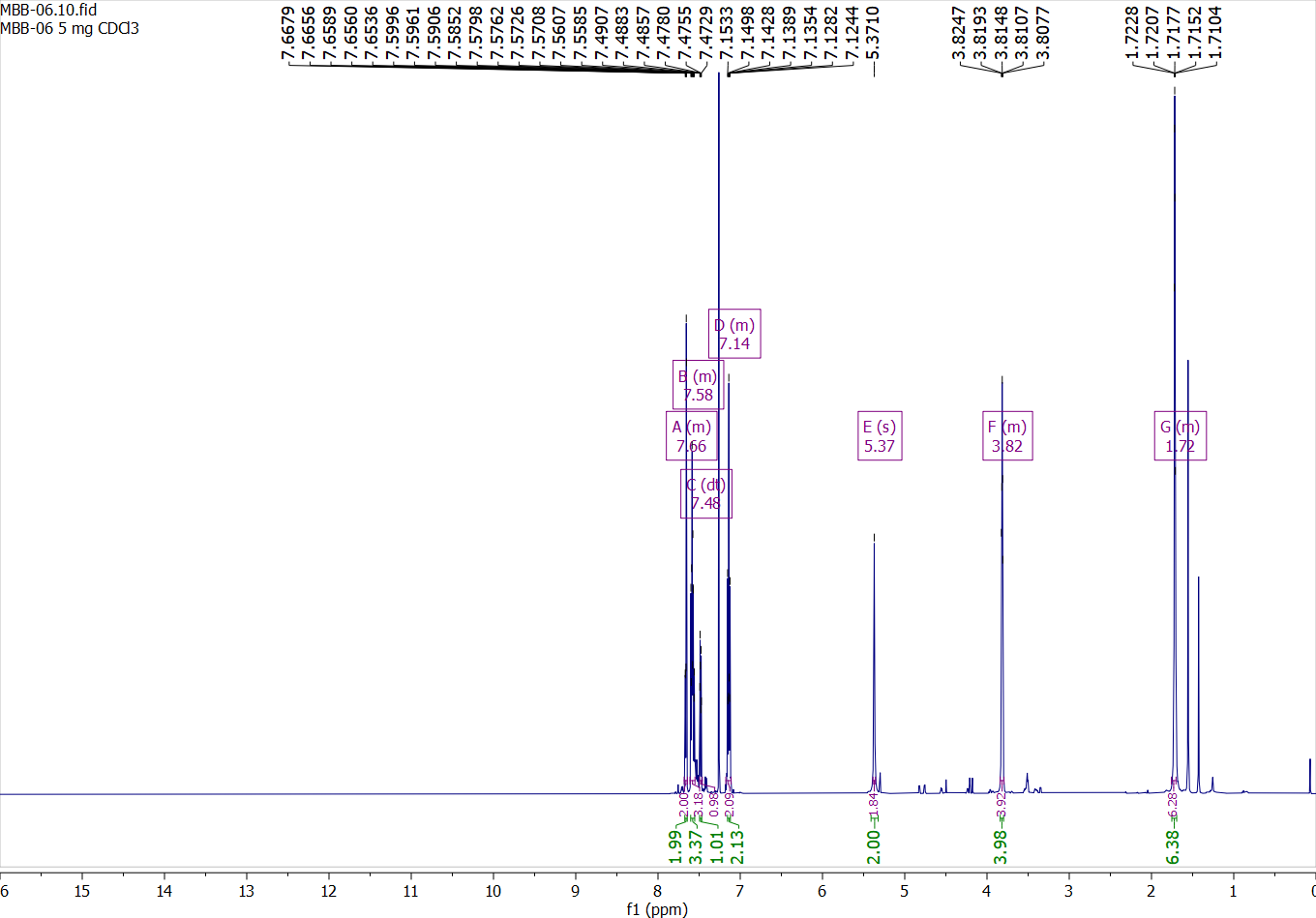

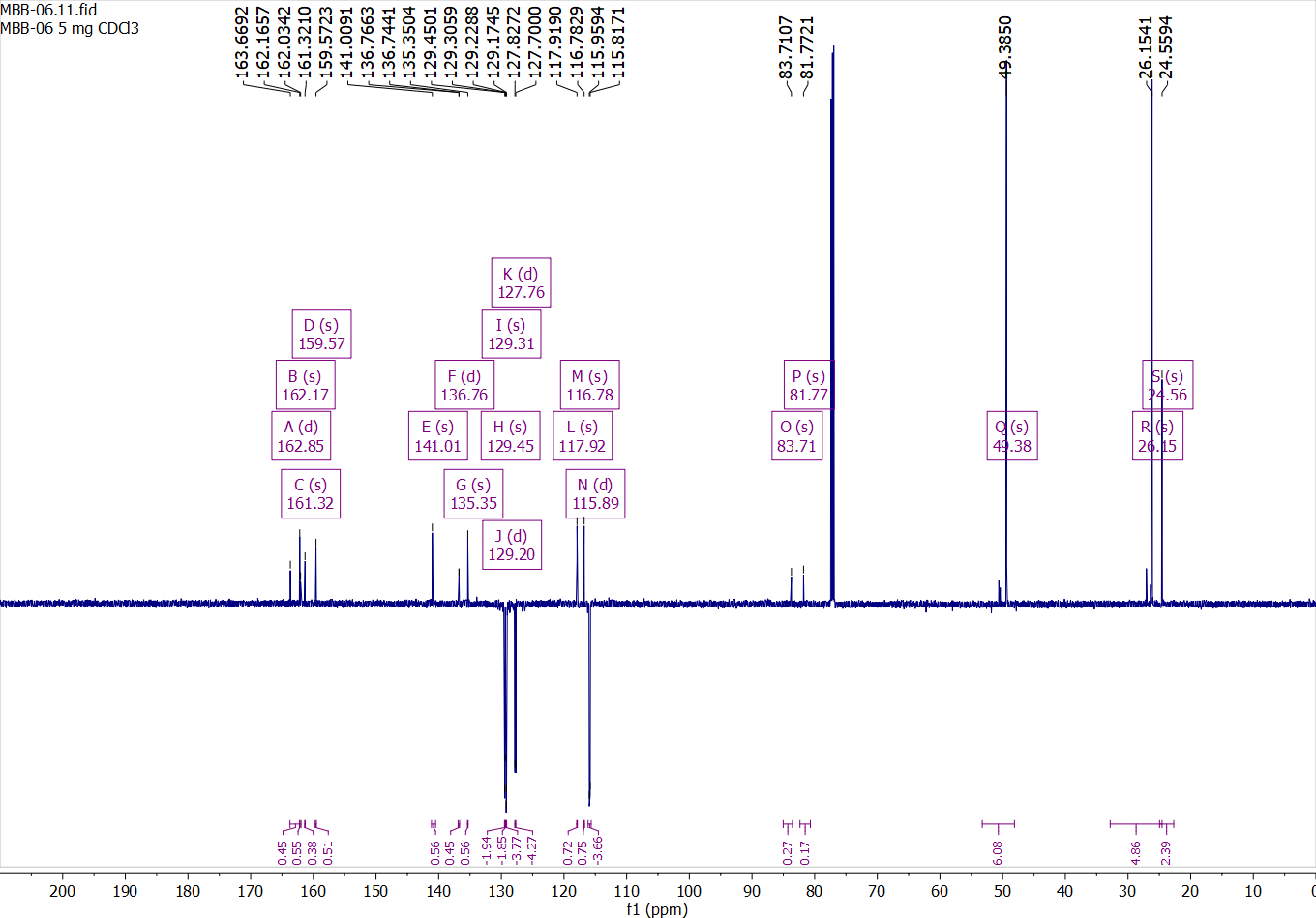

Compound 41:

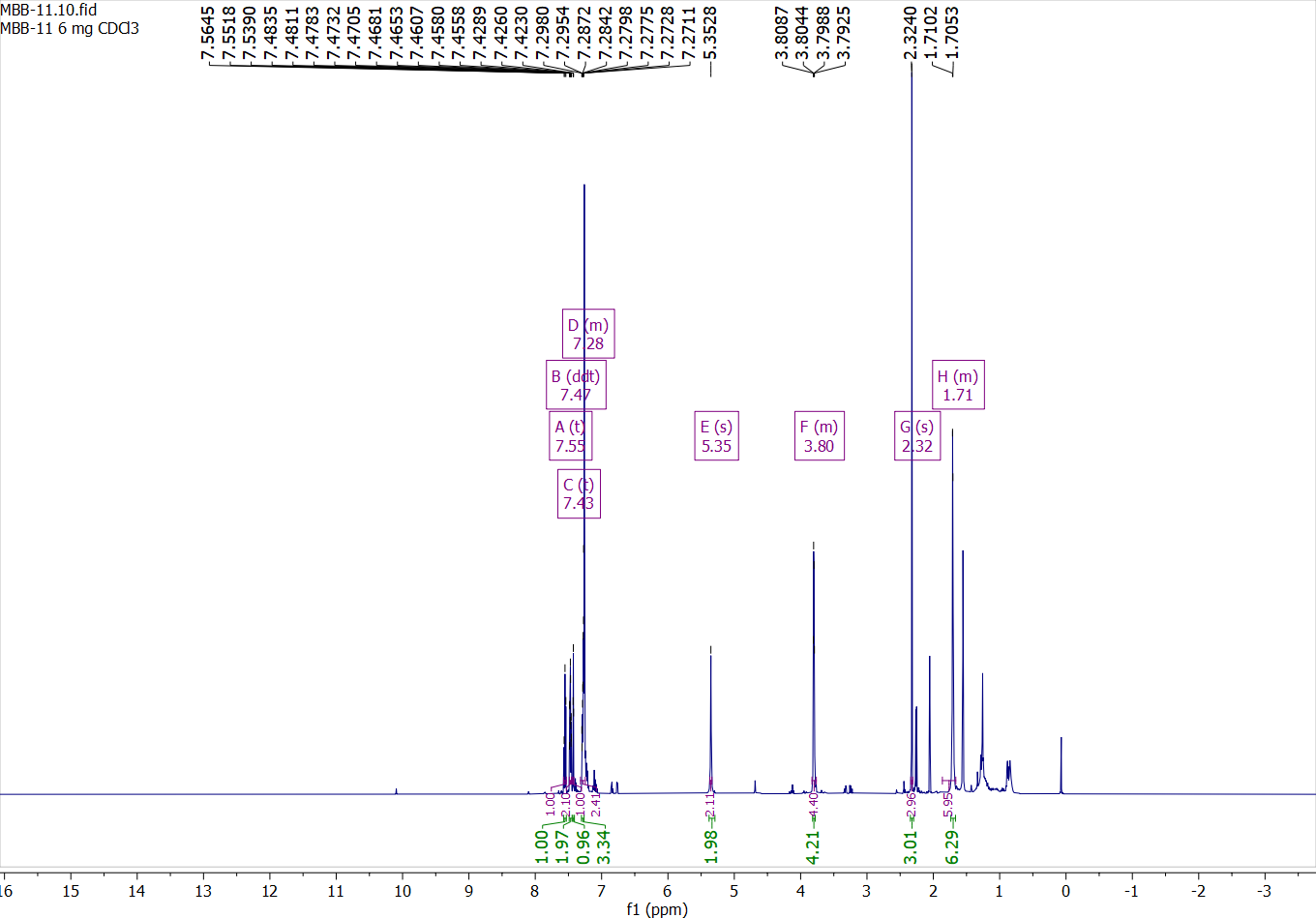

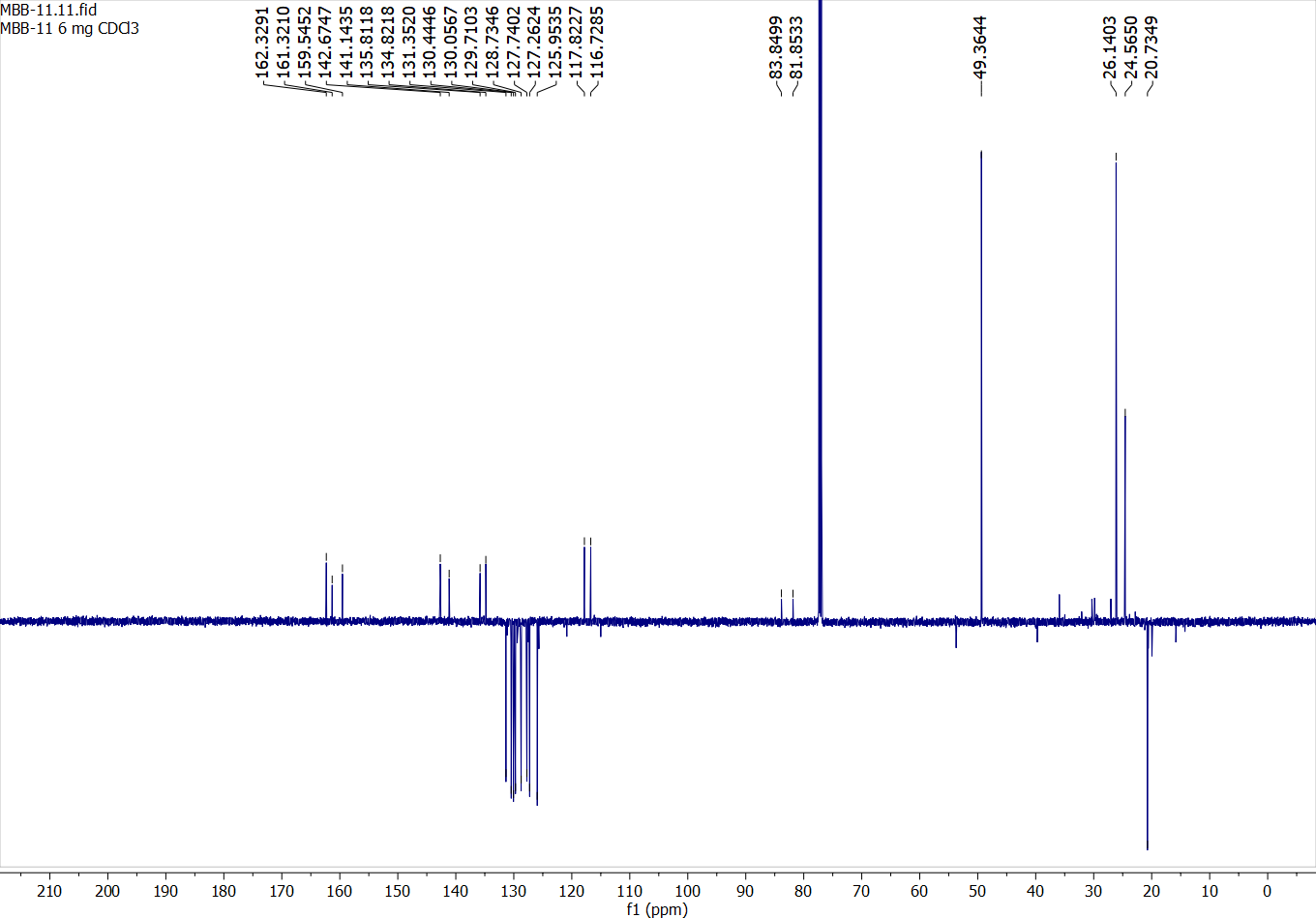

Compound 42:

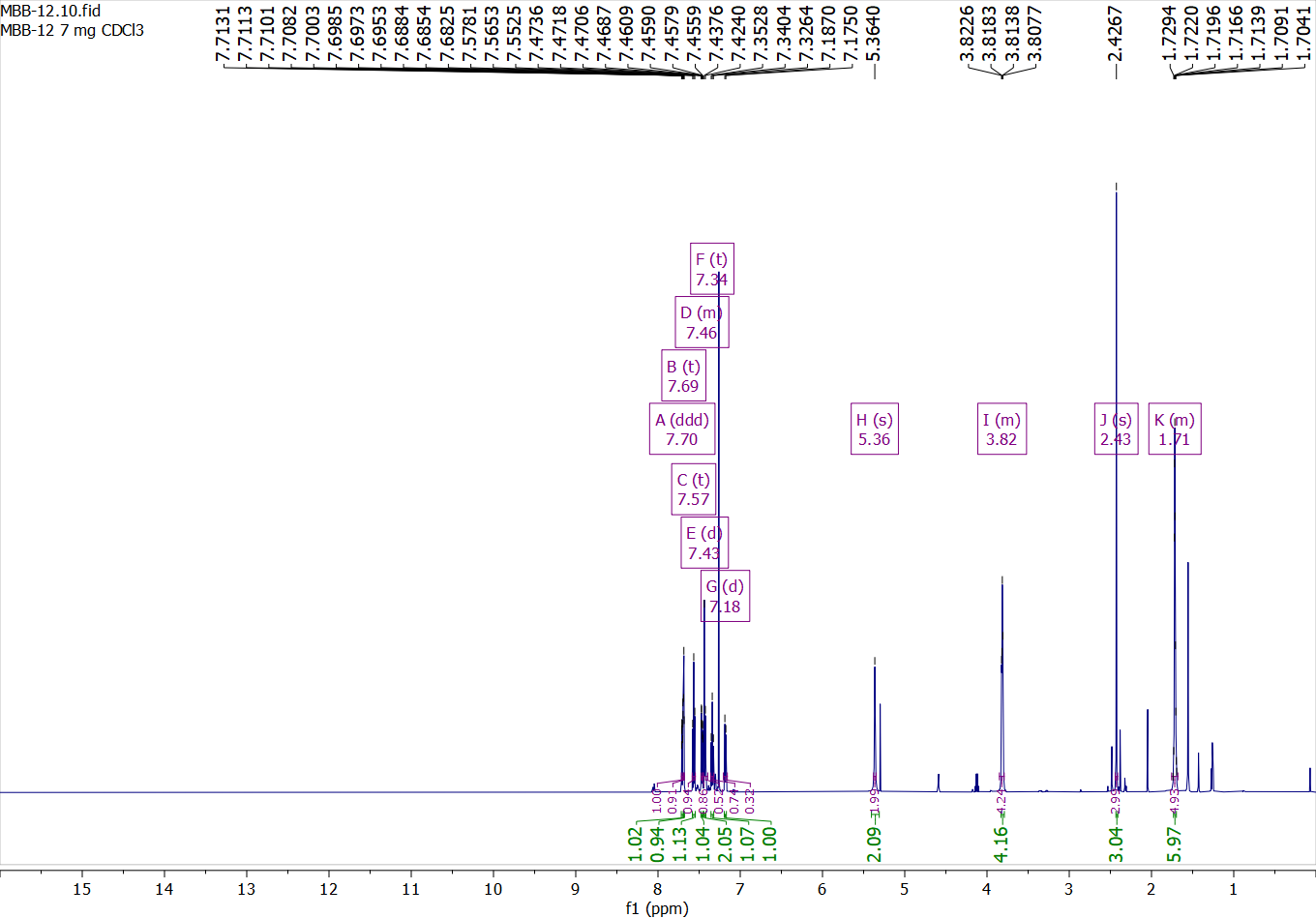

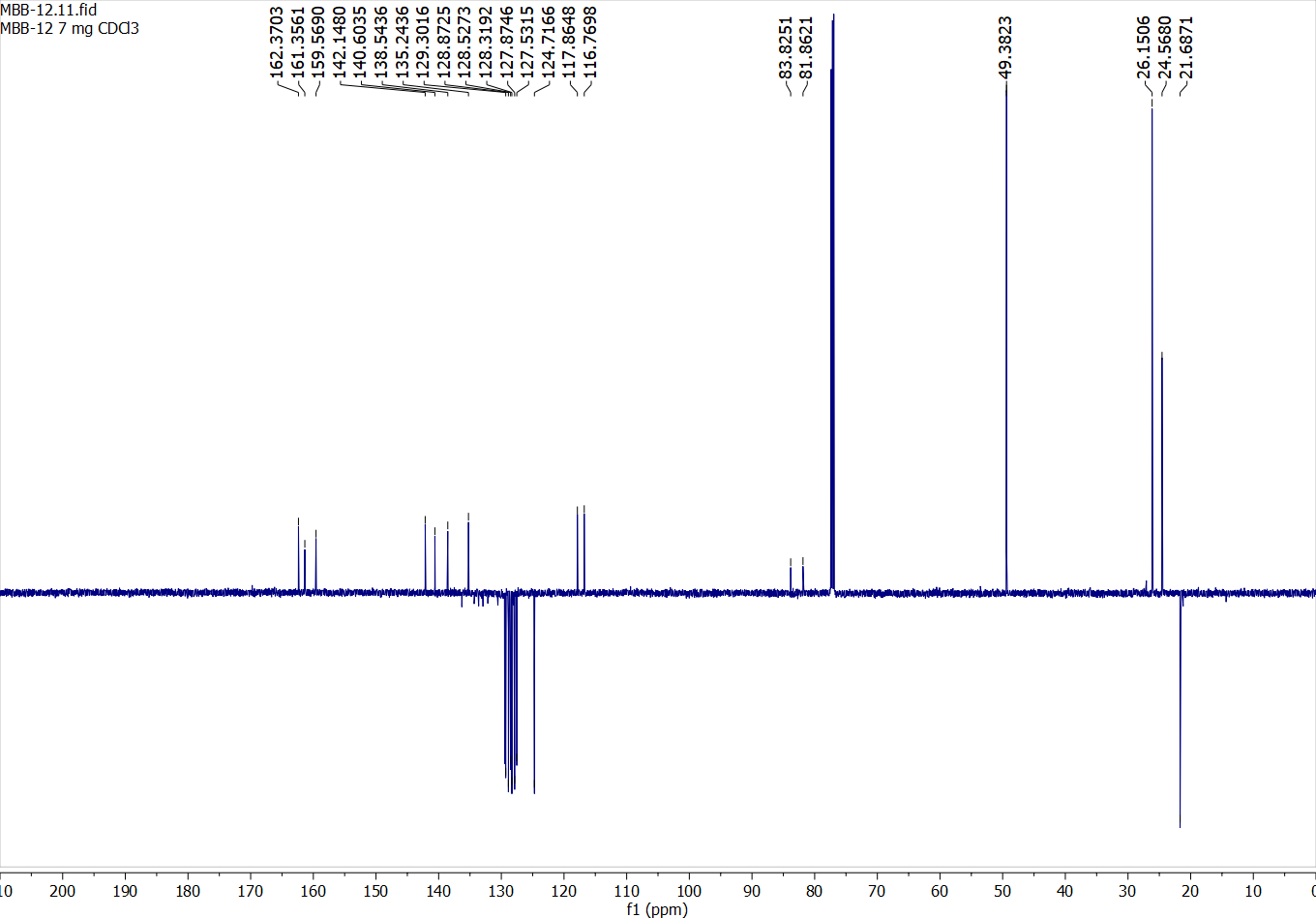

Compound 43:

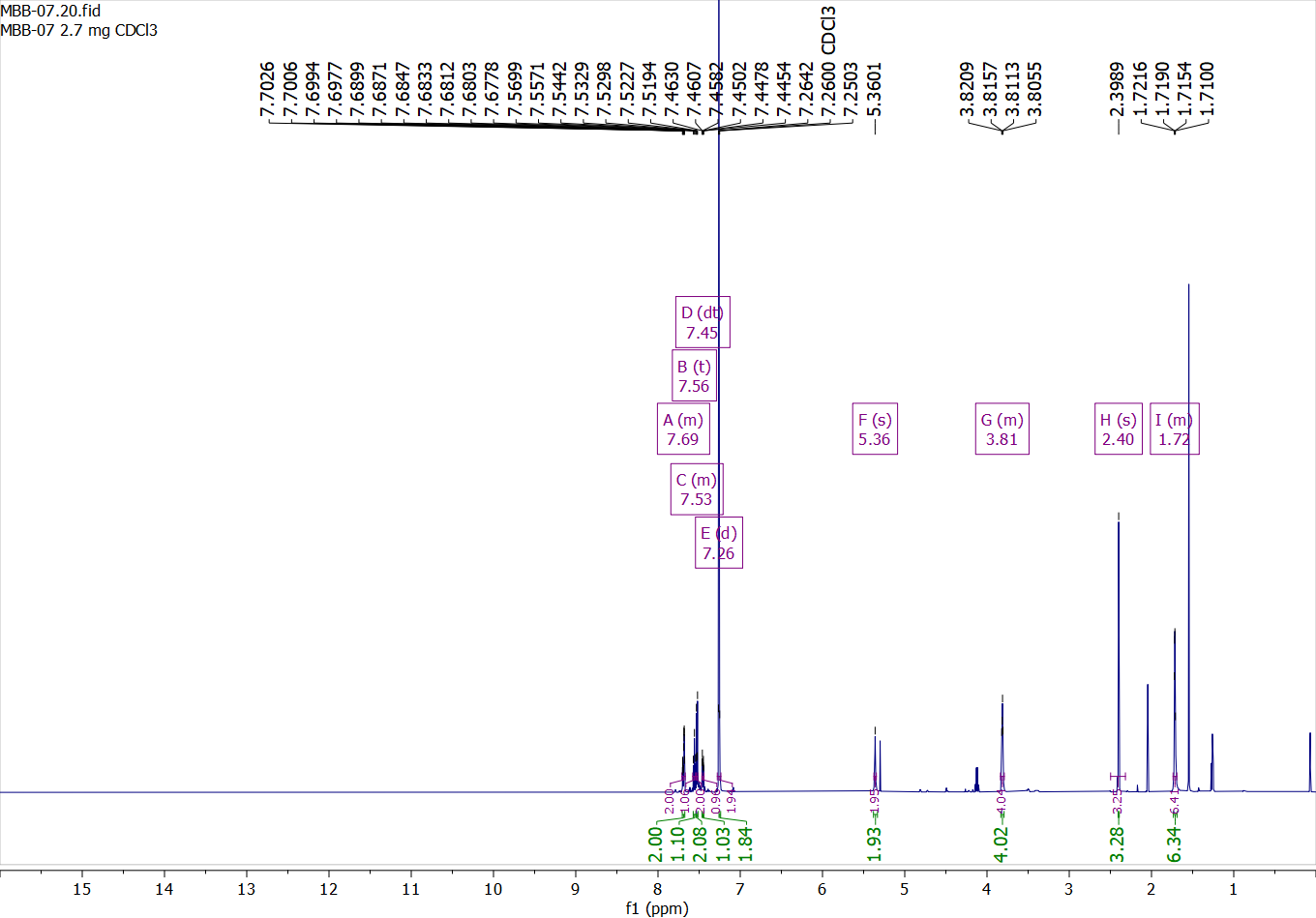

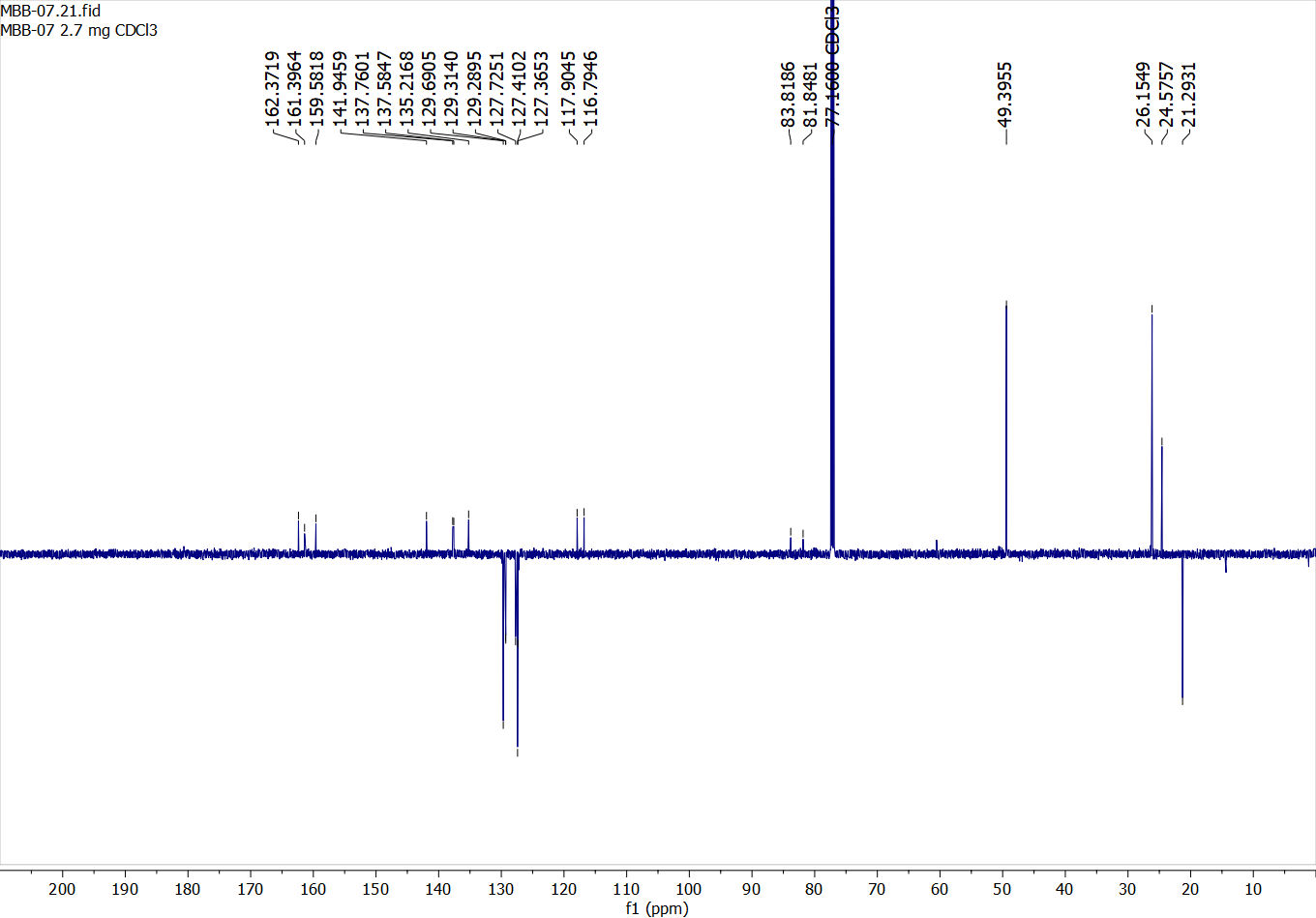

Compound 44:

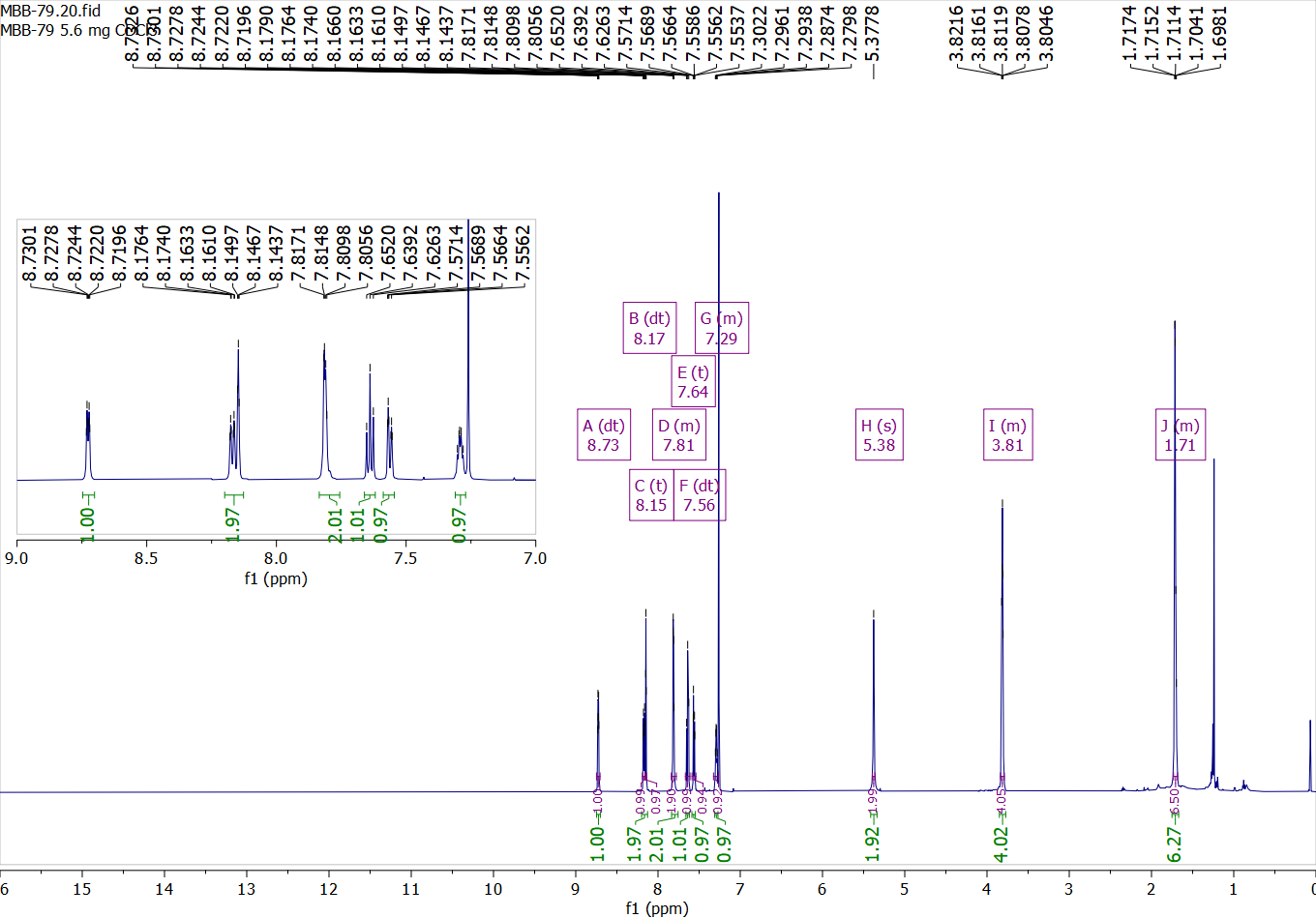

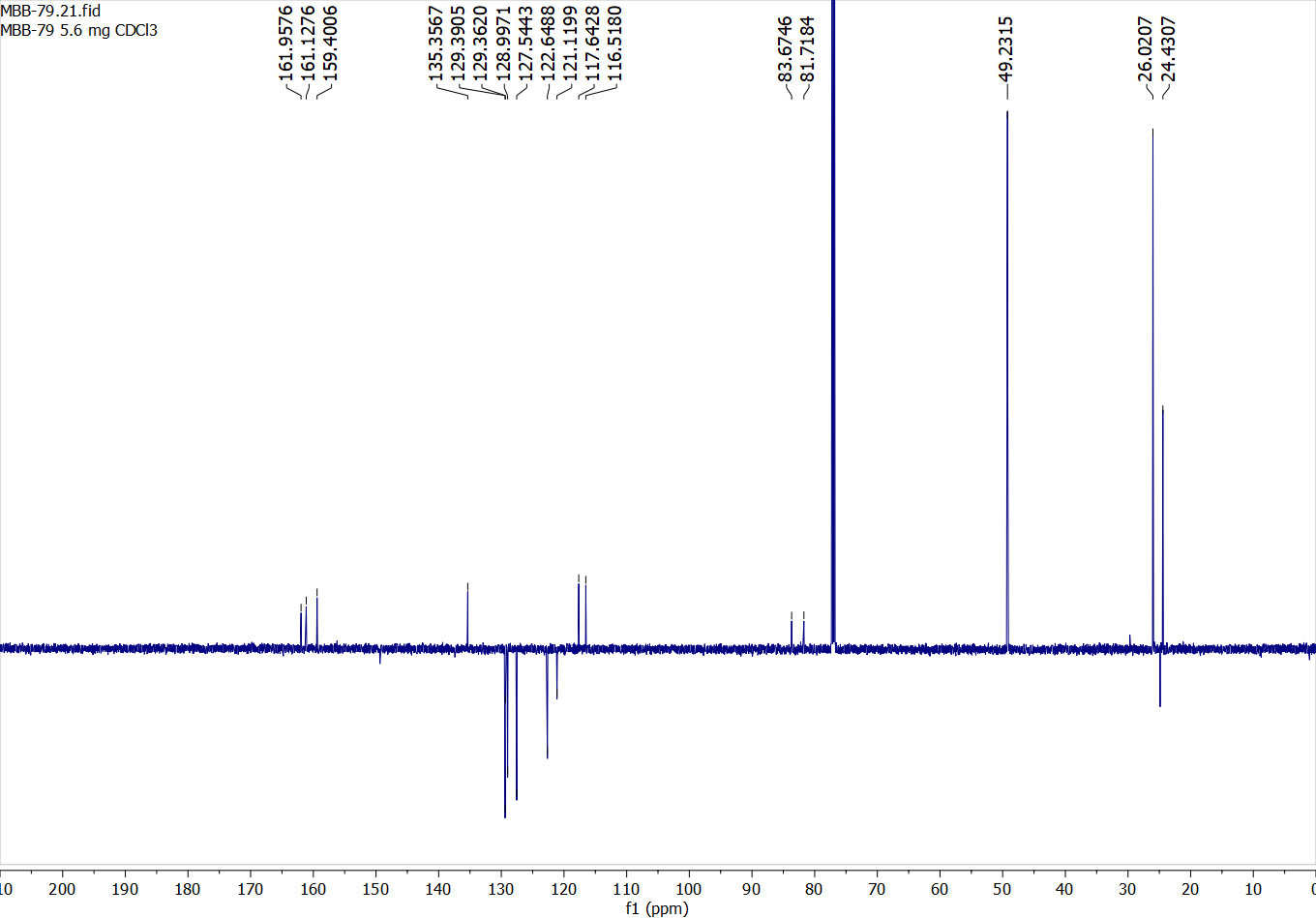

Compound 45:

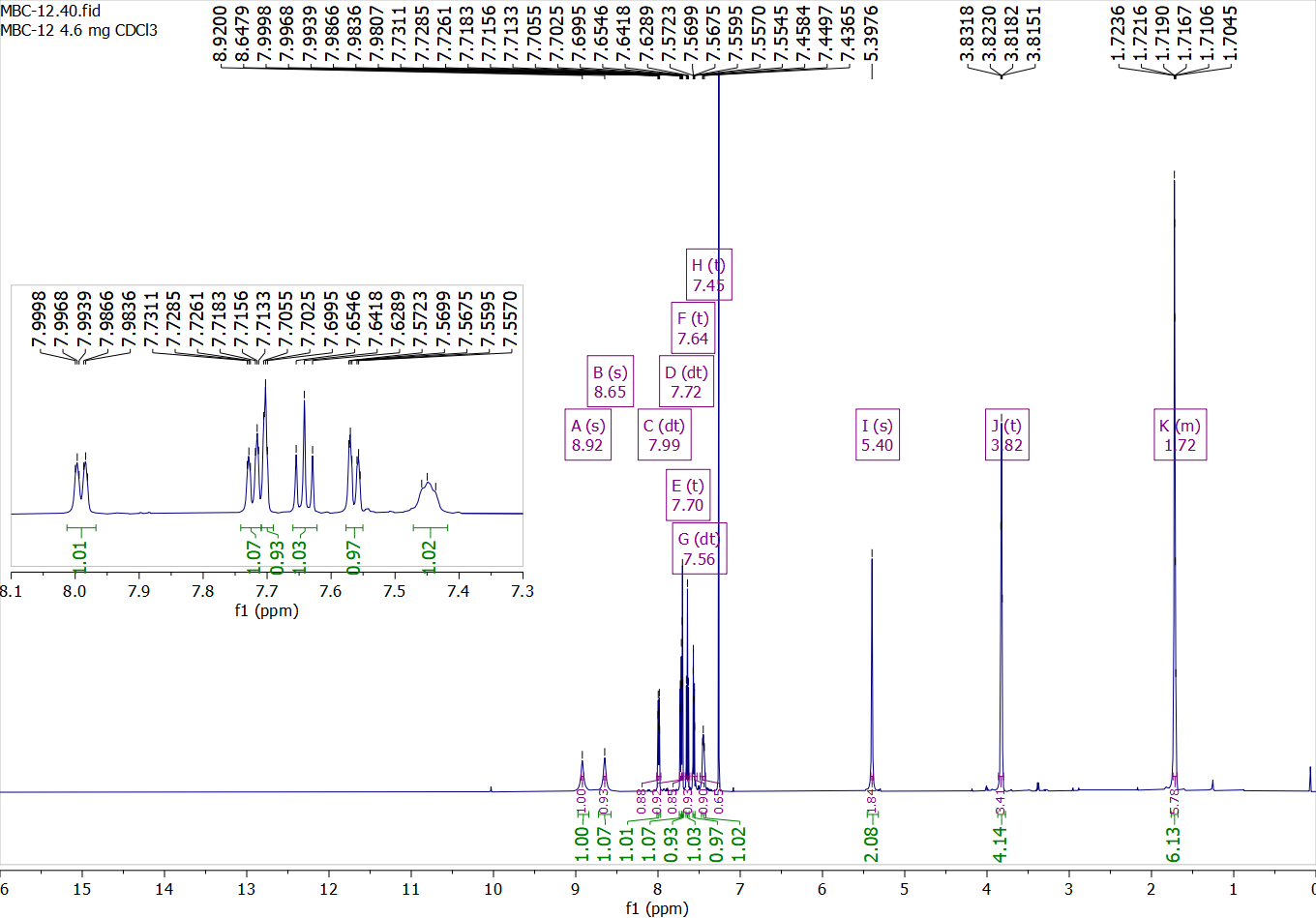

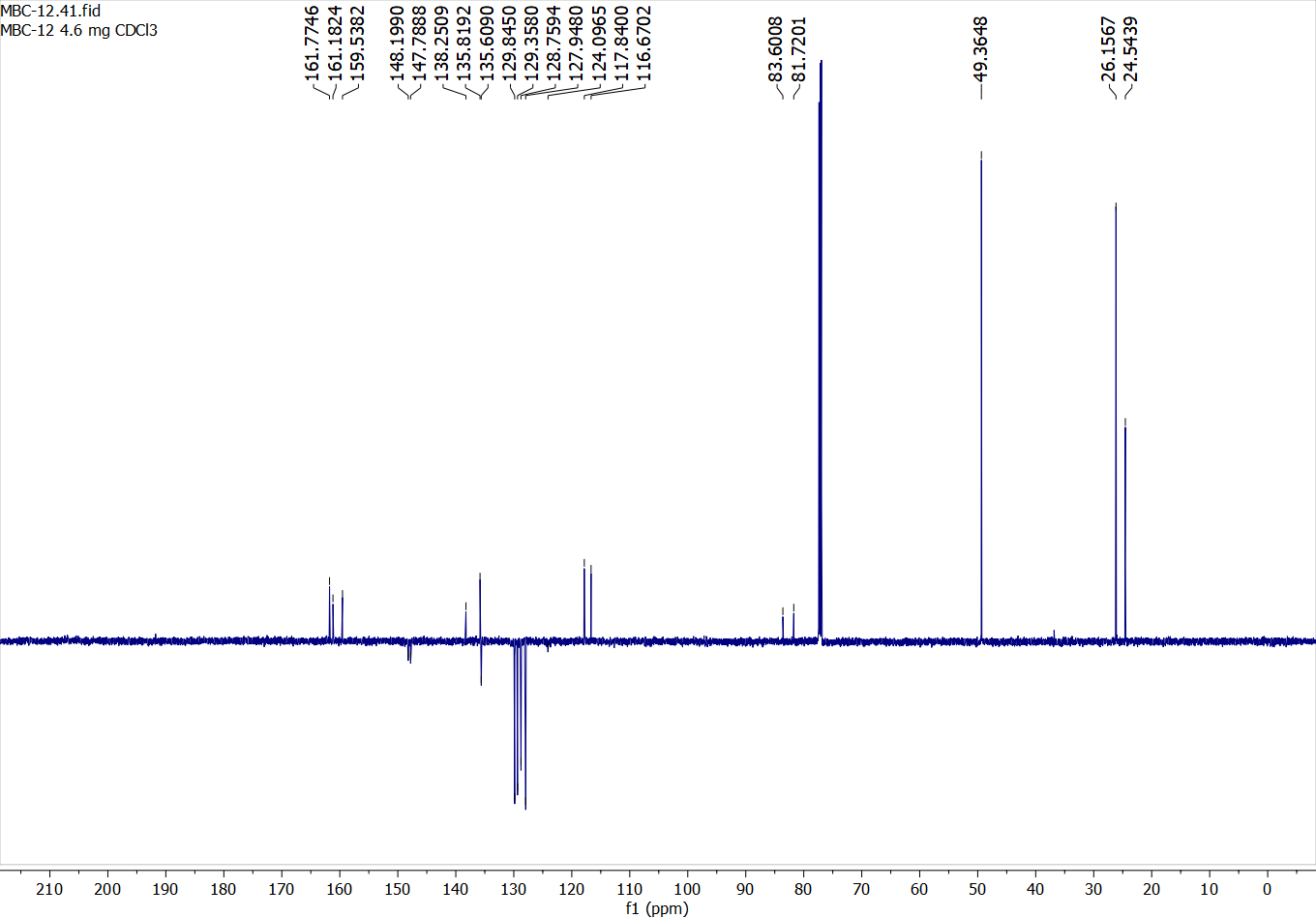

Compound 46:

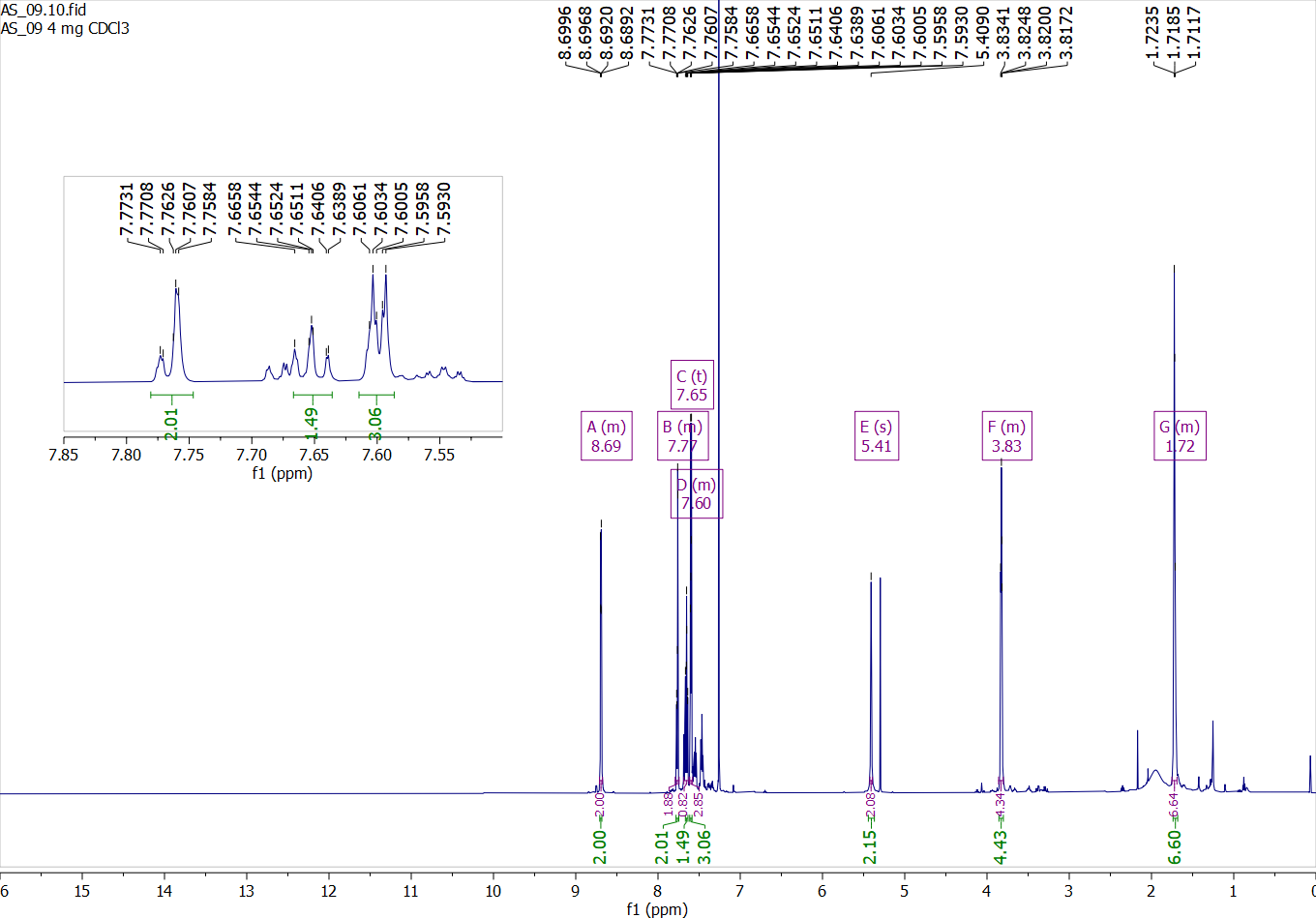

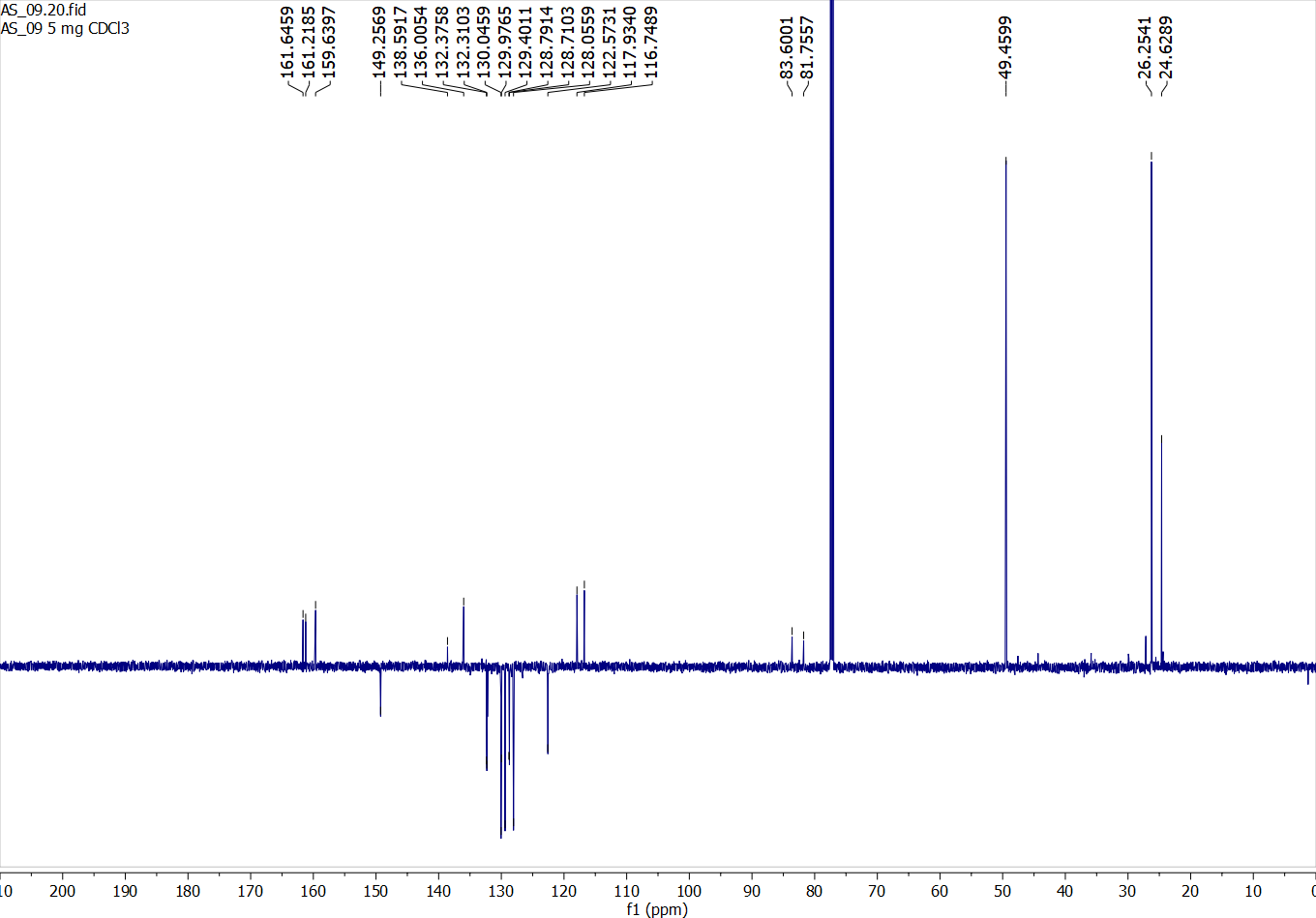

Compound 47:

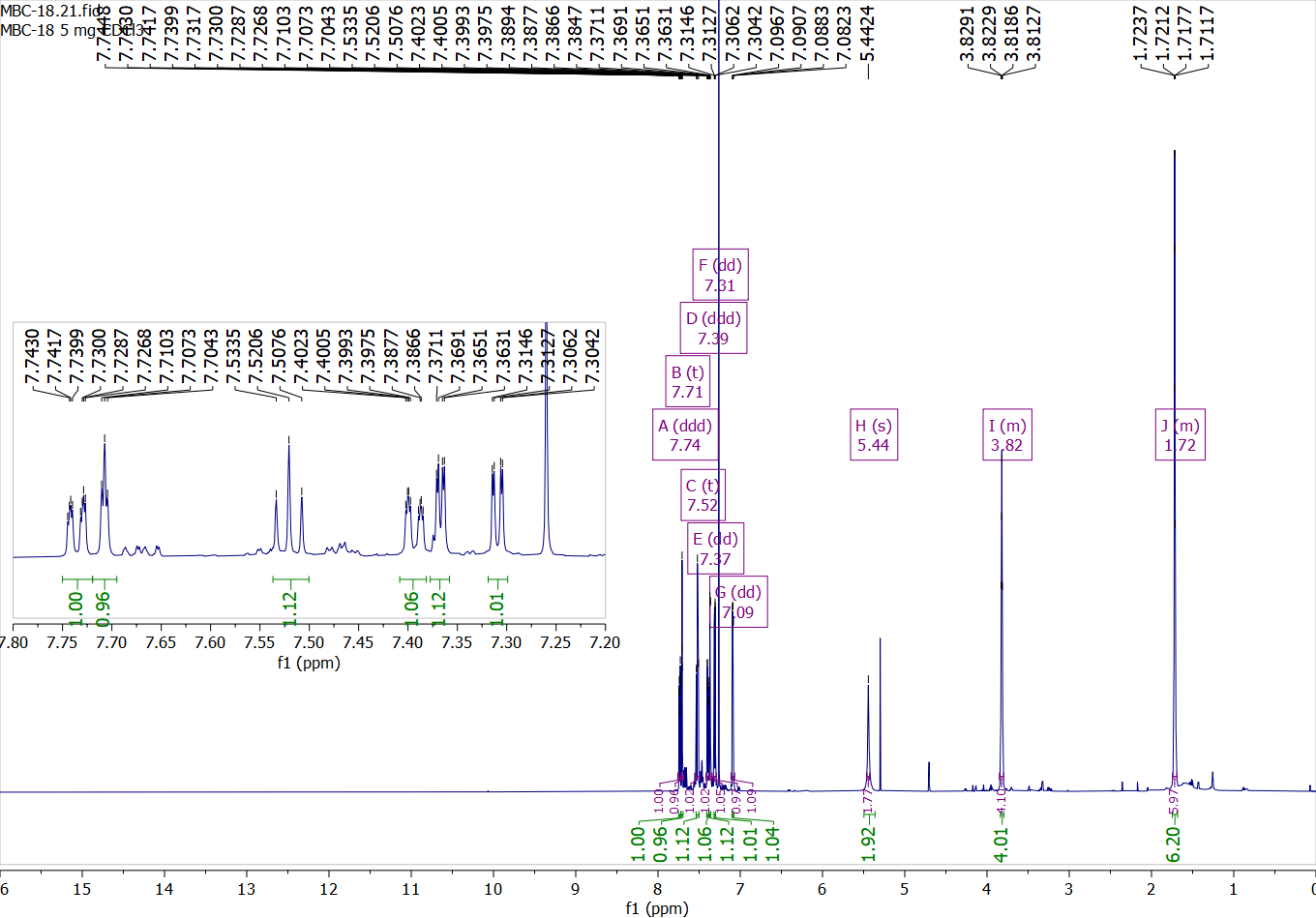

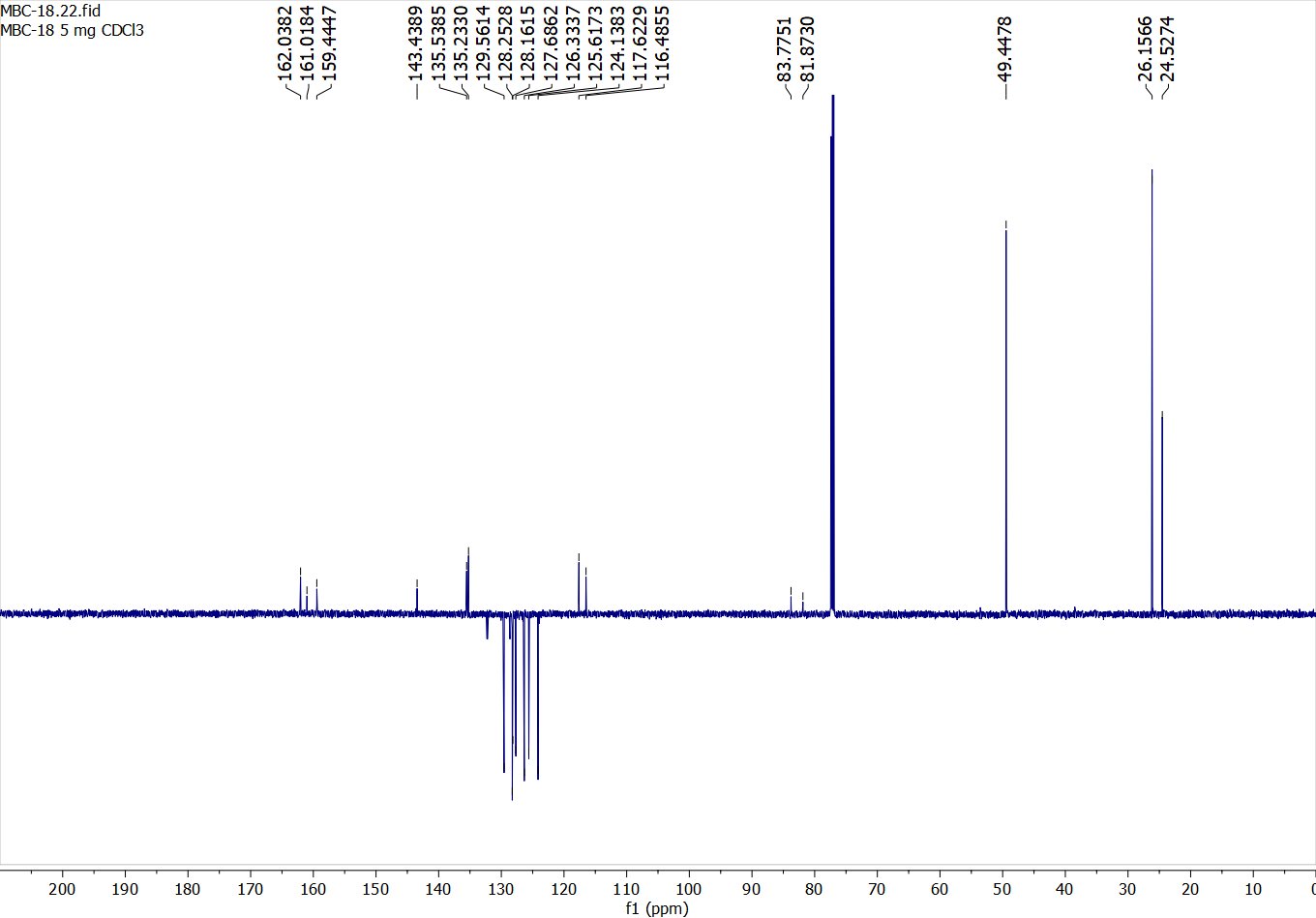

Compound 48:

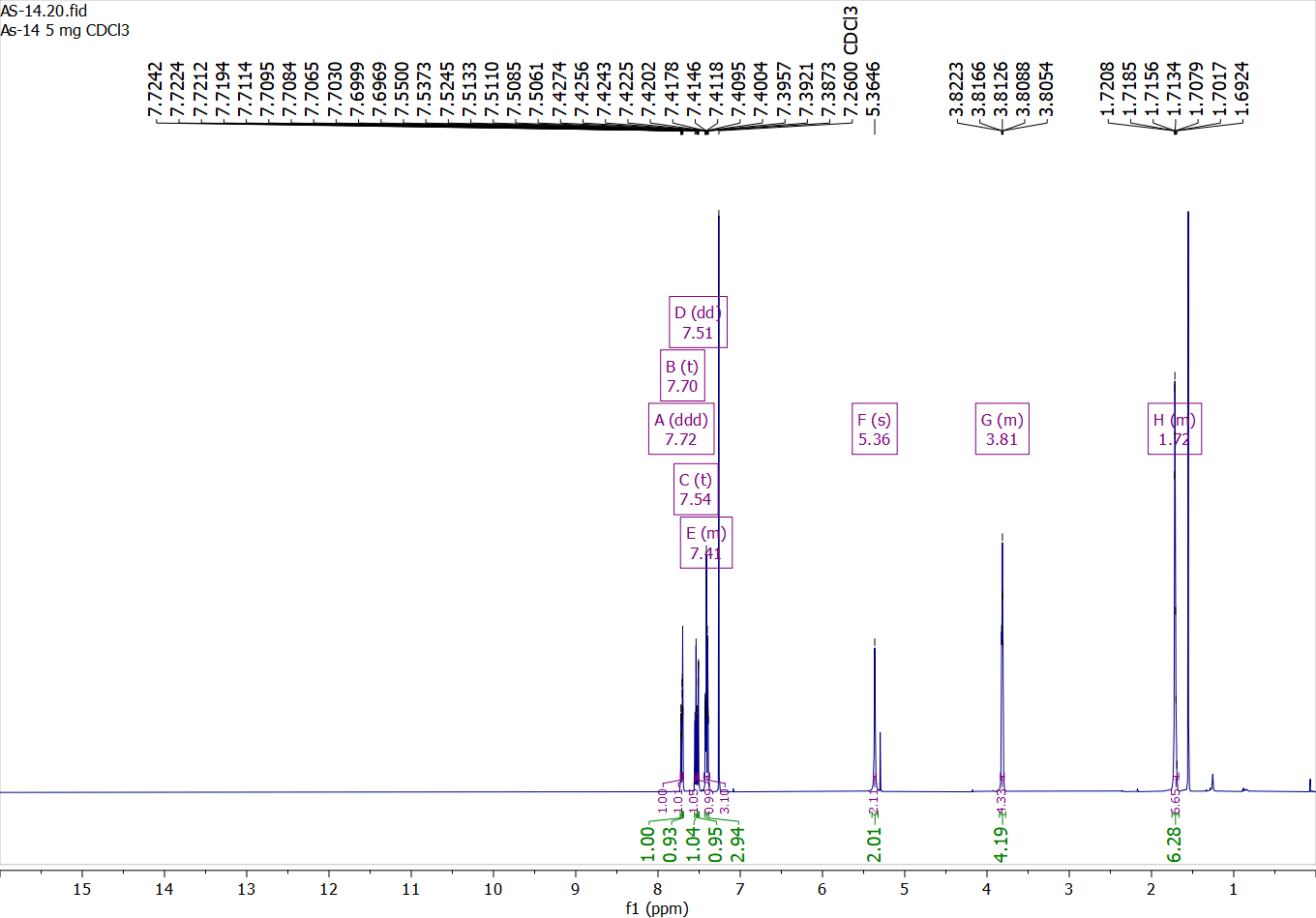

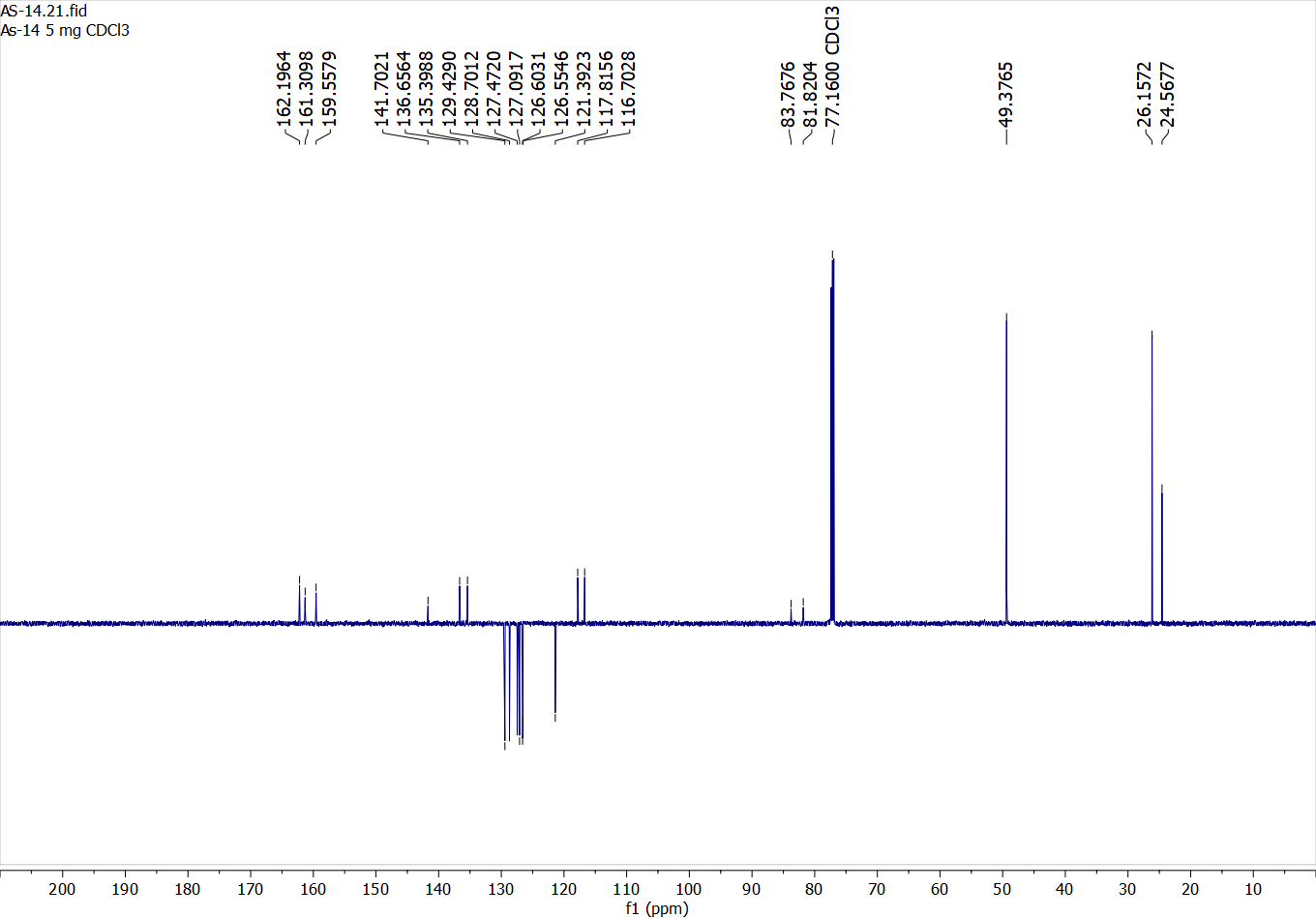

Compound 49:

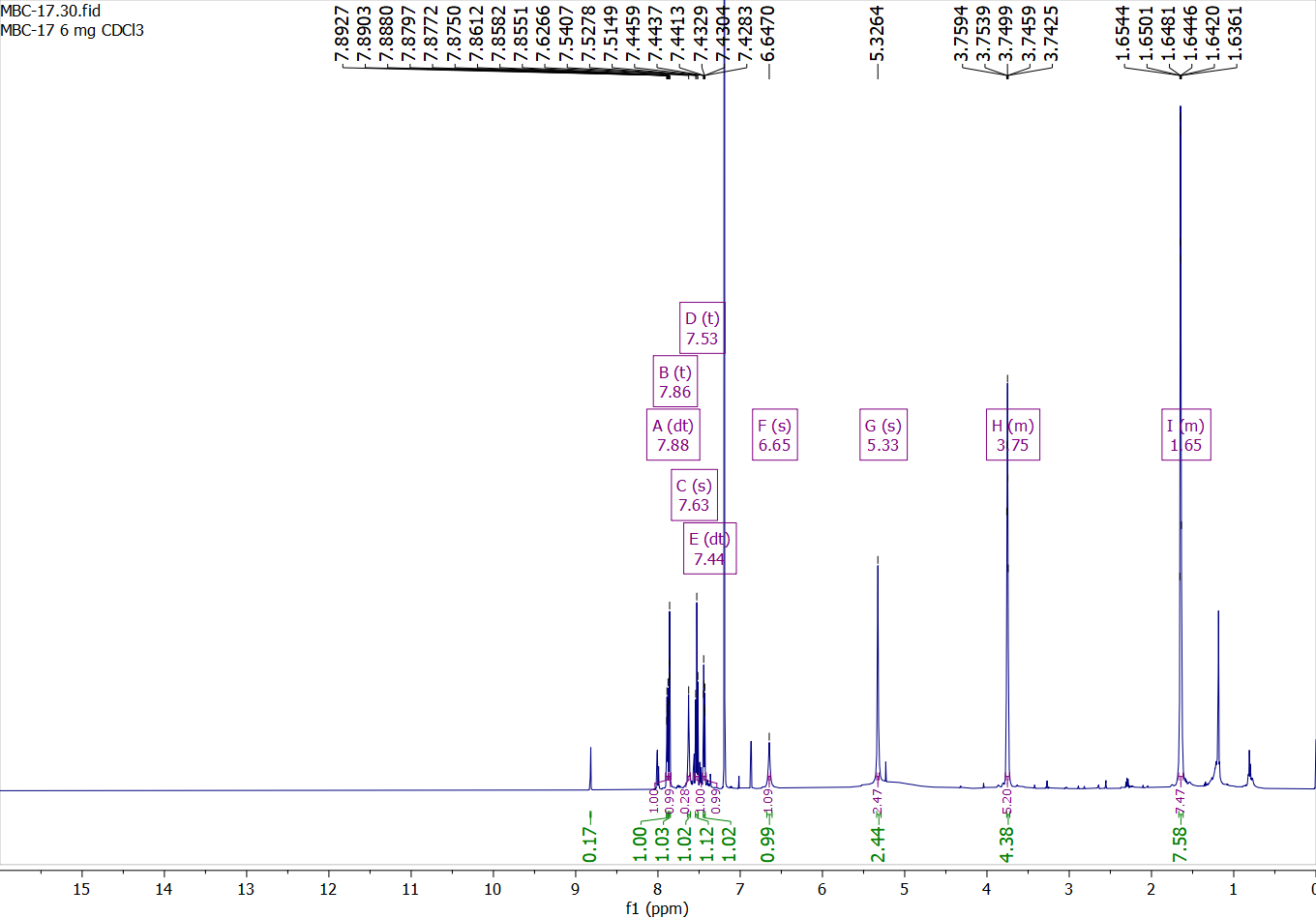

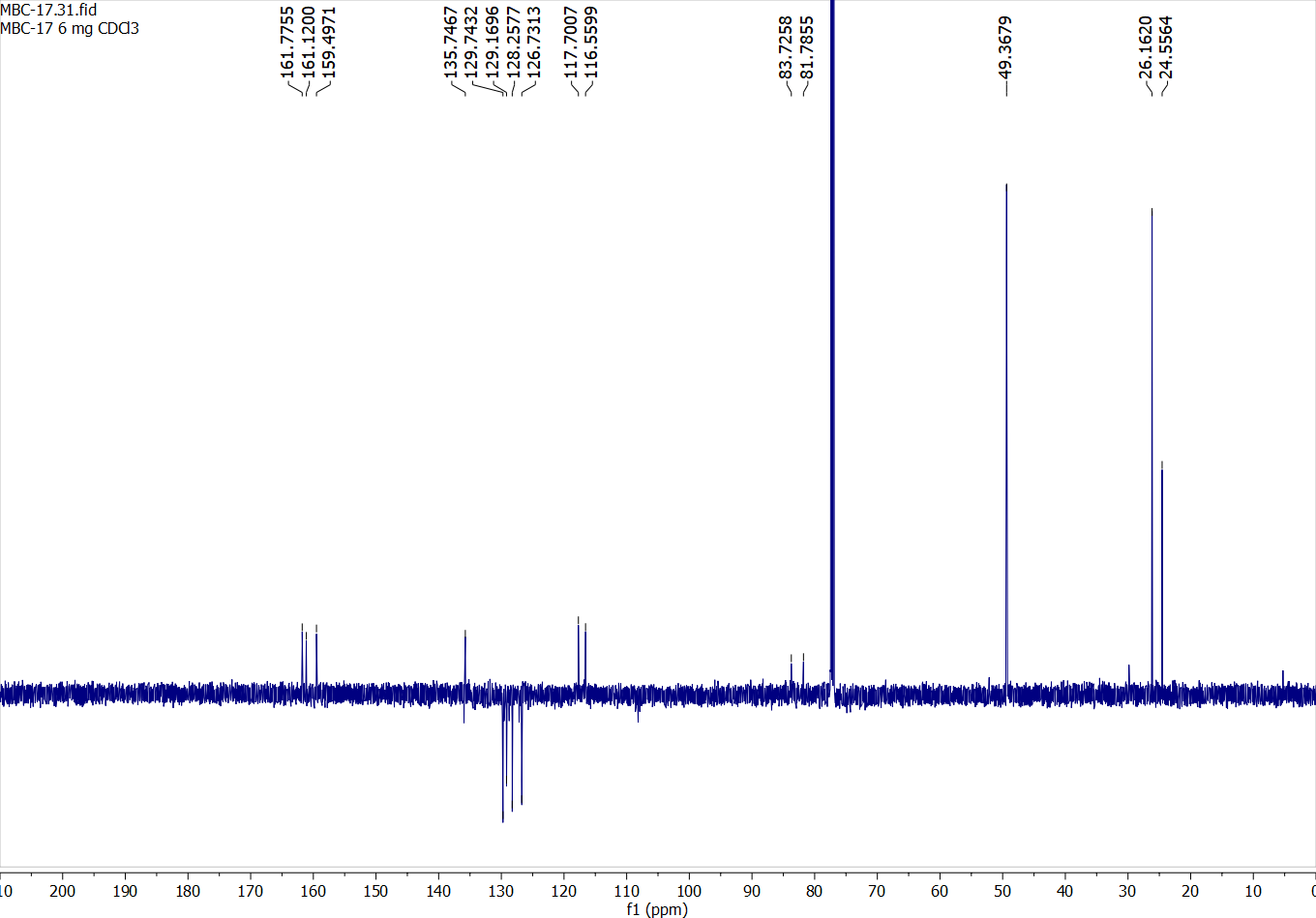

Compound 50:

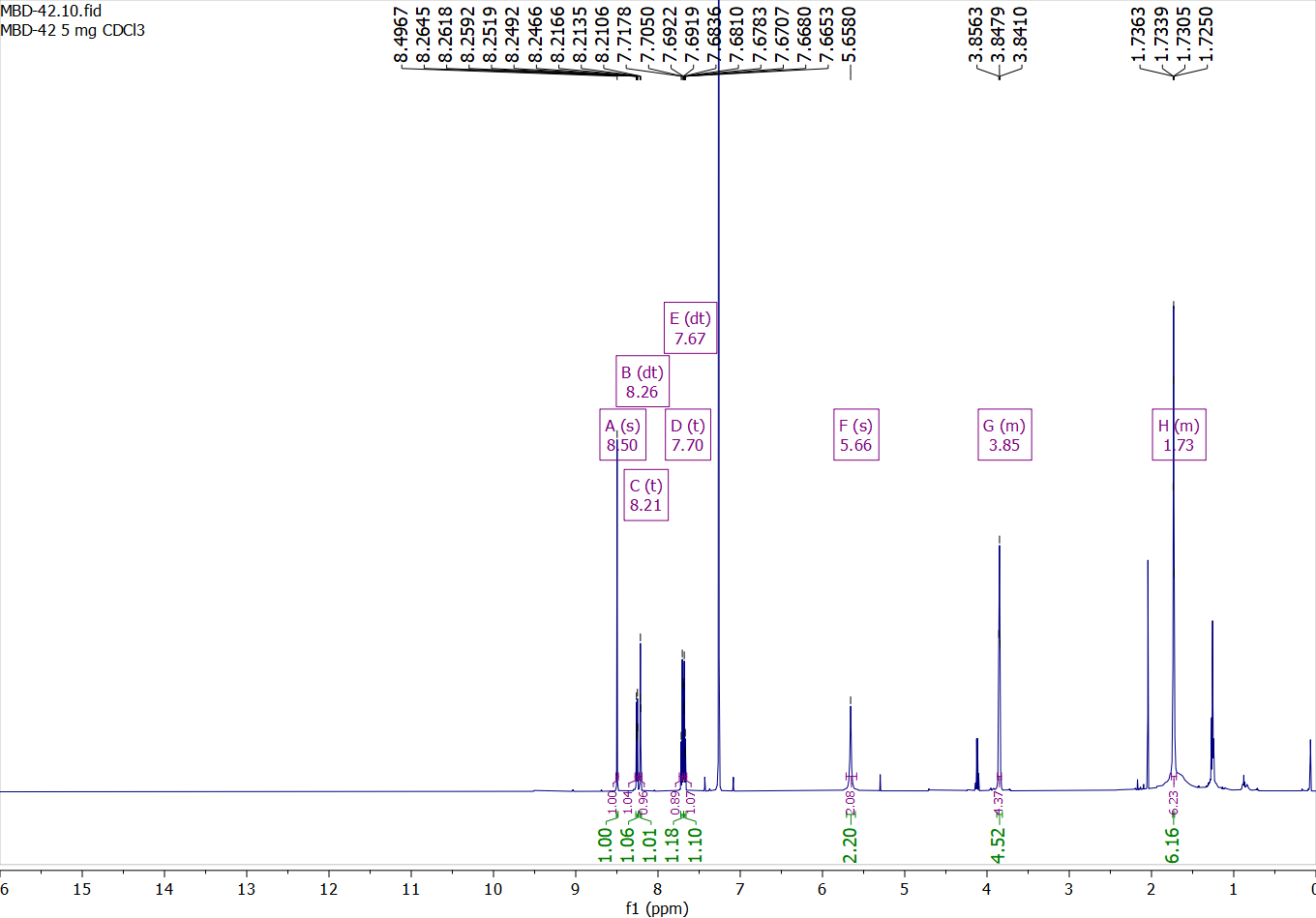

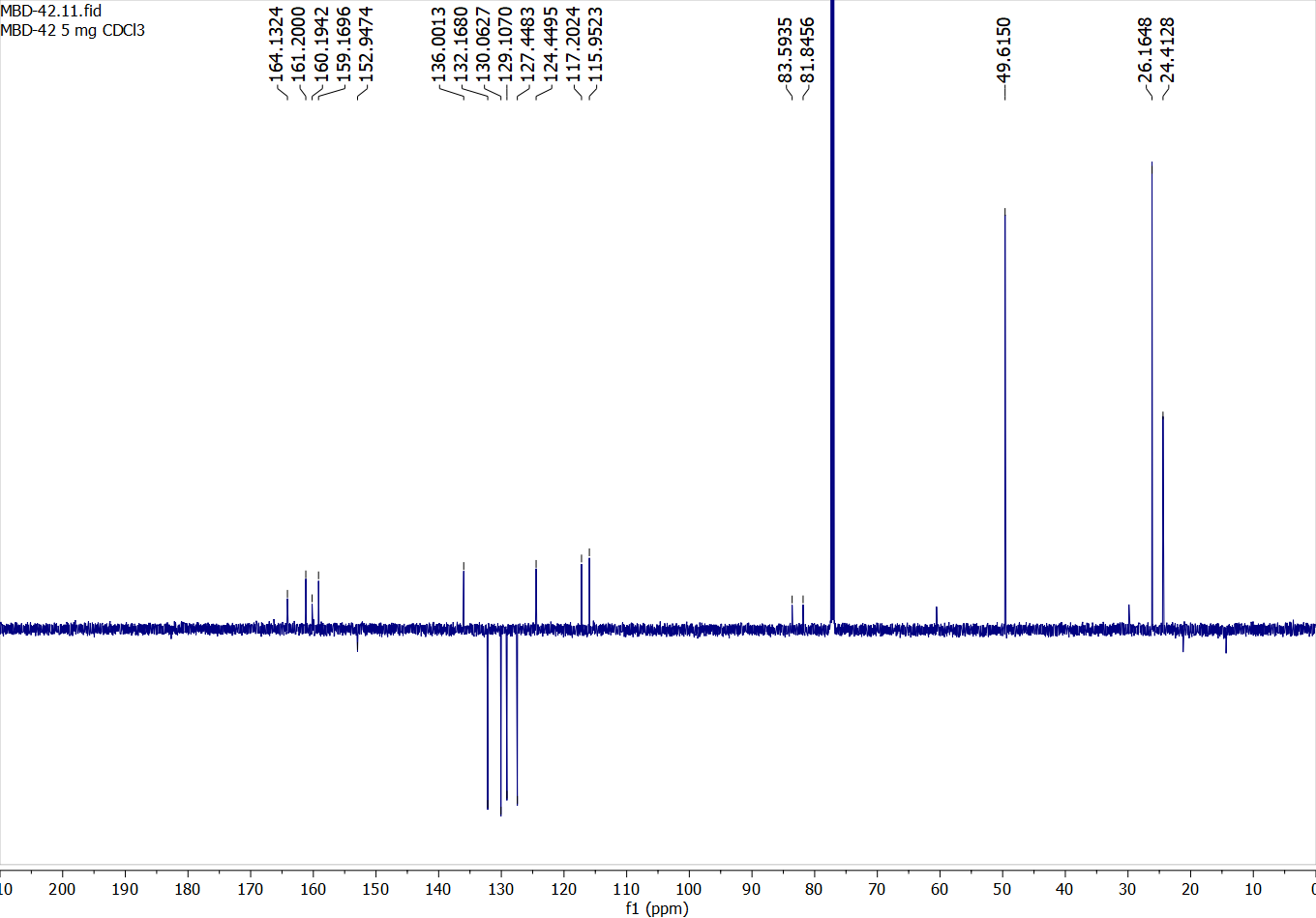

Compound 51:

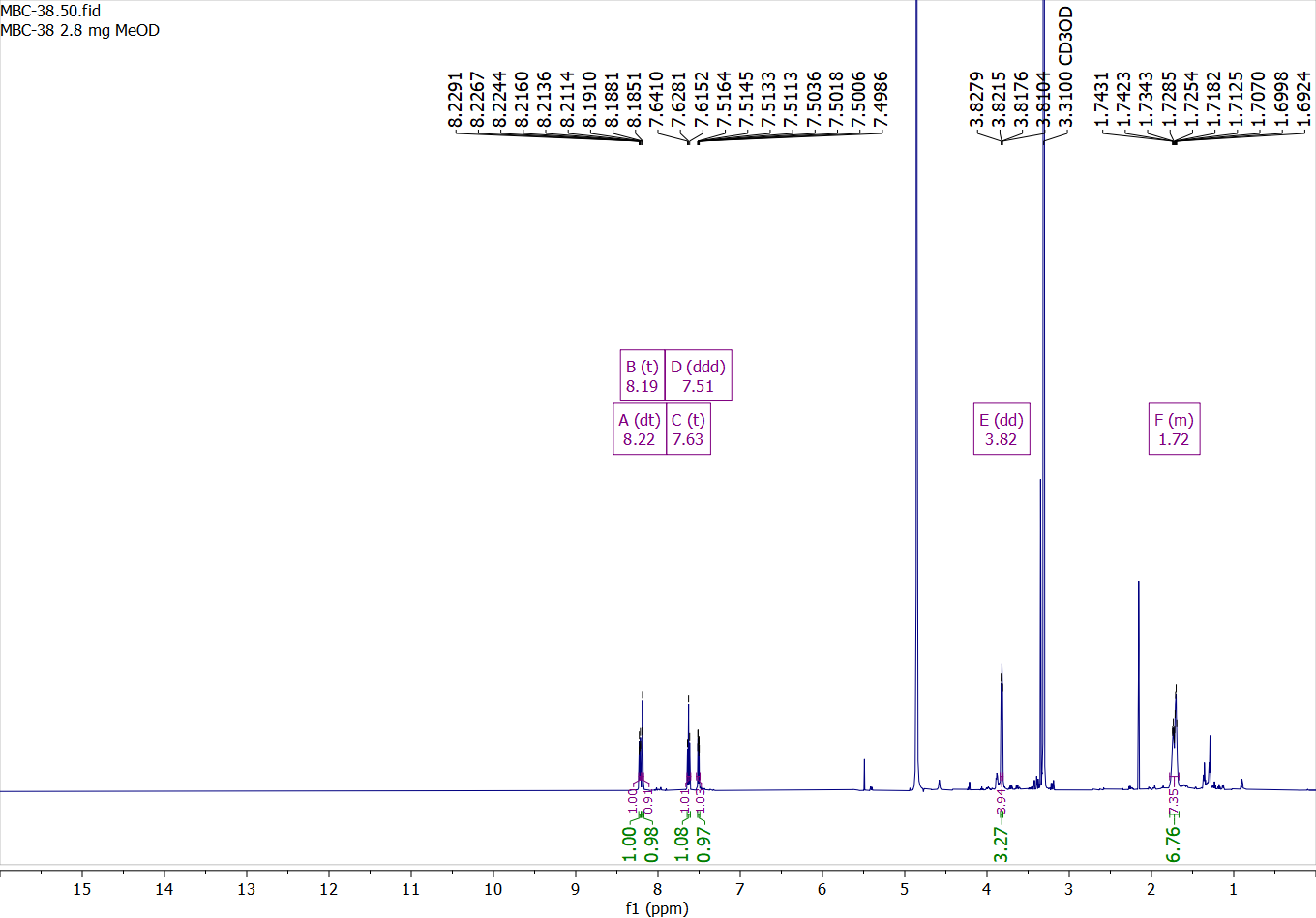

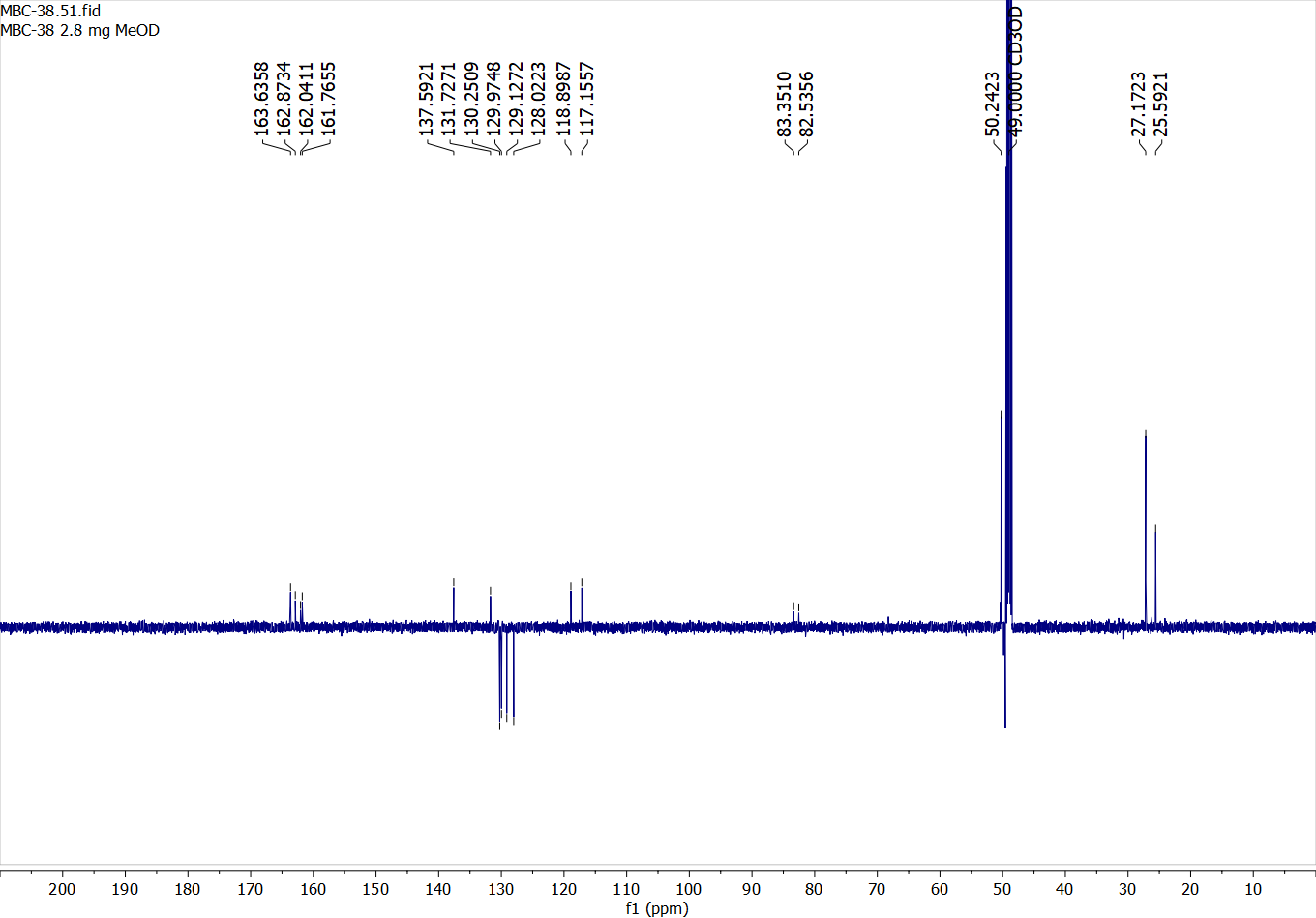

### References

[1]	Z.-G. Gao, A. P. Ijzerman, *Biochemical Pharmacology* **2000**, *60*, 669–676.

[2]	Y. Lu, H. Liu, D. Yang, L. Zhong, Y. Xin, S. Zhao, M.-W. Wang, Q. Zhou, W. Shui, *ACS Chem. Biol.* **2021**, *16*, 991–1002.
